# Antibody peptide epitope conjugates for αPD-1 therapy-resistant head and neck squamous cell carcinoma

**DOI:** 10.64898/2025.11.28.691238

**Authors:** Yifang I. Shui, David G. Millar, Lukas M. Altenburger, Taylor Chrisikos, Michael J. Kalbfleisch, Aaron Hata, Peter M. Sadow, William C. Faquin, Amanda J. Martinot, Christopher M. Snyder, Mikael J. Pittet, Sara I. Pai, Mark Cobbold, Thorsten R. Mempel

**Author notes:** Correspondence to: Thorsten R. Mempel, MD PhD, Center for Immunology and Inflammatory Diseases Massachusetts General Hospital, 55 Fruit St, Boston, MA 02467. These authors contributed equally (Yifang I. Shui, David G. Millar).

## Abstract

Poor tumor antigenicity is an important cause of non-response to immune checkpoint blockade (ICB)-therapy in many cancer patients and mandates new treatment strategies. Here we explore the use of proteolytically activated antibody-peptide epitope conjugates (APECs) to redirect the immunological effector activities of tumor-infiltrating antiviral bystander CTLs against head and neck squamous cell carcinoma (HNSCC) by loading cancer cell surface MHC I proteins with viral peptides. We find that inflationary T cell memory responses against common viral pathogens such as CMV and EBV unfold superior anti-tumor activity compared to conventional memory responses under APEC therapy in an ICB-resistant preclinical model of HNSCC. Mechanistically, APEC activation required cancer cell-intrinsic protease activity, even for proteases expressed by cells of the tumor stroma. Data mining and functional screening identified the protease activity of plasminogen activator urokinase (PLAU) as widely shared between human HNSCC cancers and as a highly efficient proteolytic activator of APECs. Furthermore, peripheral blood analysis in HNSCC patients reliably predicts the specificity, magnitude, and quality of intratumoral bystander CTL, thus allowing for the screening of HNSCC patients who may benefit most from APEC therapy.

## INTRODUCTION

Although immune checkpoint therapy has fundamentally changed the overall survival of patients with recurrent or metastatic (R/M) HNSCC, still only a minor fraction of patients benefit (1). Despite refined treatment regimens, such as adjuvant, neoadjuvant, or combination therapies (2), the low to moderate mutational load and immunogenicity of HNSCC (3) will remain a fundamental limitation to all immunotherapies that build on T cell recognition of tumor neoantigens. This presents a pressing need to develop alternative, neoantigen-independent treatment strategies.

In many cancer types the majority of tumor-infiltrating CD8^+^ T cells are not tumor-antigen-specific but represent non-specifically recruited memory and effector cells reactive to cancer-unrelated antigens, including to viral antigens introduced though prior vaccination (e.g. Flu) or prior or ongoing viral infections such as endemic chronic infections with Cytomegalovirus (CMV) or Epstein-Barr virus (EBV) (4–6). These cells are commonly characterized by high functional potential, in contrast to tumor antigen-reactive CTLs, which are often found in late stages of differentiation and have reduced residual anti-tumor activity (6–8). Recently, new treatment strategies have emerged that seek to therapeutically harness antiviral bystander T cells by redirecting their immunological effector functions against cancer cells (9–15). These strategies share the goal to engineer presentation of viral antigenic epitopes on cancer cell surface MHC I molecules in order to provoke their direct engagement with antiviral bystander CD8^+^ T cells and induce cytotoxic cancer cell killing. The most direct strategy among these is to inject free viral peptides directly into the tumor (9, 14). However, this strategy depends on the accessibility of the tumor lesions for direct injection, which may miss unidentified metastatic lesions and may be limited by off-target toxicities resulting from binding of leaked peptide to healthy tissues. We and others have therefore pursued an alternative strategy, which is to deliver viral peptides conjugated to antibodies targeting cell surface antigens over-expressed on the surface of cancer cells, using linkers that contains motifs sensitive to cancer cell-expressed proteases (10, 11, 15, 15). The activity of these APECs is predicted to be highly specific to tumor tissue through several mechanisms: 1) the targeting of the antibody to antigen(s) over-expressed on cancer cells; 2) selective cancer cell protease activity; and 3) increased density of antiviral CTLs in the tumor microenvironment (TME) relative to healthy tissues.

Preclinical studies in immune-deficient mouse and zebrafish xenograft models, where APEC target antigens are only expressed on cancer cells, have provided proof-of-concept that APECs can unfold anti-tumor activity *in vivo* (10–13, 15). However, important questions remain to be answered before first-in-human studies can be considered: How efficacious are APECs in immune-competent hosts? Here, redirected antiviral CTL activity may synergize with the effector activity of tumor-reactive CTLs. On the other hand, antiviral CTLs may also be more effectively controlled by T regulatory cells. Also, does expression of APEC target antigens on healthy tissue cause off-tumor on-target toxicity? Furthermore, which functional states of antiviral CTLs accumulate in the TME, and which show the most pronounced and sustained anti-tumor activity under APEC therapy? Does APEC activity depend on cancer cell protease activity, or can they also be cleaved by immune and stromal cell-expressed proteases? The latter could promote off-tumor activity and toxicity in inflamed healthy tissues, especially where APEC target antigens are expressed. Are some protease activities shared by a wide range of human cancer types or at least by the majority of patients with a specific cancer type? Finally, can the presence and abundance of specific antiviral CTL responses in the TME be predicted in the individual patient? The latter questions are of high practical relevance for drug development and patient selection.

Here, we seek to address these questions in the context of HNSCC. In paired analyses of CMV-, EBV-, and Flu-specific CTL in tumor tissue and blood samples from human patients we find that antiviral responses in peripheral blood are a reliable predictor of the same responses in tumor tissue, including their magnitude and composition of memory and effector-like states. Through a multistage screening approach, we furthermore identify a peptide motif cleaved by the protease PLAU to be suitable for incorporation in APECs that can be proteolytically activated with high efficiency by a panel of both HPV^−^ and HPV^+^ human HNSCC lines. Finally, we adopted an immune-competent and immune checkpoint therapy-resistant mouse model of HPV^+^ epithelial cancer to demonstrate that APECs redirecting *in vivo* primed anti-CMV CTL produce sustained tumor control, that redirected inflationary are more effective than conventional antiviral memory responses, that APEC activation requires cancer cell protease activity, and that EpCAM-targeted APECs do not produce clinical or histological signs of toxicity in EpCAM-expressing healthy tissues.

## RESULTS

### Peripheral blood analysis predicts antiviral CTL responses in the HNSCC TME

Identifying cancer patients who are candidates for APEC therapy will require knowledge of the antigen specificity and abundance of antiviral bystander CTLs in their tumor tissue. In order to assess whether antiviral CTL responses in blood predict the presence, differentiation state, and abundance of the same responses in tumors, we recruited a cohort of 43 HLA-A*02^+^ HNSCC patients, of whom 16 had HPV-positive and 27 had HPV-negative tumors and underwent surgical treatment at Massachusetts General Hospital and Massachusetts Eye and Ear between September 2020 and December 2022 (**Table S1**). Matched peripheral blood samples, tumor tissue and, for a subset of patients (n=12), matched draining lymph nodes were processed for flow cytometry analysis of CD8^+^ T cells to identify responses against two CMV epitopes (NLV, VLE), three EBV epitopes (GLC, CLG, and FLY), and one influenza (Flu) epitope (GIL) (**Fig. 1A, B**).

**Fig. 1.**
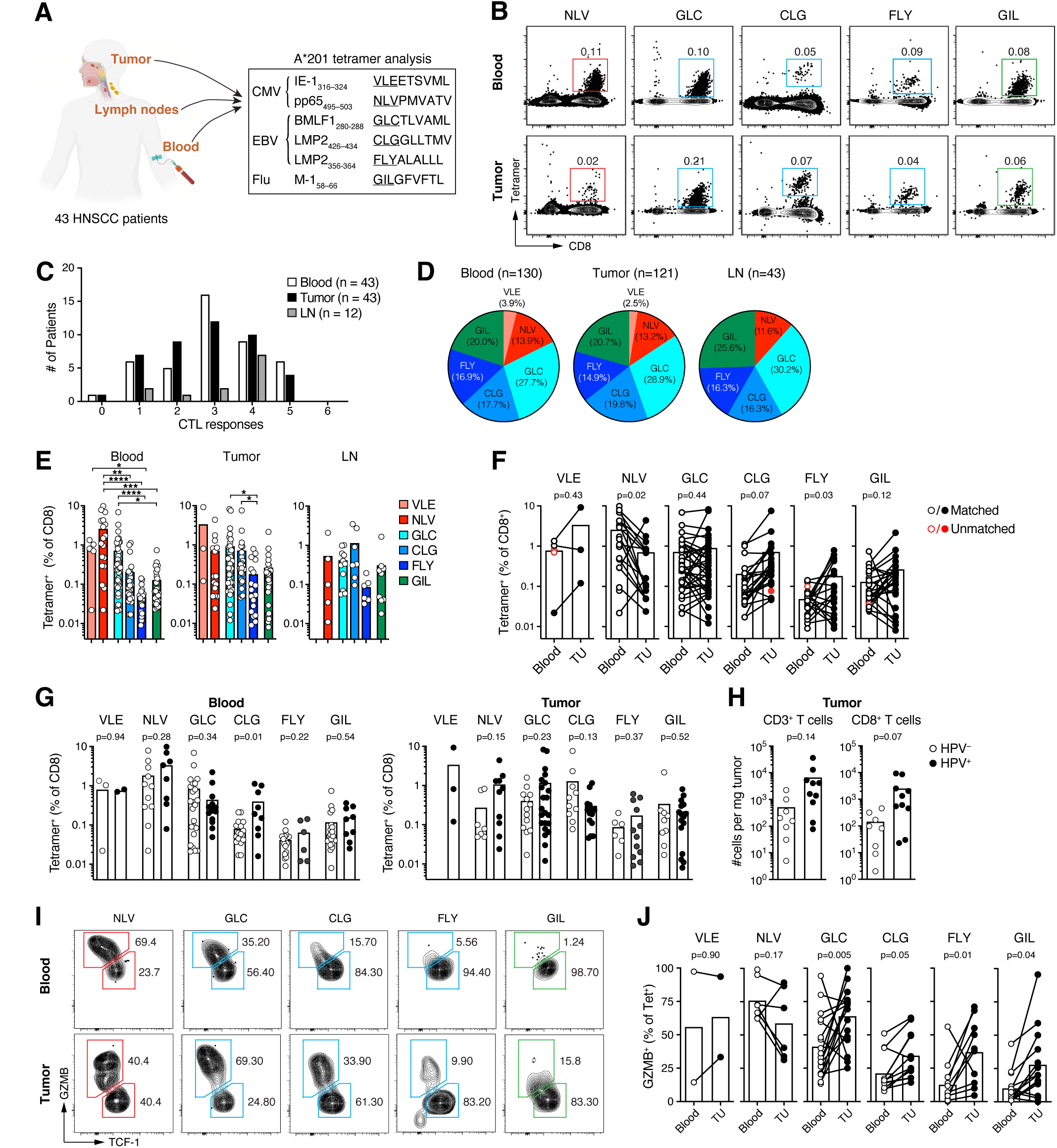
Peripheral blood analysis predicts the specificity, magnitude, and quality of intratumoral antiviral bystander CTL. (**A**) Experimental strategy. (**B**) Representative tetramer analysis of antiviral CTL in blood and tumor tissue of one HNSCC patient. (**C**) Frequency distribution of total antiviral responses detected among 6 tested in blood, tumors, and lymph nodes of individual patients (**D, E**) Frequency of patients with indicated responses (D) and magnitude of different antiviral responses (E) in blood, tumors, and lymph nodes. (**F**) Paired comparison of antiviral response magnitude in blood and tumor tissue in individual patients. Red symbols represent unmatched responses. (**G, H**) Comparison of antiviral response magnitudes (G) and of global CD3^+^ and CD8^+^ TIL abundance (H) in patients with HPV^+^ and HPV^−^ HNSCC. (**I, J**) Paired comparison of the proportion of GZMB^+^ T_EM/EFF_ states of antiviral CTL in blood and tumor tissue of individual patients. In E, P-values = 0.0332 (*), 0.0021(**), 0.0002 (***), or < 0.0001(****).

Overall, 42 out of 43 patients displayed T cell responses against at least one of the six tested epitopes. The majority responded to 2 to 4 epitopes, with none showing responses to all 6 epitopes tested (**Fig. 1C**). In the 12 lymph node samples, we detected responses to between 1 and 4 epitopes, most commonly to 4 out of the 6 CTL responses (**Fig. 1C**).

Many more patients in our cohort developed responses to the CMV pp65 antigen-derived NLV epitope, both in blood and tumors, than to the CMV immediate early-1 (IE-1) antigen-derived VLE epitope (**Fig. 1D**). Among EBV-directed responses, those targeting the lytic BMLF1 protein-derived GLC epitope were slightly more frequent than those against the latent LMP2 protein-derived CLG and FLY epitopes. All individual antiviral responses were detected at similar rates in blood, tumor, and lymph nodes (**Fig. 1D**).

Responses directed at the CMV epitopes as well as the EBV-derived GLC epitope were of the greatest magnitude in blood (**Fig. 1E**). In contrast, CMV- and EBV-directed responses were more uniform in size in tumors, except for FLY-directed responses, which were consistently the smallest along with those targeting the Flu-derived GIL epitope (**Fig. 1E**).

Paired analyses of peripheral blood and tumor samples revealed that presence in blood strongly predicted virus-specific T cell responses within the corresponding TME (**Fig. 1F**). Across all tested epitopes, 93.1% of responses (121 of 130) detected in blood were also observed in the corresponding tumor sample from the same patient. Furthermore, 36 of 37 responses (97.3%) detected in lymph nodes in the subset of 12 patients were matched in their tumor tissue (**Fig. S1A**). Interestingly, many of the EBV CLG- and FLY-directed responses were enriched in their magnitude in tumor tissue compared to blood, while CMV NLV-responses were, in 2 out of 3 cases, less abundant in tumors than in blood (**Fig. 1F**).

When we stratified patients based on tumor HPV status, we observed no consistent differences in the frequency of antiviral CTL among CD8^+^ T cells, except for EBV CLG-directed responses, which were of greater magnitude in the blood (but not in tumor tissue) of patients with HPV^+^ compared to HPV^−^ HNSCC (**Fig. 1G**). However, HPV^+^ tumors were generally more heavily infiltrated by CD8^+^ T cells (**Fig. 1H**), and hence antiviral CTLs were more abundant in those compared to HPV^−^ tumors. We did not observe consistent differences, neither in the proportion of different antiviral responses nor their magnitude, when stratifying patients by age (older versus younger than 65 years, **Fig. S1C**), or based on the anatomical location of their lesion (larynx, oral cavity, versus oropharynx, **Fig. S1D**).

Together, our results indicate that peripheral blood measurements of the repertoire of antiviral CD8^+^ T cells serve to reliably predict their presence in both HPV^−^ and HPV^+^ HNSCC TMEs, providing a noninvasive window into the intratumoral immune landscape to guide the selection of viral epitopes employed in personalized APEC therapy.

Based on the presumed mechanism of action of APEC therapy, namely the loading of cancer cell surface MHC I molecules with viral peptides to target them for destruction by bystander CTLs, we hypothesized that antiviral CD8^+^ T cells with an effector or effector memory phenotype (T_EFF/EM_) have more pronounced anti-tumor activity than stemlike memory T cells (T_SL_) when recruited into the anti-tumor response through APEC therapy. The former express Granzymes that equip them with immediate cytotoxic effector function, while the latter first need to interact with tumor-associated dendritic cells, which may not be loaded with APEC-delivered peptides, in order to acquire effector function (16). We therefore examined T cells from a subset of patients for expression of Granzyme B (GZMB), a canonical cytotoxic effector protein, as well as T cell factor (TCF)-1, the hallmark transcription factor characterizing T_SL_, both in circulating and tumor-infiltrating antiviral CTL (**Fig. 1I**). In blood, we observed the highest proportion of GZMB-expressing cells in responses against the CMV-derived NLV epitope in blood, with on average comparable frequencies in tumor tissue (**Fig. 1J**). In contrast, GZMB^+^ T_EFF/EM_ were less abundant among EBV- as well as Flu-directed responses in blood, and the observed EBV FLY-and Flu GIL-directed responses were instead dominated by TCF-1^+^ T_SL_. However, all of these responses were enriched for GZMB^+^ T_EFF/EM_ in tumor tissues compared to blood (**Fig. 1I, J**). Hence, the proportions of T_EFF/EM_ and T_SL_ in antiviral responses varied by epitope and potentially by viral antigen, but not necessarily by viral species. The CMV-derived NLV and the EBV-derived GLC epitopes emerged as attractive targets for APEC therapy in HLA-A*201^+^ HNSCC patients, because they were commonly targeted and their CTL responses were not only of large magnitude but also contained the largest proportions of T_EFF/EM_ states in tumor tissue. However, responses with a high proportion of T_SL_ states in blood were enriched for T_EFF/EM_ states in tumor and could therefore unfold APEC-induced anti-tumor activity as well.

### A screen for protease activity shared across human HNSCC cells

The design of APECs with optimal anti-tumor activity in a large proportion of patients with HNSCC will require the use of peptide linkers that are efficiently cleaved by proteases secreted or expressed at the cell surface by the majority of cancers across all patients with both HPV^+^ and HPV^−^ head and neck cancers. We hypothesized that while the full spectrum of proteolytic activities may generally vary not only between cancer types but also between individual cancer patients (17), HNSCCs may also exhibit some shared proteolytic activities that could guide the selection of peptide linkers in APECs suitable for use across most or all patients. We sought to identify such shared activities through a combined data mining and functional screening strategy (**Fig. 2A**). We first cross-referenced 718 known and putative human proteases listed in the MEROPS database (18) with a list of 7590 known and putative cell surface and secreted proteins derived from protein mass spectrometry analyses and *in silico* predictions (19) to identify 430 putative cell-surface (n=159) or secreted (n=271) proteases (**Table S2A-E**). We then screened those for over-expression in HNSCC tumors by comparing bulk RNA-Seq data sets from tumor tissue (n=520) and adjacent normal tissue (n=44) deposited in The Cancer Genome Atlas (TCGA) (20) and processed through the Broad Institute Firehose pipeline. Excluding transcripts undetectable in either malignant or healthy tissue, this identified 27 candidate proteases overexpressed more than 5-fold in malignant compared to healthy head and neck tissues of which we classified 24 as bona fide proteases (**Fig. 2B**), and for 12 of which the MEROPS database listed non-synthetic, physiological or pathological cleavage motifs (**Table S2F**). We used the MEROPS database as well as published reports referenced therein to select peptide sequences cleaved by these proteases. Based on the strength of published evidence for bioactivity and prioritizing peptide sequences described to be selectively cleaved by only one of the 12 proteases, we assembled a list of 142 peptide sequences. To those we added 40 sequences identified as targets for cancer cell proteases in our previous studies (11, 12), as well as 4 sequences cleaved by the protease ADAM10 based on its overexpression and potential role in disease progression in oral HNSCC (21, 22) (**Table S2G**). The resulting 186 peptide sequences were named after the in total 29 protease activities for which they were selected, keeping in mind that they may also be cleaved by additional proteases not included in our screen or by unknown activities of those included.

**Fig. 2.**
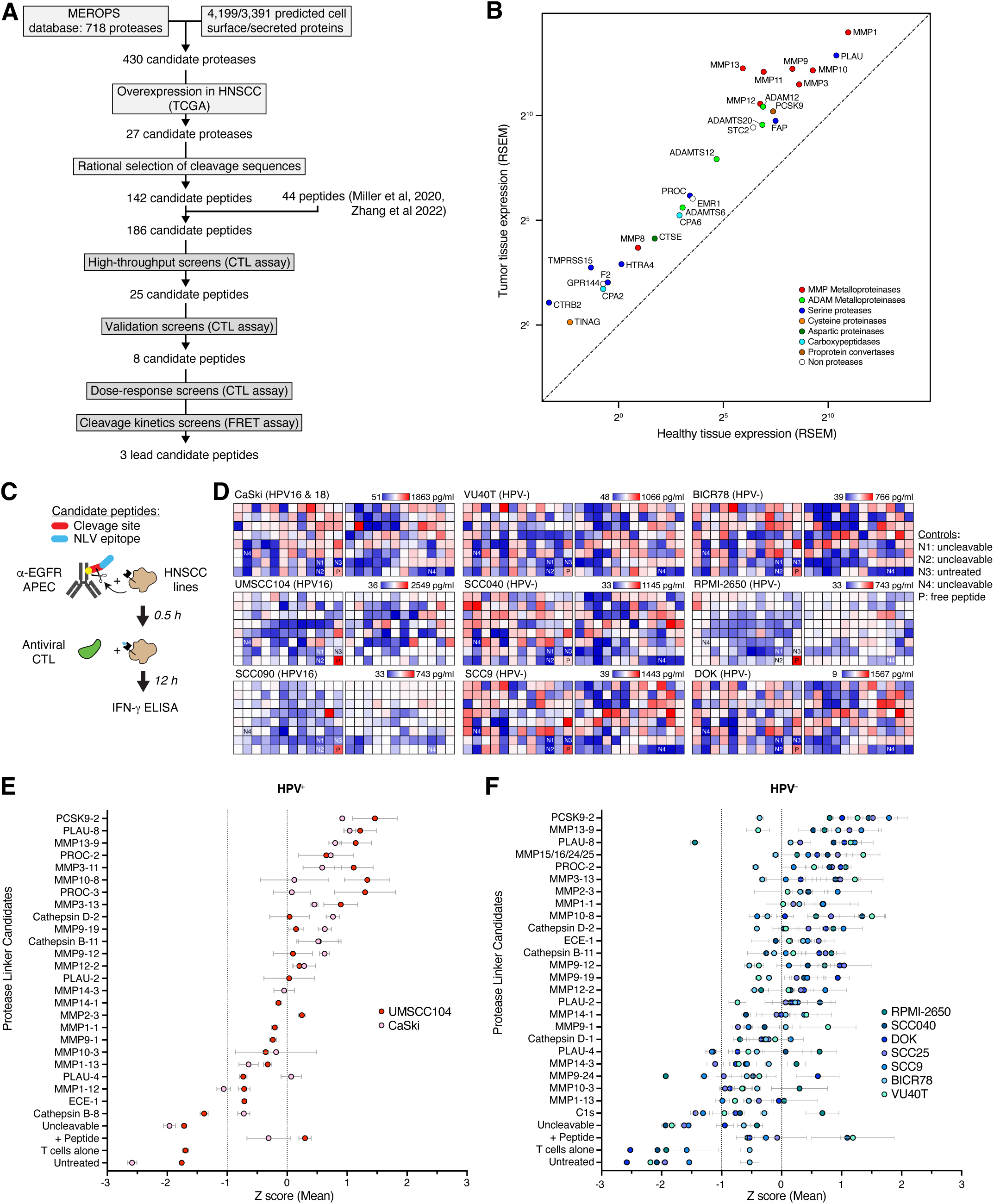
A data-mining and functional screening-based strategy to identify suitable protease activities for APEC therapy of HNSCC. (**A)** Experimental strategy. (**B**) Expression of indicated proteases based on RNA-Seq data in the TCGA. (**C**) Principle of ‘CTL assay’ to probe protease activities in HNSCC cancer cell lines based on their known cleavage motifs. (**D**) Cleavage activities determined by quantities of IFN-γ secreted by CTL in response to loading of viral peptide cleaved off APECs by cancer cell proteases. Heatmap displayed normalized for each cancer cell line. Negative and positive controls are indicated. (**E,F**) Ranking of z-scored IFN-γ quantities for top 25 peptides cleaved by HPV^+^ (E) and HPV^−^ (F) cell lines in validation screen. Each peptide was tested in 1-3 experiments. RSEM: RNA-Seq by Expectation Maximization

To assess how widely proteases expressed by cancers from different patients cleave the 186 candidate peptides we assembled a panel of 6 HPV^−^ HNSCC (VU40T, SCC040, SCC9, BICR78, RPMI-2650, DOK), 2 HPV^+^ HNSCC (UMSCC10, SCC090), and one HPV^+^ cervical cancer cell line (CaSki) for a total of 9 cancer cell lines derived from HLA A2^+^ patients to facilitate functional screening against human CTL recognizing the HLA-A2-restricted HCMV pp65 antigen-derived NLV epitope. The cervical cancer-derived CaSki line was added to compensate for the paucity of available HPV^+^ HLA-A2^+^ HNSCC lines. We selected the chimeric IgG1 α-EGFRvIII antibody Cetuximab as a backbone for the construction of HNSCC-targeted APECs, since EGFRvIII is expressed by the vast majority of both HPV^+^ and HPV^−^ HNSCC cancers, serves as an oncogenic driver due to its constitutive signaling activity, and because Cetuximab is already FDA-approved for the treatment of HNSCC in combination with chemotherapy (23), potentially lowering regulatory hurdles during future clinical development. 8-mer peptides containing the cleavage motifs were expressed as C-terminal fusions with the CMV NLV peptide epitope and conjugated to Cetuximab lysine residues using standard maleimide chemistry under conditions that yield an average stoichiometry of two peptides per antibody in order to maintain solubility of the resulting NLV-APECs (11).

For an initial, high-throughput screen, we incubated 3 HPV^+^ and 6 HPV^−^ cancer cell lines with Cetuximab-based APECs constructed with all 186 candidate peptides, removed APECs, co-cultured cancer cell lines for 12 h with NLV-specific CTL lines derived from PBMC of A2*201^+^ healthy donors, and analyzed supernatants for IFN-γ secreted by those CTLs using ELISA (’CTL-assay’, **Fig. 2C, D**), as described (11). We then re-screened the 25 top-performing peptides against the HPV^+^ lines UMSCC104 and CaSki (**Fig. 2E**) and the six HPV^−^ HNSCC lines (**Fig. 2F**) using the same assay, in many instances in multiple repeats. This yielded overlapping, but non-identical groups of top-performing peptides for HPV^+^ and HPV^−^ cancer lines (**Fig. 2E, F**). From the top-ranked peptides we selected partially overlapping sets of 8 each for a total of 12 peptides for more granular assessment on HPV^+^ and HPV^−^ cancer cells for kinetic measurements of cleavage activity. Free peptides were fluorescently tagged on their N- and C-termini to monitor their cleavage using Förster resonance energy transfer (’FRET assay’) (**Fig. 3A**). Focusing on peptide cleavage activity during the first hour after exposure to cancer cells, we found that the PLAU-cleavable peptide PLAU-8 (PGARGRAF) was consistently top-ranked based on the efficiency at which it was cleaved by all cell lines tested, followed by the MMP10-cleavable peptide MMP10-8 (RPSQRSKY) (**Fig. 3B-E**). MMP13-9 (FGVKRAYY) peptide ranked third for HPV^−^ and fifth for HPV^+^ lines (**Fig. 3B-E**), while PROC-2 (RLKKSQFL) ranked third for HPV^+^ lines but was not tested on HPV^−^ lines.

**Fig. 3.**
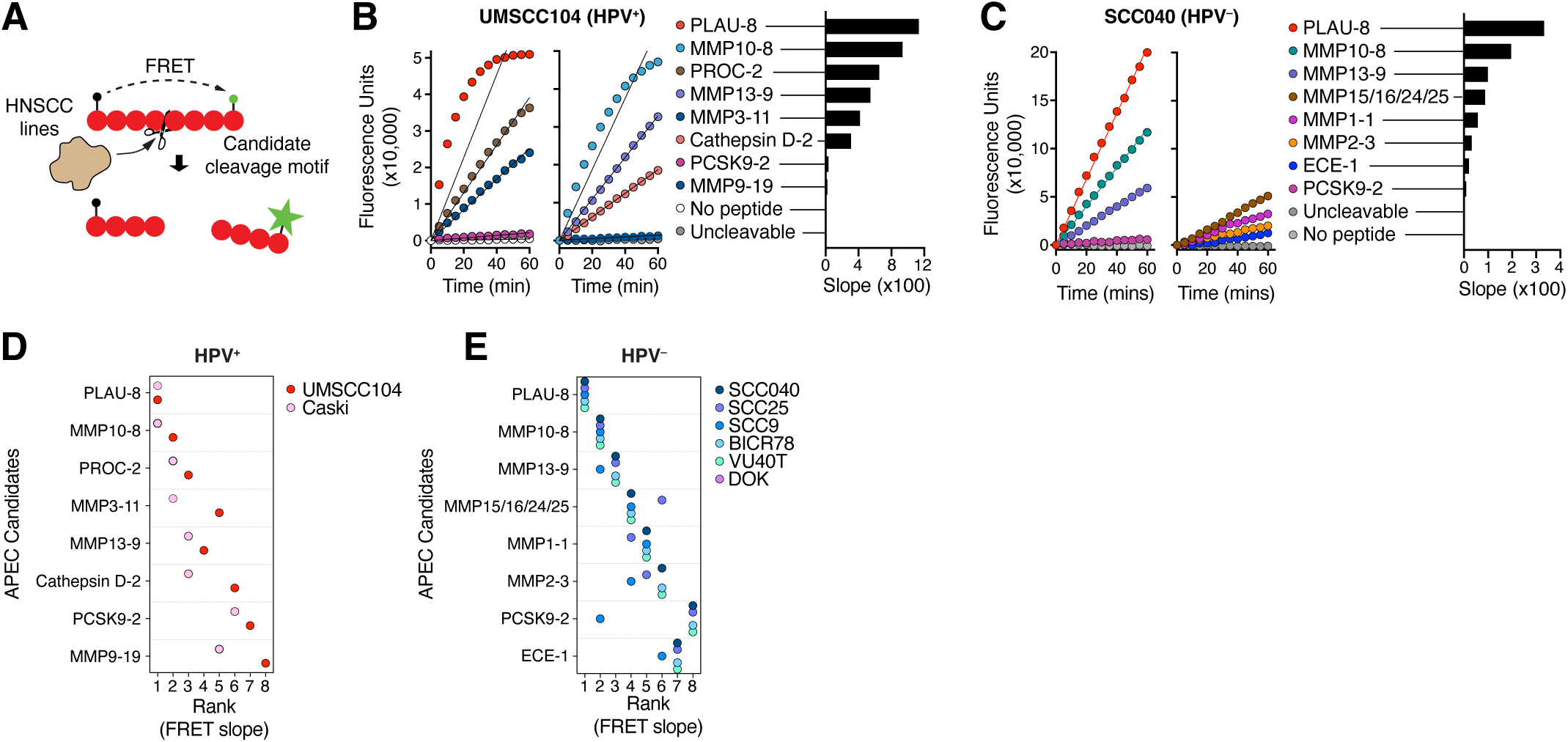
Kinetics protease activity analyses nominate PLAU, MMP10, and MMP13 protease activities for APEC therapy in HNSCC. (**A**) Principle of ‘FRET assay’ to probe protease activities in HNSCC cancer cell lines based on their known cleavage motifs. (**B, C**) Representative traces of fluorescence increase resulting from loss of FRET due to peptide cleavage of labeled peptides during the first 60 min after exposure to an HPV^+^ HNSCC line (B) and an HPV^−^ HNSCC line (C). Lines in graphs indicate linear regression functions and bar graphs at the right show slopes of regression functions as indicators of cleavage efficiencies. (**D, E**) Summary of rankings of each protease activity in each HNSCC cell line for HPV^+^ HNSCC lines (D) and HPV^−^ HNSCC lines (E).

In summary, our screen identified three peptides cleaved by the proteases PLAU, MMP10, and MMP13 proteases, all three of which are overexpressed in HNSCC tumors. These peptides thus represent promising lead candidates for the development of APECs aimed at treating both HPV^+^ and HPV^−^ HNSCCs.

### A syngeneic mouse model for antibody peptide epitope conjugate therapy

Our prior studies have demonstrated that APECs can have anti-tumor activity in murine and zebrafish xenograft preclinical cancer models adoptively transferred with *ex vivo*-primed CTLs (11–13). However, these models do not allow for the assessment of the interplay of APEC therapy with an intact host immune system or the anti-tumor activity of *in vivo*-primed antiviral CTLs in a setting of latent viral infection. They also do not enable evaluation of potential on-target, off-tumor toxicity due to the widespread expression of the APEC-targeted tumor-associated antigens in healthy tissues, a complication observed for instance in CAR adoptive cell therapies targeting such antigens (24). In order to address these open questions in an ICB therapy-resistant syngeneic mouse model of cancer, we needed to select (i) a suitable murine cancer cell line, (ii) a cancer cell-expressed cell surface antigen, (iii) an APEC backbone antibody binding this target, (iv) a cancer cell-expressed protease activity, and (v) a model of viral infection mimicking human infection leading to the formation of tumor-infiltrating antiviral CD8^+^ bystander T cells.

Given the similarities in the biology of murine CMV (MCMV) and HCMV (25, 26), we selected MCMV infection to model the formation of tumor-infiltrating anti-HCMV CTL and test their suitability for APEC therapy. We infected C57BL/6 mice with the K181 strain of MCMV through intraperitoneal injection and monitored anti-MCMV CTL responses against the previously characterized viral HGI(RNASFI), TVY(GFCLL), and SSP(PMFRV) epitopes representing three different viral antigens (27), using tetramer staining of circulating CD8^+^ T cells (**Fig. 4A, B**). We observed rapid expansion of HGI-specific cells, followed by rapid contraction and formation of a stable memory population by 3 weeks after infection, reflecting a canonical T cell response pattern (**Fig. 4C**). TVY- and SSP-specific cells, in contrast, did not contract, but persisted at elevated frequencies reached during the acute response phase, or continued to expand further, reflecting an inflationary response pattern previously described for these epitopes during MCMV responses in mice (**Fig. 4C**) (27), as well as for select epitopes of HCMV (28) and other latent viral infections, such as EBV (29).

**Fig. 4.**
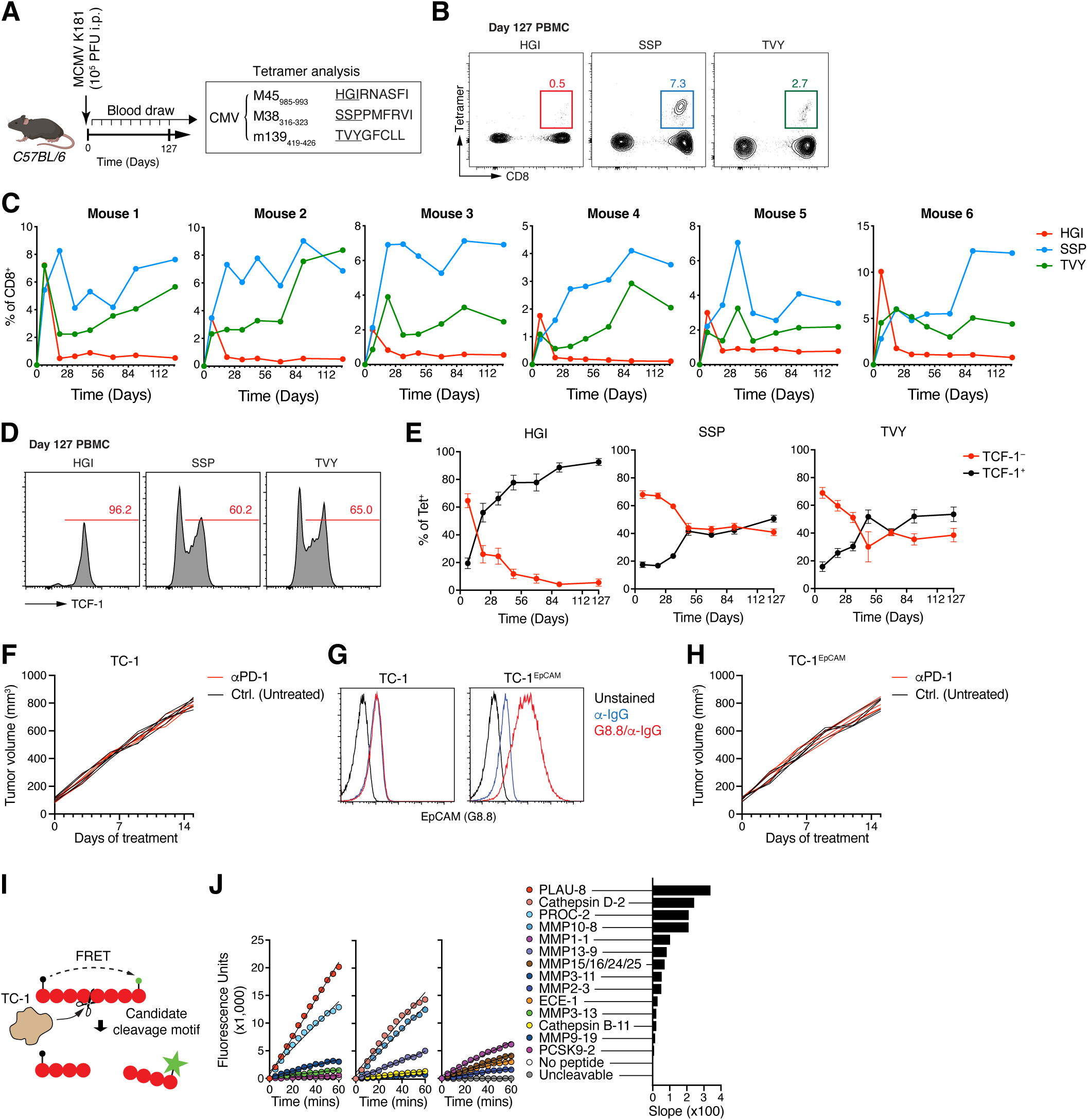
A model for EpCAM-targeted APEC therapy of ICT-resistant HPV^+^ tumors in immune-competent, MCMV-infected mice. (**A**) Experimental strategy to characterize MCMV-directed CTL responses (**B**) Representative tetramer analysis of antiviral CTL in blood 4 months after infection (**C**) Magnitude and dynamics of antiviral CTL responses in a group of 6 infected animals. (**D, E**) Representative analysis (D) and dynamics (E) of the proportion of antiviral TCF-1^−^ T_EFF/EM_ and TCF-1^+^ T_SL_ in blood 4 months after infection. (**F**) Response of TC-1 tumors to PD-1-targeted ICT (250 μg RMP1-14 i.p., 3x/week) (**G**) Expression of EpCAM on TC-1 before (left) and after (right) lentiviral transduction with murine EpCAM (**H**) Response of TC-1^EpCAM^ tumors to PD-1-targeted ICT. (**I**) Principle of ‘FRET assay’. (**J**) Traces of fluorescence resulting from loss of FRET due to peptide cleavage of labeled peptides during the first 60 min after exposure to TC-1 cells. Lines in graphs indicate linear regression functions and bar graphs at the right show slopes of regression functions as indicators of cleavage efficiencies.

Considering that the presumed mechanism of action of APECs requires immediate effector function of redirected CTLs and that the proportion of TCF-1^+^ T_SL_ and TCF-1^−^ T_EF/EM_ is variable in human antiviral CTLs (**Fig. 1I, J**), we also examined the proportion of T_SL_ and T_EFF/EM_ among the different anti-MCMV responses. While TCF-1^−^ T_EF/EM_ dominated all responses during the antiviral effector phase, their proportion rapidly declined in the HGI-directed responses, with >90% of cells adopting a TCF-1^+^ stemlike state after 4 months (**Fig. 4D, E**). In contrast, the proportion of TCF-1^−^ T_EFF/EM_ cells also initially declined but then stabilized at 40-50% for the inflationary responses directed against TVY and SSP (**Fig. 4D, E**). Hence, similar as for certain CMV- and EBV-epitope-targeted responses observed in our human studies (**Fig. 1I, J**), we predicted that APECs targeting inflationary antiviral CTL responses directed against the TVY and SSP epitopes would be more effective than those targeting canonical memory response against the HGI epitope.

The TC-1 cancer cell line derived from lung epithelial cells of C57BL/6 mice through engineered expression of Ras and the oncogenic HPV-16 antigens E6 and E7 is commonly used as a model for HPV-associated epithelial cancers, including HPV^+^ HNSCC (30). We confirmed that TC-1 is resistant to αPD-1 ICB therapy (**Fig. 4F**), as widely reported (31–34). Epithelial cell adhesion molecule (EpCAM) is a commonly selected target antigen for the development of antibody-drug conjugates as well as chimeric antigen receptor (CAR) T cell therapies based on its widespread over-expression on human epithelial cancers (35). However, its expression has been lost on many commonly used epithelial mouse cancer lines, such as MC38, ID8, as well as TC-1 (**Figs. 4G, S2A)**. We, therefore, engineered TC-1 to express mouse EpCAM through lentiviral transduction (**Figs. 4G, S2B**). TC-1^EpCAM^ tumors grew in C57BL/6 mice at a rate comparable to the parental TC-1 and retained their resistance to αPD-1 treatment (**Fig. 4H**).

To target murine EpCAM, we selected the rat IgG2a antibody clone G8.8 as an APEC backbone (36). To select a suitable protease-cleavable linker coupling viral antigen peptide epitopes to G8.8 for APEC treatment of TC-1^EpCAM^ tumors, we again used the ‘FRET assay’ to screen a panel of well-characterized protease cleavage motifs that included the top scoring motifs from our human HNSCC screen. Similar to human HNSCC lines, TC-1-expressed proteases most effectively cleaved the PLAU-8 motif, followed by motifs cleaved by Cathepsin D, PROC, and MMP-10 (**Fig. 4I, J**). We therefore selected the PLAU-8 cleavage motif to construct our G8.8-based model APEC.

### Inflationary memory responses are superior targets for APEC therapy

To examine whether *in situ* primed anti-MCMV responses could be redirected against cancer cells using EpCAM-binding APECs to produce an anti-tumor effect, we implanted TC-1^EpCAM^ tumors subcutaneously (s.c.) into mice 2 months following MCMV infection, when stable memory T cell responses had formed. Each time a tumor reached a size of ∼100 mm^3^, the animal was randomized to a group and APEC treatment initiated at a dose of 25 μg intraperitoneally (i.p.), three times a week (**Figs. 5A, S3A**). Tumors in animals treated with TVY-directed control APECs incorporating an uncleavable linker (GGGGG) grew at a similar rate as in untreated MCMV-infected animals (**Figs. 5B, S3A**). Hence, neither the binding of APECs to cancer cells nor the introduction of a viral peptide that cannot be proteolytically released from the APEC backbone had anti-tumor effects. In contrast, APECs targeting the inflationary TVY or SPP-directed responses and incorporating the PLAU-8 cleavage motif rapidly induced arrest of tumor growth, and in some cases tumor elimination, persisting for the duration of treatment (**Figs. 5B, S3A**). APECs targeting HGI-directed CTLs also showed anti-tumor activity, albeit less pronounced, and tumors eventually resumed their original growth rate (**Figs. 5B, S3A**).

**Fig. 5.**
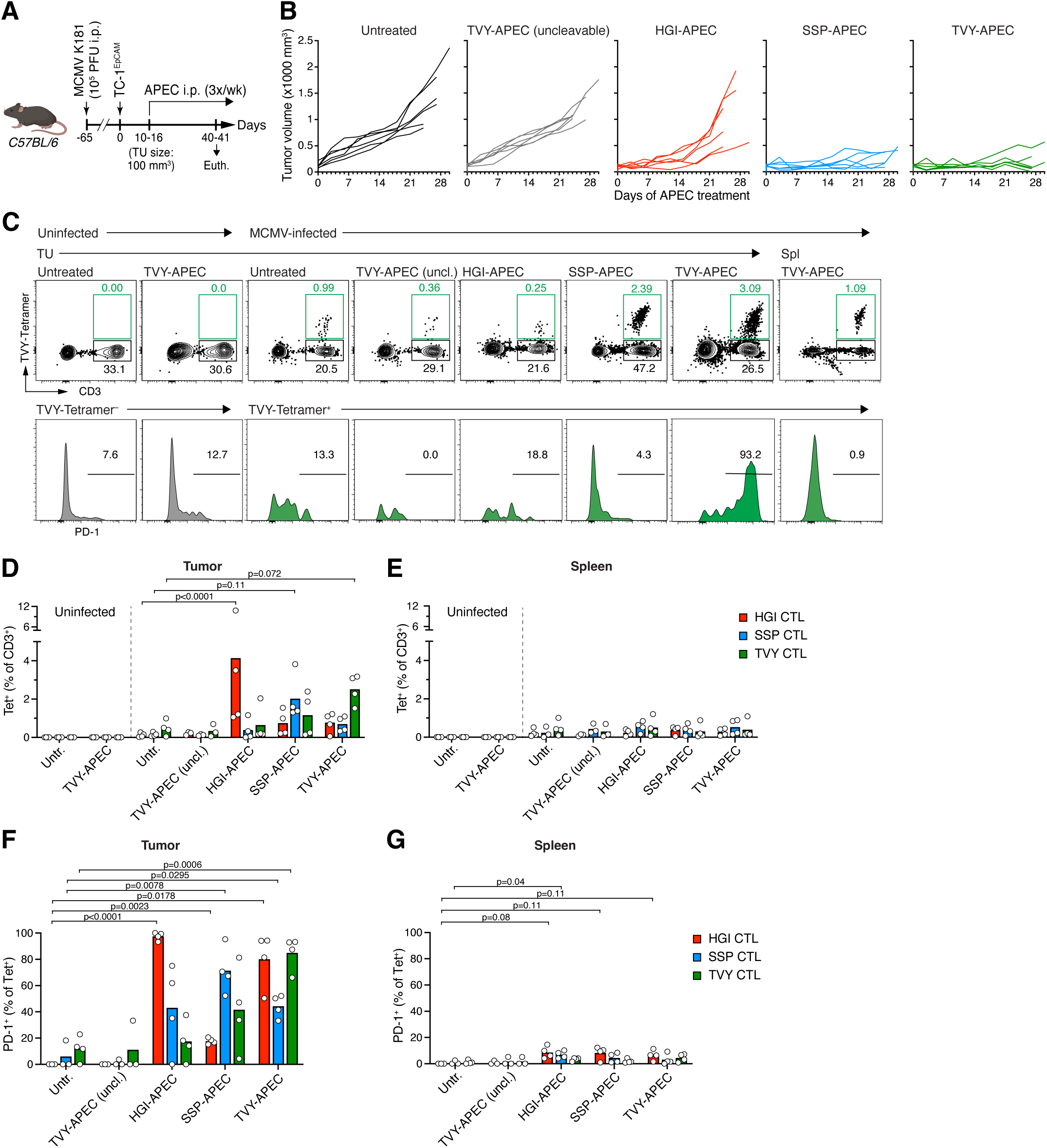
Superior anti-tumor activity of APECs targeting inflationary CTL responses. (**A**) Experimental design. (**B**) Tumor growth curves post treatment start of TC-1 tumors in individual mice treated with indicated APECs. Results are representative of two independent experiments (**C-G**) Frequency (C (top), D, E) and PD-1 expression (C (bottom), F, G) of antiviral CTL in tumors and in spleens (Spl) of MCMV-infected or uninfected animals 6 days after initiation of indicated treatments.

Accordingly, animals treated with APECs targeting the inflationary TVY or SPP-directed responses showed better survival (post tumor implantation) than those treat with HGI-directed APEC, uncleavable control APECs, or that were left untreated (**Fig. S3B**). Hence, APECs targeting inflationary antiviral memory responses, which include T_EFF/EM_ cells, can sustain robust anti-tumor activity. In contrast, targeting a conventional memory population dominated by T_SL_ results in a weaker and more transient response, potentially due to their smaller population size or limited immediate cytotoxic effector activity.

### Antiviral T cell activation is restricted to tumor-infiltrating bystander cells

To assess how effectively APEC treatment activated antiviral bystander CTLs in tumors, we analyzed tumor-infiltrating antiviral CTLs one week after treatment initiation for their expression of PD-1, a marker of recent TCR stimulation (37). No antiviral CTLs could be detected in TC-1^EpCAM^ tumors in uninfected mice, but both conventional and inflationary responses were represented in tumors of MCMV-infected animals (**Fig. 5C, D**). However, in the absence of APEC treatment or following treatment with uncleavable control APECs, antiviral T cells expressed undetectable or low levels of PD-1 (**Fig. 5C, F**), which has been suggested to occur in response to antigen-independent stimulation with type I IFN or other cytokines (38, 39). In contrast, tumor-infiltrating HGI-, SSP-, and TVY-CTLs expressed high levels of PD-1 in mice treated with HGI-, SSP-, and TVY-APECs, respectively. (**Fig. 5C, F**). Interestingly, treatment with TVY-APECs also elicited some PD-1 expression on tumor-infiltrating HGI- and SSP-CTLs, perhaps indicating enhanced expression or release of MCMV antigens in the TME, or their enhanced presentation to T cells, which could be driven by IFN-γ released by APEC-activated TVY-CTLs. Importantly, we observed no or only a minor increase in PD-1 expression on antiviral CTLs in the spleens of treated animals, indicating that APECs did not induce systemic activation of antiviral CTLs (**Fig. 5C, E, G**). Hence, APECs efficiently activate antiviral bystander CTL in the TME of the TC-1 HPV^+^ epithelial tumor model, but not systemically, facilitating sustained tumor growth control, especially when targeting inflationary antiviral CTL responses.

### APEC activity depends on cancer cell protease activity

In addition to cancer cells, various cells of the TME, and especially myeloid cell types, exert proteolytic activity. For instance, macrophages express PLAU and require PLAU for their antimicrobial function (40). In order to test if the proteolytic activity of cancer cells is redundant and non-malignant cells of the TME can activate APECs, we sought to develop a system where cancer cells cannot activate APECs. Since PLAU is the only known protease to cleave the PLAU-8 motif, other than plasmin, which derives from PLAU-mediated cleavage of plasminogen, we used CRISPR/Cas9 gene editing to delete PLAU from TC-1^EpCAM^ cells (**Fig. 6A**). PLAU^Null^ TC-1^EpCAM^ cancer cells indeed selectively lost the capacity to activate PLAU-8-based APECs in the setting of our ‘CTL-assay’, while they retained the capacity to activate MMP13-8-based APEC (**Fig. 6B, C**). Compared to PLAU-sufficient TC-1^EpCAM^, PLAU-deficient TC-1^EpCAM^ tumors grew at a slightly slower rate (**Fig. 6D, E**). Therefore, we assessed the treatment effect of PLAU-8-based APECs separately in TC-1^EpCAM^ and PLAU^Null^ TC-1^EpCAM^ tumors and found that in the latter both HGI- and TVY-APECs lost their efficacy almost completely (**Fig. 6E**). This was paralleled by a failure of antiviral CTLs to accumulate in the PLAU^Null^ TC-1^EpCAM^ TME (**Fig. 6F, G**), and those that did enter the TME failed to induce PD-1 expression (**Fig. 6F, H**). In line with the presumed mechanism of action, the activity of APECs therefore depends on the proteolytic function of cancer cells, and APEC anti-tumor activity requires matching of used peptide linkers with cancer cell protease activities.

**Fig. 6.**
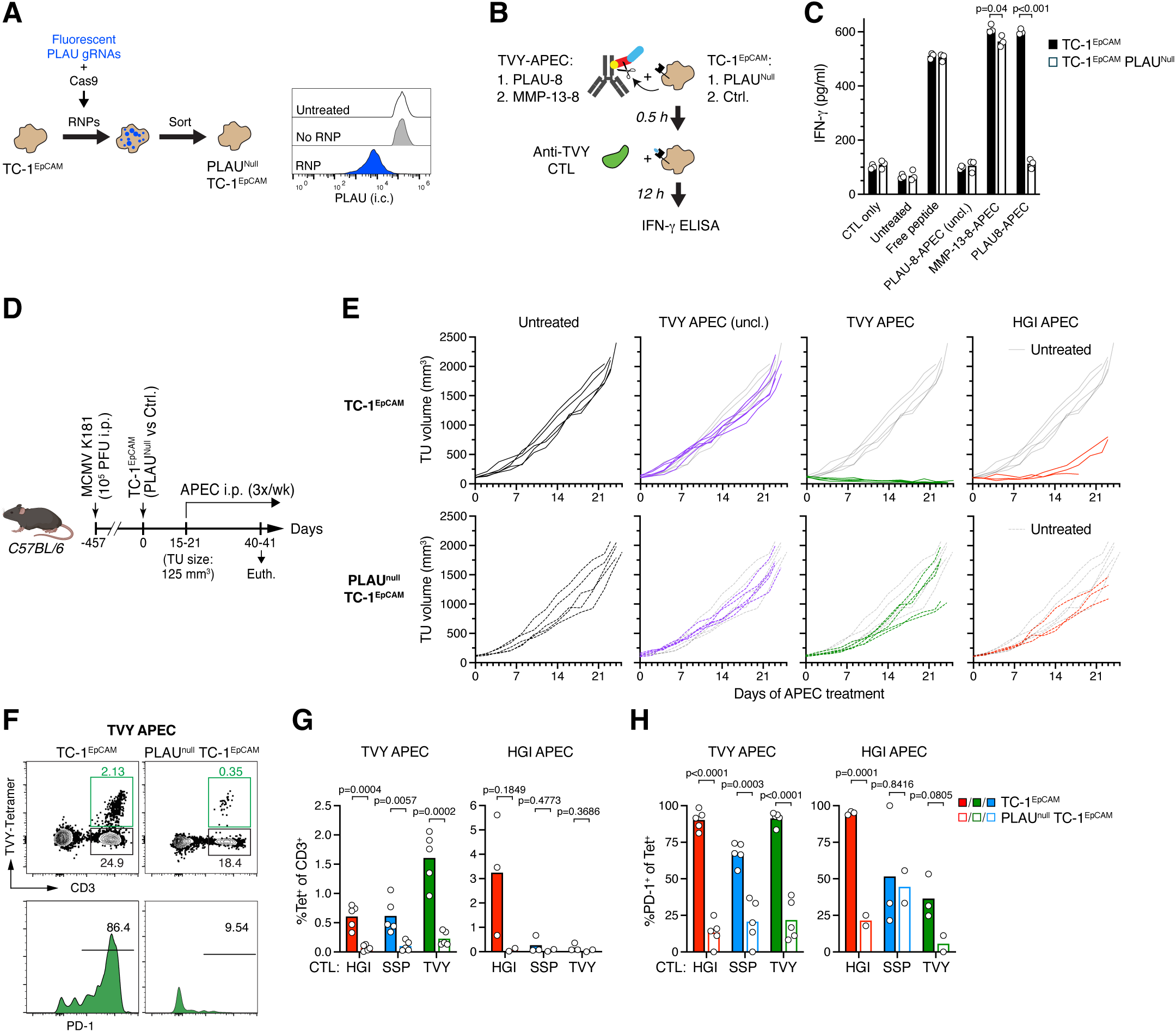
Cancer cell-expression of PLAU is required for antitumor activity of APECs incorporating a PLAU-sensitive linker. (**A**) Experimental design to generate PLAU-deficient TC-1 cells. (**B, C**) CTL assay (B) to assess activation of PLAU-8-APECs by PLAU^Null^ TC-1 and control TC-1 cancer cells (C). (**D**) Experimental design. (**E**) Growth curves of TC-1^EpCAM^ (top) and PLAU^Null^ TC-1^EpCAM^ tumors (bottom) in individual mice treated with indicated APECs. Growth curves in untreated animals are shown along growth curves from treated animals for direct comparison (grey and dashed in grey lines). (**F-H**) Frequency (F (top) and G) and PD-1 expression (F (bottom) and H) of antiviral CTL in PLAU-sufficient and PLAU^Null^ TC-1^EpCAM^ tumors 6 days after initiation of treatments with indicated APECs.

### Lack of evidence for APEC-induced on-target, off tumor toxicity

While EpCAM is overexpressed on many cancer epithelial cancers, it is also expressed at lower levels in healthy epithelial tissues (41, 42). PLAU is also expressed in some of those tissues (43), which may also be infiltrated by MCMV-reactive CD8^+^ tissue-resident memory T cells (44, 45). APEC binding could therefore also redirect tissue-resident MCMV-reactive CTLs against healthy epithelial cells, causing on-target, off-tumor toxicity. However, over the course of three independent APEC treatment studies, mortality in APEC-treated animals (4 out of 52 mice) did not exceed mortality in control animals (12 out of 50 mice). We also did not observe diarrhea, a sign of intestinal toxicity in patients undergoing EpCAM/CD3-targeted bispecific T cell engager therapy (46), increased weight loss (**Fig. 7A**), or any other clinical signs indicative of a reduction in their general state of health, compared to untreated, tumor-bearing animals (**not shown**).

**Fig. 7.**
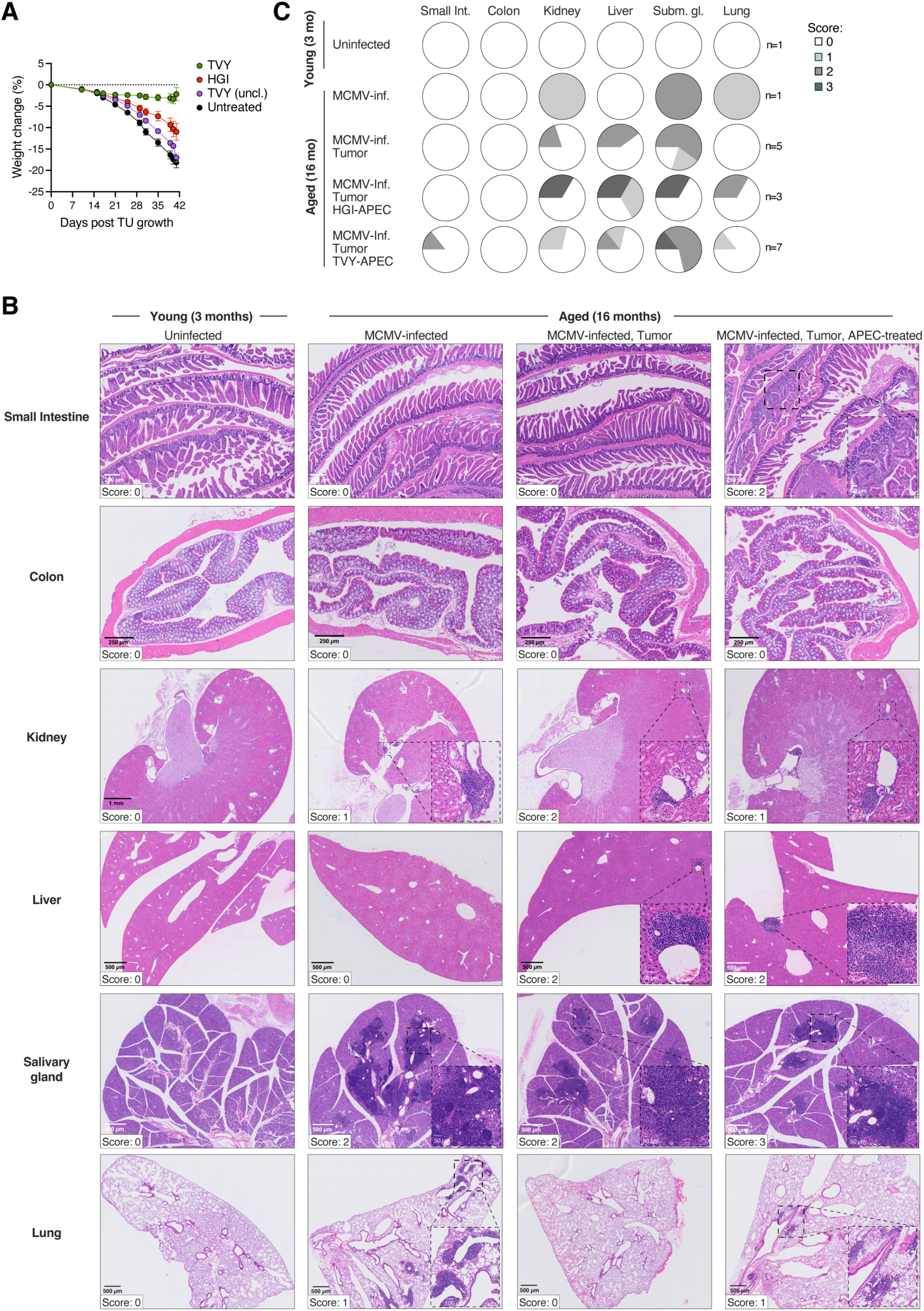
No evidence for APEC-induced off-tumor on-target toxicity in epithelial tissues. (**A**) Weight curves of 16 months old, latently MCMV-infected C57BL/6 mice under APEC therapy (as described in Fig. 6A). Means and SEM are shown. (**B, C**) Representative H&E-stained histological sections and histopathological scoring (B) and summary of histopathological scores (C) of indicated tissues from HGI-APEC-, TVY-APEC-treated, or untreated 16 months-old, latently MCMV-infected animals. Tissue from an age-matched MCMV-infected but tumor-free and one young and uninfected animal are shown for reference. Insets in B highlight lymphocytic infiltrates.

To further assess whether APEC therapy induced tissue damage or inflammatory changes in healthy tissues where EpCAM is expressed, we histologically examined epithelial tissues, including small and large intestines, kidneys, livers, salivary glands, and lungs, in animals from mice that were MCMV-infected when 2 months old and aged until 15 months before APEC treatment. Aged mice were selected for this analysis since cancer immunotherapy-induced toxicity is much more commonly observed in aged than in young mice (47, 48). Comparing APEC-treated with untreated tumor-bearing mice, we did not observe an increase in pathological changes, such as mucosal erosion or atrophy, or damage to submucosa and muscularis, in the small intestines or colons, where EpCAM is mostly highly expressed (41), with the exception of a lymphocytic infiltrate in the small intestine of one TVY-APEC-treated animal (**Fig. 7B, C**). Perivascular or peribronchial lymphocytic infiltrates were more commonly observed in the kidneys, livers, salivary glands, and lungs of a subset of animals. However, comparable changes were also observed in an aged-matched and latently MCMV-infected but tumor-free animal, but not in a young uninfected, tumor-free animal (**Fig. 7B, C**), indicating that these infiltrates were caused by a combination of advanced age (16 months), latent MCMV-infection, and tumor growth. Hence, continuous EpCAM-targeted APEC therapy over the course of three or more weeks does not appear to induce epithelial tissue damage or exacerbate inflammatory changes observed in aged and long-time MCMV-infected animals as a result of redirecting tissue-resident antiviral CTL against healthy EpCAM^+^ cells.

## DISCUSSION

In this study, we examined tissues from human HNSCC patients and developed a preclinical murine model of HNSCC cancer in the setting of latent CMV infection to explore how APEC therapy can harness the function of tumor-infiltrating antiviral bystander cells for cancer therapy. We find that in an immune-competent setting, APECs targeting inflationary CTL responses elicited the most robust and sustained control of anti-PD-1 therapy-resistant tumors and that tumor tissue of HNSCC patients was consistently populated by antiviral CTL responses whose specificity was predictable from responses found in the peripheral blood. We furthermore observed that a peptide motif targeted by the PLAU protease is efficiently cleaved by protease activity in a wide range of both HPV^−^ and HPV^+^ human HNSCC cancer lines as well as by a murine HPV^+^ epithelial cancer cell line.

The presumed mechanism of action of APECs is that upon binding to their target cell surface antigen, cancer cell-expressed proteases cleave and release their viral peptide cargo for loading onto MHC I molecules on the surface of cancer cells, but not other antigen-presenting cells of the TME, including dendritic cells (DCs). One prediction of this model is that APECs targeting antiviral CTL responses with a predominant T_EFF/EM_ state should possess greater immediate anti-tumor efficacy, since T_SL_ in the TME first require local interactions with antigen-presenting DCs to acquire cytotoxic and other anti-tumor effector functions (16). Our preclinical studies corroborated this prediction. They furthermore showed that redirected inflationary responses enriched in TCF-1^−^ T_EFF/EM_ states are more long-lived and provide more sustained anti-tumor activity under APEC therapy than canonical memory T cell responses. Viral epitopes targeted by conventional memory responses require processing through the immunoproteasome, which occurs primarily in the context of inflammation during acute viral infection. In contrast, epitopes targeted by inflationary responses are efficiently processed by the standard proteasome and can also be presented to CD8^+^ T cells by latently-infected cells such as endothelial cells when those undergo reactivation of transcription, facilitating continuous inflationary expansion of short-lived T_EM/EFF_ states (49–52). This continuous systemic replenishment may be the basis for the sustained anti-tumor activity we observed for redirected inflationary, but not conventional anti-MCMV responses under APEC therapy. Therefore, careful selection of antiviral responses for APEC targeting will likely help in optimizing therapeutic efficacy. Responses against the HLA A2-restricted NLV epitope of CMV we observed in HNSCC patients are well-known for their inflationary nature and should be prioritized for human APEC design in HLA-A2^+^ expressing patients, who represent half of the population in the US (53). Memory inflation is less well characterized in EBV infection, but has been described to occur under certain conditions (29, 54). We found that compared to CMV NLV-directed responses, CTL responses against the EBV epitopes GLC and CLG were of smaller magnitude in blood, but they were of similar magnitude to NLV-directed responses in tumor tissue of HNSCC patients, where they also comprised a large proportion of GZMB^+^ cells, suggesting parallels to inflationary anti-CMV responses. In contrast, conventional memory responses to past, cleared viral infections, such as Flu, as well as to prior vaccination appear to be increasingly dominated by T_SL_ over time, and may represent suboptimal targets for APEC therapy, even though recruitment to the TME enriches somewhat for the less abundant T_EFF/EM_ states among these responses, as observed for Flu responses in our HNSCC patients.

In further support of the presumed mechanism of action of APECs, we found that APEC-mediated activation of antiviral CTL in the TME and anti-tumor activity required cancer cells to express the relevant protease, and that expression of the same protease by other cells of the TME does not suffice to cleave APECs for activation of antiviral CTL. Our comprehensive, multi-stage screen identified PLAU as a highly active protease in multiple human HNSCC cancer lines, in addition to the murine HNSCC surrogate model TC-1. PLAU is expressed at only very low levels in healthy tissues (42), supporting tumor-focused APEC activation and low risk of systemic toxicity. As reflected by the reduced *in vivo* growth of our PLAU-deficient tumor model, PLAU serves many important functions in cancer cells. These are based both on its direct proteolytic activity and through activation not only of plasmin but also of many other proteases including MMPs, promoting extracellular matrix (ECM) break-down and release of ECM-bound growth factors to support cancer cell survival, migration, epithelial-mesenchymal transition, metastasis, as well as tumor angiogenesis (55, 56). This critical role may restrict the emergence of pre-existing PLAU-deficient cancer clones that could evade targeting by APEC therapy in genetically heterogeneous human tumors. PLAU overexpression has also been described in other epithelial cancers including breast, ovarian, and cervical carcinoma (57, 58). It will be important to assess whether its activity is a general feature of malignant transformation by examining a wider range of cancer types, including of mesenchymal, ectodermal, and hematopoietic origin. If the latter is the case, PLAU may be generally suitable for the activation not only of APECs, but also of other, protease-activated cancer therapeutics, such as antibody-drug conjugates incorporating cytotoxic compounds or biologics modified with masking domains (59).

A general concern with cancer therapies targeting cell surface antigens that are overexpressed by tumor cells but also expressed at lower levels by non-transformed cells in healthy tissues is on-target, off-tumor toxicity, which may mandate treatment discontinuation or lead to morbidity and mortality in patients if unchecked. For instance, severe diarrhea was a dose-limiting side-effect accompanied by mucosal ulceration in cancer patients in a phase I trial of EpCAM-targeting bi-specific T cell engagers (46). EpCAM-targeted CAR therapy produced severe off-tumor toxicity in the lungs in one preclinical study (60), although this observation was not reproduced in two subsequent preclinical mouse models using different CAR constructs (61, 62) or in a human phase I trial (61). In our syngenic cancer model we purposefully chose aged mice, which are most prone to develop immunotherapy-induced toxicity resembling human IrAEs (47, 48), in order to assess potential APEC treatment-induced toxicities in epithelial tissues. Although our sample size does not allow for definitive conclusions, we did not find evidence for treatment-related changes that exceeded changes observed in age-matched, latently MCMV-infected animals. Lack of toxicity may be based on the unique APEC mechanism of action that requires the local synergy of high target antigen expression, relevant protease activity, and the accumulation of antiviral CTL. However, the restricted basolateral expression pattern of EpCAM on healthy epithelium, compared to the more uniform expression on cancer cells (63), may also contribute to a lack of systemic toxicity by potentially restricting antibody access. *In vivo* studies of APECs targeting other tumor antigens, such as EGFR, in relevant preclinical models will provide additional clarity on this important question.

In summary, our observations in a relevant immune-competent preclinical model of HPV^+^ HNSCC indicate that APECs can be a highly effective therapy for immune checkpoint therapy-resistant cancers. Careful selection of targeted antiviral bystander CTL responses appears to be critical to achieve maximal and sustained therapeutic efficacy. The abundance and quality of these responses in the TME of both HPV^+^ and HPV^−^ HNSCC patients can be predicted on an individual patient level through peripheral blood analysis. PLAU-cleavable peptide linkers allow for broad and highly efficient APEC activation by cancer cells. Future APEC design improvement may include optimized peptide conjugation chemistry or genetic peptide-antibody fusions (64) but based on our findings we suggest prioritization of PLAU-cleavable peptide linkers to target inflationary CTL responses against endemic latent viral infections such as CMV and EBV for further development of APEC therapy of HNSCC and other forms of cancer.

## METHODS

### Sex as a biological variable

Our study was open to enrollment to both male and female participants. HNSCC has an estimated 3:1 higher incidence in men than in women, and an even higher imbalance in HPV-mediated HNSCCs, wherein approximately 80% of HPV-mediated HNSCC cases are in men in the United States (U.S.). The distribution of our clinical trial cohort reflects the sex prevalence of HNSCC in the U.S.

### Animals

C57BL/6/J mice were purchased from The Jackson Laboratory. Animals were maintained in specific pathogen-free facilities at the Massachusetts General Hospital (MGH), and all studies were approved and performed in accordance with guidelines and regulations implemented by the MGH Institutional Animal Care and Use Committee (IACUC).

### Cell lines and tumor growth studies

Human tumor cell lines either purchased from ATCC (SCC9, SCC090, RPMI-2650, CaSki), or obtained from Cyril Benes at MGH (DOK, BICR78), the University of Birmingham (VU40T, SCC040), and the University of Michigan (UMSCC10). All lines were grown according to directions provided by ATCC, wherever available, and either in complete Dulbecco’s modified Eagle’s medium (DMEM, Corning), DMEM/F-12 (Thermo Fisher) or RPMI (Corning), supplemented with 10% heat-inactivated fetal bovine serum (FBS, Thermo Fisher), 100 U/ml each of penicillin and streptomycin (Thermo Fisher) and 4 mM of L-glutamine (Life Technologies) at 37 °C and 5% CO2. Cells were tested regularly for both mycoplasma contamination and rodent pathogens; no cell lines tested positive at any point.

The HPV E6/7 antigen-expressing mouse lung epithelial cell line TC-1 was a gift from Dr. T.-C. Wu. TC-1 was engineered to express murine EpCAM and membrane GFP through transduction with lentiviral particles (RC201989L4V, Origene) and two rounds of FACS-based enrichment for GFP^+^ cells to generate TC-1^EpCAM^. PLAU^Null^ TC-1 and PLAU^Null^ TC-1^EpCAM^ were generated through electroporation with custom-designed RNPs composed of PLAU-specific targeting Alt-R^®^ CRISPR-Cas9 crRNAs, Alt-R^®^ CRISPR-Cas9 fluro-tracrRNA (ATTO^™^ 647 or ATTO^™^ 550), and Alt-R™ S.p. Cas9-RFP V3 or Alt-R^™^ S.p. HiFi Cas9 Nuclease V3 (all from IDT). Five different sgRNA sequences mapped to the M.mus GRCm38.p4 genome were designed using multiple online tools from IDT and Synthego and further validated *in silico* in Benchling, CHOPCHOP, and broad GPP databases (**Table S3**). RNPs were pre-assembled before electroporation maximize efficiency. crRNAs were paired and mixed with fluro-tracrRNAs and annealed at 95°C for 5 mins. Cas9 was diluted to 31 μM (5 mg/ml) with Neon buffer T. Annealed crRNA:tracrRNA were then 1:1 mixed (1 μL/test) with diluted Cas9 protein and incubated for 15 mins at room temperature to form the RNP complex. Alt-R® Cas9 electroporation enhancer was diluted at 10.8μM in Neon buffer T. A system of pre-assembled RNP (1 uL), Cas9 enhancer (2 uL) and 5 × 10^5^ cells (in 10 uL) were electroporated at 1600v, 10ms, 3 pulses using the Neon Transfection System (Thermo Fisher). Cells were sorted for gRNA fluorescence the following day to enrich for electroporated cells, as described (65). TC-1^EpCAM^ and PLAU^Null^ TC-1^EpCAM^ cells were grown in DMEM with 10% FCS and used for *in vivo* tumor growth studies in their exponential growth phase. 1.5 × 10^6^ tumor cells in 150 μL PBS were s.c. injected into the flanks of C57BL/6 mice. Animals were randomly allocated to treatment groups when their tumors reached ∼100 mm^3^, resulting in staggered treatment initiation. APECs or vehicle were injected i.p. three times per week at a dose of 25 μg in 100 μl PBS until the end of the experiment. αPD-1 mAB clone RMP1-14 was dosed i.p. at 8 mg/kg in 150 μl PBS three times per week. Tumor volumes were measured on and then every second to third day following the start of treatments and calculated as V = (length x width x 2)/2. Survival analysis was based on the time-point of euthanasia based human endpoints (tumor size or ulceration).

### Antibodies

The chimeric α-human EGFR Cetuximab (Biosimilar) was obtained from SydLabs (Syd Labs Inc.). G8.8 α-murine EpCAM was produced from a hybridoma line obtained from the Developmental Studies Hybridoma Bank (Antibody Registry ID: AB_2098655) and purified via FPLC with ÄKTA pure™ protein purification system (Cytiva) using protein G columns and filtered through glycine at pH 2.7 (peak at 2700 mAU).

The following antibodies were used for flow cytometry analyses of human cells: αCD3 APC Cy7 (clone OKT3, #317342), αCD8α BV785 (clone SKI, #344740), αCD11b FITC (clone ICRF44, # 301330), αCD19 FITC (clone HIB19, #302206) and αCD33 FITC (clone P67.6, #366620), αGZMB AF700 (clone QA16A02, # 372222), αGZMB PE/Cy7 (clone QA16A02, # 372214), (All from BioLegend, USA), and αTCF-1 AF488 (clone C63D9, 6444S), αTCF-1 PE (Clone C63D9, #14456), αTCF-1 AF647 (Clone C63D9, #6709S) (All from Cell Signaling Technologies) Human FcX was from Biolegend.

The following antibodies were used for flow cytometry analyses of mouse cells: αCD3 APC/Cy7 (clone 17A2), αCD8α APC (clone 53-6.7), αCD14 FITC (clone Sa14-2), αCD19 FITC (clone QA17A27), PD-1 BV711 (Clone 29F.1A12), H2K^b^ (Clone AF6-88.5), αTCF-1 PE (clone 7F11A10) (All from Biolegend), EpCAM PE (Clone G8.8) (eBioscience), αuPA(Plau) Biotin (clone OTI5H4) (Origene), and PD-1 BV421 (Clone 29F.1A12) (BD Bioscience).

Streptavidin BUV395, BV421, BV711, BV605, BUV737, APC, PE were all from BD Bioscience.

### Tetramers

HLA peptide monomers were a kind gift from Tetramer Core Facility, Emory University, GA, USA. Six HLA-A*0201 monomers were provided encompassing two epitopes from human cytomegalovirus (NLVPMVATV and VLEETSVML), three epitopes from Epstein-Barr virus (GLCTLVAML, CLGGLLTMV and FLYALALL) and one from Influenza (GILGFVFTL). Three biotinylated MHC I peptide monomers for the MCMV epitopes HGIRNASFI (H-2D^b^, MCMV M45 985-993), SSPPMFRV (H-2K^b^, MCMV m38 316-323), and TVYGFCLL (H-2K^b^, MCMV m139 419-426) were purchased from the NIH Tetramer Core Facility.

Biotinylated monomers containing HCMV, EBV, Flu, or MCMV epitopes (National Institute of Health Tetramer Core Facility, Emory University, USA) were diluted 1:10 in PBS (1–10 μg/ml, optimized per epitope) and tetramerized by sequentially adding streptavidin in 10 steps, each at one-tenth of the total molar ratio every 40–60 min, to maximize tetramer and minimize monomer/dimer formation. Initially, streptavidin binding sites were saturated by excess monomers; by step 9, near-equimolar conditions were reached. A final addition of streptavidin introduced slight excess to prevent unbound monomers. Tetramers were freshly prepared and stored at 4°C for up to two weeks.

### Peptides

Peptides used in this study were synthesized with Fmoc chemistry, isolated by HPLC to >90% purity and validated with mass spectrometry (Genscript). Further modifications were made to contain an *N*-terminal 3-maleimido propionic acid (3-MPA) group to allow chemical conjugation to free thiol groups and a C-terminal amide group.

### Construction of antibody peptide conjugates

Cetuximab or G8.8 antibodies were buffered using 5% v/v 1 M Tris/25 mM EDTA pH 8.0. Disulfide bonds within the antibody were reduced by equimolar addition of 10 mM Tris(2-carboxyethyl)phosphine (TCEP) for 90 min. Peptides containing an *N*-terminal 3-maleimido propionic acid (3-MPA) group were added to the reduced antibody at 3× molarity of TCEP for 60 min before the addition of 10 mM *N*-acetyl cysteine (at 3× molarity of TCEP) to quench free, unbound peptide. Antibody–peptide complexes were then purified from unbound peptide either by protein A purification (GE Healthcare) or ultrafiltration (Millipore). Complexes purified using protein A were eluted from protein A sepharose beads using 0.1 M glycine (pH 2.7) and buffered with 1 M Tris (pH 8.0). Complexes purified using ultrafiltration were diluted to 500 µl in PBS and concentrated to 25 µl. This was repeated a total of eight times before dilution to 250 µl in PBS. Protein quantitation utilized absorbance readings at 280 nm using a NanoDrop 2000 (Thermo Fisher). Purified APECs were diluted to required concentrations using PBS and stored at 4 °C for up to 2 weeks.

### Human HCMV NLV-specific T cell lines

Peripheral blood mononuclear cells from healthy donors were used to culture peptide-specific T-cell lines. Briefly, 10^7^ PBMCs (5 × 10^6^/ml) were stimulated with 10 ng/ml of NLV peptide in RPMI medium supplemented with 10% heat-inactivated FCS (Thermo Fisher), 100 U/ml each of penicillin and streptomycin (Thermo Fisher), 4 mM of L-glutamine (Life Technologies) and 1% human AB serum (Sigma). Following culture of cells for 4 d at 37 °C, 50% of the media was removed and replaced with the same media supplemented with 500 IU/ml of interleukin-2 (IL-2, Peprotech). The cells were fed every 2–3 d and expanded for up to 6 weeks. On day 21, T cells were stained with NLV/HLA-A*0201 tetramers for flow cytometric analysis and only those cell lines with >50% peptide-specific T cells were used in functional assays.

### IFN-γ ELISA-based assay cancer cell protease activity (’CTL assay’)

Tumor cells were incubated with APECs (at 10^−5^ to 1 μg/ml) in Aim-V serum-free medium (Thermo Fisher) for 30 min at room temperature and unbound APECs removed by washing in fresh medium. Tumor cells were then incubated overnight in the presence of viral peptide-specific CD8^+^ T cells in serum-free medium at a 1:10 tumor-to-T cell ratio and supernatants assayed by ELISA for human IFN-γ for T-cell stimulation (Ready-Set-Go Human IFN-γ ELISA kit, eBioscience). Results were z-scored for experiments that were repeated with different batches of tumor or T cells, or APECs to facilitate comparability.

### FRET based screens of cancer cell protease activity (’FRET assay’)

Peptides containing protease cleavage sequences were synthesized containing N-terminal Dabcyl FRET donors and C-terminal EDANS FRET acceptors (Genscript), dissolved in DMSO at 10 mM for storage, and further diluted to 100 μM in PBS and mixed with 10^5^ tumor cells to a final concentration of 10 μM in a total volume of 200 μl in 96-well flat-bottom plates. Plates were incubated overnight at 37 °C in a fluorescence plate reader (Bio-Tek), with fluorescence readings taken every 5 min at excitation and emission wavelengths of 340 and 485 nm, respectively. Cleavage efficiency was calculated as the slope of fluorescence increase (RFU/min) during the first 60 min post-exposure, determined by linear regression.

### Analysis of antiviral CTL responses in human and murine tissues

De-identified human blood and tumor tissues were obtained from HNSCC patients through the Pathology Lab at Massachusetts General Hospital/Massachusetts Eye and Ear Infirmary. All protocols were approved by the Massachusetts General Hospital Institutional Review Board and performed in accordance with Federal Law. The study was conducted according to the provisions of the Declaration of Helsinki, and all patients signed and provided their informed consent before sample collection.

Peripheral blood mononuclear cells were isolated using Ficoll-Paque Plus (GE Healthcare) density gradient centrifugation according to the manufacturer’s instructions. For long-term storage, PBMCs were resuspended in FBS with 10% DMSO and stored in liquid nitrogen at a density of 1–5 × 10^7^ cells/ml.

Human tumor samples were processed into single-cell suspensions using either enzymatic or mechanical digestion. For mechanical dissociation, the tumor was placed into RPMI 1640 in a small tube and dissected into small fragements (<1 mm) using fine forceps. Using a wide bore pipette tip (ART tips, ThermoFisher, USA), the fragments were transferred to a pre-wet filter (50 µm) and homogenized using the end of a 1 ml syringe plunger. The filter was then washed in RPMI 1640 to dislodge all single cells. For enzymatic digestion, tumors were placed in 460 μl RPMI 1640 supplemented with 42 μl Enzyme H, 21 μl Enzyme R and 5 μl Enzyme A (Tumor Dissociation Kit, human, Miltenyi Biotec, Germany). The tumor was dissected into small (<1 mm) fragments and 460 μl of RPMI 1640 added before incubation at 37 °C on a shaker for 20-30 minutes. The cell suspension was then filtered through a pre-wet 50 µm CellTrics filter (Sysmex, IL, USA) and the remaining pieces of tumor tissue homogenized using the back end of a 1 ml syringe plunger. The filter was then washed in RPMI 1640 to dislodge all remaining single cells.

Mouse tumor samples were processed using a combination of mechanical and enzymatic digestion to obtain single-cell suspensions. Tumor tissues were placed in a 6 cm Petri dish containing room temperature (RT) RPMI 1640 and minced into small fragments (approximately 1–2 mm³) using a scalpel. For enzymatic digestion, digestion medium was prepared in RPMI 1640 supplemented with 42 μL Enzyme H, 21 μL Enzyme R, and 5 μL Enzyme A (Tumor Dissociation Kit, mouse, Miltenyi Biotec, Germany). Alternatively, a digestion cocktail consisting of DNase I (0.1 mg/mL, Roche) and Collagenase IV (1 mg/mL, Worthington) was used. Minced tumor fragments were transferred into pre-warmed digestion medium in a glass jar and incubated for 30 min to 1.5 h (depending on tumor size and density) at 37°C on a shaker.

Following enzymatic digestion, the cell suspension was passed through a 40 μm nylon mesh strainer into a 50 mL conical tube to remove undigested debris. The strainer was rinsed with additional RPMI 1640 supplemented with 2% FBS to maximize cell recovery. The filtrate was centrifuged at 400 × g for 10 min at room temperature, and the supernatant was carefully aspirated.

In some cases, the cell pellet was resuspended in 1-10 ml (5–10 × the pellet volume) ACK (Ammonium-Chloride-Potassium) lysis buffer and incubated at RT for 5 min to lyse red blood cells. The reaction was quenched by adding RPMI 1640 with 2% FBS, followed by centrifugation at 400 × g for 10 min.

### RNA-Seq Data Mining from TCGA HNSC Data

Transcriptomes from 520 HNSCC tumors and 44 matched healthy tissues were obtained via the Broad Institute Firehose pipeline (stddata 2016_01_28 run). Gene expression levels were normalized using RSEM (RNA-Seq by Expectation-Maximization), which quantifies transcript abundance while accounting for transcript length and sequencing depth biases. RSEM was selected over TPM/FPKM due to its superior handling of isoform-level expression and compatibility with TCGA’s alignment pipeline. Differential expression analysis prioritized proteases with >5-fold overexpression in tumors (median RSEM-tumor/median RSEM-healthy ratio).

### Murine cytomegalovirus infection

Stocks of the K181 strain of MCMV (GenBank AM886412.1, originally provided by E. Mocarski) were produced from M2-10B4 cells (ATCC, CRL-1972) and quantified by plaque assay as previously described (66). 7-10 weeks old C57BL/6 mice were inoculated intraperitoneally with 10^5^ PFU in 100 μl of PBS.

### Generation of viral epitope-specific mouse CTL from peripheral blood samples

Peripheral blood was collected at repeated time points from MCMV-infected mice (e.g. via submandibular venipuncture) and pooled for each experiment. Peripheral blood mononuclear cells (PBMC) were isolated, washed twice, and resuspended in complete RPMI supplemented with 10% heat-inactivated FBS, 1% penicillin/streptomycin, 2 mM L-glutamine, 50 µM 2-mercaptoethanol, and recombinant murine IL-2 (50–100 U/mL). Bulk PBMC (without prior subset isolation) were stimulated in vitro with the cognate H-2K^b^–restricted TVY peptide (1 µM) and cultured at 37 °C, 5% CO₂ for 14 days, with IL-2 and medium replenished every 2–3 days as needed.

### Mouse IFN**-γ** ELISA Assay

For murine CTL activation assays, target cells (TC-1^EpCAM^ tumor cells or TC-1^EpCAM^ ^PlauNull^, 1 × 10^5^ cells/well) were plated in 96-well flat-bottom plates in RPMI 1640 supplemented with 10% heat-inactivated FCS, 100 U/ml penicillin, 100 U/ml streptomycin, and 2 mM L-glutamine. Where indicated, target cells were incubated with murine APECs TVY-PLAU8 or TVY-MMP13-8 (1 μg/ml) for 30 min at room temperature, followed by two washes in fresh medium to remove unbound APECs. Effector cells consisted of MCMV peptide-specific CD8⁺ T cells generated from peripheral blood as described above, which were added at a 1:10 target-to-effector ratio and co-cultured for 16–20 h at 37 °C in a humidified 5% CO₂ incubator. Supernatants were harvested and analyzed for murine IFN-γ using a Mouse IFN-γ Ready-Set-Go ELISA kit (eBioscience) according to the manufacturer’s instructions.

### Flow cytometry

Cells were washed in PBS with 2% FCS and placed in 5-ml FACS tubes (Corning Falcon). Tetramer staining was performed prior to antibody staining. Samples were incubated with tetramers for 30 min at room temperature, followed by two washes with 3 ml PBS before subsequent incubation with surface antibodies.

Cells were incubated for 30 min at room temperature in the dark, and excess antibody washed off. Where necessary, a secondary anti-mouse IgG PE (Thermo Fisher) was added to the cells and excess antibody removed by washing.

In some cases, intracellular antigens were stained using the Foxp3/Transcription Factor Staining Buffer Set (ThermoFisher, USA). The concentrate was dilute 4x in Fixation/Permeabilization diluent and the Permeabilization buffer was diluted 10x in distilled water. Cells were resuspended in 1ml of the diluted Fixation /Permeabilization buffer and incubated on ice for 60 minutes on ice, vortexing every 15 minutes. Cells were then washed using the Permeabiliztion buffer, pelleted at 400 x g for 4 minutes and the supernatant carefully discarded. Intracellular antibodies against TCF/-1 (Cell Signaling Technologies, USA) and Granzyme B (BioLegend, USA) were added at 1/25 dilution for 60 minutes on ice. Cells were then washed using the Permeabiliztion buffer, pelleted at 400 x g for 4 minutes and the supernatant carefully discarded. Labelled cells were analyzed using the Fortessa X-20 (Becton Dickinson, USA). Flow cytometry data was analyzed using FlowJo X software (FlowJo, USA).

### Histopathology

Tissues were harvested following intracardiac perfusion with 10% buffered formalin of euthanized animals, followed by further fixation for 48 h, trimmed, placed into microcassettes, and embedded in paraffin wax. 5 μm sections were stained with haematoxylin and eosin according to standard procedures, examined by a board-certified veterinary pathologist in a blinded manner, and scored based on the extent of lymphocytic infiltrates.

### Statistical analyses

All statistical analyses were performed using GraphPad Prism 10. Unless otherwise noted in the figure legends, data are shown as mean ± SEM. For multiple group comparisons, either one-way or two-way ANOVA were used to determine statistically significant differences between groups, and Tukey’s multiple comparison test adjusted P value < 0.033 was used to indicate a significant difference. Differences between two groups were determined by two-sided, unpaired t-test, and P < 0.05 used to indicate a significant difference. Survival curves were analyzed by a log-rank test.

## Author contributions

Conceptualization: MC, DGM, SIP, TRM

Methodology: MC, DGM, CMS, TRM

Investigation: YIS, DGM, LMA, AJM, PMS, WCF

Visualization: YIS, DGM, TRM

Funding acquisition: MC, SIP, AH, TRM

Project administration: SIP, TRM

Supervision: DGM, TRM

Writing – original draft: YIS, DGM, TRM

Writing – review & editing: YIS, TRM

## Acknowledgments

We would like to thank members of the Mempel laboratory at MGH for helpful discussions. This study was funded by National Institutes of Health grant P01 CA240239 (TRM, SIP, DGM, YIS), R01 CA278212 (SIP), R01 AI123349 (TRM) and R35 CA232103 (TRM, YIS)

## Competing interests

MC and DGM are inventors on a patent (US9402916B2) related to the design and use of APECs for cancer therapy. SIP has received consultancy payments from Abbvie, AstraZeneca/MedImmune, CUE Biopharma, Fusion Pharmaceuticals, G1 Therapeutics, Incendia, Inovio, MSD/Merck, Newlink Genetics, Oncolys Biopharma, Parthenon Therapeutics, Recurrent Respiratory Papillomatosis Foundation, Replimune, Scopus Biopharma, Sensei Biotherapeutics, and Umoja; and research grants from Abbvie, AstraZeneca/MedImmune, CUE Biopharma, Eisai, Merck, Recurrent Respiratory Papillomatosis Foundation, Sensei Biotherapeutics, and Tesaro, outside of the submitted work.

## Data and materials availability

All data are available in the main manuscript or the supplementary materials.

