## Supplementary material for "Antibody peptide epitope conjugates for αPD-1 therapy-resistant head and neck squamous cell carcinoma": Figs. S1-3 and Tables S1-3

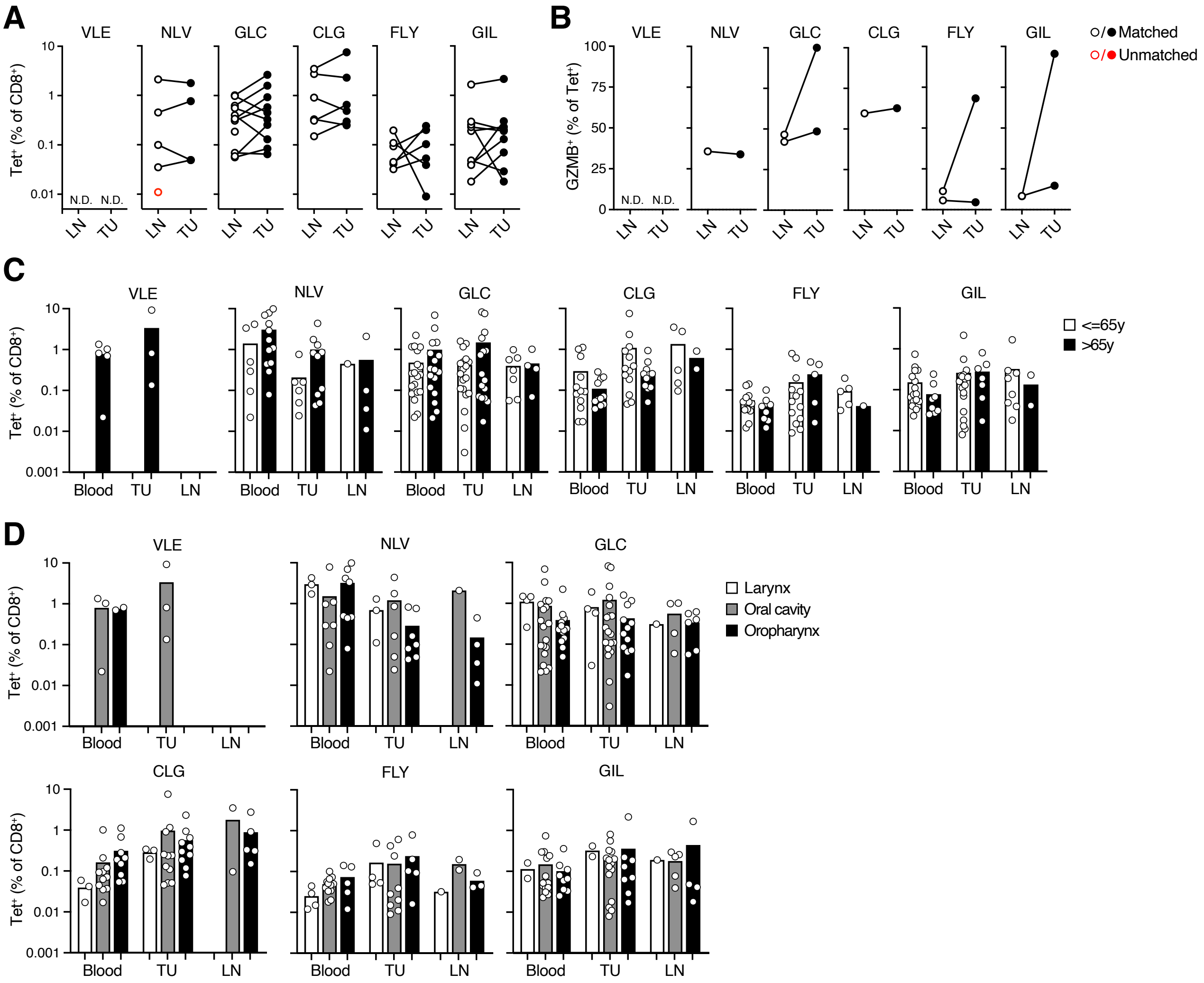
A
VLE
NLV
GLC
CLG
FLY
GIL
Tet⁺ (% of CD8⁺)
N.D.
LN
TU
B
GZMB⁺ (% of Tet⁺)
○/● Matched
○/● Unmatched
C
Blood
<=65y
>65y
D
Larynx
Oral cavity
Oropharynx

**Figure S1**

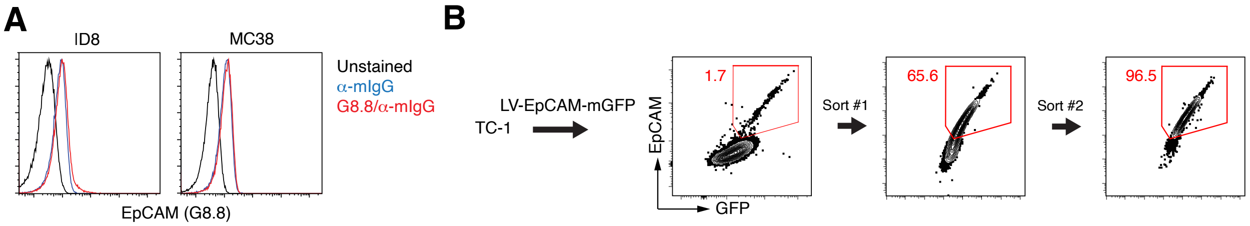

A
ID8
MC38
Unstained
α-mIgG
G8.8/α-mIgG
EpCAM (G8.8)
B
LV-EpCAM-mGFP
TC-1
EpCAM
GFP
1.7
Sort #1
65.6
Sort #2
96.5


**Figure S2**

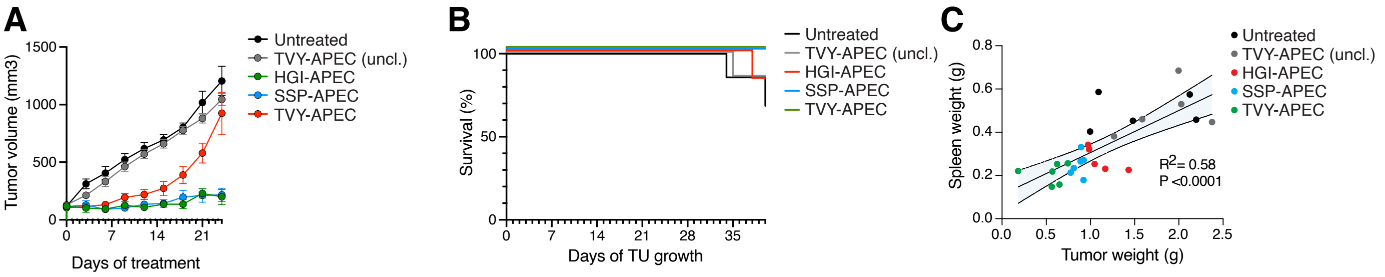
A
Tumor volume (mm3)
Days of treatment
Untreated
TVY-APEC (uncl.)
HGI-APEC
SSP-APEC
TVY-APEC
B
Survival (%)
Days of TU growth
Untreated
TVY-APEC (uncl.)
HGI-APEC
SSP-APEC
TVY-APEC
C
Spleen weight (g)
Tumor weight (g)
$R^2$= 0.58
P <0.0001
Untreated
TVY-APEC (uncl.)
HGI-APEC
SSP-APEC
TVY-APEC

**Figure S3**

| Patient ID | Age | Gender | Tumor Site | HPV status | Flu vaccination |
|---|---|---|---|---|---|
| P01.P2.10 | 60 | M | Larynx | Positive | - |
| P01.P2.26 | 65 | F | Larynx | Negative | N |
| P01.P2.35 | 69 | M | Larynx | Negative | Y - 10/30/20 |
| P01.P2.06 | 81 | M | Larynx | Negative | - |
| P01.P2.56 | 81 | M | Larynx | Negative | Y - 10/1/20 |
| P01.P2.42 | 72 | F | Larynx | Negative | N |
| P01.P2.11 | 52 | M | Oral cavity | Negative | - |
| P01.P2.14 | 56 | F | Oral cavity | Negative | - |
| P01.P2.50 | 59 | M | Oral cavity | Negative | Y - 3/11/21 |
| P01.P2.28 | 65 | M | Oral cavity | Negative | Y - 10/16/20 |
| P01.P2.43 | 66 | M | Oral cavity | Negative | Y - 12/3/20 |
| P01.P2.08 | 72 | F | Oral cavity | Negative | - |
| P01.P2.22 | 67 | M | Oral cavity | Negative | Y - 9/25/20 |
| P01.P2.52 | 81 | M | Oral cavity | Negative | Y - 9/17/20 |
| P01.P2.32 | 92 | F | Oral cavity | Negative | Y |
| P01.P2.20 | 50 | M | Oral cavity | Negative | Y |
| P01.P2.39 | 52 | M | Oral cavity | Negative | Y - 11/5/20 |
| P01.P2.05 | 55 | M | Oral cavity | Negative | - |
| P01.P2.41 | 55 | M | Oral cavity | Negative | Y - 9/23/20 |
| P01.P2.25 | 56 | M | Oral cavity | Negative | Y |
| P01.P2.47 | 62 | M | Oral cavity | Negative | Y - 9/18/20 |
| P01.P2.15 | 63 | M | Oral cavity | Negative | Y - 9/22/20 |
| P01.P2.53 | 65 | M | Oral cavity | Positive | Y - 10/1/20 |
| P01.P2.31 | 67 | F | Oral cavity | Negative | Y - 11/14/20 |
| P01.P2.57 | 67 | M | Oral cavity | Negative | Y - 12/22/20 |
| P01.P2.46 | 69 | F | Oral cavity | Negative | Y |
| P01.P2.36 | 70 | M | Oral cavity | Negative | Y - 8/31/20 |
| P01.P2.51 | 79 | F | Oral cavity | Negative | Y - 7/28/21 |
| P01.P2.23 | 53 | M | Oropharynx | Positive | Y - 12/29/20 |
| P01.P2.34 | 60 | F | Oropharynx | Negative | - |
| P01.P2.38 | 67 | M | Oropharynx | Positive | Y - 9/18/20 |
| P01.P2.18 | 71 | M | Oropharynx | Positive | Y - 9/28/20 |
| P01.P2.12 | 72 | M | Oropharynx | Positive | - |
| P01.P2.29 | 49 | M | Oropharynx | Positive | Y |
| P01.P2.16 | 55 | M | Oropharynx | Positive | N |
| P01.P2.07 | 56 | M | Oropharynx | Positive | - |
| P01.P2.17 | 58 | M | Oropharynx | Positive | Y - 10/25/20 |
| P01.P2.33 | 66 | M | Oropharynx | Positive | Y - 11/18/20 |
| P01.P2.37 | 66 | M | Oropharynx | Positive | Y - 9/16/20 |
| P01.P2.58 | 68 | M | Oropharynx | Positive | Y - 10/20/20 |
| P01.P2.40 | 71 | M | Oropharynx | Positive | Y - 10/21/20 |
| P01.P2.09 | 72 | M | Oropharynx | Positive | - |
| P01.P2.19 | 74 | M | Oropharynx | Positive | Y - 11/12/20 |

#### A. Surfacome (4199)

| No. | Gene Name |
|---|---|
| 1 | MARC1 |
| 2 | MARCH1 |
| 3 | MARC2 |
| 4 | MARCH3 |
| 5 | MARCH4 |
| 6 | MARCH5 |
| 7 | MARCH9 |
| 8 | MARCH11 |
| 9 | SEPT2 |
| 10 | SEPT6 |
| 11 | SEPT7 |
| 12 | SEPT12 |
| 13 | A4GALT |
| 14 | A4GNT |
| 15 | AADAC |
| 16 | AADACL4 |
| 17 | AAMP |
| 18 | AATK |
| 19 | ABCA10 |
| 20 | ABCA12 |
| 21 | ABCA13 |
| 22 | ABCA3 |
| 23 | ABCA6 |
| 24 | ABCA7 |
| 25 | ABCA9 |
| 26 | ABCB1 |
| 27 | ABCB11 |
| 28 | ABCB4 |
| 29 | ABCB6 |
| 30 | ABCB9 |
| 31 | ABCC1 |
| 32 | ABCC10 |
| 33 | ABCC11 |
| 34 | ABCC13 |
| 35 | ABCC2 |
| 36 | ABCC3 |
| 37 | ABCC6 |
| 38 | ABCC8 |
| 39 | ABCC9 |
| 40 | ABCD1 |
| 41 | ABCD2 |
| 42 | ABCD3 |
| 43 | ABCD4 |
| 44 | ABCG1 |
| 45 | ABCG2 |
| 46 | ABCG5 |
| 47 | ABCG8 |
| 48 | ABHD1 |
| 49 | ABHD12 |
| 50 | ABHD13 |
| 51 | ABHD14A |
| 52 | ABHD16A |
| 53 | ABHD2 |
| 54 | ABHD3 |
| 55 | ABHD6 |
| 56 | ABO |
| 57 | ACBD5 |
| 58 | ACE |
| 59 | ACE2 |
| 60 | ACHE |
| 61 | ACP2 |
| 62 | ACPT |
| 63 | ACSL1 |
| 64 | ACSL3 |
| 65 | ACSL4 |
| 66 | ACSL5 |
| 67 | ACSL6 |
| 68 | ACVR1 |
| 69 | ACVR1B |
| 70 | ACVR2A |
| 71 | ACVR2B |
| 72 | ACVRL1 |
| 73 | ADA |
| 74 | ADAM10 |
| 75 | ADAM11 |
| 76 | ADAM12 |
| 77 | ADAM15 |
| 78 | ADAM17 |
| 79 | ADAM18 |
| 80 | ADAM19 |
| 81 | ADAM2 |

| | |
|---|---|
| 82 | ADAM20 |
| 83 | ADAM21 |
| 84 | ADAM22 |
| 85 | ADAM23 |
| 86 | ADAM28 |
| 87 | ADAM29 |
| 88 | ADAM30 |
| 89 | ADAM32 |
| 90 | ADAM33 |
| 91 | ADAM7 |
| 92 | ADAM8 |
| 93 | ADAM9 |
| 94 | ADAMTS13 |
| 95 | ADAMTS7 |
| 96 | ADCK2 |
| 97 | ADCK4 |
| 98 | ADCK5 |
| 99 | ADCY1 |
| 100 | ADCY3 |
| 101 | ADCY5 |
| 102 | ADCY8 |
| 103 | ADCY9 |
| 104 | ADCYAP1R1 |
| 105 | ADIPOQ |
| 106 | ADIPOR1 |
| 107 | ADIPOR2 |
| 108 | ADORA1 |
| 109 | ADORA2B |
| 110 | ADORA3 |
| 111 | ADRA1A |
| 112 | ADRA1B |
| 113 | ADRA1D |
| 114 | ADRB1 |
| 115 | ADRB2 |
| 116 | ADRB3 |
| 117 | AFG3L2 |
| 118 | AGER |
| 119 | AGPAT1 |
| 120 | AGPAT3 |
| 121 | AGPAT4 |
| 122 | AGPAT5 |
| 123 | AGPAT6 |
| 124 | AGPAT9 |
| 125 | AGRN |
| 126 | AGTR1 |
| 127 | AGTR2 |
| 128 | AGTRAP |
| 129 | AIFM2 |
| 130 | AIG1 |
| 131 | AIM1 |
| 132 | AIMP1 |
| 133 | AJAP1 |
| 134 | ALCAM |
| 135 | ALDH3A2 |
| 136 | ALG1 |
| 137 | ALG10 |
| 138 | ALG10B |
| 139 | ALG11 |
| 140 | ALG14 |
| 141 | ALG5 |
| 142 | ALG6 |
| 143 | ALK |
| 144 | ALPP |
| 145 | AMBP |
| 146 | AMELX |
| 147 | AMHR2 |
| 148 | AMICA1 |
| 149 | AMIGO1 |
| 150 | AMIGO2 |
| 151 | AMN |
| 152 | AMOT |
| 153 | ANGPTL3 |
| 154 | ANK3 |
| 155 | ANKAR |
| 156 | ANKH |
| 157 | ANKLE2 |
| 158 | ANKRD46 |
| 159 | ANO1 |
| 160 | ANO10 |
| 161 | ANO2 |
| 162 | ANO3 |
| 163 | ANO5 |
| 164 | ANO6 |

| | |
|---|---|
| 165 | ANO7 |
| 166 | ANO9 |
| 167 | ANPEP |
| 168 | ANTXR1 |
| 169 | ANTXR2 |
| 170 | ANXA1 |
| 171 | ANXA2 |
| 172 | ANXA4 |
| 173 | ANXA5 |
| 174 | ANXA9 |
| 175 | AOC3 |
| 176 | APH1B |
| 177 | APLNR |
| 178 | APLP1 |
| 179 | APLP2 |
| 180 | APMAP |
| 181 | APOA1 |
| 182 | APOE |
| 183 | APOH |
| 184 | APOOL |
| 185 | APP |
| 186 | AQP1 |
| 187 | AQP10 |
| 188 | AQP11 |
| 189 | AQP12A |
| 190 | AQP12B |
| 191 | AQP2 |
| 192 | AQP3 |
| 193 | AQP4 |
| 194 | AQP5 |
| 195 | AQP6 |
| 196 | AQP7 |
| 197 | AQP7P3 |
| 198 | AQP8 |
| 199 | AQP9 |
| 200 | AQPEP |
| 201 | AREG |
| 202 | ARF6 |
| 203 | ARIH2OS |
| 204 | ARL6IP1 |
| 205 | ARL6IP5 |
| 206 | ARL6IP6 |
| 207 | ARMC10 |
| 208 | ARMCX1 |
| 209 | ARMCX2 |
| 210 | ARMCX3 |
| 211 | ARMCX4 |
| 212 | ARSB |
| 213 | ART1 |
| 214 | ASAH2 |
| 215 | ASGR1 |
| 216 | ASGR2 |
| 217 | ASIC1 |
| 218 | ASIC2 |
| 219 | ASIC3 |
| 220 | ASIC4 |
| 221 | ASIC5 |
| 222 | ASPH |
| 223 | ASPHD1 |
| 224 | ASPHD2 |
| 225 | ASPRV1 |
| 226 | ASTN2 |
| 227 | ATAD3A |
| 228 | ATF6 |
| 229 | ATF6B |
| 230 | ATG9A |
| 231 | ATG9B |
| 232 | ATL1 |
| 233 | ATL3 |
| 234 | ATP10A |
| 235 | ATP10B |
| 236 | ATP11A |
| 237 | ATP11B |
| 238 | ATP11C |
| 239 | ATP13A2 |
| 240 | ATP13A4 |
| 241 | ATP1A1 |
| 242 | ATP1A2 |
| 243 | ATP1A4 |
| 244 | ATP1B1 |
| 245 | ATP1B2 |
| 246 | ATP1B3 |
| 247 | ATP1B4 |

| | |
|---|---|
| 248 | ATP2B1 |
| 249 | ATP2B3 |
| 250 | ATP2C1 |
| 251 | ATP2C2 |
| 252 | ATP4A |
| 253 | ATP4B |
| 254 | ATP5B |
| 255 | ATP5G1 |
| 256 | ATP5G2 |
| 257 | ATP5G3 |
| 258 | ATP5J2 |
| 259 | ATP6AP1 |
| 260 | ATP6AP2 |
| 261 | ATP6V0A1 |
| 262 | ATP6V0A2 |
| 263 | ATP6V0A4 |
| 264 | ATP6V0B |
| 265 | ATP6V0C |
| 266 | ATP6V0E1 |
| 267 | ATP6V0E2 |
| 268 | ATP8A1 |
| 269 | ATP8A2 |
| 270 | ATP8B3 |
| 271 | ATP8B4 |
| 272 | ATP9A |
| 273 | ATP9B |
| 274 | ATPIF1 |
| 275 | ATRAID |
| 276 | ATRN |
| 277 | AUP1 |
| 278 | AVL9 |
| 279 | AVPR1A |
| 280 | AVPR1B |
| 281 | AWAT1 |
| 282 | AWAT2 |
| 283 | AXL |
| 284 | B2M |
| 285 | B3GALNT1 |
| 286 | B3GALNT2 |
| 287 | B3GALT1 |
| 288 | B3GALT2 |
| 289 | B3GALT4 |
| 290 | B3GALT5 |
| 291 | B3GALT6 |
| 292 | B3GALTL |
| 293 | B3GAT1 |
| 294 | B3GAT2 |
| 295 | B3GAT3 |
| 296 | B3GNT1 |
| 297 | B3GNT2 |
| 298 | B3GNT3 |
| 299 | B3GNT4 |
| 300 | B3GNT5 |
| 301 | B3GNT6 |
| 302 | B3GNT7 |
| 303 | B3GNT9 |
| 304 | B4GALNT1 |
| 305 | B4GALNT2 |
| 306 | B4GALNT3 |
| 307 | B4GALNT4 |
| 308 | B4GALT1 |
| 309 | B4GALT2 |
| 310 | B4GALT3 |
| 311 | B4GALT4 |
| 312 | B4GALT5 |
| 313 | B4GALT6 |
| 314 | B4GALT7 |
| 315 | BACE1 |
| 316 | BAI3 |
| 317 | BAK1 |
| 318 | BAMBI |
| 319 | BAX |
| 320 | BCAM |
| 321 | BCAP31 |
| 322 | BCL2 |
| 323 | BCL2L1 |
| 324 | BCL2L10 |
| 325 | BCL2L13 |
| 326 | BCS1L |
| 327 | BDKRB1 |
| 328 | BDKRB2 |
| 329 | BEAN1 |
| 330 | BEST1 |

| | |
|---|---|
| 331 | BEST2 |
| 332 | BEST3 |
| 333 | BEST4 |
| 334 | BET1 |
| 335 | BET1L |
| 336 | BFAR |
| 337 | BGN |
| 338 | BIK |
| 339 | BLCAP |
| 340 | BMP10 |
| 341 | BMP2 |
| 342 | BMPR1A |
| 343 | BMPR1B |
| 344 | BMPR2 |
| 345 | BNIP1 |
| 346 | BNIP3 |
| 347 | BNIP3L |
| 348 | BOC |
| 349 | BOK |
| 350 | BRAP |
| 351 | BRI3 |
| 352 | BRICD5 |
| 353 | BRS3 |
| 354 | BSG |
| 355 | BSND |
| 356 | BSPH1 |
| 357 | BST2 |
| 358 | BTBD11 |
| 359 | BTC |
| 360 | BTLA |
| 361 | BTN1A1 |
| 362 | BTN2A1 |
| 363 | BTN2A2 |
| 364 | BTN2A3P |
| 365 | BTN3A1 |
| 366 | BTN3A2 |
| 367 | BTN3A3 |
| 368 | BTNL2 |
| 369 | BTNL3 |
| 370 | BTNL8 |
| 371 | BTNL9 |
| 372 | BVES |
| 373 | C10orf105 |
| 374 | C10orf111 |
| 375 | C10orf128 |
| 376 | C10orf35 |
| 377 | C10orf54 |
| 378 | C10orf76 |
| 379 | C11orf24 |
| 380 | C11orf34 |
| 381 | C11orf75 |
| 382 | C11orf87 |
| 383 | C12orf69 |
| 384 | C12orf70 |
| 385 | C14orf1 |
| 386 | C14orf101 |
| 387 | C14orf2 |
| 388 | C14orf37 |
| 389 | C15orf27 |
| 390 | C16orf54 |
| 391 | C16orf58 |
| 392 | C16orf62 |
| 393 | C16orf92 |
| 394 | C17orf62 |
| 395 | C17orf74 |
| 396 | C17orf78 |
| 397 | C17orf80 |
| 398 | C19orf12 |
| 399 | C19orf18 |
| 400 | C19orf24 |
| 401 | C19orf26 |
| 402 | C19orf38 |
| 403 | C19orf59 |
| 404 | C19orf77 |
| 405 | C1GALT1 |
| 406 | C1GALT1C1 |
| 407 | C1orf159 |
| 408 | C1orf162 |
| 409 | C1orf204 |
| 410 | C1orf210 |
| 411 | C1orf233 |
| 412 | C1orf27 |
| 413 | C1orf43 |

| | |
|---|---|
| 414 | C1orf85 |
| 415 | C1orf95 |
| 416 | C1QBP |
| 417 | C20orf141 |
| 418 | C22orf24 |
| 419 | C22orf32 |
| 420 | C2CD2L |
| 421 | C3orf17 |
| 422 | C3orf18 |
| 423 | C3orf20 |
| 424 | C3orf33 |
| 425 | C3orf35 |
| 426 | C3orf43 |
| 427 | C3orf52 |
| 428 | C4orf21 |
| 429 | C4orf3 |
| 430 | C4orf32 |
| 431 | C4orf52 |
| 432 | C5AR1 |
| 433 | C5AR2 |
| 434 | C5orf15 |
| 435 | C5orf28 |
| 436 | C5orf4 |
| 437 | C5orf60 |
| 438 | C6orf10 |
| 439 | C6orf70 |
| 440 | C6orf89 |
| 441 | C7orf45 |
| 442 | C7orf53 |
| 443 | C9 |
| 444 | C9orf123 |
| 445 | C9orf135 |
| 446 | C9orf174 |
| 447 | C9orf57 |
| 448 | C9orf69 |
| 449 | C9orf91 |
| 450 | CA12 |
| 451 | CA14 |
| 452 | CA4 |
| 453 | CA9 |
| 454 | CABP7 |
| 455 | CACFD1 |
| 456 | CACHD1 |
| 457 | CACNA1A |
| 458 | CACNA1B |
| 459 | CACNA1C |
| 460 | CACNA1D |
| 461 | CACNA1E |
| 462 | CACNA1F |
| 463 | CACNA1G |
| 464 | CACNA1H |
| 465 | CACNA1I |
| 466 | CACNA2D1 |
| 467 | CACNA2D3 |
| 468 | CACNA2D4 |
| 469 | CACNG1 |
| 470 | CACNG2 |
| 471 | CACNG3 |
| 472 | CACNG4 |
| 473 | CACNG5 |
| 474 | CACNG6 |
| 475 | CACNG8 |
| 476 | CADM1 |
| 477 | CADM2 |
| 478 | CADM3 |
| 479 | CADM4 |
| 480 | CALHM1 |
| 481 | CALHM3 |
| 482 | CALN1 |
| 483 | CALR |
| 484 | CALY |
| 485 | CANT1 |
| 486 | CANX |
| 487 | CAPN5 |
| 488 | CASC4 |
| 489 | CASR |
| 490 | CATSPER1 |
| 491 | CATSPER3 |
| 492 | CATSPER4 |
| 493 | CATSPERD |
| 494 | CATSPERG |
| 495 | CAV3 |
| 496 | CCBP2 |

| | |
|---|---|
| 497 | CCDC108 |
| 498 | CCDC109B |
| 499 | CCDC136 |
| 500 | CCDC155 |
| 501 | CCDC167 |
| 502 | CCDC47 |
| 503 | CCDC51 |
| 504 | CCDC90A |
| 505 | CCDC90B |
| 506 | CCKAR |
| 507 | CCKBR |
| 508 | CCPG1 |
| 509 | CCR1 |
| 510 | CCR10 |
| 511 | CCR2 |
| 512 | CCR3 |
| 513 | CCR5 |
| 514 | CCR6 |
| 515 | CCR7 |
| 516 | CCR8 |
| 517 | CCR9 |
| 518 | CCRL1 |
| 519 | CCRL2 |
| 520 | CD101 |
| 521 | CD109 |
| 522 | CD14 |
| 523 | CD151 |
| 524 | CD163 |
| 525 | CD163L1 |
| 526 | CD164 |
| 527 | CD164L2 |
| 528 | CD180 |
| 529 | CD19 |
| 530 | CD1A |
| 531 | CD1B |
| 532 | CD1C |
| 533 | CD1D |
| 534 | CD1E |
| 535 | CD2 |
| 536 | CD200 |
| 537 | CD200R1 |
| 538 | CD200R1L |
| 539 | CD207 |
| 540 | CD209 |
| 541 | CD22 |
| 542 | CD226 |
| 543 | CD24 |
| 544 | CD244 |
| 545 | CD247 |
| 546 | CD248 |
| 547 | CD27 |
| 548 | CD274 |
| 549 | CD276 |
| 550 | CD28 |
| 551 | CD300A |
| 552 | CD300C |
| 553 | CD300E |
| 554 | CD300LB |
| 555 | CD300LD |
| 556 | CD300LF |
| 557 | CD300LG |
| 558 | CD302 |
| 559 | CD320 |
| 560 | CD33 |
| 561 | CD34 |
| 562 | CD36 |
| 563 | CD37 |
| 564 | CD38 |
| 565 | CD3D |
| 566 | CD3E |
| 567 | CD3G |
| 568 | CD4 |
| 569 | CD40 |
| 570 | CD40LG |
| 571 | CD44 |
| 572 | CD46 |
| 573 | CD47 |
| 574 | CD5 |
| 575 | CD53 |
| 576 | CD55 |
| 577 | CD58 |
| 578 | CD59 |
| 579 | CD6 |

| | |
|---|---|
| 580 | CD63 |
| 581 | CD68 |
| 582 | CD69 |
| 583 | CD7 |
| 584 | CD70 |
| 585 | CD72 |
| 586 | CD74 |
| 587 | CD79A |
| 588 | CD79B |
| 589 | CD80 |
| 590 | CD81 |
| 591 | CD82 |
| 592 | CD83 |
| 593 | CD84 |
| 594 | CD86 |
| 595 | CD8A |
| 596 | CD8B |
| 597 | CD9 |
| 598 | CD93 |
| 599 | CD96 |
| 600 | CD97 |
| 601 | CD99 |
| 602 | CD99L2 |
| 603 | CDAN1 |
| 604 | CDC14C |
| 605 | CDCP1 |
| 606 | CDH10 |
| 607 | CDH11 |
| 608 | CDH12 |
| 609 | CDH13 |
| 610 | CDH15 |
| 611 | CDH16 |
| 612 | CDH17 |
| 613 | CDH18 |
| 614 | CDH19 |
| 615 | CDH2 |
| 616 | CDH20 |
| 617 | CDH22 |
| 618 | CDH23 |
| 619 | CDH24 |
| 620 | CDH26 |
| 621 | CDH3 |
| 622 | CDH4 |
| 623 | CDH5 |
| 624 | CDH6 |
| 625 | CDH7 |
| 626 | CDH9 |
| 627 | CDHR2 |
| 628 | CDHR3 |
| 629 | CDHR4 |
| 630 | CDHR5 |
| 631 | CDIPT |
| 632 | CDKAL1 |
| 633 | CDON |
| 634 | CEACAM1 |
| 635 | CEACAM19 |
| 636 | CEACAM20 |
| 637 | CEACAM21 |
| 638 | CEACAM3 |
| 639 | CEACAM4 |
| 640 | CEACAM5 |
| 641 | CEND1 |
| 642 | CEP55 |
| 643 | CERS1 |
| 644 | CERS3 |
| 645 | CERS4 |
| 646 | CERS5 |
| 647 | CERS6 |
| 648 | CFTR |
| 649 | CHDC2 |
| 650 | CHL1 |
| 651 | CHODL |
| 652 | CHPF |
| 653 | CHPF2 |
| 654 | CHPT1 |
| 655 | CHRM2 |
| 656 | CHRM4 |
| 657 | CHRM5 |
| 658 | CHRNA1 |
| 659 | CHRNA3 |
| 660 | CHRNA4 |
| 661 | CHRNA5 |
| 662 | CHRNA9 |

| | |
|---|---|
| 663 | CHRNB1 |
| 664 | CHRNB2 |
| 665 | CHRNB3 |
| 666 | CHRND |
| 667 | CHRNE |
| 668 | CHST1 |
| 669 | CHST10 |
| 670 | CHST11 |
| 671 | CHST12 |
| 672 | CHST13 |
| 673 | CHST14 |
| 674 | CHST15 |
| 675 | CHST2 |
| 676 | CHST3 |
| 677 | CHST4 |
| 678 | CHST5 |
| 679 | CHST6 |
| 680 | CHST7 |
| 681 | CHST8 |
| 682 | CHST9 |
| 683 | CHSY1 |
| 684 | CHSY3 |
| 685 | CISD1 |
| 686 | CISD2 |
| 687 | CKAP4 |
| 688 | CLCA2 |
| 689 | CLCA4 |
| 690 | CLCN3 |
| 691 | CLDN1 |
| 692 | CLDN11 |
| 693 | CLDN12 |
| 694 | CLDN15 |
| 695 | CLDN16 |
| 696 | CLDN17 |
| 697 | CLDN18 |
| 698 | CLDN2 |
| 699 | CLDN20 |
| 700 | CLDN22 |
| 701 | CLDN23 |
| 702 | CLDN25 |
| 703 | CLDN3 |
| 704 | CLDN5 |
| 705 | CLDN8 |
| 706 | CLDND1 |
| 707 | CLDND2 |
| 708 | CLEC10A |
| 709 | CLEC12A |
| 710 | CLEC12B |
| 711 | CLEC14A |
| 712 | CLEC17A |
| 713 | CLEC1A |
| 714 | CLEC1B |
| 715 | CLEC2A |
| 716 | CLEC2B |
| 717 | CLEC2D |
| 718 | CLEC2L |
| 719 | CLEC4A |
| 720 | CLEC4C |
| 721 | CLEC4D |
| 722 | CLEC4E |
| 723 | CLEC4F |
| 724 | CLEC4G |
| 725 | CLEC4M |
| 726 | CLEC5A |
| 727 | CLEC6A |
| 728 | CLEC7A |
| 729 | CLEC9A |
| 730 | CLECL1 |
| 731 | CLIC1 |
| 732 | CLIC2 |
| 733 | CLIC3 |
| 734 | CLIC4 |
| 735 | CLIC5 |
| 736 | CLIC6 |
| 737 | CLMN |
| 738 | CLMP |
| 739 | CLN3 |
| 740 | CLN6 |
| 741 | CLN8 |
| 742 | CLRN1 |
| 743 | CLRN3 |
| 744 | CLSTN1 |
| 745 | CLSTN2 |

| | |
|---|---|
| 746 | CLSTN3 |
| 747 | CMKLR1 |
| 748 | CMTM1 |
| 749 | CMTM2 |
| 750 | CMTM6 |
| 751 | CMTM7 |
| 752 | CNGA1 |
| 753 | CNGB1 |
| 754 | CNGB3 |
| 755 | CNIH |
| 756 | CNIH2 |
| 757 | CNIH3 |
| 758 | CNIH4 |
| 759 | CNNM1 |
| 760 | CNNM3 |
| 761 | CNNM4 |
| 762 | CNPPD1 |
| 763 | CNR1 |
| 764 | CNST |
| 765 | CNTN2 |
| 766 | CNTNAP1 |
| 767 | CNTNAP2 |
| 768 | CNTNAP3 |
| 769 | CNTNAP3B |
| 770 | CNTNAP4 |
| 771 | CNTNAP5 |
| 772 | COA1 |
| 773 | COA3 |
| 774 | COL13A1 |
| 775 | COL17A1 |
| 776 | COL23A1 |
| 777 | COL25A1 |
| 778 | COLEC12 |
| 779 | COMT |
| 780 | COMTD1 |
| 781 | COQ2 |
| 782 | CORIN |
| 783 | COX11 |
| 784 | COX14 |
| 785 | COX6C |
| 786 | COX7A1 |
| 787 | COX7B |
| 788 | COX7B2 |
| 789 | COX7C |
| 790 | COX8A |
| 791 | COX8C |
| 792 | CPD |
| 793 | CPM |
| 794 | CPT1A |
| 795 | CPT1C |
| 796 | CR1 |
| 797 | CR2 |
| 798 | CRB1 |
| 799 | CRB2 |
| 800 | CRB3 |
| 801 | CREB3 |
| 802 | CREB3L1 |
| 803 | CREB3L2 |
| 804 | CREB3L3 |
| 805 | CREB3L4 |
| 806 | CRHR2 |
| 807 | CRIM1 |
| 808 | CRLF2 |
| 809 | CRTAM |
| 810 | CRYAB |
| 811 | CSF1 |
| 812 | CSF1R |
| 813 | CSF2RA |
| 814 | CSF2RB |
| 815 | CSF3R |
| 816 | CSGALNACT1 |
| 817 | CSGALNACT2 |
| 818 | CSMD1 |
| 819 | CSMD2 |
| 820 | CSMD3 |
| 821 | CSPG4 |
| 822 | CSPG5 |
| 823 | CST8 |
| 824 | CTAGE1 |
| 825 | CTAGE15P |
| 826 | CTAGE4 |
| 827 | CTAGE6P |
| 828 | CTAGE9 |

| | |
|---|---|
| 829 | CTDNEP1 |
| 830 | CTH |
| 831 | CTLA4 |
| 832 | CTNS |
| 833 | CTSG |
| 834 | CTSL2 |
| 835 | CTXN3 |
| 836 | CUBN |
| 837 | CWH43 |
| 838 | CX3CL1 |
| 839 | CX3CR1 |
| 840 | CXADR |
| 841 | CXCL10 |
| 842 | CXCL12 |
| 843 | CXCL16 |
| 844 | CXCL9 |
| 845 | CXCR1 |
| 846 | CXCR2 |
| 847 | CXCR3 |
| 848 | CXCR4 |
| 849 | CXCR5 |
| 850 | CXCR6 |
| 851 | CXCR7 |
| 852 | CXorf61 |
| 853 | CXorf66 |
| 854 | CXXC11 |
| 855 | CYB561 |
| 856 | CYB561D1 |
| 857 | CYB561D2 |
| 858 | CYB5A |
| 859 | CYB5B |
| 860 | CYB5R1 |
| 861 | CYBRD1 |
| 862 | CYC1 |
| 863 | CYP20A1 |
| 864 | CYP26C1 |
| 865 | CYP2W1 |
| 866 | CYP3A4 |
| 867 | CYP46A1 |
| 868 | CYP4F11 |
| 869 | CYP4F3 |
| 870 | CYP4F8 |
| 871 | CYP4V2 |
| 872 | CYP4X1 |
| 873 | CYP4Z1 |
| 874 | CYP4Z2P |
| 875 | CYP51A1 |
| 876 | CYP8B1 |
| 877 | CYSTM1 |
| 878 | CYYR1 |
| 879 | DAD1 |
| 880 | DAG1 |
| 881 | DAGLA |
| 882 | DAGLB |
| 883 | DARC |
| 884 | DBH |
| 885 | DCBLD1 |
| 886 | DCBLD2 |
| 887 | DCC |
| 888 | DCHS1 |
| 889 | DCHS2 |
| 890 | DCST1 |
| 891 | DCST2 |
| 892 | DCSTAMP |
| 893 | DCT |
| 894 | DCTN3 |
| 895 | DDR1 |
| 896 | DDR2 |
| 897 | DEFB104A |
| 898 | DEFB106A |
| 899 | DEFB108B |
| 900 | DEFB115 |
| 901 | DEFB116 |
| 902 | DEFB118 |
| 903 | DEFB119 |
| 904 | DEFB121 |
| 905 | DEFB123 |
| 906 | DEFB124 |
| 907 | DEFB125 |
| 908 | DEFB126 |
| 909 | DEFB127 |
| 910 | DEFB128 |
| 911 | DEFB129 |

| | |
|---|---|
| 912 | DEFB131 |
| 913 | DEFB132 |
| 914 | DEFB134 |
| 915 | DEFB135 |
| 916 | DEGS1 |
| 917 | DEGS2 |
| 918 | DENND5B |
| 919 | DERL1 |
| 920 | DERL2 |
| 921 | DERL3 |
| 922 | DGAT2L6 |
| 923 | DGKE |
| 924 | DHCR24 |
| 925 | DHCR7 |
| 926 | DHODH |
| 927 | DHRS7B |
| 928 | DIO1 |
| 929 | DIO2 |
| 930 | DIO3 |
| 931 | DIP2A |
| 932 | DLK1 |
| 933 | DLK2 |
| 934 | DLL1 |
| 935 | DLL3 |
| 936 | DLL4 |
| 937 | DMD |
| 938 | DMPK |
| 939 | DNAJB12 |
| 940 | DNAJB14 |
| 941 | DNAJC1 |
| 942 | DNAJC14 |
| 943 | DNAJC15 |
| 944 | DNAJC16 |
| 945 | DNAJC18 |
| 946 | DNAJC19 |
| 947 | DNAJC22 |
| 948 | DNAJC4 |
| 949 | DNER |
| 950 | DPAGT1 |
| 951 | DPCR1 |
| 952 | DPM2 |
| 953 | DPM3 |
| 954 | DPP10 |
| 955 | DPP4 |
| 956 | DPP6 |
| 957 | DPY19L1 |
| 958 | DPY19L2P1 |
| 959 | DPY19L3 |
| 960 | DRAM1 |
| 961 | DRAM2 |
| 962 | DRD4 |
| 963 | DSC2 |
| 964 | DSC3 |
| 965 | DSCAM |
| 966 | DSCAML1 |
| 967 | DSEL |
| 968 | DSG1 |
| 969 | DSG2 |
| 970 | DSG3 |
| 971 | DSG4 |
| 972 | DUOX2 |
| 973 | DYNAP |
| 974 | DYSF |
| 975 | EBAG9 |
| 976 | EBP |
| 977 | EBPL |
| 978 | ECE1 |
| 979 | ECE2 |
| 980 | ECEL1 |
| 981 | ECSCR |
| 982 | ECT2 |
| 983 | EDA2R |
| 984 | EDAR |
| 985 | EDEM1 |
| 986 | EDNRA |
| 987 | EDNRB |
| 988 | EFHA2 |
| 989 | EFNA5 |
| 990 | EFNB1 |
| 991 | EFNB2 |
| 992 | EGF |
| 993 | EGFR |
| 994 | ELANE |

| | |
|---|---|
| 995 | ELFN1 |
| 996 | ELFN2 |
| 997 | ELOVL1 |
| 998 | ELOVL2 |
| 999 | ELOVL4 |
| 1000 | ELOVL5 |
| 1001 | EMC3 |
| 1002 | EMC4 |
| 1003 | EMC6 |
| 1004 | EMCN |
| 1005 | EMD |
| 1006 | EMP1 |
| 1007 | EMP2 |
| 1008 | EMP3 |
| 1009 | EMR1 |
| 1010 | EMR2 |
| 1011 | EMR4P |
| 1012 | ENG |
| 1013 | ENOX2 |
| 1014 | ENPEP |
| 1015 | ENPP1 |
| 1016 | ENPP3 |
| 1017 | ENPP4 |
| 1018 | ENPP5 |
| 1019 | ENTPD1 |
| 1020 | ENTPD2 |
| 1021 | ENTPD3 |
| 1022 | ENTPD6 |
| 1023 | ENTPD8 |
| 1024 | EPCAM |
| 1025 | EPGN |
| 1026 | EPHA1 |
| 1027 | EPHA10 |
| 1028 | EPHA2 |
| 1029 | EPHA3 |
| 1030 | EPHA4 |
| 1031 | EPHA5 |
| 1032 | EPHA6 |
| 1033 | EPHA7 |
| 1034 | EPHA8 |
| 1035 | EPHB1 |
| 1036 | EPHB2 |
| 1037 | EPHB3 |
| 1038 | EPHB4 |
| 1039 | EPHB6 |
| 1040 | EPHX1 |
| 1041 | EPHX4 |
| 1042 | EPOR |
| 1043 | EPPIN |
| 1044 | EPT1 |
| 1045 | EQTN |
| 1046 | ERAP1 |
| 1047 | ERAP2 |
| 1048 | ERBB2 |
| 1049 | ERBB3 |
| 1050 | ERBB4 |
| 1051 | EREG |
| 1052 | ERG |
| 1053 | ERGIC1 |
| 1054 | ERGIC2 |
| 1055 | ERLIN1 |
| 1056 | ERLIN2 |
| 1057 | ERMAP |
| 1058 | ERN2 |
| 1059 | ERP29 |
| 1060 | ERP44 |
| 1061 | ERVK13-1 |
| 1062 | ERVMER34-1 |
| 1063 | ERVV-2 |
| 1064 | ESAM |
| 1065 | ESYT1 |
| 1066 | ESYT2 |
| 1067 | ESYT3 |
| 1068 | EVA1A |
| 1069 | EVA1C |
| 1070 | EVC |
| 1071 | EVC2 |
| 1072 | EVI2A |
| 1073 | EVI2B |
| 1074 | EXT1 |
| 1075 | EXT2 |
| 1076 | EXTL1 |
| 1077 | EXTL2 |

| | |
|---|---|
| 1078 | EXTL3 |
| 1079 | F10 |
| 1080 | F11R |
| 1081 | F2R |
| 1082 | F2RL1 |
| 1083 | F2RL2 |
| 1084 | F2RL3 |
| 1085 | F3 |
| 1086 | FAAH |
| 1087 | FAAH2 |
| 1088 | FADS1 |
| 1089 | FADS2 |
| 1090 | FADS3 |
| 1091 | FADS6 |
| 1092 | FAIM2 |
| 1093 | FAIM3 |
| 1094 | FAM118A |
| 1095 | FAM134A |
| 1096 | FAM134B |
| 1097 | FAM134C |
| 1098 | FAM151A |
| 1099 | FAM156A |
| 1100 | FAM159B |
| 1101 | FAM162A |
| 1102 | FAM162B |
| 1103 | FAM168B |
| 1104 | FAM171A1 |
| 1105 | FAM171A2 |
| 1106 | FAM171B |
| 1107 | FAM173A |
| 1108 | FAM173B |
| 1109 | FAM174B |
| 1110 | FAM180B |
| 1111 | FAM187B |
| 1112 | FAM189A1 |
| 1113 | FAM189A2 |
| 1114 | FAM189B |
| 1115 | FAM198B |
| 1116 | FAM200A |
| 1117 | FAM205A |
| 1118 | FAM209A |
| 1119 | FAM20B |
| 1120 | FAM210A |
| 1121 | FAM210B |
| 1122 | FAM26D |
| 1123 | FAM26E |
| 1124 | FAM57A |
| 1125 | FAM69A |
| 1126 | FAM69B |
| 1127 | FAM69C |
| 1128 | FAM73A |
| 1129 | FAM73B |
| 1130 | FAM74A3 |
| 1131 | FAM89B |
| 1132 | FAP |
| 1133 | FAR1 |
| 1134 | FAR2 |
| 1135 | FAS |
| 1136 | FASLG |
| 1137 | FAT1 |
| 1138 | FAT2 |
| 1139 | FAT3 |
| 1140 | FAT4 |
| 1141 | FATE1 |
| 1142 | FAXC |
| 1143 | FBP2 |
| 1144 | FCAMR |
| 1145 | FCAR |
| 1146 | FCER1A |
| 1147 | FCER1G |
| 1148 | FCER2 |
| 1149 | FCGR1A |
| 1150 | FCGR2A |
| 1151 | FCGR2B |
| 1152 | FCGR2C |
| 1153 | FCGR3A |
| 1154 | FCGRT |
| 1155 | FCN1 |
| 1156 | FCRL3 |
| 1157 | FCRL4 |
| 1158 | FCRL5 |
| 1159 | FCRL6 |
| 1160 | FER1L5 |

| | |
|---|---|
| 1161 | FER1L6 |
| 1162 | FERMT2 |
| 1163 | FFAR2 |
| 1164 | FGA |
| 1165 | FGB |
| 1166 | FGF10 |
| 1167 | FGF22 |
| 1168 | FGF8 |
| 1169 | FGFBP1 |
| 1170 | FGFR1 |
| 1171 | FGFR2 |
| 1172 | FGFR3 |
| 1173 | FGFR4 |
| 1174 | FGFRL1 |
| 1175 | FGG |
| 1176 | FIBCD1 |
| 1177 | FICD |
| 1178 | FIS1 |
| 1179 | FITM1 |
| 1180 | FKBP8 |
| 1181 | FKRP |
| 1182 | FKTN |
| 1183 | FLOT1 |
| 1184 | FLRT1 |
| 1185 | FLRT2 |
| 1186 | FLRT3 |
| 1187 | FLT1 |
| 1188 | FLT3 |
| 1189 | FLT3LG |
| 1190 | FLT4 |
| 1191 | FLVCR1 |
| 1192 | FMO2 |
| 1193 | FMO4 |
| 1194 | FMO6P |
| 1195 | FNDC3A |
| 1196 | FNDC3B |
| 1197 | FNDC4 |
| 1198 | FNDC5 |
| 1199 | FNDC9 |
| 1200 | FOCAD |
| 1201 | FOLH1 |
| 1202 | FOLR1 |
| 1203 | FOLR2 |
| 1204 | FOXRED1 |
| 1205 | FPR2 |
| 1206 | FRAS1 |
| 1207 | FREM2 |
| 1208 | FRMD3 |
| 1209 | FRMD5 |
| 1210 | FRRS1 |
| 1211 | FRRS1L |
| 1212 | FSD1 |
| 1213 | FSHR |
| 1214 | FTSJD1 |
| 1215 | FURIN |
| 1216 | FUT1 |
| 1217 | FUT10 |
| 1218 | FUT11 |
| 1219 | FUT2 |
| 1220 | FUT3 |
| 1221 | FUT4 |
| 1222 | FUT5 |
| 1223 | FUT6 |
| 1224 | FUT7 |
| 1225 | FUT8 |
| 1226 | FUT9 |
| 1227 | FXYD1 |
| 1228 | FXYD2 |
| 1229 | FXYD3 |
| 1230 | FXYD4 |
| 1231 | FXYD5 |
| 1232 | FXYD6 |
| 1233 | FZD1 |
| 1234 | FZD10 |
| 1235 | FZD2 |
| 1236 | FZD3 |
| 1237 | FZD4 |
| 1238 | FZD5 |
| 1239 | FZD7 |
| 1240 | FZD8 |
| 1241 | FZD9 |
| 1242 | GABBR1 |
| 1243 | GABBR2 |

| | |
|---|---|
| 1244 | GABRA1 |
| 1245 | GABRA2 |
| 1246 | GABRA3 |
| 1247 | GABRA4 |
| 1248 | GABRA5 |
| 1249 | GABRA6 |
| 1250 | GABRB1 |
| 1251 | GABRB2 |
| 1252 | GABRD |
| 1253 | GABRE |
| 1254 | GABRG1 |
| 1255 | GABRG3 |
| 1256 | GABRP |
| 1257 | GABRR3 |
| 1258 | GAL3ST1 |
| 1259 | GAL3ST2 |
| 1260 | GAL3ST3 |
| 1261 | GAL3ST4 |
| 1262 | GALNT1 |
| 1263 | GALNT10 |
| 1264 | GALNT11 |
| 1265 | GALNT12 |
| 1266 | GALNT13 |
| 1267 | GALNT14 |
| 1268 | GALNT15 |
| 1269 | GALNT16 |
| 1270 | GALNT18 |
| 1271 | GALNT2 |
| 1272 | GALNT3 |
| 1273 | GALNT4 |
| 1274 | GALNT5 |
| 1275 | GALNT6 |
| 1276 | GALNT7 |
| 1277 | GALNT8 |
| 1278 | GALNT9 |
| 1279 | GALNTL5 |
| 1280 | GALNTL6 |
| 1281 | GALR1 |
| 1282 | GALR2 |
| 1283 | GALR3 |
| 1284 | GAPT |
| 1285 | GBGT1 |
| 1286 | GCNT2 |
| 1287 | GCNT3 |
| 1288 | GCNT4 |
| 1289 | GCNT7 |
| 1290 | GDAP1 |
| 1291 | GDE1 |
| 1292 | GDPD1 |
| 1293 | GDPD2 |
| 1294 | GDPD3 |
| 1295 | GDPD4 |
| 1296 | GDPD5 |
| 1297 | GFRA3 |
| 1298 | GFRAL |
| 1299 | GGCX |
| 1300 | GGT1 |
| 1301 | GGT3P |
| 1302 | GGT5 |
| 1303 | GGT6 |
| 1304 | GGT7 |
| 1305 | GGTA1P |
| 1306 | GGTLC1 |
| 1307 | GGTLC2 |
| 1308 | GHITM |
| 1309 | GHR |
| 1310 | GHRHR |
| 1311 | GHSR |
| 1312 | GIMAP1 |
| 1313 | GIMAP2 |
| 1314 | GIMAP5 |
| 1315 | GINM1 |
| 1316 | GIPR |
| 1317 | GJA1 |
| 1318 | GJA10 |
| 1319 | GJA3 |
| 1320 | GJA4 |
| 1321 | GJA5 |
| 1322 | GJA8 |
| 1323 | GJA9 |
| 1324 | GJB1 |
| 1325 | GJB2 |
| 1326 | GJB3 |

| | |
|---|---|
| 1327 | GJB4 |
| 1328 | GJB5 |
| 1329 | GJB6 |
| 1330 | GJC1 |
| 1331 | GJC2 |
| 1332 | GJC3 |
| 1333 | GJD2 |
| 1334 | GJD3 |
| 1335 | GJD4 |
| 1336 | GLCE |
| 1337 | GLDN |
| 1338 | GLG1 |
| 1339 | GLIPR1 |
| 1340 | GLIPR1L2 |
| 1341 | GLP1R |
| 1342 | GLP2R |
| 1343 | GLRA1 |
| 1344 | GLRA2 |
| 1345 | GLRA3 |
| 1346 | GLRA4 |
| 1347 | GLRB |
| 1348 | GLT6D1 |
| 1349 | GLT8D1 |
| 1350 | GLT8D2 |
| 1351 | GNPTAB |
| 1352 | GNRHR2 |
| 1353 | GOLGA5 |
| 1354 | GOLGB1 |
| 1355 | GOLIM4 |
| 1356 | GOLM1 |
| 1357 | GOSR1 |
| 1358 | GOSR2 |
| 1359 | GP1BA |
| 1360 | GP1BB |
| 1361 | GP5 |
| 1362 | GP6 |
| 1363 | GP9 |
| 1364 | GPA33 |
| 1365 | GPAA1 |
| 1366 | GPAM |
| 1367 | GPAT2 |
| 1368 | GPBAR1 |
| 1369 | GPC4 |
| 1370 | GPER |
| 1371 | GPIHBP1 |
| 1372 | GPM6A |
| 1373 | GPM6B |
| 1374 | GPR1 |
| 1375 | GPR101 |
| 1376 | GPR108 |
| 1377 | GPR110 |
| 1378 | GPR112 |
| 1379 | GPR115 |
| 1380 | GPR116 |
| 1381 | GPR119 |
| 1382 | GPR12 |
| 1383 | GPR125 |
| 1384 | GPR126 |
| 1385 | GPR128 |
| 1386 | GPR132 |
| 1387 | GPR133 |
| 1388 | GPR135 |
| 1389 | GPR137C |
| 1390 | GPR139 |
| 1391 | GPR142 |
| 1392 | GPR143 |
| 1393 | GPR144 |
| 1394 | GPR148 |
| 1395 | GPR149 |
| 1396 | GPR15 |
| 1397 | GPR151 |
| 1398 | GPR155 |
| 1399 | GPR156 |
| 1400 | GPR157 |
| 1401 | GPR161 |
| 1402 | GPR171 |
| 1403 | GPR173 |
| 1404 | GPR18 |
| 1405 | GPR180 |
| 1406 | GPR183 |
| 1407 | GPR20 |
| 1408 | GPR21 |
| 1409 | GPR25 |

| | |
|---|---|
| 1410 | GPR3 |
| 1411 | GPR33 |
| 1412 | GPR35 |
| 1413 | GPR37 |
| 1414 | GPR37L1 |
| 1415 | GPR45 |
| 1416 | GPR50 |
| 1417 | GPR55 |
| 1418 | GPR56 |
| 1419 | GPR6 |
| 1420 | GPR61 |
| 1421 | GPR62 |
| 1422 | GPR63 |
| 1423 | GPR64 |
| 1424 | GPR65 |
| 1425 | GPR68 |
| 1426 | GPR85 |
| 1427 | GPR87 |
| 1428 | GPR98 |
| 1429 | GPRC5A |
| 1430 | GPRC5B |
| 1431 | GPRC5C |
| 1432 | GPRC5D |
| 1433 | GPRC6A |
| 1434 | GPX8 |
| 1435 | GRAMD1A |
| 1436 | GRAMD1B |
| 1437 | GRAMD1C |
| 1438 | GRAMD2 |
| 1439 | GRAMD4 |
| 1440 | GREB1 |
| 1441 | GREB1L |
| 1442 | GRIA1 |
| 1443 | GRIA2 |
| 1444 | GRIA3 |
| 1445 | GRIA4 |
| 1446 | GRID1 |
| 1447 | GRID2 |
| 1448 | GRIK1 |
| 1449 | GRIK2 |
| 1450 | GRIK3 |
| 1451 | GRIK4 |
| 1452 | GRIK5 |
| 1453 | GRIN1 |
| 1454 | GRIN2B |
| 1455 | GRIN3A |
| 1456 | GRIN3B |
| 1457 | GRINA |
| 1458 | GRM1 |
| 1459 | GRM2 |
| 1460 | GRM3 |
| 1461 | GRM4 |
| 1462 | GRM5 |
| 1463 | GRM6 |
| 1464 | GRM7 |
| 1465 | GRM8 |
| 1466 | GRPR |
| 1467 | GSR |
| 1468 | GTDC2 |
| 1469 | GUCY2C |
| 1470 | GUCY2D |
| 1471 | GUCY2F |
| 1472 | GXYLT1 |
| 1473 | GXYLT2 |
| 1474 | GYLTL1B |
| 1475 | GYPA |
| 1476 | GYPB |
| 1477 | GYPC |
| 1478 | GYPE |
| 1479 | HAS2 |
| 1480 | HAS3 |
| 1481 | HAVCR1 |
| 1482 | HAVCR2 |
| 1483 | HBEGF |
| 1484 | HBP1 |
| 1485 | HCAR2 |
| 1486 | HCN1 |
| 1487 | HCN2 |
| 1488 | HCN3 |
| 1489 | HCN4 |
| 1490 | HCST |
| 1491 | HECTD4 |
| 1492 | HEG1 |

| | |
|---|---|
| 1493 | HEPACAM |
| 1494 | HEPACAM2 |
| 1495 | HEPH |
| 1496 | HEPHL1 |
| 1497 | HERPUD1 |
| 1498 | HERPUD2 |
| 1499 | HFE |
| 1500 | HFE2 |
| 1501 | HHATL |
| 1502 | HHIP |
| 1503 | HHLA2 |
| 1504 | HIAT1 |
| 1505 | HIATL2 |
| 1506 | HIGD1A |
| 1507 | HIGD1B |
| 1508 | HIGD1C |
| 1509 | HIGD2A |
| 1510 | HIGD2B |
| 1511 | HILPDA |
| 1512 | HLA-A |
| 1513 | HLA-B |
| 1514 | HLA-C |
| 1515 | HLA-DMA |
| 1516 | HLA-DMB |
| 1517 | HLA-DOA |
| 1518 | HLA-DOB |
| 1519 | HLA-DPA1 |
| 1520 | HLA-DPB1 |
| 1521 | HLA-DQA1 |
| 1522 | HLA-DQA2 |
| 1523 | HLA-DQB1 |
| 1524 | HLA-DQB2 |
| 1525 | HLA-DRA |
| 1526 | HLA-DRB1 |
| 1527 | HLA-DRB5 |
| 1528 | HLA-E |
| 1529 | HLA-F |
| 1530 | HLA-G |
| 1531 | HLA-H |
| 1532 | HMGB1 |
| 1533 | HMGCR |
| 1534 | HMMR |
| 1535 | HNRNPU |
| 1536 | HPN |
| 1537 | HRASLS |
| 1538 | HRASLS2 |
| 1539 | HRCT1 |
| 1540 | HRK |
| 1541 | HS2ST1 |
| 1542 | HS3ST2 |
| 1543 | HS3ST3A1 |
| 1544 | HS3ST3B1 |
| 1545 | HS3ST4 |
| 1546 | HS3ST5 |
| 1547 | HS3ST6 |
| 1548 | HS6ST1 |
| 1549 | HS6ST2 |
| 1550 | HS6ST3 |
| 1551 | HSD11B1 |
| 1552 | HSD17B2 |
| 1553 | HSD17B7 |
| 1554 | HSD3B1 |
| 1555 | HSD3B2 |
| 1556 | HSD3B7 |
| 1557 | HSP90AB1 |
| 1558 | HSPA2 |
| 1559 | HSPA5 |
| 1560 | HSPD1 |
| 1561 | HTR1A |
| 1562 | HTR1B |
| 1563 | HTR1E |
| 1564 | HTR1F |
| 1565 | HTR2A |
| 1566 | HTR2B |
| 1567 | HTR2C |
| 1568 | HTR3A |
| 1569 | HTR3B |
| 1570 | HTR3C |
| 1571 | HTR3D |
| 1572 | HTR3E |
| 1573 | HTR4 |
| 1574 | HTR5A |
| 1575 | HTR6 |

| | |
|---|---|
| 1576 | HTRA2 |
| 1577 | HTT |
| 1578 | HVCN1 |
| 1579 | HYAL2 |
| 1580 | HYAL4 |
| 1581 | ICAM1 |
| 1582 | ICAM2 |
| 1583 | ICAM3 |
| 1584 | ICAM4 |
| 1585 | ICOS |
| 1586 | ICOSLG |
| 1587 | IDE |
| 1588 | IER3 |
| 1589 | IER3IP1 |
| 1590 | IFI27 |
| 1591 | IFI6 |
| 1592 | IFITM1 |
| 1593 | IFITM2 |
| 1594 | IFITM3 |
| 1595 | IFNAR2 |
| 1596 | IFNG |
| 1597 | IFNGR1 |
| 1598 | IFNGR2 |
| 1599 | IFNLR1 |
| 1600 | IGDCC3 |
| 1601 | IGDCC4 |
| 1602 | IGF1R |
| 1603 | IGF2R |
| 1604 | IGFLR1 |
| 1605 | IGLON5 |
| 1606 | IGSF1 |
| 1607 | IGSF11 |
| 1608 | IGSF23 |
| 1609 | IGSF3 |
| 1610 | IGSF6 |
| 1611 | IGSF8 |
| 1612 | IKBIP |
| 1613 | IL10RA |
| 1614 | IL10RB |
| 1615 | IL11RA |
| 1616 | IL12RB1 |
| 1617 | IL12RB2 |
| 1618 | IL13 |
| 1619 | IL13RA1 |
| 1620 | IL13RA2 |
| 1621 | IL15 |
| 1622 | IL15RA |
| 1623 | IL17A |
| 1624 | IL17RA |
| 1625 | IL17RB |
| 1626 | IL17RC |
| 1627 | IL17RD |
| 1628 | IL17RE |
| 1629 | IL18R1 |
| 1630 | IL18RAP |
| 1631 | IL1R1 |
| 1632 | IL1R2 |
| 1633 | IL1RAP |
| 1634 | IL1RAPL1 |
| 1635 | IL1RAPL2 |
| 1636 | IL1RL1 |
| 1637 | IL1RL2 |
| 1638 | IL20RA |
| 1639 | IL20RB |
| 1640 | IL21R |
| 1641 | IL23R |
| 1642 | IL27RA |
| 1643 | IL2RA |
| 1644 | IL2RB |
| 1645 | IL2RG |
| 1646 | IL31RA |
| 1647 | IL3RA |
| 1648 | IL4 |
| 1649 | IL4R |
| 1650 | IL5RA |
| 1651 | IL6 |
| 1652 | IL6R |
| 1653 | IL6ST |
| 1654 | IL7R |
| 1655 | IL9R |
| 1656 | ILDR1 |
| 1657 | ILDR2 |
| 1658 | ILVBL |

| | |
|---|---|
| 1659 | IMMP2L |
| 1660 | IMMT |
| 1661 | IMP3 |
| 1662 | IMPAD1 |
| 1663 | IMPG2 |
| 1664 | INSIG1 |
| 1665 | INSIG2 |
| 1666 | INSR |
| 1667 | INSRR |
| 1668 | INTS1 |
| 1669 | INTS2 |
| 1670 | INTS5 |
| 1671 | INTU |
| 1672 | IQGAP2 |
| 1673 | ITFG1 |
| 1674 | ITFG3 |
| 1675 | ITGA1 |
| 1676 | ITGA10 |
| 1677 | ITGA11 |
| 1678 | ITGA2 |
| 1679 | ITGA2B |
| 1680 | ITGA3 |
| 1681 | ITGA4 |
| 1682 | ITGA5 |
| 1683 | ITGA6 |
| 1684 | ITGA7 |
| 1685 | ITGA8 |
| 1686 | ITGA9 |
| 1687 | ITGAD |
| 1688 | ITGAE |
| 1689 | ITGAL |
| 1690 | ITGAM |
| 1691 | ITGAV |
| 1692 | ITGAX |
| 1693 | ITGB1 |
| 1694 | ITGB2 |
| 1695 | ITGB3 |
| 1696 | ITGB4 |
| 1697 | ITGB5 |
| 1698 | ITGB6 |
| 1699 | ITGB7 |
| 1700 | ITGB8 |
| 1701 | ITM2A |
| 1702 | ITM2B |
| 1703 | ITM2C |
| 1704 | ITPR1 |
| 1705 | ITPR3 |
| 1706 | ITPRIPL2 |
| 1707 | IYD |
| 1708 | IZUMO1 |
| 1709 | IZUMO3 |
| 1710 | JAG1 |
| 1711 | JAG2 |
| 1712 | JAM2 |
| 1713 | JAM3 |
| 1714 | JPH1 |
| 1715 | JPH2 |
| 1716 | JPH3 |
| 1717 | JPH4 |
| 1718 | JTB |
| 1719 | KCNA1 |
| 1720 | KCNA10 |
| 1721 | KCNA2 |
| 1722 | KCNA4 |
| 1723 | KCNA5 |
| 1724 | KCNA6 |
| 1725 | KCNA7 |
| 1726 | KCNB1 |
| 1727 | KCNB2 |
| 1728 | KCNC1 |
| 1729 | KCNC2 |
| 1730 | KCNC3 |
| 1731 | KCNC4 |
| 1732 | KCND1 |
| 1733 | KCND2 |
| 1734 | KCND3 |
| 1735 | KCNE1 |
| 1736 | KCNE1L |
| 1737 | KCNE2 |
| 1738 | KCNE3 |
| 1739 | KCNE4 |
| 1740 | KCNF1 |
| 1741 | KCNG1 |

| | |
|---|---|
| 1742 | KCNG2 |
| 1743 | KCNG3 |
| 1744 | KCNG4 |
| 1745 | KCNH2 |
| 1746 | KCNH5 |
| 1747 | KCNH6 |
| 1748 | KCNH7 |
| 1749 | KCNJ1 |
| 1750 | KCNJ10 |
| 1751 | KCNJ11 |
| 1752 | KCNJ12 |
| 1753 | KCNJ13 |
| 1754 | KCNJ14 |
| 1755 | KCNJ15 |
| 1756 | KCNJ16 |
| 1757 | KCNJ18 |
| 1758 | KCNJ2 |
| 1759 | KCNJ3 |
| 1760 | KCNJ4 |
| 1761 | KCNJ5 |
| 1762 | KCNJ6 |
| 1763 | KCNJ8 |
| 1764 | KCNJ9 |
| 1765 | KCNK1 |
| 1766 | KCNK10 |
| 1767 | KCNK12 |
| 1768 | KCNK13 |
| 1769 | KCNK15 |
| 1770 | KCNK17 |
| 1771 | KCNK18 |
| 1772 | KCNK2 |
| 1773 | KCNK3 |
| 1774 | KCNK4 |
| 1775 | KCNK5 |
| 1776 | KCNK7 |
| 1777 | KCNK9 |
| 1778 | KCNMA1 |
| 1779 | KCNMB1 |
| 1780 | KCNMB4 |
| 1781 | KCNN1 |
| 1782 | KCNN2 |
| 1783 | KCNN3 |
| 1784 | KCNN4 |
| 1785 | KCNQ1 |
| 1786 | KCNQ2 |
| 1787 | KCNQ3 |
| 1788 | KCNQ4 |
| 1789 | KCNQ5 |
| 1790 | KCNS1 |
| 1791 | KCNS2 |
| 1792 | KCNS3 |
| 1793 | KCNT1 |
| 1794 | KCNU1 |
| 1795 | KCNV1 |
| 1796 | KDELR1 |
| 1797 | KDELR2 |
| 1798 | KDELR3 |
| 1799 | KDR |
| 1800 | KDSR |
| 1801 | KEL |
| 1802 | KIAA0195 |
| 1803 | KIAA0247 |
| 1804 | KIAA0319 |
| 1805 | KIAA0319L |
| 1806 | KIAA0922 |
| 1807 | KIAA1024 |
| 1808 | KIAA1024L |
| 1809 | KIAA1109 |
| 1810 | KIAA1161 |
| 1811 | KIAA1244 |
| 1812 | KIAA1324 |
| 1813 | KIAA1324L |
| 1814 | KIAA1432 |
| 1815 | KIAA1467 |
| 1816 | KIAA1524 |
| 1817 | KIAA1549 |
| 1818 | KIAA1549L |
| 1819 | KIAA1644 |
| 1820 | KIAA1919 |
| 1821 | KIDINS220 |
| 1822 | KIR2DL1 |
| 1823 | KIR2DL3 |
| 1824 | KIR2DL4 |

| | |
|---|---|
| 1825 | KIR2DS4 |
| 1826 | KIR3DL1 |
| 1827 | KIR3DL2 |
| 1828 | KIR3DL3 |
| 1829 | KIRREL |
| 1830 | KIRREL2 |
| 1831 | KIRREL3 |
| 1832 | KIT |
| 1833 | KITLG |
| 1834 | KL |
| 1835 | KLB |
| 1836 | KLHDC7A |
| 1837 | KLRB1 |
| 1838 | KLRC1 |
| 1839 | KLRC2 |
| 1840 | KLRC3 |
| 1841 | KLRC4 |
| 1842 | KLRD1 |
| 1843 | KLRF1 |
| 1844 | KLRF2 |
| 1845 | KLRG1 |
| 1846 | KLRG2 |
| 1847 | KLRK1 |
| 1848 | KMO |
| 1849 | KREMEN1 |
| 1850 | KREMEN2 |
| 1851 | KRT4 |
| 1852 | KRTCAP2 |
| 1853 | KTN1 |
| 1854 | L1CAM |
| 1855 | LAG3 |
| 1856 | LAIR1 |
| 1857 | LAMP1 |
| 1858 | LAMP2 |
| 1859 | LAMP3 |
| 1860 | LAMP5 |
| 1861 | LAPTM4A |
| 1862 | LAPTM4B |
| 1863 | LARGE |
| 1864 | LAT |
| 1865 | LAT2 |
| 1866 | LAX1 |
| 1867 | LAYN |
| 1868 | LBP |
| 1869 | LBR |
| 1870 | LCLAT1 |
| 1871 | LCTL |
| 1872 | LDLR |
| 1873 | LDLRAD1 |
| 1874 | LDLRAD3 |
| 1875 | LDLRAD4 |
| 1876 | LECT1 |
| 1877 | LEMD1 |
| 1878 | LEMD3 |
| 1879 | LEPR |
| 1880 | LEPROT |
| 1881 | LEPROTL1 |
| 1882 | LETM1 |
| 1883 | LETM2 |
| 1884 | LETMD1 |
| 1885 | LFNG |
| 1886 | LGALS1 |
| 1887 | LGALS3 |
| 1888 | LGR5 |
| 1889 | LGR6 |
| 1890 | LHCGR |
| 1891 | LHFPL1 |
| 1892 | LHFPL2 |
| 1893 | LHFPL3 |
| 1894 | LHFPL4 |
| 1895 | LHFPL5 |
| 1896 | LIFR |
| 1897 | LIG1 |
| 1898 | LIG3 |
| 1899 | LILRA1 |
| 1900 | LILRA2 |
| 1901 | LILRA4 |
| 1902 | LILRA5 |
| 1903 | LILRA6 |
| 1904 | LILRB1 |
| 1905 | LILRB2 |
| 1906 | LILRB3 |
| 1907 | LILRB4 |

| | |
|---|---|
| 1908 | LILRB5 |
| 1909 | LIM2 |
| 1910 | LIMA1 |
| 1911 | LIME1 |
| 1912 | LINC00116 |
| 1913 | LINGO1 |
| 1914 | LINGO2 |
| 1915 | LINGO3 |
| 1916 | LINGO4 |
| 1917 | LMAN1 |
| 1918 | LMAN1L |
| 1919 | LMAN2 |
| 1920 | LMAN2L |
| 1921 | LMBR1 |
| 1922 | LMBR1L |
| 1923 | LMBRD1 |
| 1924 | LMBRD2 |
| 1925 | LMTK2 |
| 1926 | LMTK3 |
| 1927 | LNPEP |
| 1928 | LPAR1 |
| 1929 | LPAR2 |
| 1930 | LPAR4 |
| 1931 | LPAR5 |
| 1932 | LPAR6 |
| 1933 | LPCAT1 |
| 1934 | LPCAT2 |
| 1935 | LPCAT3 |
| 1936 | LPCAT4 |
| 1937 | LPGAT1 |
| 1938 | LPHN1 |
| 1939 | LPHN2 |
| 1940 | LPL |
| 1941 | LPPR3 |
| 1942 | LPPR4 |
| 1943 | LPPR5 |
| 1944 | LRAT |
| 1945 | LRBA |
| 1946 | LRFN1 |
| 1947 | LRFN2 |
| 1948 | LRFN3 |
| 1949 | LRFN4 |
| 1950 | LRFN5 |
| 1951 | LRIG1 |
| 1952 | LRIG2 |
| 1953 | LRIG3 |
| 1954 | LRIT1 |
| 1955 | LRIT3 |
| 1956 | LRMP |
| 1957 | LRP1 |
| 1958 | LRP10 |
| 1959 | LRP11 |
| 1960 | LRP12 |
| 1961 | LRP1B |
| 1962 | LRP2 |
| 1963 | LRP3 |
| 1964 | LRP4 |
| 1965 | LRP5 |
| 1966 | LRP6 |
| 1967 | LRP8 |
| 1968 | LRPAP1 |
| 1969 | LRRC15 |
| 1970 | LRRC19 |
| 1971 | LRRC24 |
| 1972 | LRRC25 |
| 1973 | LRRC26 |
| 1974 | LRRC32 |
| 1975 | LRRC33 |
| 1976 | LRRC37A |
| 1977 | LRRC37A2 |
| 1978 | LRRC37A3 |
| 1979 | LRRC37B |
| 1980 | LRRC38 |
| 1981 | LRRC4 |
| 1982 | LRRC4B |
| 1983 | LRRC4C |
| 1984 | LRRC55 |
| 1985 | LRRC59 |
| 1986 | LRRC66 |
| 1987 | LRRC70 |
| 1988 | LRRC8A |
| 1989 | LRRC8B |
| 1990 | LRRC8C |

| | |
|---|---|
| 1991 | LRRC8D |
| 1992 | LRRC8E |
| 1993 | LRRN1 |
| 1994 | LRRN2 |
| 1995 | LRRN3 |
| 1996 | LRRN4 |
| 1997 | LRRN4CL |
| 1998 | LRRTM2 |
| 1999 | LRRTM3 |
| 2000 | LRRTM4 |
| 2001 | LRTM1 |
| 2002 | LRTM2 |
| 2003 | LRTOMT |
| 2004 | LSAMP |
| 2005 | LSR |
| 2006 | LST1 |
| 2007 | LTB |
| 2008 | LTB4R |
| 2009 | LTB4R2 |
| 2010 | LTBR |
| 2011 | LTC4S |
| 2012 | LTK |
| 2013 | LY6D |
| 2014 | LY6G6D |
| 2015 | LY6G6F |
| 2016 | LY75 |
| 2017 | LY9 |
| 2018 | LYSMD3 |
| 2019 | LYSMD4 |
| 2020 | LYVE1 |
| 2021 | M6PR |
| 2022 | MADCAM1 |
| 2023 | MAG |
| 2024 | MAL2 |
| 2025 | MAMDC4 |
| 2026 | MAN1A1 |
| 2027 | MAN1A2 |
| 2028 | MAN1B1 |
| 2029 | MAN1C1 |
| 2030 | MAN2A1 |
| 2031 | MAN2A2 |
| 2032 | MANEA |
| 2033 | MANEAL |
| 2034 | MANSC1 |
| 2035 | MANSC4 |
| 2036 | MAOA |
| 2037 | MAOB |
| 2038 | MARCO |
| 2039 | MARVELD1 |
| 2040 | MARVELD2 |
| 2041 | MARVELD3 |
| 2042 | MAS1 |
| 2043 | MAS1L |
| 2044 | MASTL |
| 2045 | MAVS |
| 2046 | MBL2 |
| 2047 | MBOAT1 |
| 2048 | MBOAT2 |
| 2049 | MBOAT4 |
| 2050 | MBOAT7 |
| 2051 | MBTPS1 |
| 2052 | MBTPS2 |
| 2053 | MC1R |
| 2054 | MC3R |
| 2055 | MCAM |
| 2056 | MCHR1 |
| 2057 | MCL1 |
| 2058 | MCOLN1 |
| 2059 | MCOLN2 |
| 2060 | MCOLN3 |
| 2061 | MCTP2 |
| 2062 | MEGF10 |
| 2063 | MEGF11 |
| 2064 | MEGF8 |
| 2065 | MEGF9 |
| 2066 | MEN1 |
| 2067 | MEP1A |
| 2068 | MEP1B |
| 2069 | MERTK |
| 2070 | MEST |
| 2071 | METTL23 |
| 2072 | MFAP3 |
| 2073 | MFF |

| | |
|---|---|
| 2074 | MFGE8 |
| 2075 | MFI2 |
| 2076 | MFN1 |
| 2077 | MFN2 |
| 2078 | MFNG |
| 2079 | MFRP |
| 2080 | MFSD1 |
| 2081 | MFSD10 |
| 2082 | MFSD11 |
| 2083 | MFSD12 |
| 2084 | MFSD2B |
| 2085 | MFSD5 |
| 2086 | MFSD6 |
| 2087 | MFSD6L |
| 2088 | MFSD7 |
| 2089 | MFSD8 |
| 2090 | MFSD9 |
| 2091 | MGA |
| 2092 | MGAM |
| 2093 | MGARP |
| 2094 | MGAT1 |
| 2095 | MGAT2 |
| 2096 | MGAT3 |
| 2097 | MGAT4A |
| 2098 | MGAT4B |
| 2099 | MGAT4C |
| 2100 | MGAT5 |
| 2101 | MGAT5B |
| 2102 | MGST1 |
| 2103 | MGST2 |
| 2104 | MGST3 |
| 2105 | MIA3 |
| 2106 | MICA |
| 2107 | MICB |
| 2108 | MILR1 |
| 2109 | MINOS1 |
| 2110 | MIP |
| 2111 | MIR17HG |
| 2112 | MLANA |
| 2113 | MLNR |
| 2114 | MMD |
| 2115 | MMD2 |
| 2116 | MME |
| 2117 | MMEL1 |
| 2118 | MMP14 |
| 2119 | MMP15 |
| 2120 | MMP16 |
| 2121 | MMP21 |
| 2122 | MMP23A |
| 2123 | MMP24 |
| 2124 | MOG |
| 2125 | MOGAT2 |
| 2126 | MOGS |
| 2127 | MOSPD1 |
| 2128 | MOSPD2 |
| 2129 | MOSPD3 |
| 2130 | MPC1L |
| 2131 | MPDU1 |
| 2132 | MPEG1 |
| 2133 | MPL |
| 2134 | MPV17L2 |
| 2135 | MPZ |
| 2136 | MPZL1 |
| 2137 | MPZL3 |
| 2138 | MRAP |
| 2139 | MRAP2 |
| 2140 | MRC1 |
| 2141 | MRC2 |
| 2142 | MRGPRD |
| 2143 | MRGPRF |
| 2144 | MRGPRG |
| 2145 | MRGPRX1 |
| 2146 | MRGPRX2 |
| 2147 | MRGPRX3 |
| 2148 | MRGPRX4 |
| 2149 | MROH7 |
| 2150 | MRS2 |
| 2151 | MRVI1 |
| 2152 | MS4A1 |
| 2153 | MS4A15 |
| 2154 | MS4A2 |
| 2155 | MS4A3 |
| 2156 | MS4A4A |

| | |
|---|---|
| 2157 | MS4A5 |
| 2158 | MS4A6A |
| 2159 | MS4A6E |
| 2160 | MSLN |
| 2161 | MSMO1 |
| 2162 | MSR1 |
| 2163 | MST1R |
| 2164 | MTCH1 |
| 2165 | MTCH2 |
| 2166 | MTDH |
| 2167 | MTFP1 |
| 2168 | MTNR1B |
| 2169 | MTX1 |
| 2170 | MUC1 |
| 2171 | MUC12 |
| 2172 | MUC13 |
| 2173 | MUC15 |
| 2174 | MUC16 |
| 2175 | MUC17 |
| 2176 | MUC21 |
| 2177 | MUC22 |
| 2178 | MUC4 |
| 2179 | MUL1 |
| 2180 | MUSK |
| 2181 | MXRA7 |
| 2182 | MXRA8 |
| 2183 | MYADM |
| 2184 | MYH10 |
| 2185 | MYH9 |
| 2186 | MYLK |
| 2187 | MYO9A |
| 2188 | MYOF |
| 2189 | MYRF |
| 2190 | NAALAD2 |
| 2191 | NAALADL1 |
| 2192 | NAALADL2 |
| 2193 | NAT14 |
| 2194 | NAT2 |
| 2195 | NAT8 |
| 2196 | NAT8B |
| 2197 | NAT8L |
| 2198 | NCAM1 |
| 2199 | NCAM2 |
| 2200 | NCEH1 |
| 2201 | NCKAP1 |
| 2202 | NCKAP1L |
| 2203 | NCMAP |
| 2204 | NCR1 |
| 2205 | NCR2 |
| 2206 | NCR3 |
| 2207 | NCSTN |
| 2208 | NDE1 |
| 2209 | NDFIP1 |
| 2210 | NDFIP2 |
| 2211 | NDP |
| 2212 | NDST1 |
| 2213 | NDST2 |
| 2214 | NDST3 |
| 2215 | NDST4 |
| 2216 | NDUFA1 |
| 2217 | NDUFA11 |
| 2218 | NDUFA13 |
| 2219 | NDUFA3 |
| 2220 | NDUFB1 |
| 2221 | NDUFB11 |
| 2222 | NDUFB3 |
| 2223 | NDUFB4 |
| 2224 | NDUFB5 |
| 2225 | NDUFB6 |
| 2226 | NDUFB8 |
| 2227 | NDUFC2 |
| 2228 | NEGR1 |
| 2229 | NEO1 |
| 2230 | NETO1 |
| 2231 | NETO2 |
| 2232 | NF2 |
| 2233 | NFASC |
| 2234 | NFE2L1 |
| 2235 | NFXL1 |
| 2236 | NGFR |
| 2237 | NID2 |
| 2238 | NINJ1 |
| 2239 | NINJ2 |

| | |
|---|---|
| 2240 | NIPA1 |
| 2241 | NIPA2 |
| 2242 | NIPAL1 |
| 2243 | NIPAL2 |
| 2244 | NIPAL4 |
| 2245 | NKAIN1 |
| 2246 | NKAIN4 |
| 2247 | NKG7 |
| 2248 | NKPD1 |
| 2249 | NLGN1 |
| 2250 | NLGN2 |
| 2251 | NLGN3 |
| 2252 | NLGN4X |
| 2253 | NLGN4Y |
| 2254 | NMBR |
| 2255 | NMUR1 |
| 2256 | NMUR2 |
| 2257 | NNT |
| 2258 | NOC4L |
| 2259 | NOD2 |
| 2260 | NOMO1 |
| 2261 | NOMO2 |
| 2262 | NOMO3 |
| 2263 | NOTCH1 |
| 2264 | NOTCH2 |
| 2265 | NOTCH3 |
| 2266 | NOTCH4 |
| 2267 | NOV |
| 2268 | NOX1 |
| 2269 | NOX3 |
| 2270 | NOX4 |
| 2271 | NPC1 |
| 2272 | NPC1L1 |
| 2273 | NPDC1 |
| 2274 | NPFFR1 |
| 2275 | NPFFR2 |
| 2276 | NPHS1 |
| 2277 | NPIPL3 |
| 2278 | NPR1 |
| 2279 | NPR2 |
| 2280 | NPR3 |
| 2281 | NPSR1 |
| 2282 | NPTN |
| 2283 | NPTXR |
| 2284 | NPY1R |
| 2285 | NPY2R |
| 2286 | NRCAM |
| 2287 | NRD1 |
| 2288 | NRG1 |
| 2289 | NRG2 |
| 2290 | NRG3 |
| 2291 | NRG4 |
| 2292 | NRP1 |
| 2293 | NRP2 |
| 2294 | NRSN1 |
| 2295 | NRSN2 |
| 2296 | NRXN1 |
| 2297 | NRXN2 |
| 2298 | NRXN3 |
| 2299 | NSDHL |
| 2300 | NSG1 |
| 2301 | NT5E |
| 2302 | NTM |
| 2303 | NTRK1 |
| 2304 | NTRK2 |
| 2305 | NTRK3 |
| 2306 | NUP210L |
| 2307 | NXPE2 |
| 2308 | OCLN |
| 2309 | ODF4 |
| 2310 | OGFOD3 |
| 2311 | OLR1 |
| 2312 | OMA1 |
| 2313 | OPA1 |
| 2314 | OPALIN |
| 2315 | OPCML |
| 2316 | OPN1LW |
| 2317 | OPN1MW |
| 2318 | OPN1SW |
| 2319 | OPN3 |
| 2320 | OPN5 |
| 2321 | OPRK1 |
| 2322 | OR10A3 |

| | |
|---|---|
| 2323 | OR10A5 |
| 2324 | OR10A6 |
| 2325 | OR10A7 |
| 2326 | OR10G2 |
| 2327 | OR10G3 |
| 2328 | OR10G7 |
| 2329 | OR10G8 |
| 2330 | OR10G9 |
| 2331 | OR10H3 |
| 2332 | OR10P1 |
| 2333 | OR10S1 |
| 2334 | OR10V1 |
| 2335 | OR10X1 |
| 2336 | OR11A1 |
| 2337 | OR11H4 |
| 2338 | OR11L1 |
| 2339 | OR12D2 |
| 2340 | OR12D3 |
| 2341 | OR13A1 |
| 2342 | OR13C3 |
| 2343 | OR13C4 |
| 2344 | OR13C8 |
| 2345 | OR13D1 |
| 2346 | OR13F1 |
| 2347 | OR13H1 |
| 2348 | OR14A16 |
| 2349 | OR14I1 |
| 2350 | OR14J1 |
| 2351 | OR1D5 |
| 2352 | OR1E1 |
| 2353 | OR1E2 |
| 2354 | OR1F1 |
| 2355 | OR1F2P |
| 2356 | OR1G1 |
| 2357 | OR1I1 |
| 2358 | OR1J1 |
| 2359 | OR1J2 |
| 2360 | OR1J4 |
| 2361 | OR1L1 |
| 2362 | OR1L3 |
| 2363 | OR1L4 |
| 2364 | OR1L6 |
| 2365 | OR1L8 |
| 2366 | OR1N1 |
| 2367 | OR1N2 |
| 2368 | OR1Q1 |
| 2369 | OR1S1 |
| 2370 | OR1S2 |
| 2371 | OR2A4 |
| 2372 | OR2AE1 |
| 2373 | OR2AG1 |
| 2374 | OR2AG2 |
| 2375 | OR2AK2 |
| 2376 | OR2AP1 |
| 2377 | OR2AT4 |
| 2378 | OR2B2 |
| 2379 | OR2B3 |
| 2380 | OR2B6 |
| 2381 | OR2C1 |
| 2382 | OR2C3 |
| 2383 | OR2D3 |
| 2384 | OR2G2 |
| 2385 | OR2G3 |
| 2386 | OR2G6 |
| 2387 | OR2J2 |
| 2388 | OR2J3 |
| 2389 | OR2L5 |
| 2390 | OR2L8 |
| 2391 | OR2M2 |
| 2392 | OR2M3 |
| 2393 | OR2M5 |
| 2394 | OR2M7 |
| 2395 | OR2T1 |
| 2396 | OR2T11 |
| 2397 | OR2T12 |
| 2398 | OR2T2 |
| 2399 | OR2T27 |
| 2400 | OR2T29 |
| 2401 | OR2T35 |
| 2402 | OR2T4 |
| 2403 | OR2T5 |
| 2404 | OR2T8 |
| 2405 | OR2V1 |

| | |
|---|---|
| 2406 | OR2V2 |
| 2407 | OR2W3 |
| 2408 | OR2Z1 |
| 2409 | OR3A2 |
| 2410 | OR3A3 |
| 2411 | OR4A47 |
| 2412 | OR4C11 |
| 2413 | OR4C13 |
| 2414 | OR4C15 |
| 2415 | OR4C16 |
| 2416 | OR4C3 |
| 2417 | OR4C46 |
| 2418 | OR4C6 |
| 2419 | OR4D10 |
| 2420 | OR4D11 |
| 2421 | OR4D5 |
| 2422 | OR4D9 |
| 2423 | OR4E2 |
| 2424 | OR4F15 |
| 2425 | OR4M1 |
| 2426 | OR4M2 |
| 2427 | OR4N4 |
| 2428 | OR4Q3 |
| 2429 | OR4S2 |
| 2430 | OR4X1 |
| 2431 | OR51A2 |
| 2432 | OR51A4 |
| 2433 | OR51A7 |
| 2434 | OR51B2 |
| 2435 | OR51B4 |
| 2436 | OR51B5 |
| 2437 | OR51B6 |
| 2438 | OR51D1 |
| 2439 | OR51E1 |
| 2440 | OR51E2 |
| 2441 | OR51F1 |
| 2442 | OR51F2 |
| 2443 | OR51G1 |
| 2444 | OR51I1 |
| 2445 | OR51L1 |
| 2446 | OR51M1 |
| 2447 | OR51Q1 |
| 2448 | OR51S1 |
| 2449 | OR51T1 |
| 2450 | OR52A1 |
| 2451 | OR52A5 |
| 2452 | OR52B4 |
| 2453 | OR52B6 |
| 2454 | OR52D1 |
| 2455 | OR52E2 |
| 2456 | OR52E4 |
| 2457 | OR52E6 |
| 2458 | OR52E8 |
| 2459 | OR52I1 |
| 2460 | OR52I2 |
| 2461 | OR52J3 |
| 2462 | OR52K1 |
| 2463 | OR52K2 |
| 2464 | OR52L1 |
| 2465 | OR52M1 |
| 2466 | OR52N1 |
| 2467 | OR52N2 |
| 2468 | OR52N4 |
| 2469 | OR52N5 |
| 2470 | OR52R1 |
| 2471 | OR52W1 |
| 2472 | OR56A1 |
| 2473 | OR56A3 |
| 2474 | OR56A4 |
| 2475 | OR56A5 |
| 2476 | OR56B4 |
| 2477 | OR5A1 |
| 2478 | OR5AK2 |
| 2479 | OR5AP2 |
| 2480 | OR5AR1 |
| 2481 | OR5AS1 |
| 2482 | OR5B12 |
| 2483 | OR5B17 |
| 2484 | OR5B21 |
| 2485 | OR5B3 |
| 2486 | OR5C1 |
| 2487 | OR5D13 |
| 2488 | OR5D14 |

| | |
|---|---|
| 2489 | OR5D16 |
| 2490 | OR5D18 |
| 2491 | OR5F1 |
| 2492 | OR5H1 |
| 2493 | OR5H15 |
| 2494 | OR5I1 |
| 2495 | OR5J2 |
| 2496 | OR5K1 |
| 2497 | OR5K2 |
| 2498 | OR5K3 |
| 2499 | OR5L1 |
| 2500 | OR5L2 |
| 2501 | OR5M10 |
| 2502 | OR5M11 |
| 2503 | OR5M3 |
| 2504 | OR5M8 |
| 2505 | OR5M9 |
| 2506 | OR5P2 |
| 2507 | OR5P3 |
| 2508 | OR5R1 |
| 2509 | OR5T3 |
| 2510 | OR5V1 |
| 2511 | OR6A2 |
| 2512 | OR6B1 |
| 2513 | OR6B2 |
| 2514 | OR6B3 |
| 2515 | OR6C1 |
| 2516 | OR6C2 |
| 2517 | OR6C3 |
| 2518 | OR6C4 |
| 2519 | OR6C6 |
| 2520 | OR6C65 |
| 2521 | OR6C68 |
| 2522 | OR6C70 |
| 2523 | OR6C74 |
| 2524 | OR6C75 |
| 2525 | OR6C76 |
| 2526 | OR6F1 |
| 2527 | OR6K2 |
| 2528 | OR6K3 |
| 2529 | OR6K6 |
| 2530 | OR6M1 |
| 2531 | OR6N2 |
| 2532 | OR6P1 |
| 2533 | OR6Q1 |
| 2534 | OR6S1 |
| 2535 | OR6T1 |
| 2536 | OR6V1 |
| 2537 | OR6X1 |
| 2538 | OR6Y1 |
| 2539 | OR7A10 |
| 2540 | OR7A5 |
| 2541 | OR7C1 |
| 2542 | OR7C2 |
| 2543 | OR7D2 |
| 2544 | OR7E24 |
| 2545 | OR7G2 |
| 2546 | OR7G3 |
| 2547 | OR8A1 |
| 2548 | OR8B2 |
| 2549 | OR8B3 |
| 2550 | OR8B4 |
| 2551 | OR8B8 |
| 2552 | OR8D2 |
| 2553 | OR8G5 |
| 2554 | OR8H1 |
| 2555 | OR8H2 |
| 2556 | OR8H3 |
| 2557 | OR8I2 |
| 2558 | OR8J1 |
| 2559 | OR8J3 |
| 2560 | OR8K1 |
| 2561 | OR8K3 |
| 2562 | OR8K5 |
| 2563 | OR8S1 |
| 2564 | OR8U1 |
| 2565 | OR8U8 |
| 2566 | OR9A2 |
| 2567 | OR9A4 |
| 2568 | OR9G1 |
| 2569 | OR9G4 |
| 2570 | OR9I1 |
| 2571 | OR9Q1 |

| | |
|---|---|
| 2572 | OR9Q2 |
| 2573 | ORAI3 |
| 2574 | ORMDL1 |
| 2575 | ORMDL3 |
| 2576 | OSBPL5 |
| 2577 | OSMR |
| 2578 | OST4 |
| 2579 | OSTM1 |
| 2580 | OTOF |
| 2581 | OTOP2 |
| 2582 | OTOP3 |
| 2583 | OXER1 |
| 2584 | OXGR1 |
| 2585 | OXTR |
| 2586 | P2RX1 |
| 2587 | P2RX2 |
| 2588 | P2RX3 |
| 2589 | P2RX4 |
| 2590 | P2RX5 |
| 2591 | P2RX6 |
| 2592 | P2RX7 |
| 2593 | P2RY1 |
| 2594 | P2RY11 |
| 2595 | P2RY12 |
| 2596 | P2RY13 |
| 2597 | P2RY4 |
| 2598 | P2RY6 |
| 2599 | P4HTM |
| 2600 | PAF1 |
| 2601 | PAG1 |
| 2602 | PAK1 |
| 2603 | PAM |
| 2604 | PANX1 |
| 2605 | PANX2 |
| 2606 | PANX3 |
| 2607 | PAQR3 |
| 2608 | PAQR4 |
| 2609 | PAQR5 |
| 2610 | PAQR6 |
| 2611 | PAQR8 |
| 2612 | PAQR9 |
| 2613 | PAR1 |
| 2614 | PARM1 |
| 2615 | PARP16 |
| 2616 | PCDH1 |
| 2617 | PCDH10 |
| 2618 | PCDH11X |
| 2619 | PCDH11Y |
| 2620 | PCDH12 |
| 2621 | PCDH15 |
| 2622 | PCDH17 |
| 2623 | PCDH18 |
| 2624 | PCDH19 |
| 2625 | PCDH20 |
| 2626 | PCDH7 |
| 2627 | PCDH8 |
| 2628 | PCDH9 |
| 2629 | PCDHA12 |
| 2630 | PCDHA3 |
| 2631 | PCDHAC1 |
| 2632 | PCDHAC2 |
| 2633 | PCDHB1 |
| 2634 | PCDHB10 |
| 2635 | PCDHB11 |
| 2636 | PCDHB12 |
| 2637 | PCDHB13 |
| 2638 | PCDHB14 |
| 2639 | PCDHB15 |
| 2640 | PCDHB16 |
| 2641 | PCDHB18 |
| 2642 | PCDHB2 |
| 2643 | PCDHB3 |
| 2644 | PCDHB4 |
| 2645 | PCDHB5 |
| 2646 | PCDHB6 |
| 2647 | PCDHB7 |
| 2648 | PCDHB8 |
| 2649 | PCDHB9 |
| 2650 | PCDHGA10 |
| 2651 | PCDHGA11 |
| 2652 | PCDHGA12 |
| 2653 | PCDHGA2 |
| 2654 | PCDHGA3 |

| | |
|---|---|
| 2655 | PCDHGA5 |
| 2656 | PCDHGA6 |
| 2657 | PCDHGA7 |
| 2658 | PCDHGA8 |
| 2659 | PCDHGA9 |
| 2660 | PCDHGB1 |
| 2661 | PCDHGB2 |
| 2662 | PCDHGB3 |
| 2663 | PCDHGB4 |
| 2664 | PCDHGB5 |
| 2665 | PCDHGB6 |
| 2666 | PCDHGB7 |
| 2667 | PCDHGC3 |
| 2668 | PCDHGC4 |
| 2669 | PCDHGC5 |
| 2670 | PCNXL2 |
| 2671 | PCNXL3 |
| 2672 | PCNXL4 |
| 2673 | PCP2 |
| 2674 | PCSK5 |
| 2675 | PCSK6 |
| 2676 | PCSK7 |
| 2677 | PCSK9 |
| 2678 | PDCD1 |
| 2679 | PDCD1LG2 |
| 2680 | PDE3A |
| 2681 | PDGFA |
| 2682 | PDGFB |
| 2683 | PDGFC |
| 2684 | PDGFRA |
| 2685 | PDGFRB |
| 2686 | PDIA3 |
| 2687 | PDIA4 |
| 2688 | PDPN |
| 2689 | PDZK1IP1 |
| 2690 | PEAR1 |
| 2691 | PECAM1 |
| 2692 | PEMT |
| 2693 | PERP |
| 2694 | PEX11A |
| 2695 | PEX11B |
| 2696 | PEX11G |
| 2697 | PEX12 |
| 2698 | PEX13 |
| 2699 | PEX16 |
| 2700 | PEX2 |
| 2701 | PEX26 |
| 2702 | PEX3 |
| 2703 | PGAM5 |
| 2704 | PGAP1 |
| 2705 | PGBD5 |
| 2706 | PGRMC1 |
| 2707 | PGRMC2 |
| 2708 | PHEX |
| 2709 | PI16 |
| 2710 | PIEZO1 |
| 2711 | PIEZO2 |
| 2712 | PIGA |
| 2713 | PIGB |
| 2714 | PIGC |
| 2715 | PIGF |
| 2716 | PIGG |
| 2717 | PIGK |
| 2718 | PIGL |
| 2719 | PIGN |
| 2720 | PIGO |
| 2721 | PIGQ |
| 2722 | PIGR |
| 2723 | PIGS |
| 2724 | PIGT |
| 2725 | PIGX |
| 2726 | PIGZ |
| 2727 | PIK3IP1 |
| 2728 | PILRA |
| 2729 | PILRB |
| 2730 | PINK1 |
| 2731 | PIRT |
| 2732 | PITPNM1 |
| 2733 | PKD1 |
| 2734 | PKD1L1 |
| 2735 | PKD1L2 |
| 2736 | PKD1L3 |
| 2737 | PKD2 |

| | |
|---|---|
| 2738 | PKD2L2 |
| 2739 | PKDREJ |
| 2740 | PKHD1 |
| 2741 | PKHD1L1 |
| 2742 | PKN1 |
| 2743 | PKN2 |
| 2744 | PLA2G16 |
| 2745 | PLA2G1B |
| 2746 | PLA2G5 |
| 2747 | PLA2R1 |
| 2748 | PLAT |
| 2749 | PLAU |
| 2750 | PLB1 |
| 2751 | PLCD3 |
| 2752 | PLD3 |
| 2753 | PLD4 |
| 2754 | PLD5 |
| 2755 | PLD6 |
| 2756 | PLEKHG6 |
| 2757 | PLG |
| 2758 | PLK4 |
| 2759 | PLLP |
| 2760 | PLN |
| 2761 | PLP1 |
| 2762 | PLP2 |
| 2763 | PLSCR1 |
| 2764 | PLSCR2 |
| 2765 | PLSCR3 |
| 2766 | PLSCR4 |
| 2767 | PLVAP |
| 2768 | PLXDC1 |
| 2769 | PLXDC2 |
| 2770 | PLXNA1 |
| 2771 | PLXNA2 |
| 2772 | PLXNA3 |
| 2773 | PLXNA4 |
| 2774 | PLXNB1 |
| 2775 | PLXNB2 |
| 2776 | PLXNB3 |
| 2777 | PLXNC1 |
| 2778 | PLXND1 |
| 2779 | PMEL |
| 2780 | PMEPA1 |
| 2781 | PNLDC1 |
| 2782 | PNPLA2 |
| 2783 | PNPLA3 |
| 2784 | PNPLA6 |
| 2785 | PNPLA7 |
| 2786 | PNPLA8 |
| 2787 | PODXL |
| 2788 | PODXL2 |
| 2789 | POM121 |
| 2790 | POM121C |
| 2791 | POMGNT1 |
| 2792 | POMT1 |
| 2793 | POMT2 |
| 2794 | POPDC2 |
| 2795 | PORCN |
| 2796 | PPAP2A |
| 2797 | PPAP2B |
| 2798 | PPAPDC1A |
| 2799 | PPAPDC1B |
| 2800 | PPFIA2 |
| 2801 | PPFIA3 |
| 2802 | PPFIA4 |
| 2803 | PPM1L |
| 2804 | PPP1CC |
| 2805 | PPP1R3A |
| 2806 | PPP1R3F |
| 2807 | PPYR1 |
| 2808 | PQLC1 |
| 2809 | PQLC2 |
| 2810 | PQLC3 |
| 2811 | PRCD |
| 2812 | PREB |
| 2813 | PRIMA1 |
| 2814 | PRLR |
| 2815 | PRNP |
| 2816 | PROCR |
| 2817 | PROKR1 |
| 2818 | PROKR2 |
| 2819 | PROM1 |
| 2820 | PROM2 |

| | |
|---|---|
| 2821 | PRPH |
| 2822 | PRPH2 |
| 2823 | PRR3 |
| 2824 | PRR4 |
| 2825 | PRR7 |
| 2826 | PRRG1 |
| 2827 | PRRG2 |
| 2828 | PRRG3 |
| 2829 | PRRG4 |
| 2830 | PRRT1 |
| 2831 | PRRT2 |
| 2832 | PRRT4 |
| 2833 | PRSS8 |
| 2834 | PRTG |
| 2835 | PSD2 |
| 2836 | PSEN1 |
| 2837 | PSEN2 |
| 2838 | PSENEN |
| 2839 | PSTPIP1 |
| 2840 | PTCH1 |
| 2841 | PTCH2 |
| 2842 | PTCHD1 |
| 2843 | PTCHD2 |
| 2844 | PTCHD3 |
| 2845 | PTCHD4 |
| 2846 | PTCRA |
| 2847 | PTDSS1 |
| 2848 | PTDSS2 |
| 2849 | PTGDR2 |
| 2850 | PTGER3 |
| 2851 | PTGES |
| 2852 | PTGES2 |
| 2853 | PTGFR |
| 2854 | PTGFRN |
| 2855 | PTH1R |
| 2856 | PTK7 |
| 2857 | PTPLAD1 |
| 2858 | PTPLAD2 |
| 2859 | PTPRA |
| 2860 | PTPRB |
| 2861 | PTPRC |
| 2862 | PTPRCAP |
| 2863 | PTPRD |
| 2864 | PTPRE |
| 2865 | PTPRF |
| 2866 | PTPRG |
| 2867 | PTPRH |
| 2868 | PTPRJ |
| 2869 | PTPRK |
| 2870 | PTPRM |
| 2871 | PTPRN |
| 2872 | PTPRN2 |
| 2873 | PTPRO |
| 2874 | PTPRQ |
| 2875 | PTPRR |
| 2876 | PTPRS |
| 2877 | PTPRT |
| 2878 | PTPRU |
| 2879 | PTPRZ1 |
| 2880 | PTTG1IP |
| 2881 | PVR |
| 2882 | PVRL1 |
| 2883 | PVRL2 |
| 2884 | PVRL3 |
| 2885 | PVRL4 |
| 2886 | QPCTL |
| 2887 | QRFPR |
| 2888 | QSOX1 |
| 2889 | RAB11A |
| 2890 | RAB11FIP3 |
| 2891 | RAB11FIP4 |
| 2892 | RAB21 |
| 2893 | RACGAP1 |
| 2894 | RAET1E |
| 2895 | RAET1G |
| 2896 | RALA |
| 2897 | RAMP1 |
| 2898 | RAMP2 |
| 2899 | RAMP3 |
| 2900 | RARA |
| 2901 | RARRES1 |
| 2902 | RARRES3 |
| 2903 | RC3H2 |

| | |
|---|---|
| 2904 | RDH10 |
| 2905 | RDH11 |
| 2906 | RDH16 |
| 2907 | RDH8 |
| 2908 | RDX |
| 2909 | REEP2 |
| 2910 | REEP4 |
| 2911 | REEP5 |
| 2912 | REEP6 |
| 2913 | RELL1 |
| 2914 | RELL2 |
| 2915 | RELT |
| 2916 | RET |
| 2917 | RFC1 |
| 2918 | RFNG |
| 2919 | RFT1 |
| 2920 | RGMA |
| 2921 | RGS9BP |
| 2922 | RGSL1 |
| 2923 | RHAG |
| 2924 | RHBDD1 |
| 2925 | RHBDD3 |
| 2926 | RHBDF1 |
| 2927 | RHBDF2 |
| 2928 | RHBDL1 |
| 2929 | RHBDL3 |
| 2930 | RHBG |
| 2931 | RHCE |
| 2932 | RHCG |
| 2933 | RHD |
| 2934 | RHO |
| 2935 | RHOA |
| 2936 | RHOB |
| 2937 | RHOC |
| 2938 | RHOT1 |
| 2939 | RHOT2 |
| 2940 | RIC3 |
| 2941 | RMDN2 |
| 2942 | RMDN3 |
| 2943 | RNASEK |
| 2944 | RNF112 |
| 2945 | RNF121 |
| 2946 | RNF122 |
| 2947 | RNF128 |
| 2948 | RNF13 |
| 2949 | RNF130 |
| 2950 | RNF133 |
| 2951 | RNF144A |
| 2952 | RNF144B |
| 2953 | RNF148 |
| 2954 | RNF149 |
| 2955 | RNF150 |
| 2956 | RNF152 |
| 2957 | RNF167 |
| 2958 | RNF170 |
| 2959 | RNF175 |
| 2960 | RNF180 |
| 2961 | RNF182 |
| 2962 | RNF183 |
| 2963 | RNF185 |
| 2964 | RNF186 |
| 2965 | RNF19A |
| 2966 | RNF19B |
| 2967 | RNF217 |
| 2968 | RNF222 |
| 2969 | RNF223 |
| 2970 | RNF24 |
| 2971 | RNF26 |
| 2972 | RNF43 |
| 2973 | RNFT1 |
| 2974 | RNFT2 |
| 2975 | ROBO1 |
| 2976 | ROBO2 |
| 2977 | ROBO3 |
| 2978 | ROBO4 |
| 2979 | ROMO1 |
| 2980 | ROR1 |
| 2981 | ROR2 |
| 2982 | ROS1 |
| 2983 | RPN1 |
| 2984 | RPN2 |
| 2985 | RPRM |
| 2986 | RPS6KB1 |

| | |
|---|---|
| 2987 | RRBP1 |
| 2988 | RRH |
| 2989 | RRNAD1 |
| 2990 | RRP12 |
| 2991 | RSPO2 |
| 2992 | RTN2 |
| 2993 | RTN4 |
| 2994 | RTN4R |
| 2995 | RTN4RL1 |
| 2996 | RTN4RL2 |
| 2997 | RTP1 |
| 2998 | RTP2 |
| 2999 | RTP4 |
| 3000 | RYK |
| 3001 | RYR1 |
| 3002 | RYR2 |
| 3003 | S1PR1 |
| 3004 | S1PR3 |
| 3005 | SACM1L |
| 3006 | SAMD8 |
| 3007 | SAYSD1 |
| 3008 | SC5DL |
| 3009 | SCAI |
| 3010 | SCAMP1 |
| 3011 | SCAMP2 |
| 3012 | SCAMP3 |
| 3013 | SCAMP4 |
| 3014 | SCAMP5 |
| 3015 | SCARA3 |
| 3016 | SCARA5 |
| 3017 | SCARB1 |
| 3018 | SCARB2 |
| 3019 | SCARF1 |
| 3020 | SCARF2 |
| 3021 | SCD5 |
| 3022 | SCIMP |
| 3023 | SCN10A |
| 3024 | SCN11A |
| 3025 | SCN1A |
| 3026 | SCN1B |
| 3027 | SCN2A |
| 3028 | SCN2B |
| 3029 | SCN3A |
| 3030 | SCN3B |
| 3031 | SCN4A |
| 3032 | SCN4B |
| 3033 | SCN5A |
| 3034 | SCN7A |
| 3035 | SCN8A |
| 3036 | SCN9A |
| 3037 | SCNN1A |
| 3038 | SCNN1B |
| 3039 | SCNN1G |
| 3040 | SCUBE1 |
| 3041 | SCUBE3 |
| 3042 | SDC1 |
| 3043 | SDC2 |
| 3044 | SDC3 |
| 3045 | SDC4 |
| 3046 | SDHC |
| 3047 | SDK1 |
| 3048 | SDK2 |
| 3049 | SDR16C5 |
| 3050 | SEC11A |
| 3051 | SEC11C |
| 3052 | SEC22B |
| 3053 | SEC22C |
| 3054 | SEC61A1 |
| 3055 | SEC61A2 |
| 3056 | SEC61B |
| 3057 | SEC61G |
| 3058 | SEC62 |
| 3059 | SEC63 |
| 3060 | SECTM1 |
| 3061 | SEL1L |
| 3062 | SEL1L3 |
| 3063 | SELE |
| 3064 | SELK |
| 3065 | SELL |
| 3066 | SELPLG |
| 3067 | SEMA4C |
| 3068 | SEMA4D |
| 3069 | SEMA4F |

| | |
|---|---|
| 3070 | SEMA4G |
| 3071 | SEMA5A |
| 3072 | SEMA5B |
| 3073 | SEMA6A |
| 3074 | SEMA6B |
| 3075 | SEMA6D |
| 3076 | SEMA7A |
| 3077 | SERAC1 |
| 3078 | SERINC1 |
| 3079 | SERINC2 |
| 3080 | SERINC4 |
| 3081 | SERINC5 |
| 3082 | SERP1 |
| 3083 | SERP2 |
| 3084 | SERPINA5 |
| 3085 | SERPINE2 |
| 3086 | SERPINF2 |
| 3087 | SERTM1 |
| 3088 | SEZ6 |
| 3089 | SEZ6L |
| 3090 | SEZ6L2 |
| 3091 | SFRP1 |
| 3092 | SFRP4 |
| 3093 | SFT2D1 |
| 3094 | SFT2D2 |
| 3095 | SFT2D3 |
| 3096 | SFXN1 |
| 3097 | SFXN4 |
| 3098 | SGCA |
| 3099 | SGCB |
| 3100 | SGCD |
| 3101 | SGCE |
| 3102 | SGCG |
| 3103 | SGCZ |
| 3104 | SGK196 |
| 3105 | SGPL1 |
| 3106 | SGPP1 |
| 3107 | SGPP2 |
| 3108 | SHH |
| 3109 | SHISA2 |
| 3110 | SHISA3 |
| 3111 | SHISA4 |
| 3112 | SHISA5 |
| 3113 | SHISA9 |
| 3114 | SI |
| 3115 | SIGIRR |
| 3116 | SIGLEC1 |
| 3117 | SIGLEC10 |
| 3118 | SIGLEC11 |
| 3119 | SIGLEC12 |
| 3120 | SIGLEC14 |
| 3121 | SIGLEC15 |
| 3122 | SIGLEC16 |
| 3123 | SIGLEC5 |
| 3124 | SIGLEC6 |
| 3125 | SIGLEC7 |
| 3126 | SIGLEC8 |
| 3127 | SIGLEC9 |
| 3128 | SIRPA |
| 3129 | SIRPB1 |
| 3130 | SIRPB2 |
| 3131 | SIRPG |
| 3132 | SIT1 |
| 3133 | SKINTL |
| 3134 | SLAMF1 |
| 3135 | SLAMF6 |
| 3136 | SLAMF7 |
| 3137 | SLAMF8 |
| 3138 | SLAMF9 |
| 3139 | SLC10A1 |
| 3140 | SLC10A2 |
| 3141 | SLC10A3 |
| 3142 | SLC10A4 |
| 3143 | SLC10A5 |
| 3144 | SLC10A6 |
| 3145 | SLC11A1 |
| 3146 | SLC11A2 |
| 3147 | SLC12A1 |
| 3148 | SLC12A2 |
| 3149 | SLC12A3 |
| 3150 | SLC12A6 |
| 3151 | SLC12A8 |
| 3152 | SLC12A9 |

| | |
|---|---|
| 3153 | SLC13A2 |
| 3154 | SLC13A3 |
| 3155 | SLC13A4 |
| 3156 | SLC13A5 |
| 3157 | SLC15A2 |
| 3158 | SLC15A3 |
| 3159 | SLC15A4 |
| 3160 | SLC15A5 |
| 3161 | SLC16A1 |
| 3162 | SLC16A10 |
| 3163 | SLC16A11 |
| 3164 | SLC16A12 |
| 3165 | SLC16A13 |
| 3166 | SLC16A14 |
| 3167 | SLC16A5 |
| 3168 | SLC16A9 |
| 3169 | SLC17A1 |
| 3170 | SLC17A2 |
| 3171 | SLC17A3 |
| 3172 | SLC17A4 |
| 3173 | SLC17A6 |
| 3174 | SLC17A7 |
| 3175 | SLC17A8 |
| 3176 | SLC17A9 |
| 3177 | SLC18A1 |
| 3178 | SLC18A2 |
| 3179 | SLC18B1 |
| 3180 | SLC19A1 |
| 3181 | SLC19A3 |
| 3182 | SLC1A1 |
| 3183 | SLC1A2 |
| 3184 | SLC1A3 |
| 3185 | SLC1A4 |
| 3186 | SLC1A5 |
| 3187 | SLC1A7 |
| 3188 | SLC20A1 |
| 3189 | SLC20A2 |
| 3190 | SLC22A1 |
| 3191 | SLC22A11 |
| 3192 | SLC22A12 |
| 3193 | SLC22A13 |
| 3194 | SLC22A16 |
| 3195 | SLC22A17 |
| 3196 | SLC22A2 |
| 3197 | SLC22A23 |
| 3198 | SLC22A24 |
| 3199 | SLC22A3 |
| 3200 | SLC22A31 |
| 3201 | SLC22A5 |
| 3202 | SLC22A6 |
| 3203 | SLC22A8 |
| 3204 | SLC23A3 |
| 3205 | SLC24A1 |
| 3206 | SLC24A2 |
| 3207 | SLC24A4 |
| 3208 | SLC24A5 |
| 3209 | SLC24A6 |
| 3210 | SLC25A1 |
| 3211 | SLC25A10 |
| 3212 | SLC25A11 |
| 3213 | SLC25A15 |
| 3214 | SLC25A16 |
| 3215 | SLC25A17 |
| 3216 | SLC25A18 |
| 3217 | SLC25A2 |
| 3218 | SLC25A20 |
| 3219 | SLC25A22 |
| 3220 | SLC25A25 |
| 3221 | SLC25A3 |
| 3222 | SLC25A32 |
| 3223 | SLC25A39 |
| 3224 | SLC25A4 |
| 3225 | SLC25A40 |
| 3226 | SLC25A41 |
| 3227 | SLC25A43 |
| 3228 | SLC25A46 |
| 3229 | SLC25A47 |
| 3230 | SLC25A48 |
| 3231 | SLC25A5 |
| 3232 | SLC25A51 |
| 3233 | SLC25A52 |
| 3234 | SLC25A53 |
| 3235 | SLC25A6 |

| | |
|---|---|
| 3236 | SLC26A1 |
| 3237 | SLC26A10 |
| 3238 | SLC26A11 |
| 3239 | SLC26A2 |
| 3240 | SLC26A3 |
| 3241 | SLC26A4 |
| 3242 | SLC26A5 |
| 3243 | SLC26A6 |
| 3244 | SLC26A7 |
| 3245 | SLC26A8 |
| 3246 | SLC26A9 |
| 3247 | SLC27A1 |
| 3248 | SLC27A2 |
| 3249 | SLC27A3 |
| 3250 | SLC27A4 |
| 3251 | SLC28A1 |
| 3252 | SLC28A2 |
| 3253 | SLC28A3 |
| 3254 | SLC2A1 |
| 3255 | SLC2A13 |
| 3256 | SLC2A14 |
| 3257 | SLC2A2 |
| 3258 | SLC2A3 |
| 3259 | SLC2A4 |
| 3260 | SLC2A5 |
| 3261 | SLC2A6 |
| 3262 | SLC2A7 |
| 3263 | SLC2A8 |
| 3264 | SLC2A9 |
| 3265 | SLC30A1 |
| 3266 | SLC30A10 |
| 3267 | SLC30A3 |
| 3268 | SLC30A5 |
| 3269 | SLC30A6 |
| 3270 | SLC30A7 |
| 3271 | SLC30A8 |
| 3272 | SLC30A9 |
| 3273 | SLC31A1 |
| 3274 | SLC31A2 |
| 3275 | SLC34A1 |
| 3276 | SLC34A2 |
| 3277 | SLC34A3 |
| 3278 | SLC35A1 |
| 3279 | SLC35A2 |
| 3280 | SLC35B2 |
| 3281 | SLC35B3 |
| 3282 | SLC35C1 |
| 3283 | SLC35C2 |
| 3284 | SLC35D1 |
| 3285 | SLC35D2 |
| 3286 | SLC35E2 |
| 3287 | SLC35E2B |
| 3288 | SLC35F1 |
| 3289 | SLC35F2 |
| 3290 | SLC35F3 |
| 3291 | SLC35F5 |
| 3292 | SLC35F6 |
| 3293 | SLC35G2 |
| 3294 | SLC35G3 |
| 3295 | SLC35G5 |
| 3296 | SLC35G6 |
| 3297 | SLC36A2 |
| 3298 | SLC36A3 |
| 3299 | SLC37A2 |
| 3300 | SLC37A3 |
| 3301 | SLC38A1 |
| 3302 | SLC38A11 |
| 3303 | SLC38A2 |
| 3304 | SLC38A3 |
| 3305 | SLC38A4 |
| 3306 | SLC38A6 |
| 3307 | SLC39A1 |
| 3308 | SLC39A10 |
| 3309 | SLC39A11 |
| 3310 | SLC39A14 |
| 3311 | SLC39A3 |
| 3312 | SLC39A4 |
| 3313 | SLC39A5 |
| 3314 | SLC39A6 |
| 3315 | SLC39A7 |
| 3316 | SLC39A9 |
| 3317 | SLC3A1 |
| 3318 | SLC3A2 |

| | |
|---|---|
| 3319 | SLC40A1 |
| 3320 | SLC41A1 |
| 3321 | SLC41A2 |
| 3322 | SLC43A2 |
| 3323 | SLC44A1 |
| 3324 | SLC44A2 |
| 3325 | SLC44A4 |
| 3326 | SLC44A5 |
| 3327 | SLC45A1 |
| 3328 | SLC45A2 |
| 3329 | SLC46A1 |
| 3330 | SLC46A3 |
| 3331 | SLC47A1 |
| 3332 | SLC47A2 |
| 3333 | SLC4A1 |
| 3334 | SLC4A10 |
| 3335 | SLC4A2 |
| 3336 | SLC4A3 |
| 3337 | SLC4A4 |
| 3338 | SLC4A5 |
| 3339 | SLC4A9 |
| 3340 | SLC51A |
| 3341 | SLC51B |
| 3342 | SLC52A1 |
| 3343 | SLC52A2 |
| 3344 | SLC52A3 |
| 3345 | SLC5A1 |
| 3346 | SLC5A10 |
| 3347 | SLC5A12 |
| 3348 | SLC5A2 |
| 3349 | SLC5A3 |
| 3350 | SLC5A4 |
| 3351 | SLC5A5 |
| 3352 | SLC5A6 |
| 3353 | SLC5A7 |
| 3354 | SLC5A8 |
| 3355 | SLC5A9 |
| 3356 | SLC6A1 |
| 3357 | SLC6A11 |
| 3358 | SLC6A12 |
| 3359 | SLC6A13 |
| 3360 | SLC6A14 |
| 3361 | SLC6A15 |
| 3362 | SLC6A16 |
| 3363 | SLC6A17 |
| 3364 | SLC6A18 |
| 3365 | SLC6A19 |
| 3366 | SLC6A4 |
| 3367 | SLC6A6 |
| 3368 | SLC6A7 |
| 3369 | SLC6A8 |
| 3370 | SLC6A9 |
| 3371 | SLC7A1 |
| 3372 | SLC7A10 |
| 3373 | SLC7A11 |
| 3374 | SLC7A13 |
| 3375 | SLC7A14 |
| 3376 | SLC7A2 |
| 3377 | SLC7A3 |
| 3378 | SLC7A4 |
| 3379 | SLC7A5 |
| 3380 | SLC7A5P2 |
| 3381 | SLC7A6 |
| 3382 | SLC7A7 |
| 3383 | SLC7A8 |
| 3384 | SLC8A3 |
| 3385 | SLC9A1 |
| 3386 | SLC9A2 |
| 3387 | SLC9A3 |
| 3388 | SLC9A4 |
| 3389 | SLC9A5 |
| 3390 | SLC9A6 |
| 3391 | SLC9A7 |
| 3392 | SLC9A8 |
| 3393 | SLC9A9 |
| 3394 | SLC9B1 |
| 3395 | SLC9B2 |
| 3396 | SLC9C2 |
| 3397 | SLCO1A2 |
| 3398 | SLCO1B1 |
| 3399 | SLCO1B7 |
| 3400 | SLCO1C1 |
| 3401 | SLCO3A1 |

| | |
|---|---|
| 3402 | SLCO4A1 |
| 3403 | SLCO4C1 |
| 3404 | SLCO5A1 |
| 3405 | SLCO6A1 |
| 3406 | SLFN12L |
| 3407 | SLFN5 |
| 3408 | SLIT2 |
| 3409 | SLITRK1 |
| 3410 | SLITRK2 |
| 3411 | SLITRK3 |
| 3412 | SLITRK4 |
| 3413 | SLITRK5 |
| 3414 | SLITRK6 |
| 3415 | SLMAP |
| 3416 | SLN |
| 3417 | SMAGP |
| 3418 | SMCR7 |
| 3419 | SMCR7L |
| 3420 | SMIM1 |
| 3421 | SMIM11 |
| 3422 | SMIM12 |
| 3423 | SMIM14 |
| 3424 | SMIM15 |
| 3425 | SMIM9 |
| 3426 | SMLR1 |
| 3427 | SMO |
| 3428 | SMPD4 |
| 3429 | SNN |
| 3430 | SNPH |
| 3431 | SNX14 |
| 3432 | SOGA3 |
| 3433 | SORCS1 |
| 3434 | SORCS2 |
| 3435 | SORL1 |
| 3436 | SORT1 |
| 3437 | SPACA1 |
| 3438 | SPACA3 |
| 3439 | SPARC |
| 3440 | SPAST |
| 3441 | SPATA25 |
| 3442 | SPATA31A1 |
| 3443 | SPATA31A2 |
| 3444 | SPATA31A3 |
| 3445 | SPATA31A4 |
| 3446 | SPATA31A5 |
| 3447 | SPATA31A6 |
| 3448 | SPATA31A7 |
| 3449 | SPATA31C1 |
| 3450 | SPATA31C2 |
| 3451 | SPATA31D1 |
| 3452 | SPATA31D3 |
| 3453 | SPATA31D4 |
| 3454 | SPATA31E1 |
| 3455 | SPATA9 |
| 3456 | SPC25 |
| 3457 | SPCS1 |
| 3458 | SPCS2 |
| 3459 | SPCS3 |
| 3460 | SPEM1 |
| 3461 | SPG11 |
| 3462 | SPG7 |
| 3463 | SPIN1 |
| 3464 | SPINT2 |
| 3465 | SPIRE2 |
| 3466 | SPN |
| 3467 | SPNS1 |
| 3468 | SPNS3 |
| 3469 | SPPL2A |
| 3470 | SPPL3 |
| 3471 | SPTB |
| 3472 | SPTLC1 |
| 3473 | SPTLC2 |
| 3474 | SPTLC3 |
| 3475 | SPTSSA |
| 3476 | SPTSSB |
| 3477 | SQLE |
| 3478 | SRD5A1 |
| 3479 | SRD5A3 |
| 3480 | SREBF1 |
| 3481 | SREBF2 |
| 3482 | SRPRB |
| 3483 | SRPX |
| 3484 | SRPX2 |

| | |
|---|---|
| 3485 | SSH1 |
| 3486 | SSPN |
| 3487 | SSR1 |
| 3488 | SSR2 |
| 3489 | SSR4 |
| 3490 | SSTR1 |
| 3491 | SSTR3 |
| 3492 | SSTR4 |
| 3493 | SSTR5 |
| 3494 | ST14 |
| 3495 | ST3GAL1 |
| 3496 | ST3GAL2 |
| 3497 | ST3GAL3 |
| 3498 | ST3GAL4 |
| 3499 | ST3GAL5 |
| 3500 | ST3GAL6 |
| 3501 | ST6GAL1 |
| 3502 | ST6GAL2 |
| 3503 | ST6GALNAC1 |
| 3504 | ST6GALNAC2 |
| 3505 | ST6GALNAC3 |
| 3506 | ST6GALNAC4 |
| 3507 | ST6GALNAC5 |
| 3508 | ST6GALNAC6 |
| 3509 | ST7L |
| 3510 | ST8SIA1 |
| 3511 | ST8SIA2 |
| 3512 | ST8SIA3 |
| 3513 | ST8SIA4 |
| 3514 | ST8SIA5 |
| 3515 | ST8SIA6 |
| 3516 | STAB1 |
| 3517 | STAB2 |
| 3518 | STAMBP |
| 3519 | STARD3 |
| 3520 | STBD1 |
| 3521 | STEAP1B |
| 3522 | STEAP3 |
| 3523 | STEAP4 |
| 3524 | STIM1 |
| 3525 | STOML1 |
| 3526 | STOML3 |
| 3527 | STRC |
| 3528 | STS |
| 3529 | STT3A |
| 3530 | STT3B |
| 3531 | STX10 |
| 3532 | STX12 |
| 3533 | STX16 |
| 3534 | STX17 |
| 3535 | STX18 |
| 3536 | STX1A |
| 3537 | STX1B |
| 3538 | STX2 |
| 3539 | STX3 |
| 3540 | STX4 |
| 3541 | STX5 |
| 3542 | STX6 |
| 3543 | STX7 |
| 3544 | STX8 |
| 3545 | STYK1 |
| 3546 | SUCNR1 |
| 3547 | SULF1 |
| 3548 | SULF2 |
| 3549 | SUN1 |
| 3550 | SUN2 |
| 3551 | SUN3 |
| 3552 | SUN5 |
| 3553 | SURF4 |
| 3554 | SUSD1 |
| 3555 | SUSD2 |
| 3556 | SUSD3 |
| 3557 | SUSD4 |
| 3558 | SUSD5 |
| 3559 | SV2A |
| 3560 | SV2C |
| 3561 | SVIL |
| 3562 | SVOP |
| 3563 | SVOPL |
| 3564 | SYBU |
| 3565 | SYNDIG1 |
| 3566 | SYNE1 |
| 3567 | SYNE2 |

| | |
|---|---|
| 3568 | SYNE3 |
| 3569 | SYNE4 |
| 3570 | SYNGR2 |
| 3571 | SYNJ2BP |
| 3572 | SYNPR |
| 3573 | SYP |
| 3574 | SYPL1 |
| 3575 | SYPL2 |
| 3576 | SYS1 |
| 3577 | SYT1 |
| 3578 | SYT10 |
| 3579 | SYT11 |
| 3580 | SYT12 |
| 3581 | SYT13 |
| 3582 | SYT14 |
| 3583 | SYT15 |
| 3584 | SYT2 |
| 3585 | SYT3 |
| 3586 | SYT4 |
| 3587 | SYT5 |
| 3588 | SYT6 |
| 3589 | SYT7 |
| 3590 | SYT8 |
| 3591 | SYT9 |
| 3592 | TAAR1 |
| 3593 | TAAR3 |
| 3594 | TAAR8 |
| 3595 | TAAR9 |
| 3596 | TACR1 |
| 3597 | TACR2 |
| 3598 | TACSTD2 |
| 3599 | TANGO6 |
| 3600 | TAOK2 |
| 3601 | TAP1 |
| 3602 | TAPBP |
| 3603 | TAPT1 |
| 3604 | TARM1 |
| 3605 | TAS1R1 |
| 3606 | TAS1R2 |
| 3607 | TAS1R3 |
| 3608 | TAS2R1 |
| 3609 | TAS2R13 |
| 3610 | TAS2R14 |
| 3611 | TAS2R16 |
| 3612 | TAS2R19 |
| 3613 | TAS2R20 |
| 3614 | TAS2R3 |
| 3615 | TAS2R30 |
| 3616 | TAS2R31 |
| 3617 | TAS2R38 |
| 3618 | TAS2R39 |
| 3619 | TAS2R4 |
| 3620 | TAS2R40 |
| 3621 | TAS2R42 |
| 3622 | TAS2R43 |
| 3623 | TAS2R46 |
| 3624 | TAS2R50 |
| 3625 | TAS2R7 |
| 3626 | TAS2R8 |
| 3627 | TAS2R9 |
| 3628 | TBC1D20 |
| 3629 | TBC1D9B |
| 3630 | TBXA2R |
| 3631 | TBXAS1 |
| 3632 | TCIRG1 |
| 3633 | TCP11 |
| 3634 | TCTA |
| 3635 | TCTN2 |
| 3636 | TCTN3 |
| 3637 | TDGF1 |
| 3638 | TECR |
| 3639 | TEK |
| 3640 | TENM1 |
| 3641 | TENM2 |
| 3642 | TENM3 |
| 3643 | TENM4 |
| 3644 | TEX10 |
| 3645 | TEX261 |
| 3646 | TEX28 |
| 3647 | TEX29 |
| 3648 | TEX38 |
| 3649 | TF |
| 3650 | TFPI |

| | |
|---|---|
| 3651 | TFR2 |
| 3652 | TFRC |
| 3653 | TGFA |
| 3654 | TGFB1 |
| 3655 | TGFB3 |
| 3656 | TGFBR1 |
| 3657 | TGFBR2 |
| 3658 | TGFBR3 |
| 3659 | TGFBR3L |
| 3660 | TGOLN2 |
| 3661 | THBD |
| 3662 | THBS1 |
| 3663 | THSD1 |
| 3664 | THSD7A |
| 3665 | THSD7B |
| 3666 | THY1 |
| 3667 | TIE1 |
| 3668 | TIGIT |
| 3669 | TIMD4 |
| 3670 | TIMM17A |
| 3671 | TIMM17B |
| 3672 | TIMM21 |
| 3673 | TIMM22 |
| 3674 | TIMM50 |
| 3675 | TIMMDC1 |
| 3676 | TIMP2 |
| 3677 | TLCD1 |
| 3678 | TLCD2 |
| 3679 | TLN1 |
| 3680 | TLR1 |
| 3681 | TLR10 |
| 3682 | TLR2 |
| 3683 | TLR3 |
| 3684 | TLR4 |
| 3685 | TLR6 |
| 3686 | TLR7 |
| 3687 | TLR8 |
| 3688 | TLR9 |
| 3689 | TM2D1 |
| 3690 | TM2D2 |
| 3691 | TM2D3 |
| 3692 | TM4SF1 |
| 3693 | TM4SF18 |
| 3694 | TM4SF4 |
| 3695 | TM4SF5 |
| 3696 | TM6SF1 |
| 3697 | TM7SF2 |
| 3698 | TM7SF3 |
| 3699 | TM9SF1 |
| 3700 | TM9SF2 |
| 3701 | TM9SF3 |
| 3702 | TM9SF4 |
| 3703 | TMBIM1 |
| 3704 | TMBIM4 |
| 3705 | TMBIM6 |
| 3706 | TMC2 |
| 3707 | TMC3 |
| 3708 | TMC4 |
| 3709 | TMC5 |
| 3710 | TMC6 |
| 3711 | TMC7 |
| 3712 | TMC8 |
| 3713 | TMCC1 |
| 3714 | TMCO1 |
| 3715 | TMCO2 |
| 3716 | TMCO3 |
| 3717 | TMCO4 |
| 3718 | TMCO5A |
| 3719 | TMCO5B |
| 3720 | TMCO6 |
| 3721 | TMED1 |
| 3722 | TMED10 |
| 3723 | TMED2 |
| 3724 | TMED4 |
| 3725 | TMED6 |
| 3726 | TMED7 |
| 3727 | TMED9 |
| 3728 | TMEFF1 |
| 3729 | TMEFF2 |
| 3730 | TMEM100 |
| 3731 | TMEM101 |
| 3732 | TMEM102 |
| 3733 | TMEM104 |

| | |
|---|---|
| 3734 | TMEM105 |
| 3735 | TMEM106A |
| 3736 | TMEM106B |
| 3737 | TMEM107 |
| 3738 | TMEM108 |
| 3739 | TMEM109 |
| 3740 | TMEM11 |
| 3741 | TMEM110 |
| 3742 | TMEM114 |
| 3743 | TMEM115 |
| 3744 | TMEM119 |
| 3745 | TMEM120A |
| 3746 | TMEM120B |
| 3747 | TMEM121 |
| 3748 | TMEM123 |
| 3749 | TMEM125 |
| 3750 | TMEM126B |
| 3751 | TMEM127 |
| 3752 | TMEM128 |
| 3753 | TMEM129 |
| 3754 | TMEM130 |
| 3755 | TMEM131 |
| 3756 | TMEM132A |
| 3757 | TMEM132B |
| 3758 | TMEM132C |
| 3759 | TMEM132D |
| 3760 | TMEM132E |
| 3761 | TMEM133 |
| 3762 | TMEM134 |
| 3763 | TMEM135 |
| 3764 | TMEM136 |
| 3765 | TMEM138 |
| 3766 | TMEM139 |
| 3767 | TMEM143 |
| 3768 | TMEM144 |
| 3769 | TMEM145 |
| 3770 | TMEM147 |
| 3771 | TMEM14A |
| 3772 | TMEM14B |
| 3773 | TMEM14C |
| 3774 | TMEM14E |
| 3775 | TMEM150A |
| 3776 | TMEM151A |
| 3777 | TMEM151B |
| 3778 | TMEM154 |
| 3779 | TMEM156 |
| 3780 | TMEM159 |
| 3781 | TMEM161A |
| 3782 | TMEM163 |
| 3783 | TMEM167B |
| 3784 | TMEM169 |
| 3785 | TMEM170A |
| 3786 | TMEM170B |
| 3787 | TMEM171 |
| 3788 | TMEM173 |
| 3789 | TMEM174 |
| 3790 | TMEM176B |
| 3791 | TMEM177 |
| 3792 | TMEM178A |
| 3793 | TMEM179 |
| 3794 | TMEM18 |
| 3795 | TMEM180 |
| 3796 | TMEM181 |
| 3797 | TMEM182 |
| 3798 | TMEM183A |
| 3799 | TMEM183B |
| 3800 | TMEM184A |
| 3801 | TMEM184B |
| 3802 | TMEM184C |
| 3803 | TMEM185B |
| 3804 | TMEM186 |
| 3805 | TMEM189 |
| 3806 | TMEM190 |
| 3807 | TMEM191A |
| 3808 | TMEM191B |
| 3809 | TMEM192 |
| 3810 | TMEM194A |
| 3811 | TMEM194B |
| 3812 | TMEM196 |
| 3813 | TMEM199 |
| 3814 | TMEM2 |
| 3815 | TMEM200A |
| 3816 | TMEM200C |

| | |
|---|---|
| 3817 | TMEM202 |
| 3818 | TMEM203 |
| 3819 | TMEM204 |
| 3820 | TMEM205 |
| 3821 | TMEM206 |
| 3822 | TMEM207 |
| 3823 | TMEM208 |
| 3824 | TMEM209 |
| 3825 | TMEM213 |
| 3826 | TMEM214 |
| 3827 | TMEM215 |
| 3828 | TMEM216 |
| 3829 | TMEM218 |
| 3830 | TMEM219 |
| 3831 | TMEM220 |
| 3832 | TMEM221 |
| 3833 | TMEM222 |
| 3834 | TMEM223 |
| 3835 | TMEM225 |
| 3836 | TMEM229B |
| 3837 | TMEM231 |
| 3838 | TMEM233 |
| 3839 | TMEM234 |
| 3840 | TMEM235 |
| 3841 | TMEM236 |
| 3842 | TMEM238 |
| 3843 | TMEM239 |
| 3844 | TMEM241 |
| 3845 | TMEM242 |
| 3846 | TMEM245 |
| 3847 | TMEM247 |
| 3848 | TMEM252 |
| 3849 | TMEM253 |
| 3850 | TMEM254 |
| 3851 | TMEM255B |
| 3852 | TMEM256 |
| 3853 | TMEM257 |
| 3854 | TMEM258 |
| 3855 | TMEM26 |
| 3856 | TMEM27 |
| 3857 | TMEM30A |
| 3858 | TMEM30B |
| 3859 | TMEM30C |
| 3860 | TMEM31 |
| 3861 | TMEM33 |
| 3862 | TMEM35 |
| 3863 | TMEM37 |
| 3864 | TMEM38A |
| 3865 | TMEM40 |
| 3866 | TMEM42 |
| 3867 | TMEM43 |
| 3868 | TMEM45A |
| 3869 | TMEM45B |
| 3870 | TMEM48 |
| 3871 | TMEM5 |
| 3872 | TMEM50A |
| 3873 | TMEM51 |
| 3874 | TMEM52 |
| 3875 | TMEM52B |
| 3876 | TMEM53 |
| 3877 | TMEM54 |
| 3878 | TMEM55A |
| 3879 | TMEM55B |
| 3880 | TMEM56 |
| 3881 | TMEM59 |
| 3882 | TMEM59L |
| 3883 | TMEM60 |
| 3884 | TMEM61 |
| 3885 | TMEM62 |
| 3886 | TMEM63A |
| 3887 | TMEM63B |
| 3888 | TMEM63C |
| 3889 | TMEM64 |
| 3890 | TMEM65 |
| 3891 | TMEM66 |
| 3892 | TMEM67 |
| 3893 | TMEM70 |
| 3894 | TMEM71 |
| 3895 | TMEM72 |
| 3896 | TMEM74 |
| 3897 | TMEM74B |
| 3898 | TMEM80 |
| 3899 | TMEM81 |

| | |
|---|---|
| 3900 | TMEM82 |
| 3901 | TMEM86A |
| 3902 | TMEM87A |
| 3903 | TMEM87B |
| 3904 | TMEM88 |
| 3905 | TMEM89 |
| 3906 | TMEM8A |
| 3907 | TMEM8B |
| 3908 | TMEM8C |
| 3909 | TMEM9 |
| 3910 | TMEM91 |
| 3911 | TMEM92 |
| 3912 | TMEM95 |
| 3913 | TMEM98 |
| 3914 | TMEM9B |
| 3915 | TMIE |
| 3916 | TMIGD1 |
| 3917 | TMPO |
| 3918 | TMPPE |
| 3919 | TMPRSS11A |
| 3920 | TMPRSS11B |
| 3921 | TMPRSS11D |
| 3922 | TMPRSS11E |
| 3923 | TMPRSS11F |
| 3924 | TMPRSS13 |
| 3925 | TMPRSS15 |
| 3926 | TMPRSS2 |
| 3927 | TMPRSS3 |
| 3928 | TMPRSS4 |
| 3929 | TMPRSS5 |
| 3930 | TMPRSS6 |
| 3931 | TMPRSS7 |
| 3932 | TMPRSS9 |
| 3933 | TMTC1 |
| 3934 | TMTC2 |
| 3935 | TMTC4 |
| 3936 | TMUB1 |
| 3937 | TMX1 |
| 3938 | TMX2 |
| 3939 | TMX3 |
| 3940 | TMX4 |
| 3941 | TNF |
| 3942 | TNFRSF10A |
| 3943 | TNFRSF10B |
| 3944 | TNFRSF10D |
| 3945 | TNFRSF11A |
| 3946 | TNFRSF12A |
| 3947 | TNFRSF13B |
| 3948 | TNFRSF13C |
| 3949 | TNFRSF14 |
| 3950 | TNFRSF17 |
| 3951 | TNFRSF18 |
| 3952 | TNFRSF19 |
| 3953 | TNFRSF1A |
| 3954 | TNFRSF1B |
| 3955 | TNFRSF21 |
| 3956 | TNFRSF4 |
| 3957 | TNFRSF8 |
| 3958 | TNFRSF9 |
| 3959 | TNFSF10 |
| 3960 | TNFSF11 |
| 3961 | TNFSF12 |
| 3962 | TNFSF13 |
| 3963 | TNFSF13B |
| 3964 | TNFSF14 |
| 3965 | TNFSF18 |
| 3966 | TNFSF4 |
| 3967 | TNFSF8 |
| 3968 | TNFSF9 |
| 3969 | TNMD |
| 3970 | TNN |
| 3971 | TNR |
| 3972 | TNS1 |
| 3973 | TOMM20 |
| 3974 | TOMM20L |
| 3975 | TOMM22 |
| 3976 | TOMM5 |
| 3977 | TOMM7 |
| 3978 | TOMM70A |
| 3979 | TOR1AIP1 |
| 3980 | TOR1AIP2 |
| 3981 | TOR4A |
| 3982 | TP53I11 |

| | |
|---|---|
| 3983 | TPBG |
| 3984 | TPBGL |
| 3985 | TPCN2 |
| 3986 | TPO |
| 3987 | TPRA1 |
| 3988 | TPSG1 |
| 3989 | TPST1 |
| 3990 | TPST2 |
| 3991 | TRABD2B |
| 3992 | TRAF3IP3 |
| 3993 | TRAM1 |
| 3994 | TRAM1L1 |
| 3995 | TRAM2 |
| 3996 | TRAT1 |
| 3997 | TRDN |
| 3998 | TREM1 |
| 3999 | TREM2 |
| 4000 | TREML1 |
| 4001 | TREML2 |
| 4002 | TRHDE |
| 4003 | TRIM13 |
| 4004 | TRIM59 |
| 4005 | TRPA1 |
| 4006 | TRPC1 |
| 4007 | TRPC3 |
| 4008 | TRPC4 |
| 4009 | TRPC7 |
| 4010 | TRPM1 |
| 4011 | TRPM2 |
| 4012 | TRPM3 |
| 4013 | TRPM4 |
| 4014 | TRPM5 |
| 4015 | TRPM8 |
| 4016 | TRPV1 |
| 4017 | TRPV2 |
| 4018 | TRPV3 |
| 4019 | TRPV4 |
| 4020 | TRPV5 |
| 4021 | TRPV6 |
| 4022 | TSHR |
| 4023 | TSNARE1 |
| 4024 | TSPAN1 |
| 4025 | TSPAN10 |
| 4026 | TSPAN11 |
| 4027 | TSPAN12 |
| 4028 | TSPAN13 |
| 4029 | TSPAN14 |
| 4030 | TSPAN15 |
| 4031 | TSPAN16 |
| 4032 | TSPAN17 |
| 4033 | TSPAN18 |
| 4034 | TSPAN19 |
| 4035 | TSPAN2 |
| 4036 | TSPAN3 |
| 4037 | TSPAN31 |
| 4038 | TSPAN32 |
| 4039 | TSPAN33 |
| 4040 | TSPAN4 |
| 4041 | TSPAN5 |
| 4042 | TSPAN6 |
| 4043 | TSPAN7 |
| 4044 | TSPAN8 |
| 4045 | TSPAN9 |
| 4046 | TSPEAR |
| 4047 | TTYH1 |
| 4048 | TTYH2 |
| 4049 | TTYH3 |
| 4050 | TUSC3 |
| 4051 | TUSC5 |
| 4052 | TXNDC11 |
| 4053 | TXNDC15 |
| 4054 | TYR |
| 4055 | TYRO3 |
| 4056 | TYROBP |
| 4057 | TYRP1 |
| 4058 | UBAC2 |
| 4059 | UBE2J1 |
| 4060 | UBE2J2 |
| 4061 | UBIAD1 |
| 4062 | UBR3 |
| 4063 | UBR4 |
| 4064 | UBXN8 |
| 4065 | UCP1 |

| | |
|---|---|
| 4066 | UCP2 |
| 4067 | UCP3 |
| 4068 | UGT1A6 |
| 4069 | UGT1A9 |
| 4070 | UGT2A1 |
| 4071 | UGT2A2 |
| 4072 | UGT2A3 |
| 4073 | UGT2B10 |
| 4074 | UGT2B11 |
| 4075 | UGT2B15 |
| 4076 | UGT2B28 |
| 4077 | UGT2B7 |
| 4078 | UGT3A1 |
| 4079 | UGT8 |
| 4080 | ULBP2 |
| 4081 | UMODL1 |
| 4082 | UNC50 |
| 4083 | UNC5A |
| 4084 | UNC5B |
| 4085 | UNC5C |
| 4086 | UNC5CL |
| 4087 | UNC80 |
| 4088 | UNC93A |
| 4089 | UNC93B1 |
| 4090 | UPK1A |
| 4091 | UPK1B |
| 4092 | UPK2 |
| 4093 | UPK3A |
| 4094 | UPK3B |
| 4095 | UPK3BL |
| 4096 | UQCRFS1 |
| 4097 | USE1 |
| 4098 | USH2A |
| 4099 | USMG5 |
| 4100 | USP14 |
| 4101 | USP19 |
| 4102 | USP30 |
| 4103 | UST |
| 4104 | UXS1 |
| 4105 | VAMP1 |
| 4106 | VAMP2 |
| 4107 | VAMP4 |
| 4108 | VAMP5 |
| 4109 | VAMP7 |
| 4110 | VAMP8 |
| 4111 | VANGL2 |
| 4112 | VAPA |
| 4113 | VAPB |
| 4114 | VASN |
| 4115 | VAT1 |
| 4116 | VCAM1 |
| 4117 | VDAC1 |
| 4118 | VDAC2 |
| 4119 | VDAC3 |
| 4120 | VEGFA |
| 4121 | VIMP |
| 4122 | VIPR1 |
| 4123 | VLDLR |
| 4124 | VMA21 |
| 4125 | VN1R2 |
| 4126 | VN1R5 |
| 4127 | VOPP1 |
| 4128 | VRK2 |
| 4129 | VSIG1 |
| 4130 | VSIG10 |
| 4131 | VSIG10L |
| 4132 | VSIG2 |
| 4133 | VSIG4 |
| 4134 | VSIG8 |
| 4135 | VSTM1 |
| 4136 | VSTM2B |
| 4137 | VSTM4 |
| 4138 | VSTM5 |
| 4139 | VTCN1 |
| 4140 | VTI1A |
| 4141 | VTI1B |
| 4142 | VWF |
| 4143 | WBP1L |
| 4144 | WBSCR17 |
| 4145 | WDFY4 |
| 4146 | WDR11 |
| 4147 | WNT1 |
| 4148 | WNT3A |

| | |
|---|---|
| 4149 | WNT4 |
| 4150 | WNT5A |
| 4151 | WNT5B |
| 4152 | WNT6 |
| 4153 | WNT7A |
| 4154 | WSCD1 |
| 4155 | WSCD2 |
| 4156 | XBP1 |
| 4157 | XCR1 |
| 4158 | XG |
| 4159 | XK |
| 4160 | XKR4 |
| 4161 | XKR6 |
| 4162 | XKR8 |
| 4163 | XKRY |
| 4164 | XKRY2 |
| 4165 | XPR1 |
| 4166 | XXYLT1 |
| 4167 | XYLT2 |
| 4168 | YIF1A |
| 4169 | YIPF1 |
| 4170 | YIPF2 |
| 4171 | YIPF3 |
| 4172 | YIPF7 |
| 4173 | YME1L1 |
| 4174 | ZACN |
| 4175 | ZAN |
| 4176 | ZDHHC1 |
| 4177 | ZDHHC11 |
| 4178 | ZDHHC12 |
| 4179 | ZDHHC14 |
| 4180 | ZDHHC15 |
| 4181 | ZDHHC17 |
| 4182 | ZDHHC19 |
| 4183 | ZDHHC21 |
| 4184 | ZDHHC22 |
| 4185 | ZDHHC3 |
| 4186 | ZDHHC4 |
| 4187 | ZDHHC5 |
| 4188 | ZDHHC7 |
| 4189 | ZDHHC8 |
| 4190 | ZFPL1 |
| 4191 | ZFYVE19 |
| 4192 | ZFYVE27 |
| 4193 | ZMPSTE24 |
| 4194 | ZNRF3 |
| 4195 | ZP1 |
| 4196 | ZP2 |
| 4197 | ZP3 |
| 4198 | ZP4 |
| 4199 | ZPLD1 |

#### B. Secretome (3391)

| No. | Gene Name |
|---|---|
| 1 | MARCH2 |
| 2 | SEPT11 |
| 3 | SEP15 |
| 4 | A1BG |
| 5 | A2M |
| 6 | A2ML1 |
| 7 | AADACL2 |
| 8 | AARS |
| 9 | AASDHPPT |
| 10 | ABAT |
| 11 | ABHD14B |
| 12 | ABHD15 |
| 13 | ABHD8 |
| 14 | ABI1 |
| 15 | ABI3BP |
| 16 | ABP1 |
| 17 | ACAA2 |
| 18 | ACACA |
| 19 | ACADM |
| 20 | ACADSB |
| 21 | ACAN |
| 22 | ACAT1 |
| 23 | ACAT2 |
| 24 | ACLY |
| 25 | ACMSD |
| 26 | ACO1 |
| 27 | ACOT11 |
| 28 | ACOT13 |
| 29 | ACOT2 |
| 30 | ACOT7 |
| 31 | ACP1 |
| 32 | ACP5 |
| 33 | ACPP |
| 34 | ACR |
| 35 | ACRBP |
| 36 | ACRV1 |
| 37 | ACSM1 |
| 38 | ACTA1 |
| 39 | ACTA2 |
| 40 | ACTB |
| 41 | ACTBL2 |

| | |
|---|---|
| 42 | ACTC1 |
| 43 | ACTG1 |
| 44 | ACTG2 |
| 45 | ACTN1 |
| 46 | ACTN2 |
| 47 | ACTN3 |
| 48 | ACTN4 |
| 49 | ACTR1A |
| 50 | ACTR1B |
| 51 | ACTR2 |
| 52 | ACTR3 |
| 53 | ACTR3B |
| 54 | ACTR3C |
| 55 | ACY1 |
| 56 | ACY3 |
| 57 | ACYP1 |
| 58 | ADAMDEC1 |
| 59 | ADAMTS1 |
| 60 | ADAMTS10 |
| 61 | ADAMTS12 |
| 62 | ADAMTS14 |
| 63 | ADAMTS15 |
| 64 | ADAMTS16 |
| 65 | ADAMTS17 |
| 66 | ADAMTS18 |
| 67 | ADAMTS19 |
| 68 | ADAMTS2 |
| 69 | ADAMTS20 |
| 70 | ADAMTS3 |
| 71 | ADAMTS4 |
| 72 | ADAMTS5 |
| 73 | ADAMTS6 |
| 74 | ADAMTS8 |
| 75 | ADAMTS9 |
| 76 | ADAMTSL1 |
| 77 | ADAMTSL2 |
| 78 | ADAMTSL3 |
| 79 | ADAMTSL4 |
| 80 | ADAMTSL5 |
| 81 | ADCK1 |
| 82 | ADCYAP1 |
| 83 | ADH5 |
| 84 | ADH6 |

| | |
|---|---|
| 85 | ADH7 |
| 86 | ADIRF |
| 87 | ADM |
| 88 | ADM2 |
| 89 | ADM5 |
| 90 | ADNP |
| 91 | ADPGK |
| 92 | ADSS |
| 93 | AEBP1 |
| 94 | AFM |
| 95 | AFP |
| 96 | AGA |
| 97 | AGAP2 |
| 98 | AGGF1 |
| 99 | AGMAT |
| 100 | AGR2 |
| 101 | AGR3 |
| 102 | AGRP |
| 103 | AGT |
| 104 | AHCTF1 |
| 105 | AHCY |
| 106 | AHCYL1 |
| 107 | AHNAK |
| 108 | AHSA1 |
| 109 | AHSG |
| 110 | AIF1L |
| 111 | AK1 |
| 112 | AK2 |
| 113 | AK4 |
| 114 | AKR1A1 |
| 115 | AKR1B1 |
| 116 | AKR1B10 |
| 117 | AKR1C1 |
| 118 | AKR1C3 |
| 119 | AKR1C4 |
| 120 | AKR1D1 |
| 121 | AKR1E2 |
| 122 | AKR7A2 |
| 123 | AKR7A3 |
| 124 | AKR7L |
| 125 | ALAD |
| 126 | ALDH16A1 |
| 127 | ALDH1A1 |

| | |
|---|---|
| 128 | ALDH1A3 |
| 129 | ALDH1L1 |
| 130 | ALDH1L2 |
| 131 | ALDH2 |
| 132 | ALDH3A1 |
| 133 | ALDH3B1 |
| 134 | ALDH6A1 |
| 135 | ALDH7A1 |
| 136 | ALDH8A1 |
| 137 | ALDH9A1 |
| 138 | ALDOA |
| 139 | ALDOB |
| 140 | ALDOC |
| 141 | ALG2 |
| 142 | ALOX12 |
| 143 | ALOX15B |
| 144 | ALOX5 |
| 145 | ALPI |
| 146 | ALPL |
| 147 | ALPPL2 |
| 148 | ALYREF |
| 149 | AMBN |
| 150 | AMELY |
| 151 | AMH |
| 152 | AMTN |
| 153 | AMY1A |
| 154 | AMY2A |
| 155 | AMY2B |
| 156 | ANG |
| 157 | ANGPT1 |
| 158 | ANGPT2 |
| 159 | ANGPT4 |
| 160 | ANGPTL1 |
| 161 | ANGPTL2 |
| 162 | ANGPTL4 |
| 163 | ANGPTL5 |
| 164 | ANGPTL6 |
| 165 | ANGPTL7 |
| 166 | ANKFY1 |
| 167 | ANKRD19P |
| 168 | ANP32B |
| 169 | ANXA11 |
| 170 | ANXA13 |

| | |
|---|---|
| 171 | ANXA2P2 |
| 172 | ANXA3 |
| 173 | ANXA6 |
| 174 | ANXA7 |
| 175 | ANXA8L2 |
| 176 | AOAH |
| 177 | AOC2 |
| 178 | AOX1 |
| 179 | AP1M1 |
| 180 | AP1S1 |
| 181 | AP2M1 |
| 182 | AP4M1 |
| 183 | APAF1 |
| 184 | APCS |
| 185 | APEH |
| 186 | APLN |
| 187 | APOA1BP |
| 188 | APOA2 |
| 189 | APOA4 |
| 190 | APOA5 |
| 191 | APOB |
| 192 | APOBR |
| 193 | APOC1 |
| 194 | APOC2 |
| 195 | APOC3 |
| 196 | APOC4 |
| 197 | APOD |
| 198 | APOF |
| 199 | APOL1 |
| 200 | APOL2 |
| 201 | APOL3 |
| 202 | APOL4 |
| 203 | APOL5 |
| 204 | APOL6 |
| 205 | APOM |
| 206 | APPL1 |
| 207 | APPL2 |
| 208 | APRT |
| 209 | ARF1 |
| 210 | ARF3 |
| 211 | ARF4 |
| 212 | ARF5 |
| 213 | ARG1 |

| | |
|---|---|
| 214 | ARHGAP1 |
| 215 | ARHGAP23 |
| 216 | ARHGAP36 |
| 217 | ARHGDIA |
| 218 | ARHGDIB |
| 219 | ARHGEF12 |
| 220 | ARHGEF18 |
| 221 | ARL1 |
| 222 | ARL15 |
| 223 | ARL2 |
| 224 | ARL3 |
| 225 | ARL6 |
| 226 | ARL8A |
| 227 | ARL8B |
| 228 | ARMC3 |
| 229 | ARMC9 |
| 230 | ARPC1A |
| 231 | ARPC1B |
| 232 | ARPC2 |
| 233 | ARPC3 |
| 234 | ARPC4 |
| 235 | ARPC5 |
| 236 | ARPC5L |
| 237 | ARRDC1 |
| 238 | ARSA |
| 239 | ARSD |
| 240 | ARSE |
| 241 | ARSF |
| 242 | ARSG |
| 243 | ARSI |
| 244 | ARSJ |
| 245 | ARSK |
| 246 | ART3 |
| 247 | ART5 |
| 248 | ARTN |
| 249 | ASAH1 |
| 250 | ASH1L |
| 251 | ASIP |
| 252 | ASL |
| 253 | ASNA1 |
| 254 | ASPA |
| 255 | ASPN |
| 256 | ASS1 |

| | |
|---|---|
| 257 | ASTL |
| 258 | ASXL1 |
| 259 | ATAD2 |
| 260 | ATG4C |
| 261 | ATIC |
| 262 | ATP5A1 |
| 263 | ATP5C1 |
| 264 | ATP5F1 |
| 265 | ATP5H |
| 266 | ATP5L |
| 267 | ATP5O |
| 268 | ATP6V0D1 |
| 269 | ATP6V0D2 |
| 270 | ATP6V1A |
| 271 | ATP6V1B1 |
| 272 | ATP6V1B2 |
| 273 | ATP6V1C1 |
| 274 | ATP6V1C2 |
| 275 | ATP6V1D |
| 276 | ATP6V1E1 |
| 277 | ATP6V1F |
| 278 | ATP6V1G1 |
| 279 | ATP6V1H |
| 280 | ATXN10 |
| 281 | AVP |
| 282 | AZGP1 |
| 283 | AZI1 |
| 284 | AZU1 |
| 285 | BAGE |
| 286 | BAGE2 |
| 287 | BAGE3 |
| 288 | BAGE4 |
| 289 | BAGE5 |
| 290 | BAIAP2 |
| 291 | BAIAP2L1 |
| 292 | BANF1 |
| 293 | BASP1 |
| 294 | BBOX1 |
| 295 | BCAN |
| 296 | BCAS1 |
| 297 | BCHE |
| 298 | BCL2L2 |
| 299 | BCL3 |

| | |
|---|---|
| 300 | BCR |
| 301 | BDH2 |
| 302 | BDNF |
| 303 | BEND7 |
| 304 | BEX5 |
| 305 | BGLAP |
| 306 | BHLHB9 |
| 307 | BHMT |
| 308 | BHMT2 |
| 309 | BID |
| 310 | BIVM |
| 311 | BLMH |
| 312 | BLOC1S1 |
| 313 | BLVRA |
| 314 | BLVRB |
| 315 | BMP1 |
| 316 | BMP15 |
| 317 | BMP3 |
| 318 | BMP4 |
| 319 | BMP5 |
| 320 | BMP6 |
| 321 | BMP7 |
| 322 | BMP8A |
| 323 | BMP8B |
| 324 | BMPER |
| 325 | BOLA1 |
| 326 | BOLA2 |
| 327 | BOLA3 |
| 328 | BPGM |
| 329 | BPHL |
| 330 | BPI |
| 331 | BPIFA1 |
| 332 | BPIFA2 |
| 333 | BPIFA3 |
| 334 | BPIFA4P |
| 335 | BPIFB1 |
| 336 | BPIFB2 |
| 337 | BPIFB3 |
| 338 | BPIFB4 |
| 339 | BPIFB6 |
| 340 | BPIFC |
| 341 | BPNT1 |
| 342 | BPTF |

| | |
|---|---|
| 343 | BRK1 |
| 344 | BROX |
| 345 | BRPF3 |
| 346 | BST1 |
| 347 | BTBD17 |
| 348 | BTD |
| 349 | BTG2 |
| 350 | C10orf25 |
| 351 | C10orf99 |
| 352 | C11orf44 |
| 353 | C11orf45 |
| 354 | C11orf52 |
| 355 | C11orf54 |
| 356 | C11orf73 |
| 357 | C11orf80 |
| 358 | C11orf94 |
| 359 | C12orf10 |
| 360 | C12orf39 |
| 361 | C12orf49 |
| 362 | C12orf66 |
| 363 | C12orf73 |
| 364 | C14orf93 |
| 365 | C15orf61 |
| 366 | C16orf80 |
| 367 | C16orf89 |
| 368 | C17orf67 |
| 369 | C17orf77 |
| 370 | C17orf99 |
| 371 | C18orf54 |
| 372 | C19orf10 |
| 373 | C19orf80 |
| 374 | C1orf116 |
| 375 | C1orf123 |
| 376 | C1orf54 |
| 377 | C1orf56 |
| 378 | C1orf68 |
| 379 | C1QA |
| 380 | C1QB |
| 381 | C1QC |
| 382 | C1QL1 |
| 383 | C1QL2 |
| 384 | C1QL3 |
| 385 | C1QL4 |

| | |
|---|---|
| 386 | C1QTNF1 |
| 387 | C1QTNF2 |
| 388 | C1QTNF3 |
| 389 | C1QTNF4 |
| 390 | C1QTNF5 |
| 391 | C1QTNF6 |
| 392 | C1QTNF7 |
| 393 | C1QTNF8 |
| 394 | C1QTNF9 |
| 395 | C1QTNF9B |
| 396 | C1R |
| 397 | C1RL |
| 398 | C1S |
| 399 | C2 |
| 400 | C20orf195 |
| 401 | C21orf62 |
| 402 | C22orf46 |
| 403 | C2CD2 |
| 404 | C2orf16 |
| 405 | C2orf40 |
| 406 | C2orf66 |
| 407 | C2orf69 |
| 408 | C2orf82 |
| 409 | C3 |
| 410 | C3orf58 |
| 411 | C3P1 |
| 412 | C4A |
| 413 | C4B |
| 414 | C4BPA |
| 415 | C4BPB |
| 416 | C4orf26 |
| 417 | C4orf29 |
| 418 | C4orf40 |
| 419 | C4orf48 |
| 420 | C5 |
| 421 | C5orf38 |
| 422 | C5orf46 |
| 423 | C5orf55 |
| 424 | C5orf64 |
| 425 | C6 |
| 426 | C6orf1 |
| 427 | C6orf120 |
| 428 | C6orf15 |

| | |
|---|---|
| 429 | C6orf58 |
| 430 | C7 |
| 431 | C7orf34 |
| 432 | C7orf69 |
| 433 | C7orf73 |
| 434 | C8A |
| 435 | C8B |
| 436 | C8G |
| 437 | C9orf142 |
| 438 | C9orf169 |
| 439 | C9orf47 |
| 440 | C9orf72 |
| 441 | CA1 |
| 442 | CA11 |
| 443 | CA2 |
| 444 | CA6 |
| 445 | CAB39 |
| 446 | CAB39L |
| 447 | CABP1 |
| 448 | CABP4 |
| 449 | CACTIN |
| 450 | CACYBP |
| 451 | CAD |
| 452 | CALB1 |
| 453 | CALCA |
| 454 | CALCB |
| 455 | CALM1 |
| 456 | CALML3 |
| 457 | CALML5 |
| 458 | CALR3 |
| 459 | CALU |
| 460 | CAMK4 |
| 461 | CAND1 |
| 462 | CAPN1 |
| 463 | CAPN2 |
| 464 | CAPN7 |
| 465 | CAPNS1 |
| 466 | CAPS |
| 467 | CAPZA1 |
| 468 | CAPZA2 |
| 469 | CAPZB |
| 470 | CARD11 |
| 471 | CARHSP1 |

| | |
|---|---|
| 472 | CARTPT |
| 473 | CASC5 |
| 474 | CASK |
| 475 | CASP1 |
| 476 | CASP14 |
| 477 | CASQ1 |
| 478 | CASQ2 |
| 479 | CAT |
| 480 | CBLC |
| 481 | CBLN1 |
| 482 | CBLN2 |
| 483 | CBLN3 |
| 484 | CBLN4 |
| 485 | CBR1 |
| 486 | CBR3 |
| 487 | CC2D1A |
| 488 | CCBE1 |
| 489 | CCDC105 |
| 490 | CCDC126 |
| 491 | CCDC132 |
| 492 | CCDC134 |
| 493 | CCDC147 |
| 494 | CCDC25 |
| 495 | CCDC3 |
| 496 | CCDC30 |
| 497 | CCDC70 |
| 498 | CCDC80 |
| 499 | CCK |
| 500 | CCL1 |
| 501 | CCL11 |
| 502 | CCL13 |
| 503 | CCL14 |
| 504 | CCL15 |
| 505 | CCL16 |
| 506 | CCL17 |
| 507 | CCL18 |
| 508 | CCL19 |
| 509 | CCL2 |
| 510 | CCL20 |
| 511 | CCL21 |
| 512 | CCL22 |
| 513 | CCL23 |
| 514 | CCL24 |

| | |
|---|---|
| 515 | CCL25 |
| 516 | CCL26 |
| 517 | CCL27 |
| 518 | CCL28 |
| 519 | CCL3 |
| 520 | CCL3L1 |
| 521 | CCL4 |
| 522 | CCL4L1 |
| 523 | CCL5 |
| 524 | CCL7 |
| 525 | CCL8 |
| 526 | CCNY |
| 527 | CCT2 |
| 528 | CCT3 |
| 529 | CCT4 |
| 530 | CCT5 |
| 531 | CCT6A |
| 532 | CCT7 |
| 533 | CCT8 |
| 534 | CD160 |
| 535 | CD177 |
| 536 | CD2AP |
| 537 | CD48 |
| 538 | CD52 |
| 539 | CD5L |
| 540 | CDA |
| 541 | CDC37 |
| 542 | CDC42 |
| 543 | CDC42BPA |
| 544 | CDC42BPB |
| 545 | CDC42SE2 |
| 546 | CDC7 |
| 547 | CDCA8 |
| 548 | CDCP2 |
| 549 | CDK1 |
| 550 | CDK13 |
| 551 | CDK5RAP2 |
| 552 | CDKL1 |
| 553 | CDNF |
| 554 | CDSN |
| 555 | CEACAM16 |
| 556 | CEACAM18 |
| 557 | CEACAM6 |

| | |
|---|---|
| 558 | CEACAM7 |
| 559 | CEACAM8 |
| 560 | CECR1 |
| 561 | CECR5 |
| 562 | CEL |
| 563 | CELA1 |
| 564 | CELA2A |
| 565 | CELA2B |
| 566 | CELA3A |
| 567 | CELA3B |
| 568 | CEP164 |
| 569 | CEP250 |
| 570 | CEP68 |
| 571 | CER1 |
| 572 | CERCAM |
| 573 | CES1P1 |
| 574 | CES2 |
| 575 | CES3 |
| 576 | CES4A |
| 577 | CES5A |
| 578 | CETP |
| 579 | CFB |
| 580 | CFC1 |
| 581 | CFC1B |
| 582 | CFD |
| 583 | CFH |
| 584 | CFHR1 |
| 585 | CFHR2 |
| 586 | CFHR3 |
| 587 | CFHR4 |
| 588 | CFHR5 |
| 589 | CFI |
| 590 | CFL1 |
| 591 | CFL2 |
| 592 | CFP |
| 593 | CGA |
| 594 | CGB |
| 595 | CGB1 |
| 596 | CGB2 |
| 597 | CGREF1 |
| 598 | CHAD |
| 599 | CHADL |
| 600 | CHCHD3 |

| | |
|---|---|
| 601 | CHD2 |
| 602 | CHEK1 |
| 603 | CHGA |
| 604 | CHGB |
| 605 | CHI3L1 |
| 606 | CHI3L2 |
| 607 | CHIA |
| 608 | CHID1 |
| 609 | CHIT1 |
| 610 | CHMP1A |
| 611 | CHMP1B |
| 612 | CHMP2A |
| 613 | CHMP2B |
| 614 | CHMP3 |
| 615 | CHMP4A |
| 616 | CHMP4B |
| 617 | CHMP4C |
| 618 | CHMP5 |
| 619 | CHMP6 |
| 620 | CHRD |
| 621 | CHRDL1 |
| 622 | CHRDL2 |
| 623 | CIB1 |
| 624 | CIB2 |
| 625 | CILP |
| 626 | CILP2 |
| 627 | CKB |
| 628 | CKMT1A |
| 629 | CLASP1 |
| 630 | CLCA1 |
| 631 | CLCA3P |
| 632 | CLCF1 |
| 633 | CLEC11A |
| 634 | CLEC18A |
| 635 | CLEC18B |
| 636 | CLEC18C |
| 637 | CLEC19A |
| 638 | CLEC3A |
| 639 | CLEC3B |
| 640 | CLN5 |
| 641 | CLPS |
| 642 | CLPSL1 |
| 643 | CLPSL2 |

| | |
|---|---|
| 644 | CLTC |
| 645 | CLTCL1 |
| 646 | CLU |
| 647 | CLUL1 |
| 648 | CMA1 |
| 649 | CMBL |
| 650 | CMPK1 |
| 651 | CNDP1 |
| 652 | CNDP2 |
| 653 | CNFN |
| 654 | CNKSR2 |
| 655 | CNN2 |
| 656 | CNOT1 |
| 657 | CNP |
| 658 | CNPY2 |
| 659 | CNPY3 |
| 660 | CNPY4 |
| 661 | CNTF |
| 662 | CNTFR |
| 663 | CNTLN |
| 664 | CNTN1 |
| 665 | CNTN3 |
| 666 | CNTN4 |
| 667 | CNTN5 |
| 668 | CNTN6 |
| 669 | COASY |
| 670 | COBLL1 |
| 671 | COCH |
| 672 | COL10A1 |
| 673 | COL11A1 |
| 674 | COL11A2 |
| 675 | COL12A1 |
| 676 | COL14A1 |
| 677 | COL15A1 |
| 678 | COL16A1 |
| 679 | COL18A1 |
| 680 | COL19A1 |
| 681 | COL1A1 |
| 682 | COL1A2 |
| 683 | COL20A1 |
| 684 | COL21A1 |
| 685 | COL22A1 |
| 686 | COL24A1 |

| | |
|---|---|
| 687 | COL26A1 |
| 688 | COL27A1 |
| 689 | COL28A1 |
| 690 | COL2A1 |
| 691 | COL3A1 |
| 692 | COL4A1 |
| 693 | COL4A2 |
| 694 | COL4A3 |
| 695 | COL4A4 |
| 696 | COL4A5 |
| 697 | COL4A6 |
| 698 | COL5A1 |
| 699 | COL5A2 |
| 700 | COL5A3 |
| 701 | COL6A1 |
| 702 | COL6A2 |
| 703 | COL6A3 |
| 704 | COL6A5 |
| 705 | COL6A6 |
| 706 | COL7A1 |
| 707 | COL8A1 |
| 708 | COL8A2 |
| 709 | COL9A1 |
| 710 | COL9A2 |
| 711 | COL9A3 |
| 712 | COLEC10 |
| 713 | COLEC11 |
| 714 | COLQ |
| 715 | COMMD1 |
| 716 | COMMD7 |
| 717 | COMP |
| 718 | COPA |
| 719 | COPS4 |
| 720 | COPS6 |
| 721 | COPS8 |
| 722 | CORO1A |
| 723 | CORO1B |
| 724 | CORT |
| 725 | COTL1 |
| 726 | COX4I1 |
| 727 | COX5A |
| 728 | COX5B |
| 729 | COX7A2 |

| | |
|---|---|
| 730 | CP |
| 731 | CPA1 |
| 732 | CPA2 |
| 733 | CPA3 |
| 734 | CPA4 |
| 735 | CPA5 |
| 736 | CPA6 |
| 737 | CPAMD8 |
| 738 | CPB1 |
| 739 | CPB2 |
| 740 | CPE |
| 741 | CPED1 |
| 742 | CPN1 |
| 743 | CPN2 |
| 744 | CPNE1 |
| 745 | CPNE2 |
| 746 | CPNE3 |
| 747 | CPNE4 |
| 748 | CPNE5 |
| 749 | CPNE6 |
| 750 | CPNE7 |
| 751 | CPNE8 |
| 752 | CPNE9 |
| 753 | CPPED1 |
| 754 | CPQ |
| 755 | CPSF3L |
| 756 | CPVL |
| 757 | CPXM1 |
| 758 | CPXM2 |
| 759 | CPZ |
| 760 | CR1L |
| 761 | CRABP2 |
| 762 | CREB5 |
| 763 | CREG1 |
| 764 | CREG2 |
| 765 | CRELD2 |
| 766 | CRH |
| 767 | CRHBP |
| 768 | CRHR1-IT1 |
| 769 | CRISP1 |
| 770 | CRISP2 |
| 771 | CRISP3 |
| 772 | CRISPLD1 |

| | |
|---|---|
| 773 | CRISPLD2 |
| 774 | CRK |
| 775 | CRKL |
| 776 | CRLF1 |
| 777 | CRNN |
| 778 | CROCC |
| 779 | CRP |
| 780 | CRTAC1 |
| 781 | CRTAP |
| 782 | CRTC2 |
| 783 | CRY2 |
| 784 | CRYAA |
| 785 | CRYL1 |
| 786 | CRYM |
| 787 | CRYZ |
| 788 | CS |
| 789 | CSAG1 |
| 790 | CSE1L |
| 791 | CSF2 |
| 792 | CSF3 |
| 793 | CSH1 |
| 794 | CSH2 |
| 795 | CSHL1 |
| 796 | CSK |
| 797 | CSN1S1 |
| 798 | CSNK2B |
| 799 | CSRP1 |
| 800 | CST1 |
| 801 | CST11 |
| 802 | CST2 |
| 803 | CST3 |
| 804 | CST4 |
| 805 | CST5 |
| 806 | CST6 |
| 807 | CST7 |
| 808 | CST9 |
| 809 | CST9L |
| 810 | CSTA |
| 811 | CSTB |
| 812 | CSTL1 |
| 813 | CTBS |
| 814 | CTDSP1 |
| 815 | CTDSPL |

| | |
|---|---|
| 816 | CTF1 |
| 817 | CTGF |
| 818 | CTHRC1 |
| 819 | CTNNB1 |
| 820 | CTNND1 |
| 821 | CTRB1 |
| 822 | CTRB2 |
| 823 | CTRC |
| 824 | CTRL |
| 825 | CTSA |
| 826 | CTSB |
| 827 | CTSC |
| 828 | CTSD |
| 829 | CTSE |
| 830 | CTSF |
| 831 | CTSH |
| 832 | CTSK |
| 833 | CTSL1 |
| 834 | CTSO |
| 835 | CTSS |
| 836 | CTSW |
| 837 | CTSZ |
| 838 | CTTN |
| 839 | CUL3 |
| 840 | CUL4B |
| 841 | CUTA |
| 842 | CUX2 |
| 843 | CXCL1 |
| 844 | CXCL11 |
| 845 | CXCL13 |
| 846 | CXCL14 |
| 847 | CXCL17 |
| 848 | CXCL2 |
| 849 | CXCL3 |
| 850 | CXCL5 |
| 851 | CXCL6 |
| 852 | CXorf36 |
| 853 | CYB5D2 |
| 854 | CYB5R3 |
| 855 | CYFIP1 |
| 856 | CYFIP2 |
| 857 | CYP2J2 |
| 858 | CYP4A11 |

| | |
|---|---|
| 859 | CYR61 |
| 860 | CYS1 |
| 861 | CYTL1 |
| 862 | DAAM2 |
| 863 | DAB2 |
| 864 | DAB2IP |
| 865 | DAK |
| 866 | DAND5 |
| 867 | DAP |
| 868 | DARS |
| 869 | DBI |
| 870 | DBNL |
| 871 | DCD |
| 872 | DCN |
| 873 | DCTD |
| 874 | DCTN2 |
| 875 | DCXR |
| 876 | DDAH1 |
| 877 | DDAH2 |
| 878 | DDB1 |
| 879 | DDC |
| 880 | DDRGK1 |
| 881 | DDT |
| 882 | DDTL |
| 883 | DDX11 |
| 884 | DDX19B |
| 885 | DDX23 |
| 886 | DDX3X |
| 887 | DDX5 |
| 888 | DEAF1 |
| 889 | DECR1 |
| 890 | DEFA1 |
| 891 | DEFA3 |
| 892 | DEFA4 |
| 893 | DEFA5 |
| 894 | DEFA6 |
| 895 | DEFB1 |
| 896 | DEFB103A |
| 897 | DEFB105A |
| 898 | DEFB107A |
| 899 | DEFB109P1 |
| 900 | DEFB110 |
| 901 | DEFB112 |

| | |
|---|---|
| 902 | DEFB113 |
| 903 | DEFB114 |
| 904 | DEFB130 |
| 905 | DEFB133 |
| 906 | DEFB136 |
| 907 | DEFB4A |
| 908 | DERA |
| 909 | DES |
| 910 | DGCR6 |
| 911 | DHH |
| 912 | DHRS11 |
| 913 | DHRS13 |
| 914 | DHRS2 |
| 915 | DHRS4 |
| 916 | DHRS4L2 |
| 917 | DHRS7 |
| 918 | DHRS7C |
| 919 | DHRS9 |
| 920 | DHRSX |
| 921 | DHX36 |
| 922 | DIP2B |
| 923 | DKK1 |
| 924 | DKK2 |
| 925 | DKK3 |
| 926 | DKK4 |
| 927 | DKKL1 |
| 928 | DLG1 |
| 929 | DLST |
| 930 | DMBT1 |
| 931 | DMC1 |
| 932 | DMKN |
| 933 | DMXL2 |
| 934 | DNAJA1 |
| 935 | DNAJA2 |
| 936 | DNAJB1 |
| 937 | DNAJB11 |
| 938 | DNAJB3 |
| 939 | DNAJB4 |
| 940 | DNAJB9 |
| 941 | DNAJC10 |
| 942 | DNAJC11 |
| 943 | DNAJC13 |
| 944 | DNAJC3 |

| | |
|---|---|
| 945 | DNAJC5 |
| 946 | DNAJC7 |
| 947 | DNASE1 |
| 948 | DNASE1L1 |
| 949 | DNASE1L2 |
| 950 | DNASE1L3 |
| 951 | DNASE2 |
| 952 | DNASE2B |
| 953 | DNHD1 |
| 954 | DNM1 |
| 955 | DNM2 |
| 956 | DNM3 |
| 957 | DNPEP |
| 958 | DNPH1 |
| 959 | DOCK10 |
| 960 | DOCK2 |
| 961 | DOK1 |
| 962 | DOPEY2 |
| 963 | DPEP1 |
| 964 | DPEP2 |
| 965 | DPEP3 |
| 966 | DPP3 |
| 967 | DPP7 |
| 968 | DPT |
| 969 | DPYS |
| 970 | DPYSL2 |
| 971 | DPYSL3 |
| 972 | DRAXIN |
| 973 | DRG1 |
| 974 | DSP |
| 975 | DSPP |
| 976 | DST |
| 977 | DSTN |
| 978 | DUSP23 |
| 979 | DUSP26 |
| 980 | DUSP3 |
| 981 | DUT |
| 982 | DYNC1H1 |
| 983 | DYNC2H1 |
| 984 | DYNLL1 |
| 985 | DYNLRB2 |
| 986 | EBI3 |
| 987 | ECH1 |

| | |
|---|---|
| 988 | ECHDC1 |
| 989 | ECHS1 |
| 990 | ECI1 |
| 991 | ECM1 |
| 992 | ECM2 |
| 993 | EDDM3A |
| 994 | EDDM3B |
| 995 | EDEM2 |
| 996 | EDEM3 |
| 997 | EDF1 |
| 998 | EDIL3 |
| 999 | EDN1 |
| 1000 | EDN2 |
| 1001 | EDN3 |
| 1002 | EEA1 |
| 1003 | EEF1A1 |
| 1004 | EEF1E1 |
| 1005 | EEF1G |
| 1006 | EEF2 |
| 1007 | EFEMP1 |
| 1008 | EFEMP2 |
| 1009 | EFHD1 |
| 1010 | EFNA1 |
| 1011 | EFNA2 |
| 1012 | EFNA3 |
| 1013 | EFNA4 |
| 1014 | EFR3A |
| 1015 | EGFL6 |
| 1016 | EGFL7 |
| 1017 | EGFL8 |
| 1018 | EGFLAM |
| 1019 | EHD1 |
| 1020 | EHD4 |
| 1021 | EIF2A |
| 1022 | EIF2S1 |
| 1023 | EIF2S3 |
| 1024 | EIF3B |
| 1025 | EIF3E |
| 1026 | EIF3H |
| 1027 | EIF3I |
| 1028 | EIF3K |
| 1029 | EIF4A1 |
| 1030 | EIF4E |

| | |
|---|---|
| 1031 | EIF5A |
| 1032 | EIF6 |
| 1033 | ELN |
| 1034 | ELSPBP1 |
| 1035 | EMID1 |
| 1036 | EMILIN1 |
| 1037 | EMILIN2 |
| 1038 | EMILIN3 |
| 1039 | EML5 |
| 1040 | ENAM |
| 1041 | ENDOD1 |
| 1042 | ENDOU |
| 1043 | ENHO |
| 1044 | ENO1 |
| 1045 | ENO2 |
| 1046 | ENO3 |
| 1047 | ENOPH1 |
| 1048 | ENOX1 |
| 1049 | ENPP2 |
| 1050 | ENPP6 |
| 1051 | ENTPD5 |
| 1052 | EOGT |
| 1053 | EPB41L2 |
| 1054 | EPDR1 |
| 1055 | EPHX2 |
| 1056 | EPHX3 |
| 1057 | EPN3 |
| 1058 | EPO |
| 1059 | EPS8 |
| 1060 | EPS8L1 |
| 1061 | EPS8L2 |
| 1062 | EPX |
| 1063 | EPYC |
| 1064 | ERBB2IP |
| 1065 | ERLEC1 |
| 1066 | ERMN |
| 1067 | ERO1L |
| 1068 | ERO1LB |
| 1069 | ERP27 |
| 1070 | ERV3-1 |
| 1071 | ESD |
| 1072 | ESF1 |
| 1073 | ESM1 |

| | |
|---|---|
| 1074 | ESR2 |
| 1075 | ESRRA |
| 1076 | ETFA |
| 1077 | ETFB |
| 1078 | ETS1 |
| 1079 | EVPL |
| 1080 | EXOSC9 |
| 1081 | EYS |
| 1082 | EZR |
| 1083 | F11 |
| 1084 | F12 |
| 1085 | F13A1 |
| 1086 | F13B |
| 1087 | F2 |
| 1088 | F5 |
| 1089 | F7 |
| 1090 | F8 |
| 1091 | F9 |
| 1092 | FABP1 |
| 1093 | FABP3 |
| 1094 | FABP4 |
| 1095 | FABP5 |
| 1096 | FAH |
| 1097 | FAM108A1 |
| 1098 | FAM108B1 |
| 1099 | FAM129A |
| 1100 | FAM129B |
| 1101 | FAM131A |
| 1102 | FAM132A |
| 1103 | FAM150A |
| 1104 | FAM150B |
| 1105 | FAM172A |
| 1106 | FAM178A |
| 1107 | FAM180A |
| 1108 | FAM184A |
| 1109 | FAM198A |
| 1110 | FAM19A1 |
| 1111 | FAM19A2 |
| 1112 | FAM19A3 |
| 1113 | FAM19A4 |
| 1114 | FAM20A |
| 1115 | FAM20C |
| 1116 | FAM212A |

| | |
|---|---|
| 1117 | FAM213A |
| 1118 | FAM213B |
| 1119 | FAM24A |
| 1120 | FAM24B |
| 1121 | FAM3A |
| 1122 | FAM3B |
| 1123 | FAM3C |
| 1124 | FAM3D |
| 1125 | FAM49B |
| 1126 | FAM5B |
| 1127 | FAM5C |
| 1128 | FAM63A |
| 1129 | FAM65A |
| 1130 | FAN1 |
| 1131 | FASN |
| 1132 | FBL |
| 1133 | FBLN1 |
| 1134 | FBLN2 |
| 1135 | FBLN5 |
| 1136 | FBLN7 |
| 1137 | FBN1 |
| 1138 | FBN2 |
| 1139 | FBN3 |
| 1140 | FBP1 |
| 1141 | FBXO2 |
| 1142 | FCGBP |
| 1143 | FCGR3B |
| 1144 | FCN2 |
| 1145 | FCN3 |
| 1146 | FCRL1 |
| 1147 | FCRL2 |
| 1148 | FCRLA |
| 1149 | FCRLB |
| 1150 | FDCSP |
| 1151 | FERMT3 |
| 1152 | FETUB |
| 1153 | FGF1 |
| 1154 | FGF11 |
| 1155 | FGF12 |
| 1156 | FGF13 |
| 1157 | FGF14 |
| 1158 | FGF16 |
| 1159 | FGF17 |

| | |
|---|---|
| 1160 | FGF18 |
| 1161 | FGF19 |
| 1162 | FGF2 |
| 1163 | FGF20 |
| 1164 | FGF21 |
| 1165 | FGF23 |
| 1166 | FGF3 |
| 1167 | FGF4 |
| 1168 | FGF5 |
| 1169 | FGF6 |
| 1170 | FGF7 |
| 1171 | FGF9 |
| 1172 | FGFBP2 |
| 1173 | FGFBP3 |
| 1174 | FGL1 |
| 1175 | FGL2 |
| 1176 | FGR |
| 1177 | FH |
| 1178 | FHIT |
| 1179 | FIBIN |
| 1180 | FIBP |
| 1181 | FIGF |
| 1182 | FIGNL1 |
| 1183 | FJX1 |
| 1184 | FKBP10 |
| 1185 | FKBP14 |
| 1186 | FKBP1A |
| 1187 | FKBP2 |
| 1188 | FKBP4 |
| 1189 | FKBP5 |
| 1190 | FKBP7 |
| 1191 | FKBP9 |
| 1192 | FLG2 |
| 1193 | FLNA |
| 1194 | FLNB |
| 1195 | FLOT2 |
| 1196 | FMNL1 |
| 1197 | FMOD |
| 1198 | FN1 |
| 1199 | FNBP1L |
| 1200 | FNDC1 |
| 1201 | FNDC7 |
| 1202 | FOLH1B |

| | |
|---|---|
| 1203 | FOLR3 |
| 1204 | FOLR4 |
| 1205 | FONG |
| 1206 | FOXRED2 |
| 1207 | FREM1 |
| 1208 | FREM3 |
| 1209 | FRK |
| 1210 | FRMD4B |
| 1211 | FRMD7 |
| 1212 | FRMPD1 |
| 1213 | FRZB |
| 1214 | FSCN1 |
| 1215 | FSHB |
| 1216 | FST |
| 1217 | FSTL1 |
| 1218 | FSTL3 |
| 1219 | FSTL4 |
| 1220 | FSTL5 |
| 1221 | FTCD |
| 1222 | FTH1 |
| 1223 | FTL |
| 1224 | FUCA1 |
| 1225 | FUCA2 |
| 1226 | FUZ |
| 1227 | FXR2 |
| 1228 | G6PD |
| 1229 | GAA |
| 1230 | GALC |
| 1231 | GALE |
| 1232 | GALK1 |
| 1233 | GALM |
| 1234 | GALNS |
| 1235 | GALP |
| 1236 | GAMT |
| 1237 | GANAB |
| 1238 | GAPDH |
| 1239 | GAREML |
| 1240 | GARS |
| 1241 | GART |
| 1242 | GAS1 |
| 1243 | GAS6 |
| 1244 | GAST |
| 1245 | GATM |

| | |
|---|---|
| 1246 | GBA |
| 1247 | GBE1 |
| 1248 | GBP1 |
| 1249 | GBP6 |
| 1250 | GC |
| 1251 | GCA |
| 1252 | GCG |
| 1253 | GDA |
| 1254 | GDF1 |
| 1255 | GDF10 |
| 1256 | GDF11 |
| 1257 | GDF15 |
| 1258 | GDF2 |
| 1259 | GDF3 |
| 1260 | GDF5 |
| 1261 | GDF6 |
| 1262 | GDF7 |
| 1263 | GDF9 |
| 1264 | GDI2 |
| 1265 | GDNF |
| 1266 | GEMIN4 |
| 1267 | GEN1 |
| 1268 | GFER |
| 1269 | GFOD1 |
| 1270 | GFOD2 |
| 1271 | GFPT1 |
| 1272 | GFRA1 |
| 1273 | GFRA2 |
| 1274 | GFRA4 |
| 1275 | GGACT |
| 1276 | GGCT |
| 1277 | GGH |
| 1278 | GH1 |
| 1279 | GH2 |
| 1280 | GHDC |
| 1281 | GHRH |
| 1282 | GHRL |
| 1283 | GIF |
| 1284 | GIP |
| 1285 | GIPC1 |
| 1286 | GIPC2 |
| 1287 | GK |
| 1288 | GK2 |

| | |
|---|---|
| 1289 | GKN1 |
| 1290 | GKN2 |
| 1291 | GLA |
| 1292 | GLB1 |
| 1293 | GLB1L |
| 1294 | GLB1L2 |
| 1295 | GLE1 |
| 1296 | GLIPR1L1 |
| 1297 | GLIPR2 |
| 1298 | GLO1 |
| 1299 | GLOD4 |
| 1300 | GLRX |
| 1301 | GLRX3 |
| 1302 | GLT1D1 |
| 1303 | GLT25D1 |
| 1304 | GLT25D2 |
| 1305 | GLTP |
| 1306 | GLUL |
| 1307 | GLYAT |
| 1308 | GLYCAM1 |
| 1309 | GM2A |
| 1310 | GMDS |
| 1311 | GML |
| 1312 | GMPPA |
| 1313 | GMPPB |
| 1314 | GNA11 |
| 1315 | GNA13 |
| 1316 | GNA14 |
| 1317 | GNAI1 |
| 1318 | GNAI2 |
| 1319 | GNAI3 |
| 1320 | GNAL |
| 1321 | GNAQ |
| 1322 | GNAS |
| 1323 | GNAZ |
| 1324 | GNB1 |
| 1325 | GNB2 |
| 1326 | GNB2L1 |
| 1327 | GNB3 |
| 1328 | GNB4 |
| 1329 | GNG10 |
| 1330 | GNG12 |
| 1331 | GNG2 |

| | |
|---|---|
| 1332 | GNG4 |
| 1333 | GNG5 |
| 1334 | GNG7 |
| 1335 | GNL1 |
| 1336 | GNL3 |
| 1337 | GNLY |
| 1338 | GNPDA1 |
| 1339 | GNPTG |
| 1340 | GNRH1 |
| 1341 | GNRH2 |
| 1342 | GNS |
| 1343 | GOLGA4 |
| 1344 | GOLGA7 |
| 1345 | GOT1 |
| 1346 | GOT2 |
| 1347 | GP2 |
| 1348 | GPC1 |
| 1349 | GPC2 |
| 1350 | GPC3 |
| 1351 | GPC5 |
| 1352 | GPC6 |
| 1353 | GPD1 |
| 1354 | GPD1L |
| 1355 | GPHA2 |
| 1356 | GPHB5 |
| 1357 | GPI |
| 1358 | GPLD1 |
| 1359 | GPT |
| 1360 | GPX1 |
| 1361 | GPX2 |
| 1362 | GPX3 |
| 1363 | GPX4 |
| 1364 | GPX5 |
| 1365 | GPX6 |
| 1366 | GPX7 |
| 1367 | GRB2 |
| 1368 | GREM1 |
| 1369 | GREM2 |
| 1370 | GRHPR |
| 1371 | GRIPAP1 |
| 1372 | GRN |
| 1373 | GSN |
| 1374 | GSS |

| | |
|---|---|
| 1375 | GSTA1 |
| 1376 | GSTA2 |
| 1377 | GSTA3 |
| 1378 | GSTA5 |
| 1379 | GSTCD |
| 1380 | GSTK1 |
| 1381 | GSTM2 |
| 1382 | GSTM3 |
| 1383 | GSTO1 |
| 1384 | GSTO2 |
| 1385 | GSTP1 |
| 1386 | GSTT1 |
| 1387 | GSTT2 |
| 1388 | GSTT2B |
| 1389 | GUCA2A |
| 1390 | GUCA2B |
| 1391 | GUSB |
| 1392 | GYG1 |
| 1393 | GZMA |
| 1394 | GZMB |
| 1395 | GZMH |
| 1396 | GZMK |
| 1397 | GZMM |
| 1398 | H1FOO |
| 1399 | H2AFJ |
| 1400 | H2AFV |
| 1401 | H2AFX |
| 1402 | H2AFY |
| 1403 | H2AFY2 |
| 1404 | H2AFZ |
| 1405 | H3F3A |
| 1406 | H6PD |
| 1407 | HAAO |
| 1408 | HABP2 |
| 1409 | HABP4 |
| 1410 | HADHB |
| 1411 | HAGH |
| 1412 | HAMP |
| 1413 | HAO2 |
| 1414 | HAPLN1 |
| 1415 | HAPLN2 |
| 1416 | HAPLN3 |
| 1417 | HAPLN4 |

| | |
|---|---|
| 1418 | HBA1 |
| 1419 | HBB |
| 1420 | HBD |
| 1421 | HBE1 |
| 1422 | HBG2 |
| 1423 | HBM |
| 1424 | HBS1L |
| 1425 | HBZ |
| 1426 | HCG22 |
| 1427 | HCRT |
| 1428 | HDAC11 |
| 1429 | HDDC2 |
| 1430 | HDGF |
| 1431 | HDHD1 |
| 1432 | HDHD2 |
| 1433 | HDLBP |
| 1434 | HEATR5B |
| 1435 | HEBP1 |
| 1436 | HEBP2 |
| 1437 | HEXA |
| 1438 | HEXB |
| 1439 | HGD |
| 1440 | HGF |
| 1441 | HGFAC |
| 1442 | HGS |
| 1443 | HHIPL1 |
| 1444 | HHIPL2 |
| 1445 | HHLA1 |
| 1446 | HIBCH |
| 1447 | HID1 |
| 1448 | HINT1 |
| 1449 | HINT3 |
| 1450 | HIRA |
| 1451 | HIST1H1B |
| 1452 | HIST1H1E |
| 1453 | HIST1H2AA |
| 1454 | HIST1H2AB |
| 1455 | HIST1H2AC |
| 1456 | HIST1H2AD |
| 1457 | HIST1H2AG |
| 1458 | HIST1H2AH |
| 1459 | HIST1H2AI |
| 1460 | HIST1H2AJ |

| | |
|---|---|
| 1461 | HIST1H2BC |
| 1462 | HIST1H2BD |
| 1463 | HIST1H2BH |
| 1464 | HIST1H2BJ |
| 1465 | HIST1H2BK |
| 1466 | HIST1H2BL |
| 1467 | HIST1H2BM |
| 1468 | HIST1H2BN |
| 1469 | HIST1H3A |
| 1470 | HIST1H4A |
| 1471 | HIST2H2AA3 |
| 1472 | HIST2H2AB |
| 1473 | HIST2H2AC |
| 1474 | HIST2H2BE |
| 1475 | HIST2H2BF |
| 1476 | HIST2H3A |
| 1477 | HIST3H2A |
| 1478 | HIST3H3 |
| 1479 | HMCN1 |
| 1480 | HMGB2 |
| 1481 | HMOX1 |
| 1482 | HMSD |
| 1483 | HNMT |
| 1484 | HNRNPA1 |
| 1485 | HNRNPC |
| 1486 | HNRNPD |
| 1487 | HNRNPK |
| 1488 | HNRNPL |
| 1489 | HNRNPM |
| 1490 | HNRPDL |
| 1491 | HOGA1 |
| 1492 | HP |
| 1493 | HPCAL1 |
| 1494 | HPD |
| 1495 | HPGD |
| 1496 | HPR |
| 1497 | HPRT1 |
| 1498 | HPSE |
| 1499 | HPSE2 |
| 1500 | HPX |
| 1501 | HRC |
| 1502 | HRG |
| 1503 | HRNR |

| | |
|---|---|
| 1504 | HRSP12 |
| 1505 | HS3ST1 |
| 1506 | HSD11B1L |
| 1507 | HSD17B11 |
| 1508 | HSD17B13 |
| 1509 | HSD17B6 |
| 1510 | HSP90AA1 |
| 1511 | HSP90B1 |
| 1512 | HSPA12A |
| 1513 | HSPA13 |
| 1514 | HSPA1A |
| 1515 | HSPA1L |
| 1516 | HSPA4 |
| 1517 | HSPA6 |
| 1518 | HSPA7 |
| 1519 | HSPA8 |
| 1520 | HSPA9 |
| 1521 | HSPB1 |
| 1522 | HSPB11 |
| 1523 | HSPE1 |
| 1524 | HSPG2 |
| 1525 | HSPH1 |
| 1526 | HTN1 |
| 1527 | HTN3 |
| 1528 | HTRA1 |
| 1529 | HTRA3 |
| 1530 | HTRA4 |
| 1531 | HUWE1 |
| 1532 | HYAL1 |
| 1533 | HYAL3 |
| 1534 | HYOU1 |
| 1535 | IAH1 |
| 1536 | IAPP |
| 1537 | IARS |
| 1538 | IBSP |
| 1539 | IDH1 |
| 1540 | IDH2 |
| 1541 | IDS |
| 1542 | IDUA |
| 1543 | IFI27L2 |
| 1544 | IFI30 |
| 1545 | IFNA1 |
| 1546 | IFNA10 |

| | |
|---|---|
| 1547 | IFNA14 |
| 1548 | IFNA16 |
| 1549 | IFNA17 |
| 1550 | IFNA2 |
| 1551 | IFNA21 |
| 1552 | IFNA4 |
| 1553 | IFNA5 |
| 1554 | IFNA6 |
| 1555 | IFNA7 |
| 1556 | IFNA8 |
| 1557 | IFNB1 |
| 1558 | IFNE |
| 1559 | IFNK |
| 1560 | IFNL1 |
| 1561 | IFNL2 |
| 1562 | IFNL3 |
| 1563 | IFNL4 |
| 1564 | IFNW1 |
| 1565 | IFT20 |
| 1566 | IGF1 |
| 1567 | IGF2 |
| 1568 | IGFALS |
| 1569 | IGFBP1 |
| 1570 | IGFBP2 |
| 1571 | IGFBP3 |
| 1572 | IGFBP4 |
| 1573 | IGFBP5 |
| 1574 | IGFBP6 |
| 1575 | IGFBP7 |
| 1576 | IGFBPL1 |
| 1577 | IGFL1 |
| 1578 | IGFL2 |
| 1579 | IGFL3 |
| 1580 | IGFL4 |
| 1581 | IGIP |
| 1582 | IGJ |
| 1583 | IGLL1 |
| 1584 | IGLL5 |
| 1585 | IGSF10 |
| 1586 | IGSF21 |
| 1587 | IHH |
| 1588 | IK |
| 1589 | IL10 |

| | |
|---|---|
| 1590 | IL11 |
| 1591 | IL12A |
| 1592 | IL12B |
| 1593 | IL16 |
| 1594 | IL17B |
| 1595 | IL17C |
| 1596 | IL17D |
| 1597 | IL17F |
| 1598 | IL18 |
| 1599 | IL18BP |
| 1600 | IL19 |
| 1601 | IL1A |
| 1602 | IL1B |
| 1603 | IL1F10 |
| 1604 | IL1RN |
| 1605 | IL2 |
| 1606 | IL20 |
| 1607 | IL21 |
| 1608 | IL22 |
| 1609 | IL22RA2 |
| 1610 | IL23A |
| 1611 | IL24 |
| 1612 | IL25 |
| 1613 | IL26 |
| 1614 | IL27 |
| 1615 | IL3 |
| 1616 | IL31 |
| 1617 | IL32 |
| 1618 | IL33 |
| 1619 | IL34 |
| 1620 | IL36A |
| 1621 | IL36B |
| 1622 | IL36G |
| 1623 | IL36RN |
| 1624 | IL37 |
| 1625 | IL4I1 |
| 1626 | IL5 |
| 1627 | IL7 |
| 1628 | IL8 |
| 1629 | IL9 |
| 1630 | IMPA1 |
| 1631 | IMPDH2 |
| 1632 | IMPG1 |

| | |
|---|---|
| 1633 | INA |
| 1634 | INADL |
| 1635 | INHA |
| 1636 | INHBA |
| 1637 | INHBB |
| 1638 | INHBC |
| 1639 | INHBE |
| 1640 | INPP5A |
| 1641 | INPP5E |
| 1642 | INS |
| 1643 | INS-IGF2 |
| 1644 | INSL3 |
| 1645 | INSL4 |
| 1646 | INSL5 |
| 1647 | INSL6 |
| 1648 | IQCB1 |
| 1649 | IQCG |
| 1650 | IQGAP1 |
| 1651 | IRAK4 |
| 1652 | IRF2BPL |
| 1653 | IRF6 |
| 1654 | ISG15 |
| 1655 | ISLR |
| 1656 | ISM1 |
| 1657 | ISM2 |
| 1658 | ISOC1 |
| 1659 | IST1 |
| 1660 | ITCH |
| 1661 | ITGB1BP1 |
| 1662 | ITGBL1 |
| 1663 | ITIH1 |
| 1664 | ITIH2 |
| 1665 | ITIH3 |
| 1666 | ITIH4 |
| 1667 | ITIH5 |
| 1668 | ITIH6 |
| 1669 | ITLN1 |
| 1670 | ITLN2 |
| 1671 | ITPRIP |
| 1672 | ITSN2 |
| 1673 | IVL |
| 1674 | IZUMO4 |
| 1675 | JMJD8 |

| | |
|---|---|
| 1676 | JUP |
| 1677 | KAL1 |
| 1678 | KALRN |
| 1679 | KARS |
| 1680 | KAT2A |
| 1681 | KAZALD1 |
| 1682 | KCP |
| 1683 | KCTD12 |
| 1684 | KDELC1 |
| 1685 | KDELC2 |
| 1686 | KDM4D |
| 1687 | KERA |
| 1688 | KHK |
| 1689 | KIAA0100 |
| 1690 | KIAA0556 |
| 1691 | KIAA1199 |
| 1692 | KIF12 |
| 1693 | KIF23 |
| 1694 | KIF27 |
| 1695 | KIF3A |
| 1696 | KIF3B |
| 1697 | KIFAP3 |
| 1698 | KIFC3 |
| 1699 | KIR3DX1 |
| 1700 | KISS1 |
| 1701 | KLHL11 |
| 1702 | KLHL17 |
| 1703 | KLHL34 |
| 1704 | KLK1 |
| 1705 | KLK10 |
| 1706 | KLK11 |
| 1707 | KLK12 |
| 1708 | KLK13 |
| 1709 | KLK14 |
| 1710 | KLK15 |
| 1711 | KLK2 |
| 1712 | KLK3 |
| 1713 | KLK4 |
| 1714 | KLK5 |
| 1715 | KLK6 |
| 1716 | KLK7 |
| 1717 | KLK8 |
| 1718 | KLK9 |

| | |
|---|---|
| 1719 | KLKB1 |
| 1720 | KNG1 |
| 1721 | KPNA4 |
| 1722 | KPNB1 |
| 1723 | KPRP |
| 1724 | KRIT1 |
| 1725 | KRR1 |
| 1726 | KRT1 |
| 1727 | KRT10 |
| 1728 | KRT12 |
| 1729 | KRT13 |
| 1730 | KRT14 |
| 1731 | KRT15 |
| 1732 | KRT16 |
| 1733 | KRT17 |
| 1734 | KRT18 |
| 1735 | KRT19 |
| 1736 | KRT2 |
| 1737 | KRT24 |
| 1738 | KRT25 |
| 1739 | KRT26 |
| 1740 | KRT27 |
| 1741 | KRT28 |
| 1742 | KRT3 |
| 1743 | KRT31 |
| 1744 | KRT32 |
| 1745 | KRT33A |
| 1746 | KRT33B |
| 1747 | KRT34 |
| 1748 | KRT35 |
| 1749 | KRT36 |
| 1750 | KRT37 |
| 1751 | KRT38 |
| 1752 | KRT5 |
| 1753 | KRT6A |
| 1754 | KRT6B |
| 1755 | KRT6C |
| 1756 | KRT7 |
| 1757 | KRT71 |
| 1758 | KRT72 |
| 1759 | KRT73 |
| 1760 | KRT74 |
| 1761 | KRT75 |

| | |
|---|---|
| 1762 | KRT76 |
| 1763 | KRT77 |
| 1764 | KRT78 |
| 1765 | KRT79 |
| 1766 | KRT8 |
| 1767 | KRT81 |
| 1768 | KRT83 |
| 1769 | KRT84 |
| 1770 | KRT85 |
| 1771 | KRT86 |
| 1772 | KRT9 |
| 1773 | KRTDAP |
| 1774 | LACRT |
| 1775 | LAD1 |
| 1776 | LAIR2 |
| 1777 | LALBA |
| 1778 | LAMA1 |
| 1779 | LAMA2 |
| 1780 | LAMA3 |
| 1781 | LAMA4 |
| 1782 | LAMA5 |
| 1783 | LAMB1 |
| 1784 | LAMB2 |
| 1785 | LAMB3 |
| 1786 | LAMB4 |
| 1787 | LAMC1 |
| 1788 | LAMC2 |
| 1789 | LAMC3 |
| 1790 | LAMTOR1 |
| 1791 | LAMTOR2 |
| 1792 | LAMTOR3 |
| 1793 | LANCL1 |
| 1794 | LAP3 |
| 1795 | LASP1 |
| 1796 | LCAT |
| 1797 | LCK |
| 1798 | LCN1 |
| 1799 | LCN10 |
| 1800 | LCN12 |
| 1801 | LCN15 |
| 1802 | LCN2 |
| 1803 | LCN6 |
| 1804 | LCN8 |

| | |
|---|---|
| 1805 | LCN9 |
| 1806 | LCP1 |
| 1807 | LDHA |
| 1808 | LDHAL6A |
| 1809 | LDHB |
| 1810 | LDHC |
| 1811 | LEAP2 |
| 1812 | LECT2 |
| 1813 | LEFTY1 |
| 1814 | LEFTY2 |
| 1815 | LEP |
| 1816 | LEPRE1 |
| 1817 | LEPREL1 |
| 1818 | LEPREL2 |
| 1819 | LGALS3BP |
| 1820 | LGALS7 |
| 1821 | LGALS8 |
| 1822 | LGALS9 |
| 1823 | LGI1 |
| 1824 | LGI2 |
| 1825 | LGI3 |
| 1826 | LGI4 |
| 1827 | LGMN |
| 1828 | LHB |
| 1829 | LIF |
| 1830 | LILRA3 |
| 1831 | LIN7A |
| 1832 | LIN7C |
| 1833 | LINC00305 |
| 1834 | LIPA |
| 1835 | LIPC |
| 1836 | LIPF |
| 1837 | LIPG |
| 1838 | LIPH |
| 1839 | LIPI |
| 1840 | LIPK |
| 1841 | LIPM |
| 1842 | LIPN |
| 1843 | LMCD1 |
| 1844 | LOX |
| 1845 | LOXL1 |
| 1846 | LOXL2 |
| 1847 | LOXL3 |

| | |
|---|---|
| 1848 | LOXL4 |
| 1849 | LPA |
| 1850 | LPAL2 |
| 1851 | LPO |
| 1852 | LRCH3 |
| 1853 | LRCOL1 |
| 1854 | LRG1 |
| 1855 | LRRC16A |
| 1856 | LRRC17 |
| 1857 | LRRC57 |
| 1858 | LRRK2 |
| 1859 | LRSAM1 |
| 1860 | LSM6 |
| 1861 | LSP1 |
| 1862 | LTA |
| 1863 | LTA4H |
| 1864 | LTBP1 |
| 1865 | LTBP2 |
| 1866 | LTBP3 |
| 1867 | LTBP4 |
| 1868 | LTF |
| 1869 | LUM |
| 1870 | LUZP1 |
| 1871 | LUZP2 |
| 1872 | LXN |
| 1873 | LY6E |
| 1874 | LY6G5B |
| 1875 | LY6G5C |
| 1876 | LY6G6C |
| 1877 | LY6H |
| 1878 | LY6K |
| 1879 | LY86 |
| 1880 | LY96 |
| 1881 | LYG1 |
| 1882 | LYG2 |
| 1883 | LYN |
| 1884 | LYNX1 |
| 1885 | LYPD1 |
| 1886 | LYPD2 |
| 1887 | LYPD3 |
| 1888 | LYPD4 |
| 1889 | LYPD5 |
| 1890 | LYPD6 |

| | |
|---|---|
| 1891 | LYPD6B |
| 1892 | LYPD8 |
| 1893 | LYPLA1 |
| 1894 | LYPLA2 |
| 1895 | LYPLAL1 |
| 1896 | LYRM1 |
| 1897 | LYZ |
| 1898 | LYZL1 |
| 1899 | LYZL2 |
| 1900 | LYZL4 |
| 1901 | LYZL6 |
| 1902 | MAGEF1 |
| 1903 | MAL |
| 1904 | MAMDC2 |
| 1905 | MAN2B1 |
| 1906 | MAN2B2 |
| 1907 | MANBA |
| 1908 | MANF |
| 1909 | MAP2K1 |
| 1910 | MAP2K2 |
| 1911 | MAP4 |
| 1912 | MAPK1 |
| 1913 | MAPK14 |
| 1914 | MAPK15 |
| 1915 | MAPK3 |
| 1916 | MAPKAPK2 |
| 1917 | MARCKS |
| 1918 | MARCKSL1 |
| 1919 | MARK3 |
| 1920 | MARS |
| 1921 | MASP1 |
| 1922 | MASP2 |
| 1923 | MAT2B |
| 1924 | MATN1 |
| 1925 | MATN2 |
| 1926 | MATN3 |
| 1927 | MATN4 |
| 1928 | MB |
| 1929 | MBD5 |
| 1930 | MBLAC2 |
| 1931 | MBP |
| 1932 | MCF2 |
| 1933 | MCF2L |

| | |
|---|---|
| 1934 | MCFD2 |
| 1935 | MDGA1 |
| 1936 | MDGA2 |
| 1937 | MDH1 |
| 1938 | MDH2 |
| 1939 | MDK |
| 1940 | MDP1 |
| 1941 | MDS2 |
| 1942 | MECP2 |
| 1943 | MEGF6 |
| 1944 | MEPE |
| 1945 | MESDC2 |
| 1946 | MESP2 |
| 1947 | METRN |
| 1948 | METRNL |
| 1949 | METTL24 |
| 1950 | METTL7A |
| 1951 | METTL7B |
| 1952 | METTL9 |
| 1953 | MFAP1 |
| 1954 | MFAP2 |
| 1955 | MFAP4 |
| 1956 | MFAP5 |
| 1957 | MGP |
| 1958 | MGRN1 |
| 1959 | MIA |
| 1960 | MIA2 |
| 1961 | MID2 |
| 1962 | MIEN1 |
| 1963 | MIF |
| 1964 | MINK1 |
| 1965 | MINPP1 |
| 1966 | MIOX |
| 1967 | MITD1 |
| 1968 | MLL5 |
| 1969 | MLLT3 |
| 1970 | MLN |
| 1971 | MLPH |
| 1972 | MMP1 |
| 1973 | MMP10 |
| 1974 | MMP11 |
| 1975 | MMP12 |
| 1976 | MMP13 |

| | |
|---|---|
| 1977 | MMP17 |
| 1978 | MMP19 |
| 1979 | MMP2 |
| 1980 | MMP20 |
| 1981 | MMP25 |
| 1982 | MMP26 |
| 1983 | MMP27 |
| 1984 | MMP28 |
| 1985 | MMP3 |
| 1986 | MMP7 |
| 1987 | MMP8 |
| 1988 | MMP9 |
| 1989 | MMRN1 |
| 1990 | MMRN2 |
| 1991 | MNDA |
| 1992 | MOB1A |
| 1993 | MOB1B |
| 1994 | MON2 |
| 1995 | MOV10 |
| 1996 | MOXD2P |
| 1997 | MPHOSPH8 |
| 1998 | MPI |
| 1999 | MPO |
| 2000 | MPP5 |
| 2001 | MPP6 |
| 2002 | MPST |
| 2003 | MRAS |
| 2004 | MRPL18 |
| 2005 | MSMB |
| 2006 | MSMP |
| 2007 | MSN |
| 2008 | MSRA |
| 2009 | MSRB3 |
| 2010 | MST1L |
| 2011 | MSTN |
| 2012 | MTAP |
| 2013 | MTHFD1 |
| 2014 | MTHFD2 |
| 2015 | MTM1 |
| 2016 | MTMR11 |
| 2017 | MTMR2 |
| 2018 | MTMR4 |
| 2019 | MTPN |

| | |
|---|---|
| 2020 | MTRNR2L1 |
| 2021 | MTRNR2L10 |
| 2022 | MTRNR2L2 |
| 2023 | MTRNR2L3 |
| 2024 | MTRNR2L4 |
| 2025 | MTRNR2L5 |
| 2026 | MTRNR2L6 |
| 2027 | MTRNR2L7 |
| 2028 | MTRNR2L8 |
| 2029 | MTTP |
| 2030 | MTUS1 |
| 2031 | MUC2 |
| 2032 | MUC20 |
| 2033 | MUC5B |
| 2034 | MUC6 |
| 2035 | MUC7 |
| 2036 | MUCL1 |
| 2037 | MUM1L1 |
| 2038 | MVB12A |
| 2039 | MVB12B |
| 2040 | MVK |
| 2041 | MVP |
| 2042 | MXRA5 |
| 2043 | MYH11 |
| 2044 | MYH13 |
| 2045 | MYH14 |
| 2046 | MYH3 |
| 2047 | MYL12A |
| 2048 | MYL12B |
| 2049 | MYL6 |
| 2050 | MYL6B |
| 2051 | MYLK4 |
| 2052 | MYO15A |
| 2053 | MYO1B |
| 2054 | MYO1C |
| 2055 | MYO1D |
| 2056 | MYO1E |
| 2057 | MYO1G |
| 2058 | MYO5A |
| 2059 | MYO5B |
| 2060 | MYO5C |
| 2061 | MYO6 |
| 2062 | MYO7B |

| | |
|---|---|
| 2063 | MYOC |
| 2064 | MYZAP |
| 2065 | MZB1 |
| 2066 | N4BP2L2 |
| 2067 | N6AMT2 |
| 2068 | NAA16 |
| 2069 | NAA50 |
| 2070 | NAAA |
| 2071 | NACA |
| 2072 | NAGA |
| 2073 | NAGK |
| 2074 | NAGLU |
| 2075 | NAIP |
| 2076 | NAMPT |
| 2077 | NANS |
| 2078 | NAPA |
| 2079 | NAPB |
| 2080 | NAPEPLD |
| 2081 | NAPG |
| 2082 | NAPRT1 |
| 2083 | NAPSA |
| 2084 | NARS |
| 2085 | NAV2 |
| 2086 | NBL1 |
| 2087 | NBR1 |
| 2088 | NCALD |
| 2089 | NCAN |
| 2090 | NCCRP1 |
| 2091 | NCL |
| 2092 | NCOA3 |
| 2093 | NCOA5 |
| 2094 | NCS1 |
| 2095 | NDNF |
| 2096 | NDRG1 |
| 2097 | NDRG2 |
| 2098 | NDRG3 |
| 2099 | NDUFA4 |
| 2100 | NDUFAF7 |
| 2101 | NDUFB10 |
| 2102 | NDUFB9 |
| 2103 | NEB |
| 2104 | NEBL |
| 2105 | NEDD4 |

| | |
|---|---|
| 2106 | NEDD4L |
| 2107 | NEDD8 |
| 2108 | NEK2 |
| 2109 | NELL1 |
| 2110 | NELL2 |
| 2111 | NENF |
| 2112 | NEU1 |
| 2113 | NGF |
| 2114 | NGRN |
| 2115 | NHLRC3 |
| 2116 | NID1 |
| 2117 | NIPBL |
| 2118 | NIT1 |
| 2119 | NIT2 |
| 2120 | NKX6-1 |
| 2121 | NLRP4 |
| 2122 | NMB |
| 2123 | NME1 |
| 2124 | NME2 |
| 2125 | NME3 |
| 2126 | NMS |
| 2127 | NMU |
| 2128 | NODAL |
| 2129 | NOG |
| 2130 | NOTCH2NL |
| 2131 | NOTUM |
| 2132 | NPAP1 |
| 2133 | NPB |
| 2134 | NPC2 |
| 2135 | NPEPPS |
| 2136 | NPFF |
| 2137 | NPHS2 |
| 2138 | NPNT |
| 2139 | NPPA |
| 2140 | NPPB |
| 2141 | NPPC |
| 2142 | NPS |
| 2143 | NPTX1 |
| 2144 | NPTX2 |
| 2145 | NPVF |
| 2146 | NPW |
| 2147 | NPY |
| 2148 | NQO1 |

| | |
|---|---|
| 2149 | NQO2 |
| 2150 | NR2C2AP |
| 2151 | NRAS |
| 2152 | NRF1 |
| 2153 | NRN1 |
| 2154 | NRN1L |
| 2155 | NRTN |
| 2156 | NSF |
| 2157 | NT5C |
| 2158 | NTF3 |
| 2159 | NTF4 |
| 2160 | NTN1 |
| 2161 | NTN3 |
| 2162 | NTN4 |
| 2163 | NTN5 |
| 2164 | NTNG1 |
| 2165 | NTNG2 |
| 2166 | NTPCR |
| 2167 | NTS |
| 2168 | NUBP1 |
| 2169 | NUCB1 |
| 2170 | NUCB2 |
| 2171 | NUDCD2 |
| 2172 | NUDT1 |
| 2173 | NUDT10 |
| 2174 | NUDT14 |
| 2175 | NUDT3 |
| 2176 | NUDT5 |
| 2177 | NUDT9 |
| 2178 | NUMA1 |
| 2179 | NUTF2 |
| 2180 | NXPE1 |
| 2181 | NXPE3 |
| 2182 | NXPE4 |
| 2183 | NXPH1 |
| 2184 | NXPH2 |
| 2185 | NXPH3 |
| 2186 | NXPH4 |
| 2187 | NYX |
| 2188 | OAF |
| 2189 | OAS1 |
| 2190 | OAS3 |
| 2191 | OAZ3 |

| | |
|---|---|
| 2192 | OBP2A |
| 2193 | OBP2B |
| 2194 | OC90 |
| 2195 | OCRL |
| 2196 | ODAM |
| 2197 | OGN |
| 2198 | OIT3 |
| 2199 | OLA1 |
| 2200 | OLFM1 |
| 2201 | OLFM2 |
| 2202 | OLFM3 |
| 2203 | OLFM4 |
| 2204 | OLFML1 |
| 2205 | OLFML2A |
| 2206 | OLFML2B |
| 2207 | OLFML3 |
| 2208 | OMD |
| 2209 | OMG |
| 2210 | OPTC |
| 2211 | ORM1 |
| 2212 | ORM2 |
| 2213 | OS9 |
| 2214 | OSBPL1A |
| 2215 | OSCAR |
| 2216 | OSM |
| 2217 | OSTF1 |
| 2218 | OSTN |
| 2219 | OTOA |
| 2220 | OTOGL |
| 2221 | OTOL1 |
| 2222 | OTOR |
| 2223 | OTOS |
| 2224 | OTUB1 |
| 2225 | OVCH1 |
| 2226 | OVCH2 |
| 2227 | OVGP1 |
| 2228 | OXNAD1 |
| 2229 | OXSR1 |
| 2230 | OXT |
| 2231 | P4HA1 |
| 2232 | P4HA2 |
| 2233 | P4HA3 |
| 2234 | P4HB |

| | |
|---|---|
| 2235 | PA2G4 |
| 2236 | PABPC1 |
| 2237 | PABPC1L |
| 2238 | PABPC1L2A |
| 2239 | PABPC3 |
| 2240 | PACSIN2 |
| 2241 | PACSIN3 |
| 2242 | PADI1 |
| 2243 | PADI2 |
| 2244 | PAEP |
| 2245 | PAFAH1B1 |
| 2246 | PAFAH1B2 |
| 2247 | PAFAH1B3 |
| 2248 | PAH |
| 2249 | PAICS |
| 2250 | PAMR1 |
| 2251 | PAPL |
| 2252 | PAPLN |
| 2253 | PAPPA |
| 2254 | PAPPA2 |
| 2255 | PARD6B |
| 2256 | PARK7 |
| 2257 | PARP4 |
| 2258 | PATE1 |
| 2259 | PATE2 |
| 2260 | PATE3 |
| 2261 | PATE4 |
| 2262 | PBLD |
| 2263 | PCBD1 |
| 2264 | PCBP1 |
| 2265 | PCBP2 |
| 2266 | PCBP3 |
| 2267 | PCIF1 |
| 2268 | PCK1 |
| 2269 | PCK2 |
| 2270 | PCLO |
| 2271 | PCMT1 |
| 2272 | PCNA |
| 2273 | PCNT |
| 2274 | PCOLCE |
| 2275 | PCOLCE2 |
| 2276 | PCSK1 |
| 2277 | PCSK1N |

| | |
|---|---|
| 2278 | PCSK2 |
| 2279 | PCYOX1 |
| 2280 | PCYOX1L |
| 2281 | PDCD10 |
| 2282 | PDCD2 |
| 2283 | PDCD5 |
| 2284 | PDCD6 |
| 2285 | PDCD6IP |
| 2286 | PDDC1 |
| 2287 | PDE4C |
| 2288 | PDE8A |
| 2289 | PDF |
| 2290 | PDGFD |
| 2291 | PDGFRL |
| 2292 | PDHB |
| 2293 | PDIA2 |
| 2294 | PDIA5 |
| 2295 | PDIA6 |
| 2296 | PDILT |
| 2297 | PDLIM2 |
| 2298 | PDXK |
| 2299 | PDXP |
| 2300 | PDYN |
| 2301 | PDZD11 |
| 2302 | PDZD2 |
| 2303 | PDZD7 |
| 2304 | PDZK1 |
| 2305 | PEBP1 |
| 2306 | PEBP4 |
| 2307 | PEF1 |
| 2308 | PENK |
| 2309 | PEPD |
| 2310 | PEX1 |
| 2311 | PF4 |
| 2312 | PF4V1 |
| 2313 | PFAS |
| 2314 | PFDN2 |
| 2315 | PFKL |
| 2316 | PFKM |
| 2317 | PFKP |
| 2318 | PFN1 |
| 2319 | PFN2 |
| 2320 | PGA3 |

| | |
|---|---|
| 2321 | PGA4 |
| 2322 | PGA5 |
| 2323 | PGAM1 |
| 2324 | PGAM2 |
| 2325 | PGAM4 |
| 2326 | PGC |
| 2327 | PGD |
| 2328 | PGF |
| 2329 | PGK1 |
| 2330 | PGK2 |
| 2331 | PGLS |
| 2332 | PGLYRP1 |
| 2333 | PGLYRP2 |
| 2334 | PGLYRP3 |
| 2335 | PGLYRP4 |
| 2336 | PGM1 |
| 2337 | PGM2 |
| 2338 | PHB |
| 2339 | PHB2 |
| 2340 | PHF15 |
| 2341 | PHF17 |
| 2342 | PHGDH |
| 2343 | PHIP |
| 2344 | PHLDA3 |
| 2345 | PHPT1 |
| 2346 | PI15 |
| 2347 | PI3 |
| 2348 | PI4KA |
| 2349 | PIK3C2A |
| 2350 | PIN4 |
| 2351 | PINLYP |
| 2352 | PIP |
| 2353 | PIP4K2C |
| 2354 | PITPNA |
| 2355 | PITPNB |
| 2356 | PKDCC |
| 2357 | PKLR |
| 2358 | PKM |
| 2359 | PKP1 |
| 2360 | PLA1A |
| 2361 | PLA2G10 |
| 2362 | PLA2G12A |
| 2363 | PLA2G12B |

| | |
|---|---|
| 2364 | PLA2G15 |
| 2365 | PLA2G2A |
| 2366 | PLA2G2C |
| 2367 | PLA2G2D |
| 2368 | PLA2G2E |
| 2369 | PLA2G2F |
| 2370 | PLA2G3 |
| 2371 | PLA2G4B |
| 2372 | PLA2G7 |
| 2373 | PLAA |
| 2374 | PLAC1 |
| 2375 | PLAC1L |
| 2376 | PLAC9 |
| 2377 | PLAUR |
| 2378 | PLBD1 |
| 2379 | PLBD2 |
| 2380 | PLCB1 |
| 2381 | PLCD1 |
| 2382 | PLCG2 |
| 2383 | PLEC |
| 2384 | PLEK |
| 2385 | PLEKHA1 |
| 2386 | PLEKHA7 |
| 2387 | PLEKHH3 |
| 2388 | PLGLA |
| 2389 | PLGLB1 |
| 2390 | PLIN2 |
| 2391 | PLOD1 |
| 2392 | PLOD2 |
| 2393 | PLOD3 |
| 2394 | PLS1 |
| 2395 | PLTP |
| 2396 | PM20D1 |
| 2397 | PM20D2 |
| 2398 | PMCH |
| 2399 | PMCHL1 |
| 2400 | PMM2 |
| 2401 | PMP2 |
| 2402 | PMPCA |
| 2403 | PMVK |
| 2404 | PNLIP |
| 2405 | PNLIPRP1 |
| 2406 | PNLIPRP2 |

| | |
|---|---|
| 2407 | PNLIPRP3 |
| 2408 | PNOC |
| 2409 | PNP |
| 2410 | PNPO |
| 2411 | PODN |
| 2412 | PODNL1 |
| 2413 | POFUT1 |
| 2414 | POFUT2 |
| 2415 | POGLUT1 |
| 2416 | POLG |
| 2417 | POLG2 |
| 2418 | POMC |
| 2419 | PON1 |
| 2420 | PON2 |
| 2421 | PON3 |
| 2422 | POSTN |
| 2423 | POTEE |
| 2424 | POTEF |
| 2425 | PPA1 |
| 2426 | PPA2 |
| 2427 | PPBP |
| 2428 | PPCS |
| 2429 | PPFIBP2 |
| 2430 | PPIA |
| 2431 | PPIAL4A |
| 2432 | PPIB |
| 2433 | PPIC |
| 2434 | PPIL1 |
| 2435 | PPL |
| 2436 | PPP1CA |
| 2437 | PPP1CB |
| 2438 | PPP1R13L |
| 2439 | PPP1R1A |
| 2440 | PPP1R7 |
| 2441 | PPP2CA |
| 2442 | PPP2CB |
| 2443 | PPP2R1A |
| 2444 | PPP2R1B |
| 2445 | PPP2R4 |
| 2446 | PPP3CC |
| 2447 | PPT1 |
| 2448 | PPT2 |
| 2449 | PPY |

| | |
|---|---|
| 2450 | PRADC1 |
| 2451 | PRAP1 |
| 2452 | PRB1 |
| 2453 | PRB2 |
| 2454 | PRB3 |
| 2455 | PRB4 |
| 2456 | PRCP |
| 2457 | PRDX1 |
| 2458 | PRDX2 |
| 2459 | PRDX3 |
| 2460 | PRDX4 |
| 2461 | PRDX5 |
| 2462 | PRDX6 |
| 2463 | PRELP |
| 2464 | PRF1 |
| 2465 | PRG1 |
| 2466 | PRG2 |
| 2467 | PRG3 |
| 2468 | PRG4 |
| 2469 | PRH1 |
| 2470 | PRKACA |
| 2471 | PRKACB |
| 2472 | PRKAG1 |
| 2473 | PRKAG2 |
| 2474 | PRKAG3 |
| 2475 | PRKAR2A |
| 2476 | PRKAR2B |
| 2477 | PRKCA |
| 2478 | PRKCB |
| 2479 | PRKCD |
| 2480 | PRKCH |
| 2481 | PRKCI |
| 2482 | PRKCSH |
| 2483 | PRKCZ |
| 2484 | PRKRIP1 |
| 2485 | PRL |
| 2486 | PRLH |
| 2487 | PRND |
| 2488 | PRNT |
| 2489 | PROC |
| 2490 | PROK1 |
| 2491 | PROK2 |
| 2492 | PROL1 |

| | |
|---|---|
| 2493 | PROS1 |
| 2494 | PROSC |
| 2495 | PROZ |
| 2496 | PRPS2 |
| 2497 | PRRC2A |
| 2498 | PRSS1 |
| 2499 | PRSS12 |
| 2500 | PRSS16 |
| 2501 | PRSS2 |
| 2502 | PRSS21 |
| 2503 | PRSS22 |
| 2504 | PRSS23 |
| 2505 | PRSS27 |
| 2506 | PRSS3 |
| 2507 | PRSS33 |
| 2508 | PRSS35 |
| 2509 | PRSS36 |
| 2510 | PRSS37 |
| 2511 | PRSS38 |
| 2512 | PRSS3P2 |
| 2513 | PRSS41 |
| 2514 | PRSS42 |
| 2515 | PRSS48 |
| 2516 | PRSS50 |
| 2517 | PRSS53 |
| 2518 | PRSS54 |
| 2519 | PRSS56 |
| 2520 | PRSS57 |
| 2521 | PRSS58 |
| 2522 | PRTN3 |
| 2523 | PSAP |
| 2524 | PSAPL1 |
| 2525 | PSAT1 |
| 2526 | PSCA |
| 2527 | PSG1 |
| 2528 | PSG11 |
| 2529 | PSG2 |
| 2530 | PSG3 |
| 2531 | PSG4 |
| 2532 | PSG5 |
| 2533 | PSG6 |
| 2534 | PSG7 |
| 2535 | PSG8 |

| | |
|---|---|
| 2536 | PSG9 |
| 2537 | PSMA1 |
| 2538 | PSMA2 |
| 2539 | PSMA3 |
| 2540 | PSMA4 |
| 2541 | PSMA5 |
| 2542 | PSMA6 |
| 2543 | PSMA7 |
| 2544 | PSMA8 |
| 2545 | PSMB1 |
| 2546 | PSMB2 |
| 2547 | PSMB3 |
| 2548 | PSMB4 |
| 2549 | PSMB5 |
| 2550 | PSMB6 |
| 2551 | PSMB7 |
| 2552 | PSMB8 |
| 2553 | PSMB9 |
| 2554 | PSMC5 |
| 2555 | PSMC6 |
| 2556 | PSMD1 |
| 2557 | PSMD11 |
| 2558 | PSMD12 |
| 2559 | PSMD13 |
| 2560 | PSMD14 |
| 2561 | PSMD2 |
| 2562 | PSMD3 |
| 2563 | PSMD6 |
| 2564 | PSMD7 |
| 2565 | PSMD8 |
| 2566 | PSME1 |
| 2567 | PSME2 |
| 2568 | PSORS1C2 |
| 2569 | PSPN |
| 2570 | PTBP1 |
| 2571 | PTEN |
| 2572 | PTER |
| 2573 | PTGDS |
| 2574 | PTGES3 |
| 2575 | PTGR1 |
| 2576 | PTGR2 |
| 2577 | PTGS1 |
| 2578 | PTGS2 |

| | |
|---|---|
| 2579 | PTH |
| 2580 | PTH2 |
| 2581 | PTHLH |
| 2582 | PTMA |
| 2583 | PTN |
| 2584 | PTP4A1 |
| 2585 | PTP4A2 |
| 2586 | PTPN13 |
| 2587 | PTPN23 |
| 2588 | PTPN6 |
| 2589 | PTRHD1 |
| 2590 | PTX3 |
| 2591 | PTX4 |
| 2592 | PVALB |
| 2593 | PXDN |
| 2594 | PXDNL |
| 2595 | PYGB |
| 2596 | PYGL |
| 2597 | PYGM |
| 2598 | PYY |
| 2599 | PYY2 |
| 2600 | PZP |
| 2601 | QDPR |
| 2602 | QPCT |
| 2603 | QPRT |
| 2604 | QRFP |
| 2605 | R3HDML |
| 2606 | RAB10 |
| 2607 | RAB11B |
| 2608 | RAB13 |
| 2609 | RAB14 |
| 2610 | RAB15 |
| 2611 | RAB17 |
| 2612 | RAB18 |
| 2613 | RAB19 |
| 2614 | RAB1A |
| 2615 | RAB1B |
| 2616 | RAB22A |
| 2617 | RAB23 |
| 2618 | RAB25 |
| 2619 | RAB27A |
| 2620 | RAB27B |
| 2621 | RAB2A |

| | |
|---|---|
| 2622 | RAB2B |
| 2623 | RAB33B |
| 2624 | RAB34 |
| 2625 | RAB35 |
| 2626 | RAB3B |
| 2627 | RAB3D |
| 2628 | RAB3GAP1 |
| 2629 | RAB41 |
| 2630 | RAB43 |
| 2631 | RAB4A |
| 2632 | RAB5A |
| 2633 | RAB5B |
| 2634 | RAB5C |
| 2635 | RAB6A |
| 2636 | RAB7A |
| 2637 | RAB7L1 |
| 2638 | RAB8A |
| 2639 | RAB8B |
| 2640 | RAB9A |
| 2641 | RAC1 |
| 2642 | RAC2 |
| 2643 | RAC3 |
| 2644 | RAET1L |
| 2645 | RALB |
| 2646 | RALGAPA2 |
| 2647 | RAN |
| 2648 | RAP1A |
| 2649 | RAP1B |
| 2650 | RAP1GDS1 |
| 2651 | RAP2A |
| 2652 | RAP2B |
| 2653 | RAP2C |
| 2654 | RAPGEF3 |
| 2655 | RARRES2 |
| 2656 | RARS |
| 2657 | RASAL3 |
| 2658 | RASSF9 |
| 2659 | RBBP8NL |
| 2660 | RBBP9 |
| 2661 | RBKS |
| 2662 | RBL2 |
| 2663 | RBM44 |
| 2664 | RBMX |

| | |
|---|---|
| 2665 | RBP3 |
| 2666 | RBP4 |
| 2667 | RBP5 |
| 2668 | RCN1 |
| 2669 | RCN2 |
| 2670 | RCN3 |
| 2671 | RDH5 |
| 2672 | RECK |
| 2673 | REG1A |
| 2674 | REG1B |
| 2675 | REG3A |
| 2676 | REG3G |
| 2677 | REG4 |
| 2678 | RELN |
| 2679 | REN |
| 2680 | RENBP |
| 2681 | RESP18 |
| 2682 | RETN |
| 2683 | RETNLB |
| 2684 | RETSAT |
| 2685 | RFTN1 |
| 2686 | RFXANK |
| 2687 | RGMB |
| 2688 | RHEB |
| 2689 | RHOBTB3 |
| 2690 | RHOF |
| 2691 | RHOG |
| 2692 | RHOJ |
| 2693 | RHOQ |
| 2694 | RILPL2 |
| 2695 | RIMS2 |
| 2696 | RLN1 |
| 2697 | RLN2 |
| 2698 | RLN3 |
| 2699 | RNASE1 |
| 2700 | RNASE10 |
| 2701 | RNASE11 |
| 2702 | RNASE12 |
| 2703 | RNASE13 |
| 2704 | RNASE2 |
| 2705 | RNASE3 |
| 2706 | RNASE4 |
| 2707 | RNASE6 |

| | |
|---|---|
| 2708 | RNASE7 |
| 2709 | RNASE8 |
| 2710 | RNASE9 |
| 2711 | RNASET2 |
| 2712 | RND3 |
| 2713 | RNF11 |
| 2714 | RNLS |
| 2715 | RNPEP |
| 2716 | RP1L1 |
| 2717 | RP2 |
| 2718 | RPE |
| 2719 | RPL10A |
| 2720 | RPL11 |
| 2721 | RPL12 |
| 2722 | RPL13AP3 |
| 2723 | RPL14 |
| 2724 | RPL15 |
| 2725 | RPL22 |
| 2726 | RPL23 |
| 2727 | RPL23A |
| 2728 | RPL24 |
| 2729 | RPL26 |
| 2730 | RPL26L1 |
| 2731 | RPL27 |
| 2732 | RPL28 |
| 2733 | RPL3 |
| 2734 | RPL30 |
| 2735 | RPL31 |
| 2736 | RPL34 |
| 2737 | RPL35A |
| 2738 | RPL37A |
| 2739 | RPL39 |
| 2740 | RPL4 |
| 2741 | RPL5 |
| 2742 | RPL7 |
| 2743 | RPL7A |
| 2744 | RPLP0 |
| 2745 | RPLP1 |
| 2746 | RPLP2 |
| 2747 | RPS10 |
| 2748 | RPS11 |
| 2749 | RPS13 |
| 2750 | RPS14 |

| | |
|---|---|
| 2751 | RPS15A |
| 2752 | RPS16 |
| 2753 | RPS17 |
| 2754 | RPS18 |
| 2755 | RPS19 |
| 2756 | RPS2 |
| 2757 | RPS20 |
| 2758 | RPS25 |
| 2759 | RPS26 |
| 2760 | RPS27A |
| 2761 | RPS28 |
| 2762 | RPS29 |
| 2763 | RPS3 |
| 2764 | RPS3A |
| 2765 | RPS4X |
| 2766 | RPS5 |
| 2767 | RPS7 |
| 2768 | RPS8 |
| 2769 | RPS9 |
| 2770 | RPSA |
| 2771 | RPTN |
| 2772 | RRAS |
| 2773 | RRAS2 |
| 2774 | RREB1 |
| 2775 | RRM1 |
| 2776 | RRM2B |
| 2777 | RS1 |
| 2778 | RSPO1 |
| 2779 | RSPO3 |
| 2780 | RSPO4 |
| 2781 | RSPRY1 |
| 2782 | RSU1 |
| 2783 | RTBDN |
| 2784 | RTN3 |
| 2785 | RUNX1 |
| 2786 | RUSC2 |
| 2787 | RUVBL1 |
| 2788 | RUVBL2 |
| 2789 | S100A10 |
| 2790 | S100A11 |
| 2791 | S100A12 |
| 2792 | S100A13 |
| 2793 | S100A14 |

| | |
|---|---|
| 2794 | S100A16 |
| 2795 | S100A4 |
| 2796 | S100A6 |
| 2797 | S100A7 |
| 2798 | S100A8 |
| 2799 | S100A9 |
| 2800 | S100B |
| 2801 | S100P |
| 2802 | SAA1 |
| 2803 | SAA2 |
| 2804 | SAA4 |
| 2805 | SAFB2 |
| 2806 | SAMD1 |
| 2807 | SAMM50 |
| 2808 | SAR1A |
| 2809 | SARS |
| 2810 | SAT2 |
| 2811 | SBSN |
| 2812 | SBSPON |
| 2813 | SCEL |
| 2814 | SCG2 |
| 2815 | SCG3 |
| 2816 | SCG5 |
| 2817 | SCGB1A1 |
| 2818 | SCGB1C1 |
| 2819 | SCGB1D1 |
| 2820 | SCGB1D2 |
| 2821 | SCGB1D4 |
| 2822 | SCGB2A1 |
| 2823 | SCGB2A2 |
| 2824 | SCGB2B2 |
| 2825 | SCGB3A1 |
| 2826 | SCGB3A2 |
| 2827 | SCGN |
| 2828 | SCIN |
| 2829 | SCLT1 |
| 2830 | SCPEP1 |
| 2831 | SCRG1 |
| 2832 | SCRIB |
| 2833 | SCRN2 |
| 2834 | SCT |
| 2835 | SCUBE2 |
| 2836 | SDCBP |

| | |
|---|---|
| 2837 | SDCBP2 |
| 2838 | SDF2 |
| 2839 | SDF2L1 |
| 2840 | SDF4 |
| 2841 | SDHB |
| 2842 | SDSL |
| 2843 | SEC13 |
| 2844 | SEC14L2 |
| 2845 | SEC14L3 |
| 2846 | SELENBP1 |
| 2847 | SELM |
| 2848 | SELP |
| 2849 | SELT |
| 2850 | SEMA3A |
| 2851 | SEMA3B |
| 2852 | SEMA3C |
| 2853 | SEMA3D |
| 2854 | SEMA3E |
| 2855 | SEMA3F |
| 2856 | SEMA3G |
| 2857 | SEMG1 |
| 2858 | SEMG2 |
| 2859 | SEPN1 |
| 2860 | SEPP1 |
| 2861 | SERBP1 |
| 2862 | SERPINA1 |
| 2863 | SERPINA10 |
| 2864 | SERPINA11 |
| 2865 | SERPINA12 |
| 2866 | SERPINA13P |
| 2867 | SERPINA3 |
| 2868 | SERPINA4 |
| 2869 | SERPINA6 |
| 2870 | SERPINA7 |
| 2871 | SERPINA9 |
| 2872 | SERPINB1 |
| 2873 | SERPINB10 |
| 2874 | SERPINB11 |
| 2875 | SERPINB12 |
| 2876 | SERPINB13 |
| 2877 | SERPINB2 |
| 2878 | SERPINB3 |
| 2879 | SERPINB4 |

| | |
|---|---|
| 2880 | SERPINB5 |
| 2881 | SERPINB6 |
| 2882 | SERPINB7 |
| 2883 | SERPINB8 |
| 2884 | SERPINB9 |
| 2885 | SERPINC1 |
| 2886 | SERPIND1 |
| 2887 | SERPINE1 |
| 2888 | SERPINE3 |
| 2889 | SERPINF1 |
| 2890 | SERPING1 |
| 2891 | SERPINH1 |
| 2892 | SERPINI1 |
| 2893 | SERPINI2 |
| 2894 | SFRP2 |
| 2895 | SFRP5 |
| 2896 | SFTA2 |
| 2897 | SFTPA1 |
| 2898 | SFTPA2 |
| 2899 | SFTPB |
| 2900 | SFTPC |
| 2901 | SFTPD |
| 2902 | SGSH |
| 2903 | SH3BGRL |
| 2904 | SH3BGRL2 |
| 2905 | SH3BGRL3 |
| 2906 | SH3BP4 |
| 2907 | SH3D21 |
| 2908 | SH3GL2 |
| 2909 | SH3GL3 |
| 2910 | SH3GLB1 |
| 2911 | SHBG |
| 2912 | SHMT1 |
| 2913 | SHMT2 |
| 2914 | SHROOM2 |
| 2915 | SIAE |
| 2916 | SIL1 |
| 2917 | SIN3A |
| 2918 | SIPA1L3 |
| 2919 | SIRPD |
| 2920 | SKP1 |
| 2921 | SLC9A3R1 |
| 2922 | SLC9A3R2 |

| | |
|---|---|
| 2923 | SLIT1 |
| 2924 | SLIT3 |
| 2925 | SLK |
| 2926 | SLPI |
| 2927 | SLURP1 |
| 2928 | SMAP2 |
| 2929 | SMARCA4 |
| 2930 | SMC2 |
| 2931 | SMC3 |
| 2932 | SMOC1 |
| 2933 | SMOC2 |
| 2934 | SMPD1 |
| 2935 | SMPDL3A |
| 2936 | SMPDL3B |
| 2937 | SMR3A |
| 2938 | SMR3B |
| 2939 | SMS |
| 2940 | SMURF1 |
| 2941 | SNAP23 |
| 2942 | SNCA |
| 2943 | SNCG |
| 2944 | SND1 |
| 2945 | SNED1 |
| 2946 | SNF8 |
| 2947 | SNRNP25 |
| 2948 | SNRPB |
| 2949 | SNRPD1 |
| 2950 | SNRPD2 |
| 2951 | SNRPD3 |
| 2952 | SNRPE |
| 2953 | SNTB2 |
| 2954 | SNUPN |
| 2955 | SNX12 |
| 2956 | SNX18 |
| 2957 | SNX2 |
| 2958 | SNX29 |
| 2959 | SNX3 |
| 2960 | SNX9 |
| 2961 | SOD1 |
| 2962 | SOD2 |
| 2963 | SOD3 |
| 2964 | SOGA1 |
| 2965 | SOGA2 |

| | |
|---|---|
| 2966 | SORD |
| 2967 | SOST |
| 2968 | SOSTDC1 |
| 2969 | SOWAHA |
| 2970 | SP4 |
| 2971 | SPACA4 |
| 2972 | SPACA5 |
| 2973 | SPACA7 |
| 2974 | SPAG11A |
| 2975 | SPAG11B |
| 2976 | SPAG9 |
| 2977 | SPAM1 |
| 2978 | SPARCL1 |
| 2979 | SPATA20 |
| 2980 | SPATA6 |
| 2981 | SPEN |
| 2982 | SPESP1 |
| 2983 | SPHKAP |
| 2984 | SPINK1 |
| 2985 | SPINK13 |
| 2986 | SPINK14 |
| 2987 | SPINK2 |
| 2988 | SPINK4 |
| 2989 | SPINK5 |
| 2990 | SPINK6 |
| 2991 | SPINK7 |
| 2992 | SPINK8 |
| 2993 | SPINK9 |
| 2994 | SPINT1 |
| 2995 | SPINT3 |
| 2996 | SPINT4 |
| 2997 | SPOCK1 |
| 2998 | SPOCK2 |
| 2999 | SPOCK3 |
| 3000 | SPON1 |
| 3001 | SPON2 |
| 3002 | SPP1 |
| 3003 | SPP2 |
| 3004 | SPR |
| 3005 | SPRN |
| 3006 | SPRR1B |
| 3007 | SPRR3 |
| 3008 | SPTAN1 |

| | |
|---|---|
| 3009 | SPTBN1 |
| 3010 | SPTBN2 |
| 3011 | SPTBN4 |
| 3012 | SPTBN5 |
| 3013 | SQSTM1 |
| 3014 | SRA1 |
| 3015 | SRC |
| 3016 | SRGN |
| 3017 | SRI |
| 3018 | SRL |
| 3019 | SRP14 |
| 3020 | SRP9 |
| 3021 | SRPR |
| 3022 | SRSF1 |
| 3023 | SRSF2 |
| 3024 | SRSF7 |
| 3025 | SSBP1 |
| 3026 | SSC5D |
| 3027 | SSH2 |
| 3028 | SSPO |
| 3029 | SST |
| 3030 | ST13 |
| 3031 | ST13P4 |
| 3032 | STAG3 |
| 3033 | STATH |
| 3034 | STAU1 |
| 3035 | STC1 |
| 3036 | STC2 |
| 3037 | STK10 |
| 3038 | STK11 |
| 3039 | STK24 |
| 3040 | STK25 |
| 3041 | STMN1 |
| 3042 | STOM |
| 3043 | STRIP1 |
| 3044 | STUB1 |
| 3045 | STXBP1 |
| 3046 | STXBP2 |
| 3047 | STXBP3 |
| 3048 | STXBP4 |
| 3049 | SUB1 |
| 3050 | SUCLA2 |
| 3051 | SUCLG1 |

| | |
|---|---|
| 3052 | SULT2B1 |
| 3053 | SUMF1 |
| 3054 | SUMF2 |
| 3055 | SUMO3 |
| 3056 | SUPT20HL2 |
| 3057 | SVEP1 |
| 3058 | SVIP |
| 3059 | SYAP1 |
| 3060 | SYCN |
| 3061 | SYTL1 |
| 3062 | SZT2 |
| 3063 | TAB3 |
| 3064 | TAC1 |
| 3065 | TAC3 |
| 3066 | TAC4 |
| 3067 | TAF1B |
| 3068 | TAF6L |
| 3069 | TAGLN2 |
| 3070 | TALDO1 |
| 3071 | TAOK1 |
| 3072 | TARS |
| 3073 | TAX1BP1 |
| 3074 | TAX1BP3 |
| 3075 | TBC1D10A |
| 3076 | TBC1D15 |
| 3077 | TBC1D21 |
| 3078 | TBC1D4 |
| 3079 | TBCA |
| 3080 | TCEB2 |
| 3081 | TCEB3 |
| 3082 | TCN1 |
| 3083 | TCN2 |
| 3084 | TCP1 |
| 3085 | TCTN1 |
| 3086 | TDP2 |
| 3087 | TECTA |
| 3088 | TECTB |
| 3089 | TEKT3 |
| 3090 | TEPP |
| 3091 | TEX101 |
| 3092 | TEX14 |
| 3093 | TEX264 |
| 3094 | TFF1 |

| | |
|---|---|
| 3095 | TFF2 |
| 3096 | TFF3 |
| 3097 | TFG |
| 3098 | TFIP11 |
| 3099 | TFPI2 |
| 3100 | TG |
| 3101 | TGFB2 |
| 3102 | TGFBI |
| 3103 | TGM1 |
| 3104 | TGM2 |
| 3105 | TGM3 |
| 3106 | TGM4 |
| 3107 | TGS1 |
| 3108 | THAP11 |
| 3109 | THBS2 |
| 3110 | THBS3 |
| 3111 | THBS4 |
| 3112 | THEM6 |
| 3113 | THNSL2 |
| 3114 | THOC1 |
| 3115 | THPO |
| 3116 | THRAP3 |
| 3117 | THRB |
| 3118 | THSD4 |
| 3119 | TIAL1 |
| 3120 | TIAM2 |
| 3121 | TIMM8B |
| 3122 | TIMP1 |
| 3123 | TIMP3 |
| 3124 | TIMP4 |
| 3125 | TINAG |
| 3126 | TINAGL1 |
| 3127 | TIRAP |
| 3128 | TKT |
| 3129 | TLE2 |
| 3130 | TLL1 |
| 3131 | TLL2 |
| 3132 | TMEM155 |
| 3133 | TMSB4X |
| 3134 | TNC |
| 3135 | TNFAIP2 |
| 3136 | TNFAIP3 |
| 3137 | TNFAIP6 |

| | |
|---|---|
| 3138 | TNFRSF10C |
| 3139 | TNFRSF11B |
| 3140 | TNFRSF6B |
| 3141 | TNIK |
| 3142 | TNPO1 |
| 3143 | TNS4 |
| 3144 | TNXA |
| 3145 | TNXB |
| 3146 | TOLLIP |
| 3147 | TOM1 |
| 3148 | TOM1L1 |
| 3149 | TOM1L2 |
| 3150 | TOMM40 |
| 3151 | TOR1A |
| 3152 | TOR1B |
| 3153 | TOR2A |
| 3154 | TOR3A |
| 3155 | TP53I3 |
| 3156 | TP73-AS1 |
| 3157 | TPI1 |
| 3158 | TPM3 |
| 3159 | TPM4 |
| 3160 | TPMT |
| 3161 | TPPP3 |
| 3162 | TPRG1L |
| 3163 | TPSAB1 |
| 3164 | TPSB2 |
| 3165 | TPSD1 |
| 3166 | TPT1 |
| 3167 | TRAP1 |
| 3168 | TREH |
| 3169 | TREML4 |
| 3170 | TRH |
| 3171 | TRIM23 |
| 3172 | TRIM36 |
| 3173 | TRIM45 |
| 3174 | TRIP10 |
| 3175 | TRMT10A |
| 3176 | TRMT112 |
| 3177 | TSG101 |
| 3178 | TSHB |
| 3179 | TSKU |
| 3180 | TSLP |

| | |
|---|---|
| 3181 | TST |
| 3182 | TSTA3 |
| 3183 | TTBK2 |
| 3184 | TTC18 |
| 3185 | TTC38 |
| 3186 | TTN |
| 3187 | TTR |
| 3188 | TUB |
| 3189 | TUBA1A |
| 3190 | TUBA1B |
| 3191 | TUBA4A |
| 3192 | TUBB |
| 3193 | TUBB1 |
| 3194 | TUBB2A |
| 3195 | TUBB3 |
| 3196 | TUBB4A |
| 3197 | TUBB4B |
| 3198 | TUBB6 |
| 3199 | TUBB8 |
| 3200 | TUBGCP6 |
| 3201 | TUFM |
| 3202 | TUFT1 |
| 3203 | TULP1 |
| 3204 | TULP2 |
| 3205 | TULP3 |
| 3206 | TUT1 |
| 3207 | TWF2 |
| 3208 | TWSG1 |
| 3209 | TXLNA |
| 3210 | TXN |
| 3211 | TXNDC12 |
| 3212 | TXNDC16 |
| 3213 | TXNDC17 |
| 3214 | TXNDC5 |
| 3215 | TXNDC8 |
| 3216 | TXNL1 |
| 3217 | TXNRD1 |
| 3218 | TYK2 |
| 3219 | UACA |
| 3220 | UBA1 |
| 3221 | UBA52 |
| 3222 | UBAC1 |
| 3223 | UBASH3A |

| | |
|---|---|
| 3224 | UBB |
| 3225 | UBC |
| 3226 | UBE2D2 |
| 3227 | UBE2D3 |
| 3228 | UBE2G1 |
| 3229 | UBE2K |
| 3230 | UBE2L3 |
| 3231 | UBE2M |
| 3232 | UBE2N |
| 3233 | UBE2NL |
| 3234 | UBE2V1 |
| 3235 | UBE2V2 |
| 3236 | UBL3 |
| 3237 | UBN2 |
| 3238 | UBXN6 |
| 3239 | UCHL1 |
| 3240 | UCHL3 |
| 3241 | UCMA |
| 3242 | UCN |
| 3243 | UCN2 |
| 3244 | UCN3 |
| 3245 | UEVLD |
| 3246 | UFC1 |
| 3247 | UFM1 |
| 3248 | UFSP1 |
| 3249 | UGDH |
| 3250 | UGGT1 |
| 3251 | UGGT2 |
| 3252 | UGP2 |
| 3253 | ULBP1 |
| 3254 | ULBP3 |
| 3255 | UMOD |
| 3256 | UNC119 |
| 3257 | UPB1 |
| 3258 | UQCR10 |
| 3259 | UQCRC2 |
| 3260 | URM1 |
| 3261 | USPL1 |
| 3262 | UTP11L |
| 3263 | UTRN |
| 3264 | UTS2 |
| 3265 | UTS2D |
| 3266 | VASH1 |

| | |
|---|---|
| 3267 | VASP |
| 3268 | VAV3 |
| 3269 | VCAN |
| 3270 | VCL |
| 3271 | VCP |
| 3272 | VEGFB |
| 3273 | VEGFC |
| 3274 | VGF |
| 3275 | VIL1 |
| 3276 | VIM |
| 3277 | VIP |
| 3278 | VIT |
| 3279 | VMO1 |
| 3280 | VNN1 |
| 3281 | VNN2 |
| 3282 | VNN3 |
| 3283 | VPREB1 |
| 3284 | VPREB3 |
| 3285 | VPS13C |
| 3286 | VPS13D |
| 3287 | VPS25 |
| 3288 | VPS26A |
| 3289 | VPS28 |
| 3290 | VPS29 |
| 3291 | VPS35 |
| 3292 | VPS36 |
| 3293 | VPS37B |
| 3294 | VPS37C |
| 3295 | VPS37D |
| 3296 | VPS4A |
| 3297 | VPS4B |
| 3298 | VSTM2A |
| 3299 | VSTM2L |
| 3300 | VTA1 |
| 3301 | VTN |
| 3302 | VWA1 |
| 3303 | VWA2 |
| 3304 | VWA3A |
| 3305 | VWA5B1 |
| 3306 | VWA7 |
| 3307 | VWA8 |
| 3308 | VWC2 |
| 3309 | VWC2L |

| | |
|---|---|
| 3310 | VWCE |
| 3311 | VWDE |
| 3312 | WARS |
| 3313 | WAS |
| 3314 | WASF2 |
| 3315 | WASF3 |
| 3316 | WASL |
| 3317 | WDR1 |
| 3318 | WDR60 |
| 3319 | WFDC1 |
| 3320 | WFDC10A |
| 3321 | WFDC10B |
| 3322 | WFDC11 |
| 3323 | WFDC12 |
| 3324 | WFDC13 |
| 3325 | WFDC2 |
| 3326 | WFDC3 |
| 3327 | WFDC5 |
| 3328 | WFDC6 |
| 3329 | WFDC8 |
| 3330 | WFDC9 |
| 3331 | WFIKKN1 |
| 3332 | WFIKKN2 |
| 3333 | WIF1 |
| 3334 | WISP1 |
| 3335 | WISP2 |
| 3336 | WISP3 |
| 3337 | WIZ |
| 3338 | WNT10A |
| 3339 | WNT10B |
| 3340 | WNT11 |
| 3341 | WNT16 |
| 3342 | WNT2 |
| 3343 | WNT2B |
| 3344 | WNT3 |
| 3345 | WNT7B |
| 3346 | WNT8A |
| 3347 | WNT8B |
| 3348 | WNT9A |
| 3349 | WNT9B |
| 3350 | WWP1 |
| 3351 | WWP2 |
| 3352 | XCL1 |

| | |
|---|---|
| 3353 | XCL2 |
| 3354 | XDH |
| 3355 | XPA |
| 3356 | XPC |
| 3357 | XPNPEP1 |
| 3358 | XPNPEP2 |
| 3359 | XPNPEP3 |
| 3360 | XYLB |
| 3361 | YARS |
| 3362 | YBX1 |
| 3363 | YES1 |
| 3364 | YKT6 |
| 3365 | YWHAB |
| 3366 | YWHAE |
| 3367 | YWHAG |
| 3368 | YWHAH |
| 3369 | YWHAQ |
| 3370 | YWHAZ |
| 3371 | ZBTB18 |
| 3372 | ZBTB38 |
| 3373 | ZBTB8OS |
| 3374 | ZCCHC11 |
| 3375 | ZFC3H1 |
| 3376 | ZG16 |
| 3377 | ZG16B |
| 3378 | ZNF114 |
| 3379 | ZNF177 |
| 3380 | ZNF420 |
| 3381 | ZNF445 |
| 3382 | ZNF446 |
| 3383 | ZNF486 |
| 3384 | ZNF649 |
| 3385 | ZNF711 |
| 3386 | ZNF763 |
| 3387 | ZNHIT6 |
| 3388 | ZPBP |
| 3389 | ZPBP2 |
| 3390 | ZSCAN23 |
| 3391 | ZSWIM5 |

#### C. MEROPS Proteases (718)

| No. | Gene Name |
|---|---|
| 1 | AADAC |
| 2 | ABHD10 |
| 3 | ABHD11 |
| 4 | ABHD12 |
| 5 | ABHD12B |
| 6 | ABHD13 |
| 7 | ABHD14A |
| 8 | ABHD14A-ACY1 |
| 9 | ABHD14B |
| 10 | ABHD16A |
| 11 | ABHD16B |
| 12 | ABHD2 |
| 13 | ABHD4 |
| 14 | ABHD5 |
| 15 | ABHD6 |
| 16 | ABHD8 |
| 17 | ACE |
| 18 | ACE2 |
| 19 | ACHE |
| 20 | ACOT2 |
| 21 | ACOT4 |
| 22 | ACOT6 |
| 23 | ACR |
| 24 | ACY1 |
| 25 | ADAM1 |
| 26 | ADAM10 |
| 27 | ADAM11 |
| 28 | ADAM12 |
| 29 | ADAM15 |
| 30 | ADAM17 |
| 31 | ADAM18 |
| 32 | ADAM19 |
| 33 | ADAM2 |
| 34 | ADAM20 |
| 35 | ADAM21 |
| 36 | ADAM22 |
| 37 | ADAM23 |
| 38 | ADAM28 |
| 39 | ADAM29 |
| 40 | ADAM30 |
| 41 | ADAM32 |

| | |
|---|---|
| 42 | ADAM33 |
| 43 | ADAM3A |
| 44 | ADAM3B |
| 45 | ADAM7 |
| 46 | ADAM8 |
| 47 | ADAM9 |
| 48 | ADAMDEC1 |
| 49 | ADAMTS1 |
| 50 | ADAMTS10 |
| 51 | ADAMTS12 |
| 52 | ADAMTS13 |
| 53 | ADAMTS14 |
| 54 | ADAMTS15 |
| 55 | ADAMTS16 |
| 56 | ADAMTS17 |
| 57 | ADAMTS18 |
| 58 | ADAMTS19 |
| 59 | ADAMTS2 |
| 60 | ADAMTS20 |
| 61 | ADAMTS3 |
| 62 | ADAMTS4 |
| 63 | ADAMTS5 |
| 64 | ADAMTS6 |
| 65 | ADAMTS7 |
| 66 | ADAMTS8 |
| 67 | ADAMTS9 |
| 68 | ADGB |
| 69 | ADGRA3 |
| 70 | ADGRB1 |
| 71 | ADGRB2 |
| 72 | ADGRF4 |
| 73 | ADGRG1 |
| 74 | ADGRG2 |
| 75 | ADGRG7 |
| 76 | ADGRL2 |
| 77 | AEBP1 |
| 78 | AFG3L1P |
| 79 | AFG3L2 |
| 80 | AFMID |
| 81 | AGA |
| 82 | AGBL1 |
| 83 | AGBL2 |
| 84 | AGBL3 |

| | |
|---|---|
| 85 | AGBL4 |
| 86 | AGTPBP1 |
| 87 | ALG13 |
| 88 | AMDHD1 |
| 89 | AMZ1 |
| 90 | AMZ2 |
| 91 | ANPEP |
| 92 | APEH |
| 93 | ARHGAP21 |
| 94 | ARHGAP23 |
| 95 | ASAH1 |
| 96 | ASNS |
| 97 | ASPRV1 |
| 98 | ASRGL1 |
| 99 | ASTL |
| 100 | ATG4A |
| 101 | ATG4B |
| 102 | ATG4C |
| 103 | ATG4D |
| 104 | ATXN3 |
| 105 | ATXN3L |
| 106 | AZU1 |
| 107 | BAAT |
| 108 | BACE1 |
| 109 | BACE2 |
| 110 | BAI3 |
| 111 | BAP1 |
| 112 | BCHE |
| 113 | BLMH |
| 114 | BMP1 |
| 115 | BPHL |
| 116 | BRCC3 |
| 117 | C1R |
| 118 | C1RL |
| 119 | C1S |
| 120 | C2 |
| 121 | C21orf33 |
| 122 | C9orf3 |
| 123 | CAD |
| 124 | CAD |
| 125 | CAPN1 |
| 126 | CAPN10 |
| 127 | CAPN11 |

| | |
|---|---|
| 128 | CAPN12 |
| 129 | CAPN13 |
| 130 | CAPN14 |
| 131 | CAPN2 |
| 132 | CAPN3 |
| 133 | CAPN5 |
| 134 | CAPN6 |
| 135 | CAPN7 |
| 136 | CAPN8 |
| 137 | CAPN9 |
| 138 | CAPN9 |
| 139 | CARD8 |
| 140 | CASP1 |
| 141 | CASP10 |
| 142 | CASP12 |
| 143 | CASP14 |
| 144 | CASP2 |
| 145 | CASP3 |
| 146 | CASP4 |
| 147 | CASP5 |
| 148 | CASP6 |
| 149 | CASP7 |
| 150 | CASP8 |
| 151 | CASP9 |
| 152 | CD97 |
| 153 | CEL |
| 154 | CELA1 |
| 155 | CELA2A |
| 156 | CELA2B |
| 157 | CELA3A |
| 158 | CELA3B |
| 159 | CELSR1 |
| 160 | CELSR2 |
| 161 | CES1 |
| 162 | CES1P1 |
| 163 | CES2 |
| 164 | CES3 |
| 165 | CES5A |
| 166 | CFB |
| 167 | CFD |
| 168 | CFH |
| 169 | CFI |
| 170 | CFLAR |

| | |
|---|---|
| 171 | CLCA1 |
| 172 | CLPP |
| 173 | CMA1 |
| 174 | CNDP1 |
| 175 | CNDP2 |
| 176 | COPS5 |
| 177 | COPS6 |
| 178 | CORIN |
| 179 | CPA1 |
| 180 | CPA2 |
| 181 | CPA3 |
| 182 | CPA4 |
| 183 | CPA5 |
| 184 | CPA6 |
| 185 | CPB1 |
| 186 | CPB2 |
| 187 | CPD |
| 188 | CPE |
| 189 | CPM |
| 190 | CPN1 |
| 191 | CPO |
| 192 | CPS1 |
| 193 | CPVL |
| 194 | CPXM1 |
| 195 | CPXM2 |
| 196 | CPZ |
| 197 | CRMP1 |
| 198 | CTPS1 |
| 199 | CTRB1 |
| 200 | CTRB2 |
| 201 | CTRC |
| 202 | CTRL |
| 203 | CTSA |
| 204 | CTSB |
| 205 | CTSC |
| 206 | CTSD |
| 207 | CTSE |
| 208 | CTSF |
| 209 | CTSG |
| 210 | CTSH |
| 211 | CTSK |
| 212 | CTSL1 |
| 213 | CTSL1P1 |

| | |
|---|---|
| 214 | CTSL1P2 |
| 215 | CTSO |
| 216 | CTSS |
| 217 | CTSV |
| 218 | CTSW |
| 219 | CTSZ |
| 220 | CYLD |
| 221 | CYMP |
| 222 | DAG1 |
| 223 | DDI1 |
| 224 | DDI2 |
| 225 | DESI2 |
| 226 | DHH |
| 227 | DNPEP |
| 228 | DPEP1 |
| 229 | DPEP2 |
| 230 | DPEP3 |
| 231 | DPP10 |
| 232 | DPP3 |
| 233 | DPP4 |
| 234 | DPP6 |
| 235 | DPP7 |
| 236 | DPP8 |
| 237 | DPP9 |
| 238 | DPYS |
| 239 | DPYSL2 |
| 240 | DPYSL3 |
| 241 | DPYSL4 |
| 242 | DPYSL5 |
| 243 | ECE1 |
| 244 | ECE2 |
| 245 | ECEL1 |
| 246 | EIF3F |
| 247 | EIF3H |
| 248 | ELANE |
| 249 | ELTD1 |
| 250 | EMR1 |
| 251 | EMR2 |
| 252 | EMR3 |
| 253 | EMR4P |
| 254 | ENPEP |
| 255 | EPB42 |
| 256 | EPHX1 |

| | |
|---|---|
| 257 | EPHX2 |
| 258 | EPHX3 |
| 259 | EPHX4 |
| 260 | ERAP1 |
| 261 | ERAP2 |
| 262 | ERMP1 |
| 263 | ESD |
| 264 | ESPL1 |
| 265 | F10 |
| 266 | F11 |
| 267 | F12 |
| 268 | F13A1 |
| 269 | F2 |
| 270 | F7 |
| 271 | F9 |
| 272 | FAM105A |
| 273 | FAM105B |
| 274 | FAM108A1 |
| 275 | FAM108B1 |
| 276 | FAM108C1 |
| 277 | FAP |
| 278 | FOLH1 |
| 279 | FOLH1B |
| 280 | FURIN |
| 281 | GDA |
| 282 | GFPT1 |
| 283 | GFPT2 |
| 284 | GGH |
| 285 | GGT1 |
| 286 | GGT2 |
| 287 | GGT5 |
| 288 | GGT6 |
| 289 | GGT7 |
| 290 | GGTLC2 |
| 291 | GMPS |
| 292 | GPR110 |
| 293 | GPR112 |
| 294 | GPR113 |
| 295 | GPR114 |
| 296 | GPR116 |
| 297 | GPR126 |
| 298 | GPR133 |
| 299 | GPR144 |

| | |
|---|---|
| 300 | GPR97 |
| 301 | GZMA |
| 302 | GZMB |
| 303 | GZMH |
| 304 | GZMK |
| 305 | GZMM |
| 306 | HABP2 |
| 307 | HGF |
| 308 | HGFAC |
| 309 | HM13 |
| 310 | HP |
| 311 | HPN |
| 312 | HPR |
| 313 | HTRA1 |
| 314 | HTRA2 |
| 315 | HTRA3 |
| 316 | HTRA4 |
| 317 | IDE |
| 318 | IHH |
| 319 | IMMP1L |
| 320 | IMMP2L |
| 321 | JOSD1 |
| 322 | JOSD2 |
| 323 | KEL |
| 324 | KLK1 |
| 325 | KLK10 |
| 326 | KLK11 |
| 327 | KLK12 |
| 328 | KLK13 |
| 329 | KLK14 |
| 330 | KLK15 |
| 331 | KLK2 |
| 332 | KLK3 |
| 333 | KLK4 |
| 334 | KLK5 |
| 335 | KLK6 |
| 336 | KLK7 |
| 337 | KLK8 |
| 338 | KLK9 |
| 339 | KLKB1 |
| 340 | KLKP1 |
| 341 | KY |
| 342 | LACTB |

| | |
|---|---|
| 343 | LAP3 |
| 344 | LGMN |
| 345 | LGMN2P |
| 346 | LIPA |
| 347 | LIPE |
| 348 | LIPF |
| 349 | LIPJ |
| 350 | LMLN |
| 351 | LNPEP |
| 352 | LONP1 |
| 353 | LONP2 |
| 354 | LPA |
| 355 | LPHN1 |
| 356 | LPHN3 |
| 357 | LTA4H |
| 358 | LTF |
| 359 | LYPLA2 |
| 360 | MALT1 |
| 361 | MASP1 |
| 362 | MASP1 |
| 363 | MASP2 |
| 364 | MBTPS1 |
| 365 | MBTPS2 |
| 366 | MEP1A |
| 367 | MEP1B |
| 368 | MEST |
| 369 | METAP1 |
| 370 | METAP2 |
| 371 | MFI2 |
| 372 | MFI2 |
| 373 | MGLL |
| 374 | MINDY1 |
| 375 | MINDY2 |
| 376 | MIPEP |
| 377 | MME |
| 378 | MMEL1 |
| 379 | MMP1 |
| 380 | MMP10 |
| 381 | MMP11 |
| 382 | MMP12 |
| 383 | MMP13 |
| 384 | MMP14 |
| 385 | MMP15 |

| | |
|---|---|
| 386 | MMP16 |
| 387 | MMP17 |
| 388 | MMP19 |
| 389 | MMP2 |
| 390 | MMP20 |
| 391 | MMP21 |
| 392 | MMP23B |
| 393 | MMP24 |
| 394 | MMP25 |
| 395 | MMP26 |
| 396 | MMP27 |
| 397 | MMP28 |
| 398 | MMP3 |
| 399 | MMP7 |
| 400 | MMP8 |
| 401 | MMP9 |
| 402 | MMPL1~withdrawn |
| 403 | MPND |
| 404 | MPP4 |
| 405 | MST1 |
| 406 | MST1P9 |
| 407 | MUC1 |
| 408 | MUC3B |
| 409 | MYSM1 |
| 410 | NAAA |
| 411 | NAALAD2 |
| 412 | NAALADL1 |
| 413 | NAALADL2 |
| 414 | NAPSA |
| 415 | NAPSB |
| 416 | NCEH1 |
| 417 | NCLN |
| 418 | NDRG1 |
| 419 | NDRG2 |
| 420 | NDRG3 |
| 421 | NDRG4 |
| 422 | NEK2P1 |
| 423 | NLGN1 |
| 424 | NLGN2 |
| 425 | NLGN3 |
| 426 | NLGN4X |
| 427 | NLGN4Y |
| 428 | NLN |

| | |
|---|---|
| 429 | NLRP1 |
| 430 | NPEPL1 |
| 431 | NPEPPS |
| 432 | NPEPPS |
| 433 | NRD1/NRDC |
| 434 | NRDC |
| 435 | NRIP2 |
| 436 | NRIP3 |
| 437 | NUP98 |
| 438 | OMA1 |
| 439 | OTUB1 |
| 440 | OTUB2 |
| 441 | OTUD1 |
| 442 | OTUD3 |
| 443 | OTUD4 |
| 444 | OTUD5 |
| 445 | OTUD6B |
| 446 | OTUD7A |
| 447 | OTUD7B |
| 448 | OVCH1 |
| 449 | OVCH2 |
| 450 | PA2G4 |
| 451 | PAMR1 |
| 452 | PAN2 |
| 453 | PAPPA |
| 454 | PAPPA2 |
| 455 | PARK7 |
| 456 | PARL |
| 457 | PCSK1 |
| 458 | PCSK2 |
| 459 | PCSK4 |
| 460 | PCSK5 |
| 461 | PCSK6 |
| 462 | PCSK7 |
| 463 | PCSK9 |
| 464 | PDDC1 |
| 465 | PEPD |
| 466 | PFAS |
| 467 | PGA3 |
| 468 | PGA4 |
| 469 | PGA5 |
| 470 | PGC |
| 471 | PGPEP1 |

| | |
|---|---|
| 472 | PGPEP1L |
| 473 | PHEX |
| 474 | PIDD1 |
| 475 | PIDD1 |
| 476 | PIGK |
| 477 | PITRM1 |
| 478 | PKD1 |
| 479 | PKD1L2 |
| 480 | PKD1L3 |
| 481 | PLAT |
| 482 | PLAU |
| 483 | PLBD2 |
| 484 | PLG |
| 485 | PM20D1 |
| 486 | PM20D2 |
| 487 | PMPCA |
| 488 | PMPCB |
| 489 | PPAT |
| 490 | PPME1 |
| 491 | PRCP |
| 492 | PREP |
| 493 | PREPL |
| 494 | PROC |
| 495 | PROZ |
| 496 | PRSS1 |
| 497 | PRSS12 |
| 498 | PRSS16 |
| 499 | PRSS2 |
| 500 | PRSS21 |
| 501 | PRSS22 |
| 502 | PRSS23 |
| 503 | PRSS27 |
| 504 | PRSS3 |
| 505 | PRSS30P |
| 506 | PRSS33 |
| 507 | PRSS35 |
| 508 | PRSS36 |
| 509 | PRSS36 |
| 510 | PRSS36 |
| 511 | PRSS41 |
| 512 | PRSS43 |
| 513 | PRSS44 |
| 514 | PRSS53 |

| | |
|---|---|
| 515 | PRSS53 |
| 516 | PRSS55 |
| 517 | PRSS56 |
| 518 | PRSS57 |
| 519 | PRSS8 |
| 520 | PRTN3 |
| 521 | PSEN1 |
| 522 | PSEN2 |
| 523 | PSMA1 |
| 524 | PSMA2 |
| 525 | PSMA3 |
| 526 | PSMA4 |
| 527 | PSMA5 |
| 528 | PSMA6 |
| 529 | PSMA7 |
| 530 | PSMA8 |
| 531 | PSMB1 |
| 532 | PSMB10 |
| 533 | PSMB11 |
| 534 | PSMB2 |
| 535 | PSMB3 |
| 536 | PSMB3P |
| 537 | PSMB3P2 |
| 538 | PSMB4 |
| 539 | PSMB5 |
| 540 | PSMB6 |
| 541 | PSMB7 |
| 542 | PSMB8 |
| 543 | PSMB9 |
| 544 | PSMD14 |
| 545 | PSMD7 |
| 546 | QPCT |
| 547 | QPCTL |
| 548 | RBP3 |
| 549 | RCE1 |
| 550 | REN |
| 551 | RHBDD1 |
| 552 | RHBDD2 |
| 553 | RHBDD3 |
| 554 | RHBDF1 |
| 555 | RHBDF2 |
| 556 | RHBDL1 |
| 557 | RHBDL2 |

| | |
|---|---|
| 558 | RHBDL4~withdrawn |
| 559 | RNPEP |
| 560 | RNPEPL1 |
| 561 | SCPEP1 |
| 562 | SCRN1 |
| 563 | SCRN2 |
| 564 | SCRN3 |
| 565 | SEC11A |
| 566 | SEC11B |
| 567 | SEC11C |
| 568 | SENP1 |
| 569 | SENP2 |
| 570 | SENP3 |
| 571 | SENP5 |
| 572 | SENP6 |
| 573 | SENP7 |
| 574 | SENP8 |
| 575 | SERHL |
| 576 | SHH |
| 577 | SOLH |
| 578 | SPG7 |
| 579 | SPPL2A |
| 580 | SPPL2B |
| 581 | SPPL2C |
| 582 | SPPL3 |
| 583 | SPRTN |
| 584 | ST14 |
| 585 | STAMBP |
| 586 | STAMBPL1 |
| 587 | STC2 |
| 588 | SUPT16H |
| 589 | TAF2 |
| 590 | TASP1 |
| 591 | TCAF1 |
| 592 | TCAF2 |
| 593 | TEX30 |
| 594 | TF |
| 595 | TFR2 |
| 596 | TFRC |
| 597 | TG |
| 598 | TGM1 |
| 599 | TGM2 |
| 600 | TGM3 |

| | |
|---|---|
| 601 | TGM4 |
| 602 | TGM5 |
| 603 | TGM6 |
| 604 | TGM7 |
| 605 | THOP1 |
| 606 | TINAG |
| 607 | TINAGL1 |
| 608 | TLL1 |
| 609 | TLL2 |
| 610 | TMPRSS11A |
| 611 | TMPRSS11B |
| 612 | TMPRSS11D |
| 613 | TMPRSS11E |
| 614 | TMPRSS11F |
| 615 | TMPRSS12 |
| 616 | TMPRSS13 |
| 617 | TMPRSS15 |
| 618 | TMPRSS2 |
| 619 | TMPRSS3 |
| 620 | TMPRSS4 |
| 621 | TMPRSS5 |
| 622 | TMPRSS6 |
| 623 | TMPRSS7 |
| 624 | TMPRSS9 |
| 625 | TMPRSS9 |
| 626 | TMPRSS9 |
| 627 | TNFAIP3 |
| 628 | TPP1 |
| 629 | TPP2 |
| 630 | TPSAB1 |
| 631 | TPSD1 |
| 632 | TPSG1 |
| 633 | TRABD2A |
| 634 | TRABD2B |
| 635 | TRHDE |
| 636 | TYSND1 |
| 637 | UCHL1 |
| 638 | UCHL3 |
| 639 | UCHL5 |
| 640 | UFSP1 |
| 641 | UFSP2 |
| 642 | UQCRC1 |
| 643 | UQCRC2 |

| | |
|---|---|
| 644 | USP1 |
| 645 | USP10 |
| 646 | USP11 |
| 647 | USP12 |
| 648 | USP12 |
| 649 | USP13 |
| 650 | USP14 |
| 651 | USP15 |
| 652 | USP16 |
| 653 | USP17 |
| 654 | USP17L12 |
| 655 | USP17L15 |
| 656 | USP17L17 |
| 657 | USP17L18 |
| 658 | USP17L7 |
| 659 | USP18 |
| 660 | USP19 |
| 661 | USP19 |
| 662 | USP2 |
| 663 | USP20 |
| 664 | USP21 |
| 665 | USP22 |
| 666 | USP22 |
| 667 | USP24 |
| 668 | USP25 |
| 669 | USP26 |
| 670 | USP27X |
| 671 | USP28 |
| 672 | USP28 |
| 673 | USP29 |
| 674 | USP3 |
| 675 | USP30 |
| 676 | USP31 |
| 677 | USP31 |
| 678 | USP32 |
| 679 | USP33 |
| 680 | USP34 |
| 681 | USP35 |
| 682 | USP36 |
| 683 | USP37 |
| 684 | USP38 |
| 685 | USP39 |
| 686 | USP4 |

| | |
|---|---|
| 687 | USP40 |
| 688 | USP41 |
| 689 | USP42 |
| 690 | USP43 |
| 691 | USP44 |
| 692 | USP45 |
| 693 | USP46 |
| 694 | USP47 |
| 695 | USP48 |
| 696 | USP49 |
| 697 | USP5 |
| 698 | USP50 |
| 699 | USP51 |
| 700 | USP53 |
| 701 | USP54 |
| 702 | USP6 |
| 703 | USP7 |
| 704 | USP8 |
| 705 | USP8P1 |
| 706 | USP9X |
| 707 | USP9Y |
| 708 | USPL1 |
| 709 | VCPIP1 |
| 710 | XPNPEP1 |
| 711 | XPNPEP2 |
| 712 | XPNPEP3 |
| 713 | XRCC6BP1 |
| 714 | YME1L1 |
| 715 | YOD1 |
| 716 | ZMPSTE24 |
| 717 | ZRANB1 |
| 718 | ZUP1 |

**D. Surfaceome x Proteases (1**

| No. | Gene Name |
|---|---|
| 1 | AADAC |
| 2 | ABHD12 |
| 3 | ABHD13 |
| 4 | ABHD14A |
| 5 | ABHD16A |
| 6 | ABHD2 |
| 7 | ABHD6 |
| 8 | ACE |
| 9 | ACE2 |
| 10 | ACHE |
| 11 | ADAM10 |
| 12 | ADAM11 |
| 13 | ADAM12 |
| 14 | ADAM15 |
| 15 | ADAM17 |
| 16 | ADAM18 |
| 17 | ADAM19 |
| 18 | ADAM2 |
| 19 | ADAM20 |
| 20 | ADAM21 |
| 21 | ADAM22 |
| 22 | ADAM23 |
| 23 | ADAM28 |
| 24 | ADAM29 |
| 25 | ADAM30 |
| 26 | ADAM32 |
| 27 | ADAM33 |
| 28 | ADAM7 |
| 29 | ADAM8 |
| 30 | ADAM9 |
| 31 | ADAMTS13 |
| 32 | ADAMTS7 |
| 33 | AFG3L2 |
| 34 | ANPEP |
| 35 | ASPRV1 |
| 36 | BACE1 |
| 37 | BAI3 |
| 38 | CAPN5 |
| 39 | CD97 |
| 40 | CORIN |
| 41 | CPD |

| | |
|---|---|
| 42 | CPM |
| 43 | CTSG |
| 44 | DAG1 |
| 45 | DPP10 |
| 46 | DPP4 |
| 47 | DPP6 |
| 48 | ECE1 |
| 49 | ECE2 |
| 50 | ECEL1 |
| 51 | ELANE |
| 52 | EMR1 |
| 53 | EMR2 |
| 54 | EMR4P |
| 55 | ENPEP |
| 56 | EPHX1 |
| 57 | EPHX4 |
| 58 | ERAP1 |
| 59 | ERAP2 |
| 60 | F10 |
| 61 | FAP |
| 62 | FOLH1 |
| 63 | FURIN |
| 64 | GGT1 |
| 65 | GGT5 |
| 66 | GGT6 |
| 67 | GGT7 |
| 68 | GGTLC2 |
| 69 | GPR110 |
| 70 | GPR112 |
| 71 | GPR116 |
| 72 | GPR126 |
| 73 | GPR133 |
| 74 | GPR144 |
| 75 | HPN |
| 76 | HTRA2 |
| 77 | IDE |
| 78 | IMMP2L |
| 79 | KEL |
| 80 | LNPEP |
| 81 | LPHN1 |
| 82 | MBTPS1 |
| 83 | MBTPS2 |
| 84 | MEP1A |

| | |
|---|---|
| 85 | MEP1B |
| 86 | MEST |
| 87 | MFI2 |
| 88 | MME |
| 89 | MMEL1 |
| 90 | MMP14 |
| 91 | MMP15 |
| 92 | MMP16 |
| 93 | MMP21 |
| 94 | MMP24 |
| 95 | MUC1 |
| 96 | NAALAD2 |
| 97 | NAALADL1 |
| 98 | NAALADL2 |
| 99 | NCEH1 |
| 100 | NLGN1 |
| 101 | NLGN2 |
| 102 | NLGN3 |
| 103 | NLGN4X |
| 104 | NLGN4Y |
| 105 | OMA1 |
| 106 | PCSK5 |
| 107 | PCSK6 |
| 108 | PCSK7 |
| 109 | PCSK9 |
| 110 | PHEX |
| 111 | PIGK |
| 112 | PKD1 |
| 113 | PKD1L2 |
| 114 | PKD1L3 |
| 115 | PLAT |
| 116 | PLAU |
| 117 | PLG |
| 118 | PRSS8 |
| 119 | PSEN1 |
| 120 | PSEN2 |
| 121 | QPCTL |
| 122 | RHBDD1 |
| 123 | RHBDD3 |
| 124 | RHBDF1 |
| 125 | RHBDF2 |
| 126 | RHBDL1 |
| 127 | SEC11A |

| | |
|---|---|
| 128 | SEC11C |
| 129 | SHH |
| 130 | SPG7 |
| 131 | SPPL2A |
| 132 | SPPL3 |
| 133 | ST14 |
| 134 | STAMBP |
| 135 | TF |
| 136 | TFR2 |
| 137 | TFRC |
| 138 | TMPRSS11A |
| 139 | TMPRSS11B |
| 140 | TMPRSS11D |
| 141 | TMPRSS11E |
| 142 | TMPRSS11F |
| 143 | TMPRSS13 |
| 144 | TMPRSS15 |
| 145 | TMPRSS2 |
| 146 | TMPRSS3 |
| 147 | TMPRSS4 |
| 148 | TMPRSS5 |
| 149 | TMPRSS6 |
| 150 | TMPRSS7 |
| 151 | TMPRSS9 |
| 152 | TPSG1 |
| 153 | TRABD2B |
| 154 | TRHDE |
| 155 | USP14 |
| 156 | USP19 |
| 157 | USP30 |
| 158 | YME1L1 |
| 159 | ZMPSTE24 |

**E. Secretome x Proteases (271) x Proteases (159)**

| No. | Gene Name |
|---|---|
| 1 | ABHD14B |
| 2 | ABHD8 |
| 3 | ACOT2 |
| 4 | ACR |
| 5 | ACY1 |
| 6 | ADAMDEC1 |
| 7 | ADAMTS1 |
| 8 | ADAMTS10 |
| 9 | ADAMTS12 |
| 10 | ADAMTS14 |
| 11 | ADAMTS15 |
| 12 | ADAMTS16 |
| 13 | ADAMTS17 |
| 14 | ADAMTS18 |
| 15 | ADAMTS19 |
| 16 | ADAMTS2 |
| 17 | ADAMTS20 |
| 18 | ADAMTS3 |
| 19 | ADAMTS4 |
| 20 | ADAMTS5 |
| 21 | ADAMTS6 |
| 22 | ADAMTS8 |
| 23 | ADAMTS9 |
| 24 | AEBP1 |
| 25 | AGA |
| 26 | APEH |
| 27 | ARHGAP23 |
| 28 | ASAH1 |
| 29 | ASTL |
| 30 | ATG4C |
| 31 | AZU1 |
| 32 | BCHE |
| 33 | BLMH |
| 34 | BMP1 |
| 35 | BPHL |
| 36 | C1R |
| 37 | C1RL |
| 38 | C1S |
| 39 | C2 |
| 40 | CAD |
| 41 | CAPN1 |
| 42 | CAPN2 |
| 43 | CAPN7 |
| 44 | CASP1 |
| 45 | CASP14 |
| 46 | CEL |
| 47 | CELA1 |
| 48 | CELA2A |
| 49 | CELA2B |
| 50 | CELA3A |
| 51 | CELA3B |
| 52 | CES1P1 |
| 53 | CES2 |
| 54 | CES3 |
| 55 | CES5A |
| 56 | CFB |
| 57 | CFD |
| 58 | CFH |
| 59 | CFI |
| 60 | CLCA1 |
| 61 | CMA1 |
| 62 | CNDP1 |
| 63 | CNDP2 |
| 64 | COPS6 |
| 65 | CPA1 |
| 66 | CPA2 |
| 67 | CPA3 |
| 68 | CPA4 |
| 69 | CPA5 |
| 70 | CPA6 |
| 71 | CPB1 |
| 72 | CPB2 |
| 73 | CPE |
| 74 | CPN1 |
| 75 | CPVL |
| 76 | CPXM1 |
| 77 | CPXM2 |
| 78 | CPZ |
| 79 | CTRB1 |
| 80 | CTRB2 |
| 81 | CTRC |

| | |
|---|---|
| 82 | CTRL |
| 83 | CTSA |
| 84 | CTSB |
| 85 | CTSC |
| 86 | CTSD |
| 87 | CTSE |
| 88 | CTSF |
| 89 | CTSH |
| 90 | CTSK |
| 91 | CTSL1 |
| 92 | CTSO |
| 93 | CTSS |
| 94 | CTSW |
| 95 | CTSZ |
| 96 | DHH |
| 97 | DNPEP |
| 98 | DPEP1 |
| 99 | DPEP2 |
| 100 | DPEP3 |
| 101 | DPP3 |
| 102 | DPP7 |
| 103 | DPYS |
| 104 | DPYSL2 |
| 105 | DPYSL3 |
| 106 | EIF3H |
| 107 | EPHX2 |
| 108 | EPHX3 |
| 109 | ESD |
| 110 | F11 |
| 111 | F12 |
| 112 | F13A1 |
| 113 | F2 |
| 114 | F7 |
| 115 | F9 |
| 116 | FAM108A1 |
| 117 | FAM108B1 |
| 118 | FOLH1B |
| 119 | GDA |
| 120 | GFPT1 |
| 121 | GGH |
| 122 | GZMA |
| 123 | GZMB |
| 124 | GZMH |
| 125 | GZMK |
| 126 | GZMM |
| 127 | HABP2 |
| 128 | HGF |
| 129 | HGFAC |
| 130 | HP |
| 131 | HPR |
| 132 | HTRA1 |
| 133 | HTRA3 |
| 134 | HTRA4 |
| 135 | IHH |
| 136 | KLK1 |
| 137 | KLK10 |
| 138 | KLK11 |
| 139 | KLK12 |
| 140 | KLK13 |
| 141 | KLK14 |
| 142 | KLK15 |
| 143 | KLK2 |
| 144 | KLK3 |
| 145 | KLK4 |
| 146 | KLK5 |
| 147 | KLK6 |
| 148 | KLK7 |
| 149 | KLK8 |
| 150 | KLK9 |
| 151 | KLKB1 |
| 152 | LAP3 |
| 153 | LGMN |
| 154 | LIPA |
| 155 | LIPF |
| 156 | LPA |
| 157 | LTA4H |
| 158 | LTF |
| 159 | LYPLA2 |
| 160 | MASP1 |
| 161 | MASP2 |
| 162 | MMP1 |
| 163 | MMP10 |
| 164 | MMP11 |

| | |
|---|---|
| 165 | MMP12 |
| 166 | MMP13 |
| 167 | MMP17 |
| 168 | MMP19 |
| 169 | MMP2 |
| 170 | MMP20 |
| 171 | MMP25 |
| 172 | MMP26 |
| 173 | MMP27 |
| 174 | MMP28 |
| 175 | MMP3 |
| 176 | MMP7 |
| 177 | MMP8 |
| 178 | MMP9 |
| 179 | NAAA |
| 180 | NAPSA |
| 181 | NDRG1 |
| 182 | NDRG2 |
| 183 | NDRG3 |
| 184 | NPEPPS |
| 185 | OTUB1 |
| 186 | OVCH1 |
| 187 | OVCH2 |
| 188 | PA2G4 |
| 189 | PAMR1 |
| 190 | PAPPA |
| 191 | PAPPA2 |
| 192 | PARK7 |
| 193 | PCSK1 |
| 194 | PCSK2 |
| 195 | PDDC1 |
| 196 | PEPD |
| 197 | PFAS |
| 198 | PGA3 |
| 199 | PGA4 |
| 200 | PGA5 |
| 201 | PGC |
| 202 | PLBD2 |
| 203 | PM20D1 |
| 204 | PM20D2 |
| 205 | PMPCA |
| 206 | PRCP |
| 207 | PROC |
| 208 | PROZ |
| 209 | PRSS1 |
| 210 | PRSS12 |
| 211 | PRSS16 |
| 212 | PRSS2 |
| 213 | PRSS21 |
| 214 | PRSS22 |
| 215 | PRSS23 |
| 216 | PRSS27 |
| 217 | PRSS3 |
| 218 | PRSS33 |
| 219 | PRSS35 |
| 220 | PRSS36 |
| 221 | PRSS41 |
| 222 | PRSS53 |
| 223 | PRSS56 |
| 224 | PRSS57 |
| 225 | PRTN3 |
| 226 | PSMA1 |
| 227 | PSMA2 |
| 228 | PSMA3 |
| 229 | PSMA4 |
| 230 | PSMA5 |
| 231 | PSMA6 |
| 232 | PSMA7 |
| 233 | PSMA8 |
| 234 | PSMB1 |
| 235 | PSMB2 |
| 236 | PSMB3 |
| 237 | PSMB4 |
| 238 | PSMB5 |
| 239 | PSMB6 |
| 240 | PSMB7 |
| 241 | PSMB8 |
| 242 | PSMB9 |
| 243 | PSMD14 |
| 244 | PSMD7 |
| 245 | QPCT |
| 246 | RBP3 |
| 247 | REN |

| | |
|---|---|
| 248 | RNPEP |
| 249 | SCPEP1 |
| 250 | SCRN2 |
| 251 | STC2 |
| 252 | TG |
| 253 | TGM1 |
| 254 | TGM2 |
| 255 | TGM3 |
| 256 | TGM4 |
| 257 | TINAG |
| 258 | TINAGL1 |
| 259 | TLL1 |
| 260 | TLL2 |
| 261 | TNFAIP3 |
| 262 | TPSAB1 |
| 263 | TPSD1 |
| 264 | UCHL1 |
| 265 | UCHL3 |
| 266 | UFSP1 |
| 267 | UQCRC2 |
| 268 | USPL1 |
| 269 | XPNPEP1 |
| 270 | XPNPEP2 |
| 271 | XPNPEP3 |

### F. HNSCC overexpressed

| No. | Gene Name | Cancer expr. | Healthy expr. | Ratio | Cleavage motifs |
|---|---|---|---|---|---|
| 1 | MMP13 | 4868.4 | 60.8 | 80.116 | 103 |
| 2 | ADAMTS20 | 17.3 | 0.3 | 67.119 | 0 |
| 3 | MMP11 | 4341 | 121.3 | 35.789 | 4 |
| 4 | TMPRSS15 | 6.7 | 0.4 | 18.464 | 4 |
| 5 | MMP9 | 4800.1 | 313.6 | 15.305 | 326 |
| 6 | CTRB2 | 2.1 | 0.1 | 14.695 | 0 |
| 7 | MMP12 | 1509.2 | 108.2 | 13.948 | 218 |
| 8 | ADAM12 | 1372.6 | 118.9 | 11.544 | 0 |
| 9 | ADAMTS12 | 241.9 | 25.6 | 9.437 | 0 |
| 10 | MMP1 | 16049 | 1986.3 | 8.08 | 65 |
| 11 | STC2 | 689.6 | 85.7 | 8.05 | 0 |
| 12 | MMP10 | 4561.1 | 615.9 | 7.406 | 18 |
| 13 | MMP3 | 2866.3 | 392.3 | 7.306 | 2451 |
| 14 | PCSK9 | 1174.9 | 165.8 | 7.086 | 3 |
| 15 | PROC | 72.3 | 10.6 | 6.82 | 11 |
| 16 | GPR144 | 3.9 | 0.6 | 6.774 | 0 |
| 17 | HTRA4 | 7.5 | 1.1 | 6.684 | 0 |
| 18 | MMP8 | 12.9 | 1.9 | 6.618 | 0 |
| 19 | TINAG | 1.1 | 0.2 | 6.12 | 0 |
| 20 | F2 | 4.1 | 0.7 | 5.898 | 0 |
| 21 | ADAMTS6 | 48.8 | 8.3 | 5.858 | 0 |
| 22 | CPA2 | 3.3 | 0.6 | 5.731 | 0 |
| 23 | EMR1 | 65.4 | 11.6 | 5.656 | 0 |
| 24 | FAP | 756.2 | 135.9 | 5.563 | 4 |
| 25 | PLAU | 7462.6 | 1344.4 | 5.551 | 10 |
| 26 | CTSE | 17.5 | 3.3 | 5.249 | 0 |
| 27 | CPA6 | 37.7 | 7.5 | 5.024 | 0 |

G. Protease substr. seq.  (186)

| No | Protease (Gene / Protein name) | Cleavage motif | Peptide sequence | Peptide ID | Protease/Peptide Source |
|---|---|---|---|---|---|
| 1 | ADAM10 / ADAM10 | SANAVVSQ | SANAVVSQNLVPMVATV | ADAM10-1 | Literature |
| | | HANHMAAQ | HANHMAAQNLVPMVATV | ADAM10-2 | Literature |
| | | HPSHVLSS | HPSHVLSSNLVPMVATV | ADAM10-3 | Literature |
| | | SAATLIIVN | SAATLIIVNLVPMVATV | ADAM10-4 | Literature |
| 2 | ADAM28 / ADAM28 | KPAKFFRL | KPAKFFRLNLVPMVATV{NH2} | ADAM28 | Prior screens |
| 3 | C1A / C1s | YLGRSYKV | YLGRSYKVNLVPMVATV | C1s | Prior screens |
| 4 | CTSB / Cathepsin B | FRNLVPMVATV | FRNLVPMVATV | Cathepsin B-1 | Prior screens |
| | | PMKRLTLG | PMKRLTLGNLVPMVATV | Cathepsin B-10 | HNSCC screen |
| | | HLVEALYL | HLVEALYLNLVPMVATV | Cathepsin B-11 | HNSCC screen |
| | | EVDLLIGS | EVDLLIGSNLVPMVATV | Cathepsin B-12 | HNSCC screen |
| | | PRFKIIGG | PRFKIIGGNLVPMVATV | Cathepsin B-13 | HNSCC screen |
| | | FRFRFR | FRFRFRNLVPMVATV | Cathepsin B-2 | Prior screens |
| | | ASVRA | ASVRANLVPMVATV | Cathepsin B-3 | Prior screens |
| | | VRA | VRANLVPMVATV | Cathepsin B-4 | Prior screens |
| | | VR | VRNLVPMVATV | Cathepsin B-5 | Prior screens |
| | | FRL | FRLNLVPMVATVFRLNLVPMVATVFRLNLVPMVATV | Cathepsin B-6 | Prior screens |
| | | FK | FKNLVPMVATV | Cathepsin B-7 | Prior screens |
| | | GGGGF | GGGGFNLVPMVATV | Cathepsin B-8 | HNSCC screen |
| | | QVVAG | QVVAGNLVPMVATV | Cathepsin B-9 | HNSCC screen |
| 5 | CTSD / Cathepsin D | PRSFFRLGK | PRSFFRLGKNLVPMVATV | Cathepsin D-1 | Prior screens |
| | | GSTFF | GSTFFNLVPMVATV | Cathepsin D-2 | HNSCC screen |
| | | EVLLSWAV | EVLLSWAVNLVPMVATV | Cathepsin G-1 | HNSCC screen |
| | | PVSLSYRC | PVSLSYRCNLVPMVATV | Cathepsin G-2 | HNSCC screen |
| 6 | ECE1 / Endothelin-converting enzyme 1 | TPEHVVPY | TPEHVVPYNLVPMVATV | ECE-1 | Prior screens |
| 7 | FAP / Fibroblast activation protein, alpha | PNQEQ | PNQEQNLVPMVATV | FAP-1 | HNSCC screen |
| | | AMEPLGRQ | AMEPLGRQNLVPMVATV | FAP-2 | HNSCC screen |
| | | TSGPNQEQ | TSGPNQEQNLVPMVATV | FAP-3 | HNSCC screen |
| 8 | KLK2 / Kallikrein Related Peptidase 2 | GKAFRRL | GKAFRRLNLVPMVATV | KLK2 | Prior screens |
| 9 | LGMN / Legumain | AANL | AANLNLVPMVATV | Legumain | Prior screens |
| 10 | MASP2 / MBL associated serine protease 2 | SLGRKIQI | SLGRKIQINLVPMVATV | MASP2 | Prior screens |
| 11 | MMP1 / MMP1 | AIPVSLR | AIPVSLRNLVPMVATV | MMP1-1 | Prior screens |
| | | RVAEMRGE | RVAEMRGENLVPMVATV | MMP1-2 | HNSCC screen |
| | | GPQGLLGA | GPQGLLGANLVPMVATV | MMP1-3 | HNSCC screen |
| | | GPQGLAGQ | GPQGLAGQNLVPMVATV | MMP1-4 | HNSCC screen |
| | | GPQGIAGI | GPQGIAGINLVPMVATV | MMP1-5 | HNSCC screen |
| | | LYVGSKTK | LYVGSKTKNLVPMVATV | MMP1-6 | HNSCC screen |
| | | TAVAQKTV | TAVAQKTVNLVPMVATV | MMP1-7 | HNSCC screen |
| | | QPDAINAP | QPDAINAPNLVPMVATV | MMP1-8 | HNSCC screen |
| | | QPVGINTS | QPVGINTSNLVPMVATV | MMP1-9 | HNSCC screen |
| | | GPLGIAGI | GPLGIAGINLVPMVATV | MMP1-10 | HNSCC screen |

| | | | | | |
|---|---|---|---|---|---|
| | | LRAYLLPA | LRAYLLPANLVPMVATV | MMP1-11 | HNSCC screen |
| | | VSRLRAYL | VSRLRAYLNLVPMVATV | MMP1-12 | HNSCC screen |
| | | DAETLKVM | DAETLKVMNLVPMVATV | MMP1-13 | HNSCC screen |
| | | TLKAMRTP | TLKAMRTPNLVPMVATV | MMP1-14 | HNSCC screen |
| | | AYSDMREA | AYSDMREANLVPMVATV | MMP1-15 | HNSCC screen |
| 12 | **MMP2 / MMP2** | GPLGVRGK | GPLGVRGKNLVPMVATV | MMP2-1 | Prior screens |
| | | AIPVSLR | AIPVSLRNLVPMVATV | MMP2-2 | Prior screens |
| | | HPVGLLAR | HPVGLLARNLVPMVATV | MMP2-3 | Prior screens |
| | | PLGLAG | PLGLAGNLVPMVATV | MMP2-4 | HNSCC screen |
| | | PLGLWA | PLGLWANLVPMVATV | MMP2-5 | HNSCC screen |
| | | KGPLGVRG | KGPLGVRGNLVPMVATV | MMP2-6 | HNSCC screen |
| 12, 15 | **MMP2, 9 / MMP2, 9** | PLGLYL | PLGLYLNLVPMVATV | MMP2/9-1 | Prior screens |
| | | GPLGIAGQ | GPLGIAGQNLVPMVATV | MMP2/9-2 | Prior screens |
| | | PVGLIG | PVGLIGNLVPMVATV | MMP2/9-3 | Prior screens |
| | | GPLGLWAQ | GPLGLWAQNLVPMVATV | MMP2/9-4 | Prior screens |
| 13 | **MMP3 / MMP3** | IPENFFGV | IPENFFGVNLVPMVATV | MMP3-1 | Prior screens |
| | | VASSSTAV | VASSSTAVNLVPMVATV | MMP3-2 | HNSCC screen |
| | | STAVIVSA | STAVIVSANLVPMVATV | MMP3-3 | HNSCC screen |
| | | EAIPMSIP | EAIPMSIPNLVPMVATV | MMP3-4 | HNSCC screen |
| | | GPKGARGD | GPKGARGDNLVPMVATV | MMP3-5 | HNSCC screen |
| | | GPNGFNGD | GPNGFNGDNLVPMVATV | MMP3-6 | HNSCC screen |
| | | GPPGLTGP | GPPGLTGPNLVPMVATV | MMP3-7 | HNSCC screen |
| | | GPRGRSGE | GPRGRSGENLVPMVATV | MMP3-8 | HNSCC screen |
| | | DMSAFAGL | DMSAFAGLNLVPMVATV | MMP3-9 | HNSCC screen |
| | | PPAHSHRD | PPAHSHRDNLVPMVATV | MMP3-10 | HNSCC screen |
| | | YPRGIHTL | YPRGIHTLNLVPMVATV | MMP3-11 | HNSCC screen |
| | | SLGLFHSA | SLGLFHSANLVPMVATV | MMP3-12 | HNSCC screen |
| | | LTYRIVNY | LTYRIVNYNLVPMVATV | MMP3-13 | HNSCC screen |
| | | IRGNEVRA | IRGNEVRANLVPMVATV | MMP3-14 | HNSCC screen |
| | | LPKDAVDS | LPKDAVDSNLVPMVATV | MMP3-15 | HNSCC screen |
| | | DPALSFDA | DPALSFDANLVPMVATV | MMP3-16 | HNSCC screen |
| | | GIPKWRKT | GIPKWRKTNLVPMVATV | MMP3-17 | HNSCC screen |
| 14 | **MMP7 / MMP7** | GPLGLARK | GPLGLARKNLVPMVATV | MMP7-1 | Prior screens |
| | | RPLALWRS | RPLALWRSNLVPMVATV | MMP7-2 | Prior screens |
| | | AVSRLRAY | AVSRLRAYNLVPMVATV | MMP7-3 | Prior screens |
| | | VVYGLRSK | VVYGLRSKNLVPMVATV | MMP7-4 | Prior screens |
| | | LRELHLDN | LRELHLDNNLVPMVATV | MMP7-5 | HNSCC screen |
| | | MLEDEASG | MLEDEASGNLVPMVATV | MMP7-6 | HNSCC screen |
| 15 | **MMP9 / MMP9** | KPVSLSYR | KPVSLSYRNLVPMVATV | MMP9-1 | Prior screens |
| | | GPQGIAGQR | GPQGIAGQRNLVPMVATV | MMP9-2 | Prior screens |
| | | AVRWLLTA | AVRWLLTANLVPMVATV | MMP9-3 | Prior screens |
| | | PRALM | PRALMNLVPMVATV | MMP9-4 | Prior screens |
| | | PLGMTS | PLGMTSNLVPMVATV | MMP9-5 | HNSCC screen |

| | | | | | |
|---|---|---|---|---|---|
| | | PRGMAS | PRGMASNLVPMVATV | MMP9-6 | HNSCC screen |
| | | GPQGARGQ | GPQGARGQNLVPMVATV | MMP9-7 | HNSCC screen |
| | | AAALGNVAP | AAALGNVAPNLVPMVATV | MMP9-8 | HNSCC screen |
| | | PQGLAG | PQGLAGNLVPMVATV | MMP9-9 | HNSCC screen |
| | | LLESDLFP | LLESDLFPNLVPMVATV | MMP9-10 | HNSCC screen |
| | | KPAVTAAP | KPAVTAAPNLVPMVATV | MMP9-11 | HNSCC screen |
| | | NPLENSGF | NPLENSGFNLVPMVATV | MMP9-12 | HNSCC screen |
| | | GPGGVVGP | GPGGVVGPNLVPMVATV | MMP9-13 | HNSCC screen |
| | | GPRGLPGE | GPRGLPGENLVPMVATV | MMP9-14 | HNSCC screen |
| | | GPEGAQGP | GPEGAQGPNLVPMVATV | MMP9-15 | HNSCC screen |
| | | GPSGKDGA | GPSGKDGANLVPMVATV | MMP9-16 | HNSCC screen |
| | | GLRGLPGK | GLRGLPGKNLVPMVATV | MMP9-17 | HNSCC screen |
| | | AIPGGVPG | AIPGGVPGNLVPMVATV | MMP9-18 | HNSCC screen |
| | | LPVNVTDY | LPVNVTDYNLVPMVATV | MMP9-19 | HNSCC screen |
| | | SVYTVKDD | SVYTVKDDNLVPMVATV | MMP9-20 | HNSCC screen |
| | | GPAGANGE | GPAGANGENLVPMVATV | MMP9-21 | HNSCC screen |
| | | GYPIKAPK | GYPIKAPKNLVPMVATV | MMP9-22 | HNSCC screen |
| | | LATASTMD | LATASTMDNLVPMVATV | MMP9-23 | HNSCC screen |
| | | LVYMVTGN | LVYMVTGNNLVPMVATV | MMP9-24 | HNSCC screen |
| | | GVFYPGAG | GVFYPGAGNLVPMVATV | MMP9-25 | HNSCC screen |
| 16 | **MMP10 / MMP10** | PPAQAVLQ | PPAQAVLQNLVPMVATV | MMP10-1 | HNSCC screen |
| | | GPHLLVEA | GPHLLVEANLVPMVATV | MMP10-2 | HNSCC screen |
| | | RGLSLSRF | RGLSLSRFNLVPMVATV | MMP10-3 | HNSCC screen |
| | | SPMARRGG | SPMARRGGNLVPMVATV | MMP10-4 | HNSCC screen |
| | | LPQKSQHG | LPQKSQHGNLVPMVATV | MMP10-5 | HNSCC screen |
| | | GSLPQKSQ | GSLPQKSQNLVPMVATV | MMP10-6 | HNSCC screen |
| | | DPKNNWQG | DPKNNWQGNLVPMVATV | MMP10-7 | HNSCC screen |
| | | RPSQRSKY | RPSQRSKYNLVPMVATV | MMP10-8 | HNSCC screen |
| | | RSKYLATA | RSKYLATANLVPMVATV | MMP10-9 | HNSCC screen |
| | | LPRHRDTG | LPRHRDTGNLVPMVATV | MMP10-10 | HNSCC screen |
| 17 | **MMP11 / MMP11** | AAATSIAM | AAATSIAMNLVPMVATV | MMP11-1 | Prior screens |
| | | AAGAMFLE | AAGAMFLENLVPMVATV | MMP11-2 | Prior screens |
| | | EAAAATSI | EAAAATSINLVPMVATV | MMP11-3 | HNSCC screen |
| | | PAAALILA | PAAALILANLVPMVATV | MMP11-4 | HNSCC screen |
| | | AAGAFLME | AAGAFLMENLVPMVATV | MMP11-5 | HNSCC screen |
| | | KALHVTNI | KALHVTNINLVPMVATV | MMP11-6 | HNSCC screen |
| 18 | **MMP12 / MMP12** | GPKGARGD | GPKGARGDNLVPMVATV | MMP12-1 | HNSCC screen |
| | | GPPGATGF | GPPGATGFNLVPMVATV | MMP12-2 | HNSCC screen |
| | | GIGGIAGV | GIGGIAGVNLVPMVATV | MMP12-3 | HNSCC screen |
| | | NPLKSSGI | NPLKSSGINLVPMVATV | MMP12-4 | HNSCC screen |
| | | GLGALGGV | GLGALGGVNLVPMVATV | MMP12-5 | HNSCC screen |
| | | PGAGLGAL | PGAGLGALNLVPMVATV | MMP12-6 | HNSCC screen |
| | | GQFPLGGV | GQFPLGGVNLVPMVATV | MMP12-7 | HNSCC screen |

| | | | | | |
|---|---|---|---|---|---|
| | | PGARFPGV | PGARFPGVNLVPMVATV | MMP12-8 | HNSCC screen |
| | | GLSPIFPG | GLSPIFPGNLVPMVATV | MMP12-9 | HNSCC screen |
| | | RLEHLYLN | RLEHLYLNNLVPMVATV | MMP12-10 | HNSCC screen |
| | | KAAQFGLV | KAAQFGLVNLVPMVATV | MMP12-11 | HNSCC screen |
| | | GALVPGGV | GALVPGGVNLVPMVATV | MMP12-12 | HNSCC screen |
| | | LPYGYGPG | LPYGYGPGNLVPMVATV | MMP12-13 | HNSCC screen |
| | | GPVEVFIT | GPVEVFITNLVPMVATV | MMP12-14 | HNSCC screen |
| | | LVFLYMEK | LVFLYMEKNLVPMVATV | MMP12-15 | HNSCC screen |
| 19 | **MMP13 / MMP13** | TVKPVFEV | TVKPVFEVNLVPMVATV | MMP13-1 | HNSCC screen |
| | | TVKPIFEV | TVKPIFEVNLVPMVATV | MMP13-2 | HNSCC screen |
| | | LLSALVET | LLSALVETNLVPMVATV | MMP13-3 | HNSCC screen |
| | | VPKGVFSG | VPKGVFSGNLVPMVATV | MMP13-4 | HNSCC screen |
| | | DISELRKD | DISELRKDNLVPMVATV | MMP13-5 | HNSCC screen |
| | | SDLGLKSV | SDLGLKSVNLVPMVATV | MMP13-6 | HNSCC screen |
| | | SEAKLTGI | SEAKLTGINLVPMVATV | MMP13-7 | HNSCC screen |
| | | QAIELEDL | QAIELEDLNLVPMVATV | MMP13-8 | HNSCC screen |
| | | FGVKRAYY | FGVKRAYYNLVPMVATV | MMP13-9 | HNSCC screen |
| | | NPVPYWEV | NPVPYWEVNLVPMVATV | MMP13-10 | HNSCC screen |
| | | APVGTREV | APVGTREVNLVPMVATV | MMP13-11 | HNSCC screen |
| | | VVGGLVAL | VVGGLVALNLVPMVATV | MMP13-12 | HNSCC screen |
| | | GLAGQRGI | GLAGQRGINLVPMVATV | MMP13-13 | HNSCC screen |
| | | GPPGSNGN | GPPGSNGNNLVPMVATV | MMP13-14 | HNSCC screen |
| | | GPSGLAGP | GPSGLAGPNLVPMVATV | MMP13-15 | HNSCC screen |
| | | GAPGLRGL | GAPGLRGLNLVPMVATV | MMP13-16 | HNSCC screen |
| | | DIKDIVGP | DIKDIVGPNLVPMVATV | MMP13-17 | HNSCC screen |
| | | GAPGLRGG | GAPGLRGGNLVPMVATV | MMP13-18 | HNSCC screen |
| | | GPPGKNGE | GPPGKNGENLVPMVATV | MMP13-19 | HNSCC screen |
| | | LPARITPG | LPARITPGNLVPMVATV | MMP13-20 | HNSCC screen |
| | | TARPWRAD | TARPWRADNLVPMVATV | MMP13-21 | HNSCC screen |
| | | SLRELHLD | SLRELHLDNLVPMVATV | MMP13-22 | HNSCC screen |
| | | GPSGLLAH | GPSGLLAHNLVPMVATV | MMP13-23 | HNSCC screen |
| | | HLRYLRLD | HLRYLRLDNLVPMVATV | MMP13-24 | HNSCC screen |
| | | NPVQVEVG | NPVQVEVGNLVPMVATV | MMP13-25 | HNSCC screen |
| 20 | **MMP14 / MMP14** | PRHLR | PRHLRNLVPMVATV | MMP14-1 | Prior screens |
| | | PRSAKELR | PRSAKELRNLVPMVATV | MMP14-2 | Prior screens |
| | | GPLPLR | GPLPLRNLVPMVATV | MMP14-3 | HNSCC screen |
| 21, 22, 23, 24 | **MMP15, 16, 24, 25 / MMP15, 16, 24, 25** | PRGLRK | PRGLRKNLVPMVATV | MMP15/16/24/25 | Prior screens |
| | | PRWLRS | PRWLRSNLVPMVATV | MMP15/16/24/25-1 | HNSCC screen |
| | | PRGLRP | PRGLRPNLVPMVATV | MMP15/16/24/25-2 | HNSCC screen |
| 25 | **PCSK9 / Proprotein Convertase Subtilisin/Kexin Type 9** | VVVLKEET | VVVLKEETNLVPMVATV | PCSK9-1 | HNSCC screen |
| | | VFAQSIPW | VFAQSIPWNLVPMVATV | PCSK9-2 | HNSCC screen |
| 26 | **PLG / Plasminogen (–> Plasmin)** | GGR | GGRNLVPMVATV | Plasmin | Prior screens |

| | | | | | |
|---|---|---|---|---|---|
| 27 | **PLAU / Plasminogen activator, urokinase (uPA)** | SGRSANAK | SGRSANAKNLVPMVATV | PLAU-1 | Prior screens |
| | | NSGRAVTY | NSGRAVTYNLVPMVATV | PLAU-2 | Prior screens |
| | | SRRRVNSL | SRRRVNSLNLVPMVATV | PLAU-3 | Prior screens |
| | | TYSRSRYL | TYSRSRYLNLVPMVATV | PLAU-4 | HNSCC screen |
| | | RGSVILTV | RGSVILTVNLVPMVATV | PLAU-5 | HNSCC screen |
| | | PSSRRRVN | PSSRRRVNNLVPMVATV | PLAU-6 | HNSCC screen |
| | | SSSRGPYH | SSSRGPYHNLVPMVATV | PLAU-7 | HNSCC screen |
| | | PGARGRAF | PGARGRAFNLVPMVATV | PLAU-8 | HNSCC screen |
| | | VSNKYFSN | VSNKYFSNNLVPMVATV | PLAU-9 | HNSCC screen |
| | | KQLRVVNG | KQLRVVNGNLVPMVATV | PLAU-10 | HNSCC screen |
| 28 | **PROC / protein C** | PQLRMKNN | PQLRMKNNNLVPMVATV | PROC-1 | HNSCC screen |
| | | RLKKSQFL | RLKKSQFLNLVPMVATV | PROC-2 | HNSCC screen |
| | | LDQRGVQR | LDQRGVQRNLVPMVATV | PROC-3 | HNSCC screen |
| | | VDQRGNQI | VDQRGNQINLVPMVATV | PROC-4 | HNSCC screen |
| | | IEPRSFSQ | IEPRSFSQNLVPMVATV | PROC-5 | HNSCC screen |
| 29 | **TMPRSS15 / Transmembrane serine protease 15** | GESKSLGP | GESKSLGPNLVPMVATV | TMPRSS15-1 | HNSCC screen |
| | | DDDKIVGG | DDDKIVGGNLVPMVATV | TMPRSS15-2 | HNSCC screen |
| | | DDDK | DDDKNLVPMVATV | TMPRSS15-3 | HNSCC screen |
| | **Controls** | | NLVPMVATV | Free peptide | |
| | | GGGGG | GGGGGNLVPMVATV | Uncleavable linker | |
| | | GGGS | GGGSNLVPMVATV | Uncleavable linker | |
| | | | NLVPMVATV | Untreated | |
| | | | Uncleavable (MPA113) | No linker | |

#### A. RNP sequences

| ID | gRNA-1 | gRNA-2 | gRNA-3 | gRNA-4 | gRNA-5 |
|---|---|---|---|---|---|
| **Design Tool** | IDT | IDT | IDT | Synthego | Synthego |
| **Guide RNA sequence** | CCGACCTTTG GTATC | TGGTAGATGG CTGCGAACCA | GATGCTATTA GCCTAGGCCT | GCAAACUGU GGCUGUCAG AA | AACUGUGGC UGUCAGAAC GG |
| **Strand** | Sense(-) | Sense(-) | Antisense(+) | Sense(-) | Sense(-) |
| **PAM** | TGG | GGG | GGG | CGG | AGG |
| **Locus** | Exon 4 | Exon 5 | Exon 4 | Exon 4 | Exon 4 |
| **DSB Location** | 20838566 | 20839444 | 20838646 | 20837794 | 20837797 |
| **crRNA seq** | /AltR1/rCrC rGrArC rCrUrU rUrGrG rUrArU rCrArG rUrGrU rGrUrU rUrUrA rGrArG rCrUrA rUrGrC rU/AltR2/ | /AltR1/rUrG rGrUrA rGrArU rGrGrC rUrGrC rGrArA rCrCrA rGrUrU rUrUrA rGrArG rCrUrA rUrGrC rU/AltR2/ | /AltR1/rGrA rUrGrC rUrArU rUrArG rCrCrU rArGrG rCrCrU rGrUrU rUrUrA rGrArG rCrUrA rUrGrC rU/AltR2/ | /AltR1/rGrC rArArA rCrUrG rUrGrG rCrUrG rUrCrA rGrArA rGrUrU rUrUrA rGrArG rCrUrA rUrGrC rU/AltR2/ | /AltR1/rArA rCrUrG rUrGrG rCrUrG rUrCrA rGrArA rCrGrG rGrUrU rUrUrA rGrArG rCrUrA rUrGrC rU/AltR2/ |

**Anti-human antibodies**

| Antigen | Flurochrome | Clone | Cat# | Vendor | Dilution |
|---|---|---|---|---|---|
| CD3 | APC-Cy7 | OKT3 | 317342 | Biolegend | 1:200 |
| CD8a | BV785 | SKI | 344740 | Biolegend | 1:200 |
| CD11b | FITC | ICRF44 | 301330 | Biolegend | 1:200 |
| CD19 | FITC | HIB19 | 302206 | Biolegend | 1:200 |
| CD33 | FITC | P67.6 | 366620 | Biolegend | 1:200 |
| GzmB | AF700 | QA16A02 | 372222 | Biolegend | 1:50 |
| GzmB | AF700 | QA16A02 | 372214 | Biolegend | 1:50 |
| TCF-1 | AF488 | C63D9 | 6444S | Cell Signaling | 1:25 |
| TCF-1 | PE | C63D9 | 14456 | Cell Signaling | 1:25 |
| TCF-1 | PE | C63D9 | 6709S | Cell Signaling | 1:25 |
| Human FcX | N/A | N/A | 422302 | Biolegend | 1:100 |

**Anti-mouse antibodies**

| Antigen | Flurochrome | Clone | Cat# | Vendor | Dilution |
|---|---|---|---|---|---|
| CD3 | APC/Cy7 | 17A2 | 100222 | Biolegend | 1:200 |
| CD8a | APC | 53-6.7 | 100712 | Biolegend | 1:200 |
| CD8a | PE | 53-6.7 | 12-0081-83 | Biolegend | 1:200 |
| CD19 | FITC | QA17A27 | 159808 | Biolegend | 1:200 |
| CD14 | FITC | Sa14-2 | 123307 | Biolegend | 1:200 |
| GzmB | PE/Cy7 | QA16A02 | 372214 | Biolegend | 1:50 |
| TCF-1 | PE | 7F11A10 | 655207 | Biolegend | 1:50 |
| PD-1 | BV711 | 29F.1A12 | 135231 | Biolegend | 1:100 |
| PD-1 | BV421 | 29F.1A12 | 135221 | Biolegend | 1:100 |
| EpCAM | PE | G8.8 | 12-5791-81 | eBioscience | 1:100 |
| H2Kb | AF647 | AF6-88.5 | 116512 | Biolegend | 1:100 |
| uPA | (Biotin) | OTI5H4 | TA805243AM | Origene | 1:50 |

**Streptavidin**

| Target | Flurochrome | Clone | Cat# | Vendor | |
|---|---|---|---|---|---|
| Biotin | BUV395 | N/A | 564176 | BD Bioscience | 1:200 |
| Biotin | BV421 | N/A | 563259 | BD Bioscience | 1:200 |
| Biotin | BV711 | N/A | 563262 | BD Bioscience | 1:200 |
| Biotin | BV605 | N/A | 563260 | BD Bioscience | 1:200 |
| Biotin | BUV737 | N/A | 612775 | BD Bioscience | 1:200 |
| Biotin | APC | N/A | 554067 | BD Bioscience | 1:200 |
| Biotin | PE | N/A | 554061 | BD Bioscience | 1:200 |

**Other reagents**

| | | | | | |
|---|---|---|---|---|---|
| LiveDead | Aqua | N/A | L34957 | Thermo Fisher | 1:200 |
